# Profiling Myelodysplastic Syndromes by Mass Cytometry Demonstrates Abnormal Progenitor Cell Phenotype and Differentiation

**DOI:** 10.1101/604397

**Authors:** Gregory K. Behbehani, Rachel Finck, Nikolay Samusik, Kunju Sridhar, Wendy J. Fantl, Peter L. Greenberg, Garry P. Nolan

## Abstract

**Purpose:** We sought to enhance the cytometric analysis of MDS by performing a pilot study of a single cell mass cytometry (MCM) assay to more comprehensively analyze patterns of surface marker expression in patients with MDS.

**Experimental Design:** Twenty-three MDS and five healthy donor bone marrow samples were studied using a 34-parameter mass cytometry panel utilizing barcoding and internal reference standards. The resulting data were analyzed by both traditional gating and high-dimensional clustering.

**Results:** This high-dimensional assay provided three major benefits relative to traditional cytometry approaches: First, MCM enabled detection of aberrant surface maker at high resolution, detecting aberrancies in 27/31 surface markers, encompassing almost every previously reported MDS surface marker aberrancy. Additionally, three previously unrecognized aberrancies in MDS were detected in multiple samples at least one developmental stage: increased CD321 and CD99; and decreased CD47. Second, analysis of the stem and progenitor cell compartment (HSPCs), demonstrated aberrant expression in 21 of the 23 MDS samples, which were not detected in three samples from patients with idiopathic cytopenia of undetermined significance (ICUS). These immunophenotypically abnormal HSPCs were also the single most significant distinguishing feature between clinical risk groups. Third, unsupervised clustering of high-parameter MCM data allowed identification of abnormal differentiation patterns associated with immunophenotypically aberrant myeloid cells similar to myeloid derived suppressor cells.

**Conclusions:** These results demonstrate that high-parameter cytometry methods that enable simultaneous analysis of all bone marrow cell types could enhance the diagnostic utility of immunophenotypic analysis in MDS.

**Statement of Significance:**

- High-dimensional mass cytometry enables high-resolution characterization of abnormal maker expression and myeloid development in MDS.
- This technology could enhance MDS diagnosis and therapeutic monitoring and merits further research.

**Statement Translational Relevance:** In spite of several studies suggesting the utility of flow cytometry in the diagnosis of myelodysoplastic syndrome (MDS), this technique has not been widely adopted. We sought to enhance the utility of cytometry in MDS by performing the first high-dimensional mass cytometry characterization of a cohort of MDS patients. High-dimensional mass cytometry allowed all bone marrow cell populations to be simultaneously analyzed enabling high-resolution characterization of abnormal maker expression and myeloid development in MDS. This approach could identify almost all previously identified aberrant surface marker expression patterns in MDS while simultaneously enabling analysis by unsupervised clustering. Additionally, this mass cytometry analysis approach enabled the modeling of abnormal differentiation in MDS. This technology could enhance MDS diagnosis and therapeutic monitoring and merits further research.

## Introduction

The myelodysplastic syndromes (MDS) are characterized by ineffective hematopoiesis, dysplasia and peripheral blood cytopenias ^1,2^. The clinical course is heterogeneous with the disease evolving rapidly to acute myeloid leukemia (AML) in some patients, whereas in others symptoms are mild and patient survival is prolonged. Although well-characterized at the level of cytogenetic changes and gene mutations, the pathogenesis of MDS still remains incompletely understood.

Using fluorescence-based flow cytometry, several previous studies demonstrated that the type and degree of immunophenotypic aberrancy observed in MDS patient samples correlated both with diagnostic subtype and with overall survival ^3,4^. The European LeukemiaNet has recently developed standardized criteria for the classification of MDS by flow cytometry that encompasses many of these recent findings and provides a basis for the analysis of MDS by flow cytometry ^5,6^. Our investigation extends these studies by utilizing the novel technology of mass cytometry (MCM) ^7–11^, simultaneously analyzing 33 metal-labeled antibodies to characterize the immunophenotypic and functional variation of marrow cell populations. Using this method we evaluated up to 37 dimensions per single cell in bone marrow samples from MDS patients, patients with ICUS ^12^, and normal donors.

Mass cytometry is a recent innovation that enables the creation of highly multi-parametric data by conjugating antibodies against cell surface and intracellular markers to atoms of heavy metal rather than fluorophores. The binding of these antibodies to cells can then be detected and quantitated by vaporizing and ionizing the cell and measuring the amount of each heavy metal atom bound (along with its conjugated antibody) to each cell by time-of-flight mass spectrometry (ICP-MS). The much higher resolution of ICP-MS for detecting single Dalton differences in atomic mass enables up to 50 measurement channels per cell and could theoretically allow up to 120 channels. The data generated in this study allowed for the creation of high-dimensional models of aberrant hematopoietic development in MDS. This approach was facilitated by our recent development of methodologies for the assessment clinical samples by mass cytometry with barcoding techniques that permitted multiple samples to be stained and analyzed with high precision ^13,14^. The results of these analyses suggest that simultaneous analysis of all cell populations in the bone marrow of patients with MDS can yield additional diagnostic and prognostic insights and could allow for a more objective phenotypic classification of MDS.

## Methods

### Antibodies

Antibodies, isotope conjugates, manufacturers, and concentrations are listed in Supplementary Table 1. Primary antibody transition metal-conjugates were either purchased or conjugated using the MaxPAR antibody conjugation kit (Fluidigm) according to the manufacturer’s instructions. Following conjugation, antibodies were diluted to 100x working concentration in Candor PBS Antibody Stabilization solution (Candor Bioscience GmbH) and stored at 4 °C.

### Human samples

Fresh bone marrow aspirates were collected within 24 hours after aspiration into heparinized tubes and fixed using a fixation/stabilization buffer (Smart Tube, Inc.), according to the manufacturer’s instructions and frozen at −80 °C for up to 36 months prior to analysis. Samples were collected from patients at Stanford University Hospital undergoing routine bone marrow aspiration and who provided informed consent to donate a portion of the sample for tissue banking as part of a protocol approved by the Stanford University Institutional Review Board in accordance with the Declaration of Helsinki. The clinical characteristics of each patient are shown in Supplementary Table 2, with the risk-based clinical status of MDS patients indicated by IPSS category and morphologic characteristics ^15^. Five healthy control samples were obtained from Allcells, Inc. using the same protocol. Normal bone marrow samples came from 3 women and 2 men (ages: 38, 23, 23, 18, 41). Bone marrow cell samples were thawed just prior to analysis in a 4 °C water bath, and red cells were lysed using a hypotonic lysis buffer (Smart Tube).

### Antibody staining

Prior to antibody staining, mass tag cellular barcoding was performed as previously described ^13,14^ and detailed in the Supplemental Methods. Barcoding allowed groups of 20 MDS and healthy control samples to be stained with surface and intracellular antibodies in a single tube and simultaneously analyzed by mass cytometry as a single mixed sample. Barcoded cells were incubated with anti-surface marker antibodies in 2 mL of cell staining medium (CSM; 1xPBS with 0.5% bovine serum albumin and 0.02% sodium azide) for 50 minutes with continuous mixing. Cells were washed twice with CSM, and surface antibodies were fixed in place by a 15-minute incubation with 1.5% paraformaldehyde (PFA; Electron Microscopy Sciences). Subsequently, after a 15-minute incubation at −20 °C in methanol, cells were washed once with PBS and CSM prior to incubation with antibodies against intracellular signaling proteins for 50 minutes at room temperature ^16^. After completion of antibody staining, cells were washed twice with CSM and then incubated 12-36 hours in PBS with a 1:5000 dilution of the iridium intercalator pentamethylcyclopentadienyl-Ir(III)-dipyridophenazine (Fluidigm) and 1.5% paraformaldehyde. Excess intercalator was then washed away and samples were run on a CyTOF™ mass cytometer (Fluidigm) ^17^.

### Immunophenotypic aberrancy analysis

Aberrant immunophenotype analysis was performed by first gating the normal populations into developmental immunophenotypic subsets on the basis of standard surface markers (as in Supplementary Figure 1). MDS cells in each population were compared to the normal samples across 31 surface markers. Since normal healthy donor and MDS samples were all stained and analyzed in the same tube simultaneously, the same exact gates were used to identify immunophenotypic populations from all samples. The median expression level of each marker in each gated population from each patient sample was then calculated. MDS sample aberrancy was defined as an MDS sample median expression level greater than or less than the median of the similar healthy bone marrow cell population plus or minus twice the absolute variance of the healthy control samples. The summed number of markers with aberrant expression patterns was calculated for each gated immunophenotypic population from each patient sample. Statistical comparisons of median marker expression levels and population frequencies between sample groups were performed with the Mann-Whitney U test. This process is fully detailed in the Supplemental Methods.

### Data analysis

Immunophenotypic assignments were based on previous studies from our laboratory ^8,18^ and others ^19^. All gating and extraction of median expression levels was performed using Cytobank (www.cytobank.org). SPADE and viSNE analyses were performed as previously described ^20,21^, clustering markers are indicated in Supplementary Table 1. Both analyses were performed using Cytobank; SPADE analysis was performed on all analyzed events, while data files were sampled to ≤5,000 events each for viSNE analyses. K-means binning analysis of CD34^+^CD38^low^ cells was performed utilizing the same sampled cells and same surface markers employed in the viSNE analysis. Binning analysis of total cell populations was performed similarly on all cells and the same surface markers using the K-mediods algorithm with K=100. X-shift clustering was performed in accordance with the previously published methods ^22^; briefly, 10,000 cells from each sample were sampled from each MDS or controls sample and all sampled events were pooled into a single X-shift analysis to generate minimum spanning trees for each MDS and control sample. These analyses are further described in the Supplemental Methods.

## Results

### Consistent immunophenotypic measurements by mass cytometry

Thirty-one whole bone marrow aspirate samples were collected from 9 patients with higher-risk MDS (HR-MDS; IPSS = Int2/High/RAEB-T), 12 with lower-risk MDS (LR-MDS; IPSS = Low/Int1), and 3 patients with ICUS (a total of 26 samples from 24 patients). In addition, 5 BM samples from normal donors were simultaneously analyzed as internal reference comparisons (Supplemental Table 2). Two similar antibody panels incorporating 34 different antibodies were used for analysis (Supplementary Table 1). The staining panels included 31 markers of cell surface proteins and 3 markers of intracellular signaling. All samples were barcoded, such that 20 samples (MDS and healthy) could be combined into a single tube for simultaneous antibody staining and analysis. These protocols produced highly reproducible measurements of surface marker expression levels. Across replicates of the normal samples, the average coefficient of variation (CV) was 11.9%, with 29 of the 31 evaluable antibodies having CVs of less than 20% (Supplementary Table 1) ^13^. These data are consistent with prior studies ^8,21,23^ and confirmed that mass cytometry can be used with a high degree of reproducibility and accuracy for the analysis of clinical samples from patients with MDS.

### High-dimensional characterization of surface marker expression enables high-resolution aberrant immunophenotype assessment

To perform immunophenotypic analysis of the mass cytometry data, both traditional gating of cell populations based on standard surface markers and SPADE high-dimensional clustering of the data were performed using 19 of the surface markers. SPADE (spanning tree progression analysis of density normalized events) allows cells to be grouped into clusters of immunophenotypically similar cells with each cell cluster connected to its most related neighboring clusters across all clustering dimensions and represented in a minimum-spanning tree. The cell type corresponding to each cluster or group of clusters is then manually annotated by analysis of the relevant surface markers (e.g., CD3) of the cells in the cluster. The resulting SPADE analysis of normal bone marrow yielded cell groupings (nodes) that corresponded to commonly defined immunophenotypic subsets of normal hematopoietic development ^8,20^ and was consistent across all of the healthy donors; an example from one healthy donor is shown in Figure 1A and Supplementary Figure 2. As shown in Figure 1B, the fold difference in each marker could then be compared between each MDS sample and the healthy donors at each cell node. This enabled visualization of 15 common, previously identified, aberrant expression patterns (CD44, CD15, CD34, HLA-DR, CD11b, CD38, CD33, CD117, CD45, CD64, CD99, CD16, CD7, CD56, CD123)^4,5,24–28^ in samples from MDS patients from both high- and low-risk patient groups. In addition, our results demonstrated that aberrant increased CD321 and decreased CD47 expression could also be observed in MDS (Figure 1B and Supplemental Figure 3). Finally, we noted aberrant increased expression of CD99 in the erythroid progenitor populations in 4 of 11 samples from patients with lower-risk MDS, an aberrancy not previously reported in low-risk MDS (Figure 1B; and Supplemental Figure 3C). For comparison, the variance of each individual normal sample to the average of all normal samples is shown in Supplemental Figure 4. Raw median intensity levels for each population are shown in Supplemental Table 3.

**Figure 1:**
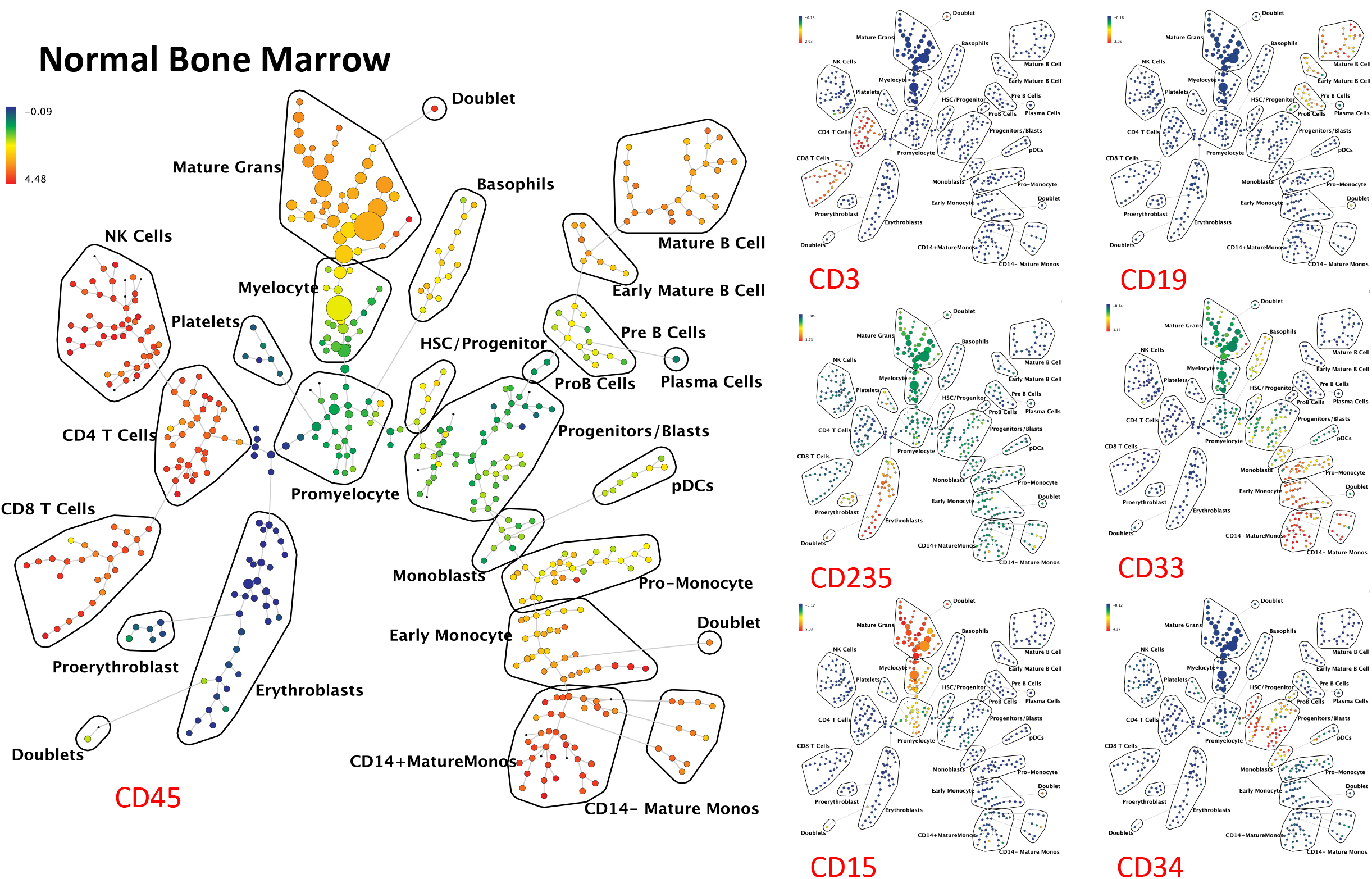
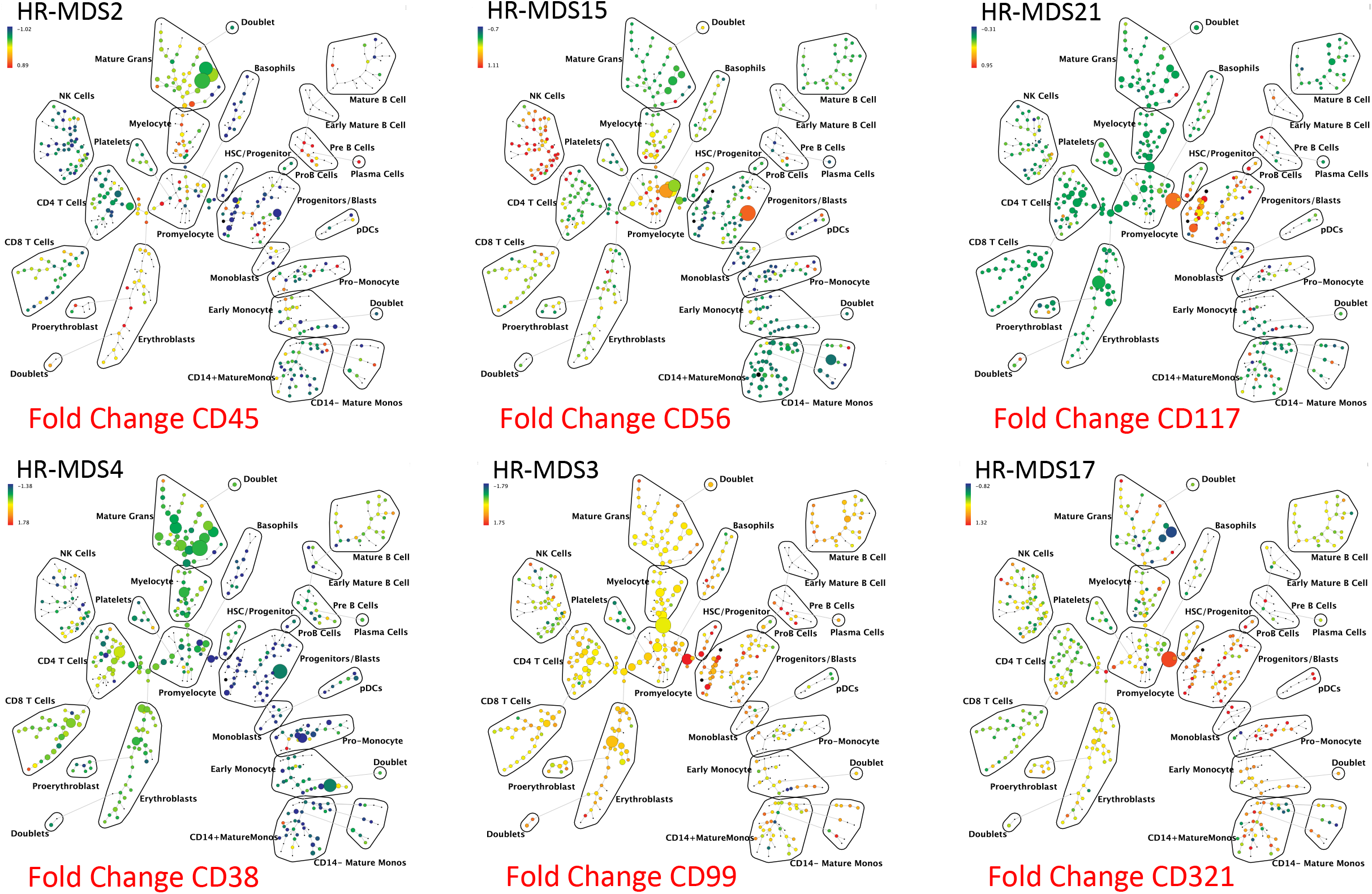
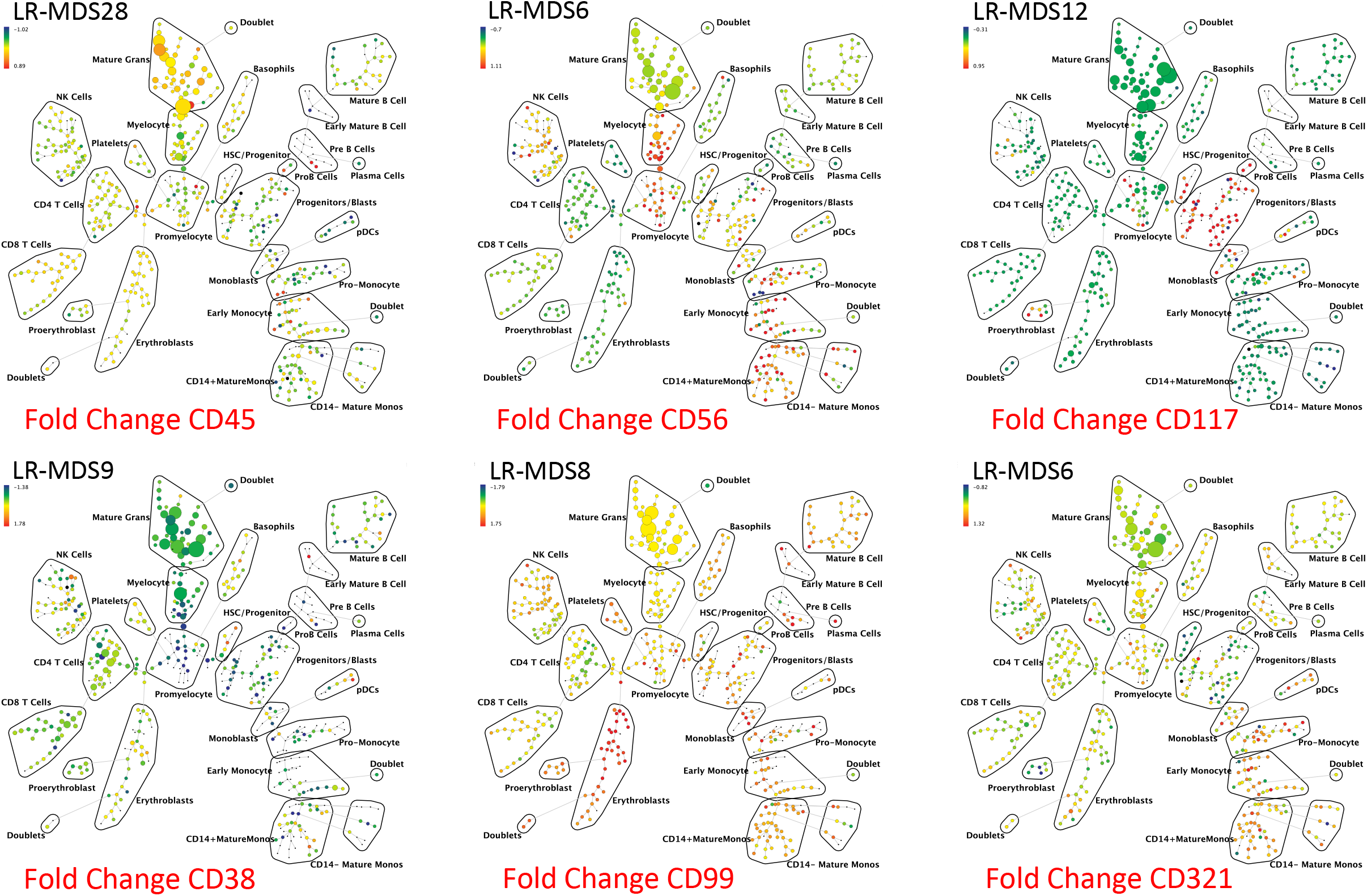
SPADE analysis enables detection of aberrant surface marker expression patterns. **(A)** SPADE plots of normal bone marrow sample #6. SPADE clustering was performed on all samples (normal and MDS) simultaneously to generate a single tree structure for all samples. All of the cell events from each sample were then mapped to the common tree structure. Each cell cluster (node) of the SPADE tree in (A) is colored for the median expression of the indicated markers from low (blue) to high (red). **(B)** SPADE tree colored for the fold change in each cell node for each of the indicated markers relative to the average of the 8 healthy donor samples for the same node. Cell nodes in (B) are colored from lowest expression relative to normal (blue) to highest expression relative to normal (red); yellow and light green colors indicate no change in expression relative the average of the control samples. These can be compared to the normal samples shown in Supplemental Figure 4. HR indicates higher-risk MDS (IPSS of Int-2 or High) LR indicates lower-risk MDS (IPSS of low or Int-1). The 8 replicate control samples came from 5 healthy donors. The size of each node is correlated to the fraction of cells mapping to the node; however, a minimum size was enforced for most nodes to allow visualization of node color. Immunophenotypic grouping of nodes was performed manually on the basis of the median marker expression level of each node, and based on analysis of the relevant biaxial plots (e.g., CD38 vs. CD34).

The simultaneous measurement of 31 markers also allowed analysis of aberrant marker expression by manual gating of each immunophenotypic population. Surface marker expression in MDS samples was evaluated compared to normal samples by first gating cells from the normal samples into immunophenotypic developmental subsets based on standard surface markers (as shown in Supplemental Figure 1). Of note, the immunophenotypic analysis demonstrated the presence of a dim-mid CD33^+^ population within the lin^-^CD34^+^CD38^low^ cells, this population was gated separately as “CD33^+^MPP” cells as this level of CD33 expression was significantly higher than the HSC population or other MPP cells. Each population from each of the normal and MDS samples was then compared across the 31 surface markers as shown in Figure 2A. Since the normal and MDS samples were stained and analyzed in the same tubes simultaneously, levels of each surface maker in each of 30 immunophenotypic cell populations of each patient sample could be reliably assessed. The summed number of markers (of the 31) with aberrant expression levels was calculated for each gated immunophenotypic population in each patient sample (a summary of aberrancies is shown in Figure 2B). Immunophenotypic aberrancies were detected at every stage of myeloid development, spanning from HSCs to mature myeloid populations in almost all MDS samples. In the majority of patients, aberrancies were also detected in most non-myeloid immunophenotypic populations (including B, T, and NK cells), suggesting that MDS has wide-ranging effects in the bone marrow. In total, changes in at least one gated HSPC or myeloid population in at least one sample were noted for 27 of the 31 surface markers in the panel (Supplemental Tables 4 and 5). As shown in Supplemental Table 4, the markers with aberrant expression varied by cell population, with marker aberrancies generally found in the cell populations that normally express each marker. The most specific changes were noted were in the hematopoietic stem and progenitor cell compartments (HSC, MPP, CD33^+^MPP, and CMP/GMP) where all MDS samples had at least one abnormality in one of these populations, while only one sample from a patient with a ICUS (ICUS18; Figure 2B) exhibited a single abnormality in the CMP/GMP population (Figure 2B and Supplemental Tables 4 and 5). These findings suggest that high parameter mass cytometry could potentially be much more sensitive than traditional flow cytometry approaches for MDS diagnosis.

**Figure 2:**
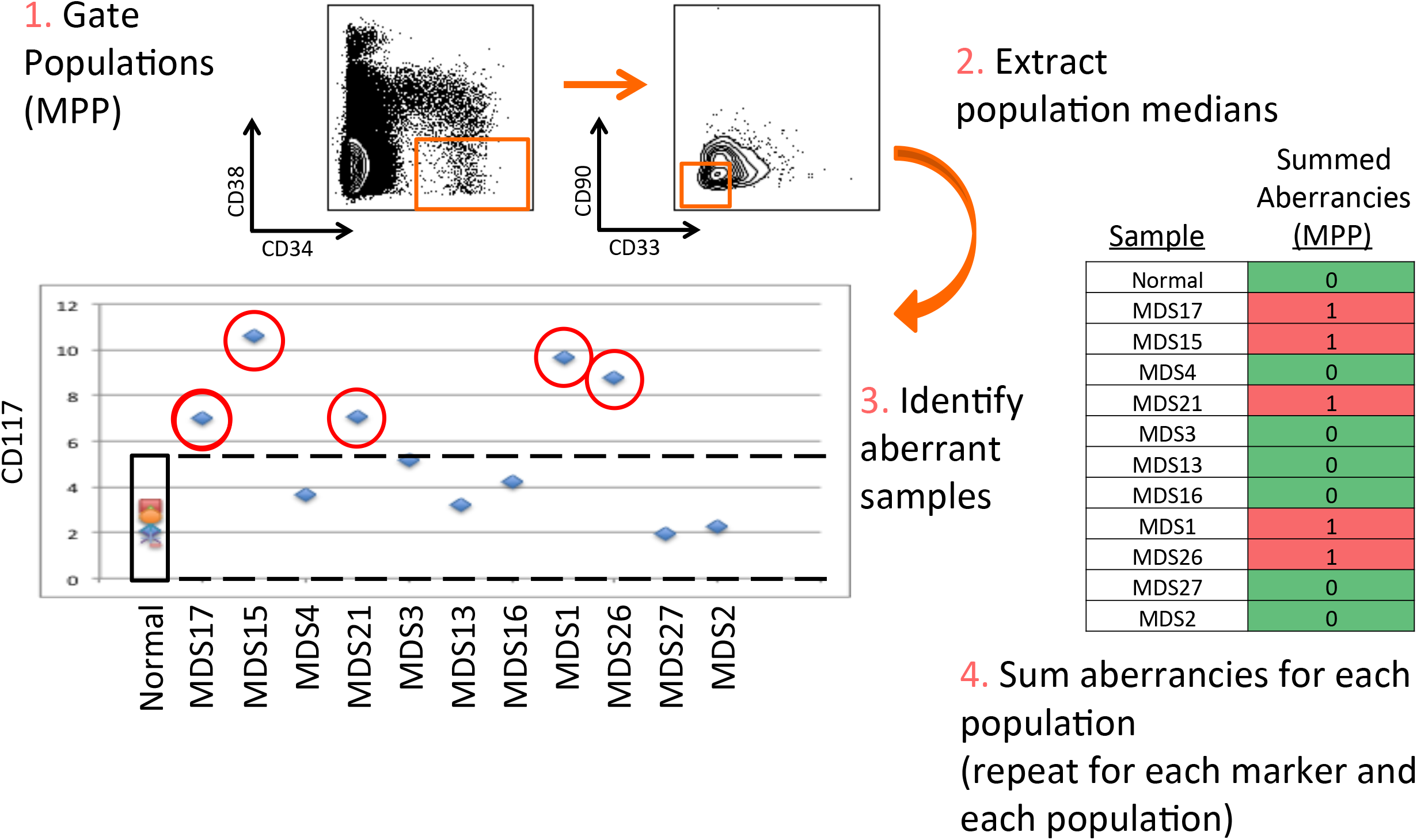
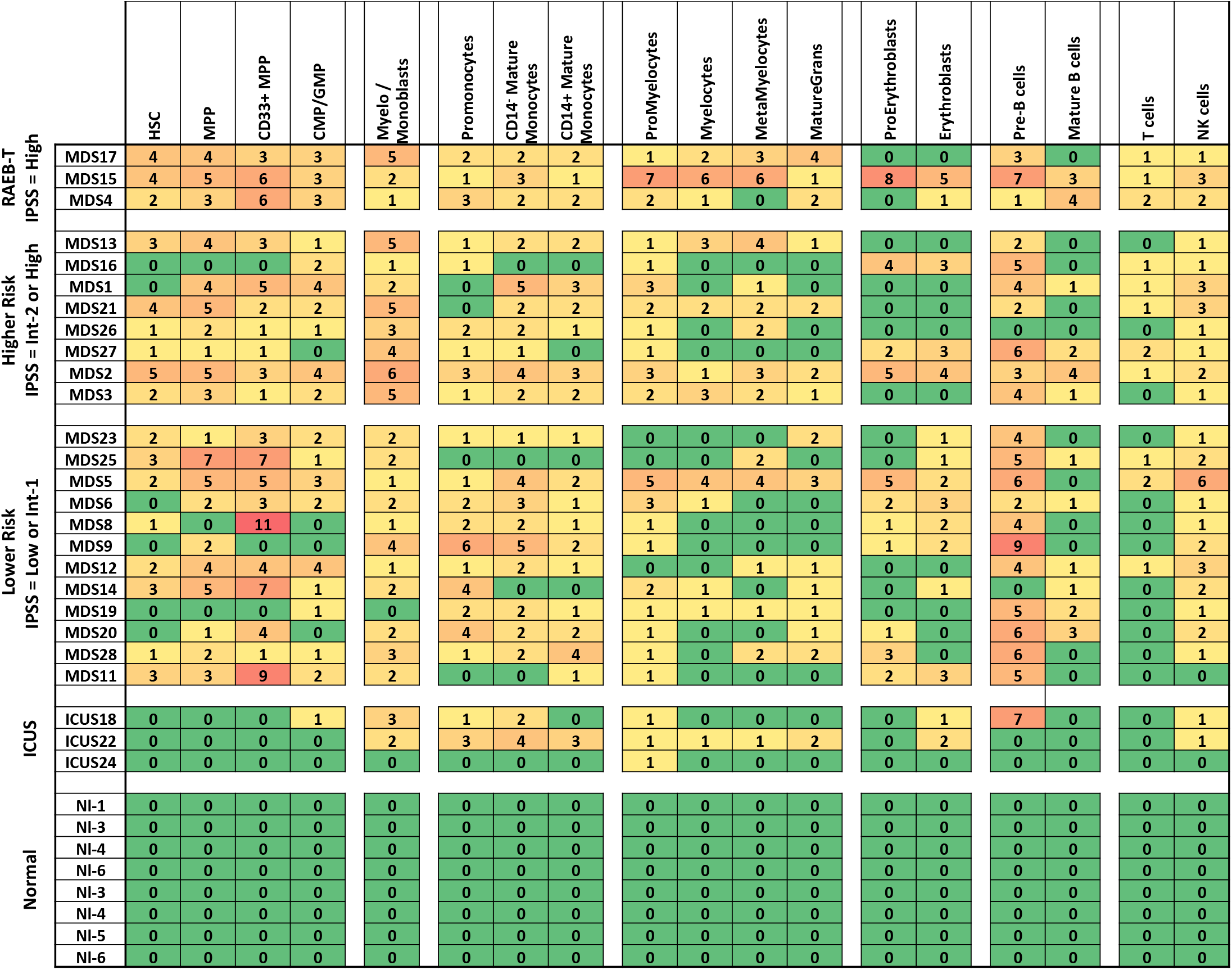
Systematic detection of multiple small aberrancies defines large immunophenotypic changes across hematopoiesis in patients with MDS. **A)** Method for defining aberrant marker expression. All immunophenotypic gates were defined on the basis of the normal samples and the same gates were applied to the each of the MDS samples. Once each population was gated, the median expression of each of the 31 surface markers was extracted and compared to the median expression in the 8 samples from five healthy donors for each gated population. MDS samples that were outside 2-fold the total variance of the normal samples were considered to be aberrant for that marker (CD117 is shown as an example). **B)** The total number of aberrant markers (of 31 measured markers) was summed for each population and each patient. Each box is colored for the number of the 31 markers that was aberrant for each patient (rows) in each gated immunophenotypic population (columns). The color scale ranges from green indicating no aberrant marker expression to the highest numbers of aberrancies colored red. The exact number of aberrant markers expressed (of the 31 tested) is printed in each box. The high rates of aberrancy observed in the pre-B cell population may be due in part to contamination of this gate with dimly CD19-positive malignant myeloid cells due to the limited number of markers defining this immunophenotypic subset (CD19 and CD10) and to the relatively dim staining of these antibodies in normal cells. Because the normal reference range used for this analysis was larger than the absolute variance of the healthy donor samples; no aberrancies were observed in the healthy donor control samples by definition. Note that samples MDS3, MDS13, and MDS21 come from serial biopsies of the same patient (each several months apart) and demonstrate consistent properties.

### Aberrant MDS immunophenotypes could be detected with multiple analysis approaches

The immunophenotypic aberrancies detected by mass cytometry could be detected regardless of the approach used to analyze the MCM data. Almost all changes noted in the manual gating analysis (Figure 2) could also be noted in the SPADE clustering (Figure 1; Supplemental Figure 3). These same abnormalities could also be appreciated in a completely independent clustering approach based on Voronoi clustering as previously described ^22^. This approach, known as X-shift, uses weighted K-nearest neighbor density estimation to find local density maxima, which then become the basis for the clustering as shown in Supplemental Figure 5. Similar to the SPADE and manual gating approaches, once cell events were organized into immunophenotypic clusters, abnormal increases or decreases in surface marker expression could be readily detected with much higher sensitivity. Almost all abnormalities could be defined in each of the three approaches (manual gating, SPADE clustering, and X-shift clustering).

Finally, we performed a manual analysis of our gated mass cytometry data in using an approximation of the European LeukemiaNET recommendations ^5,6^ as shown in Supplementary Table 6. Performing this analysis required significant modification, since side scatter is not measurable by mass cytometry, these modifications are detailed in the Supplemental Methods. Using this approach, 19 of the 23 MDS samples demonstrated abnormalities in 2 or more of the approximated parameters. These comparisons demonstrate that abnormal surface marker expression patterns detected by mass cytometry are not restricted to the approach used to analyze the data, and can be detected by a variety of novel and established analysis approaches.

### MDS stem and progenitor cells exhibit abnormal high-dimensional immunophenotypic patterns

Further analyses were focused on the stem and progenitor cell compartment (HSC, MPP, CD33+MPP), in which 21 of 23 MDS samples exhibited at least one aberrancy (average = 2.7) in one of the three populations. By contrast, no aberrancies were detected within these populations in the three samples from patients with ICUS. In addition to analysis of the total number of immunophenotypic aberrancies, specific aberrancies were more frequently found in patients with higher-risk MDS. These aberrancies were particularly informative in the immunophenotypic hematopoietic stem and progenitor cell (HSPC) compartment (lin^-^CD34^+^CD38^low^). Significant increases (~2-fold) in median expression of CD117 (p=0.007) and HLA-DR (p=0.02) were shown when comparing HSPCs from all MDS samples to the HSPCs from healthy donor samples. MDS HSPCs also exhibited a (~40%) decrease in CD34 expression (p=0.002) as compared to healthy donor HSPCs. Differences in CD117, HLA-DR, and CD34 were also demonstrated as aberrant expression patterns within gated immunophenotypic populations of cells (outside 4-fold the variance of normal) in 11/23, 13/23, and 16/23 samples, respectively. Comparison of marker expression within the HSPCs between patients with HR-MDS and LR-MDS also revealed significant differences (Supplemental Figure 6). HR-MDS HSPCs (lin^-^CD34^+^CD38^low^) were characterized by a ~2-fold increase in CD99 compared to LR-MDS (p=0.0018) and a ~3-fold decrease in CD45 compared to LR-MDS (p=8.8×10^-5^; Supplemental Figure 6). Differences in CD99 and CD45 could also be appreciated as aberrant expression patterns in 4/11 and 6/11 of the HR-MDS samples, respectively. Analysis of aberrancy in CD117, HLA-DR, CD34, CD45, and CD99 were sufficient to identify at least one aberrancy in 21 of the 23 samples, suggesting that these markers would be the best candidates for inclusion in smaller cytometry panels (Supplementary Tables 4 and 5). Three other markers, CD44, CD47, and CD321 were also commonly abnormal in the MDS HSPC population (aberrant expressed in 11, 8, and 7 samples, respectively), however, these markers were variably either high or low in MDS or (in the case of CD321) altered in a smaller fraction of cells and thus not statistically different in MDS as a whole (Figure 2B, Supplemental Figure 6, and Supplemental Tables 3, 4 and 5).

In order to simultaneously view and compare the entire aberrant expression pattern within the HSPC populations as a function of MDS risk, a visualization tool called viSNE was utilized. ViSNE is a modification of t-SNE (t-stochastic neighbour embedding) that employs a nonlinear, iterative process of single cell alignment to minimize the multidimensional distance between events and represents the high-dimensional distribution of cell events in a two-dimensional map ^21,29^. The HSPC population (lin^-^CD34^+^CD38^low^) of each sample was analyzed along with the five healthy donor samples. Healthy donor samples exhibited a consistent localization on the viSNE plot and patients with low-risk MDS consistently demonstrated viSNE patterns with small differences compared to healthy controls. Samples from patients with the RA-RS MDS subtype were most similar to the viSNE patterns exhibited by the control samples (Figure 3). By contrast, most patients with Int-1 and all patients with higher-risk MDS exhibited viSNE patterns that were clearly different from the healthy donor samples (Figure 3). These differences in HSPC immunophenotype were confirmed by an independent multidimensional binning approach (Supplemental Figure 7) that also demonstrated that the HSPCs from samples of healthy controls and two of the three patients with ICUS clustered together in a region of the dendrogram distinct from the MDS samples. To further assess the relevance of these global differences in surface marker expression pattern, the sum total fold increase in median expression relative to normal was calculated for the HSPC population (as detailed in Supplemental methods) and this sum was compared to survival for the 11 patients (13 samples) for whom survival endpoints have been reached. The sum total of HSPC marker abnormality was inversely correlated with survival time from biopsy R=-0.56 with a p<0.05, supporting that metrics of surface marker expression in the HSPC population could have relevance for monitoring clinical disease progression in MDS (Supplemental Figure 8), as has been seen in other cytometric analyses of MDS ^26^.

**Figure 3:**
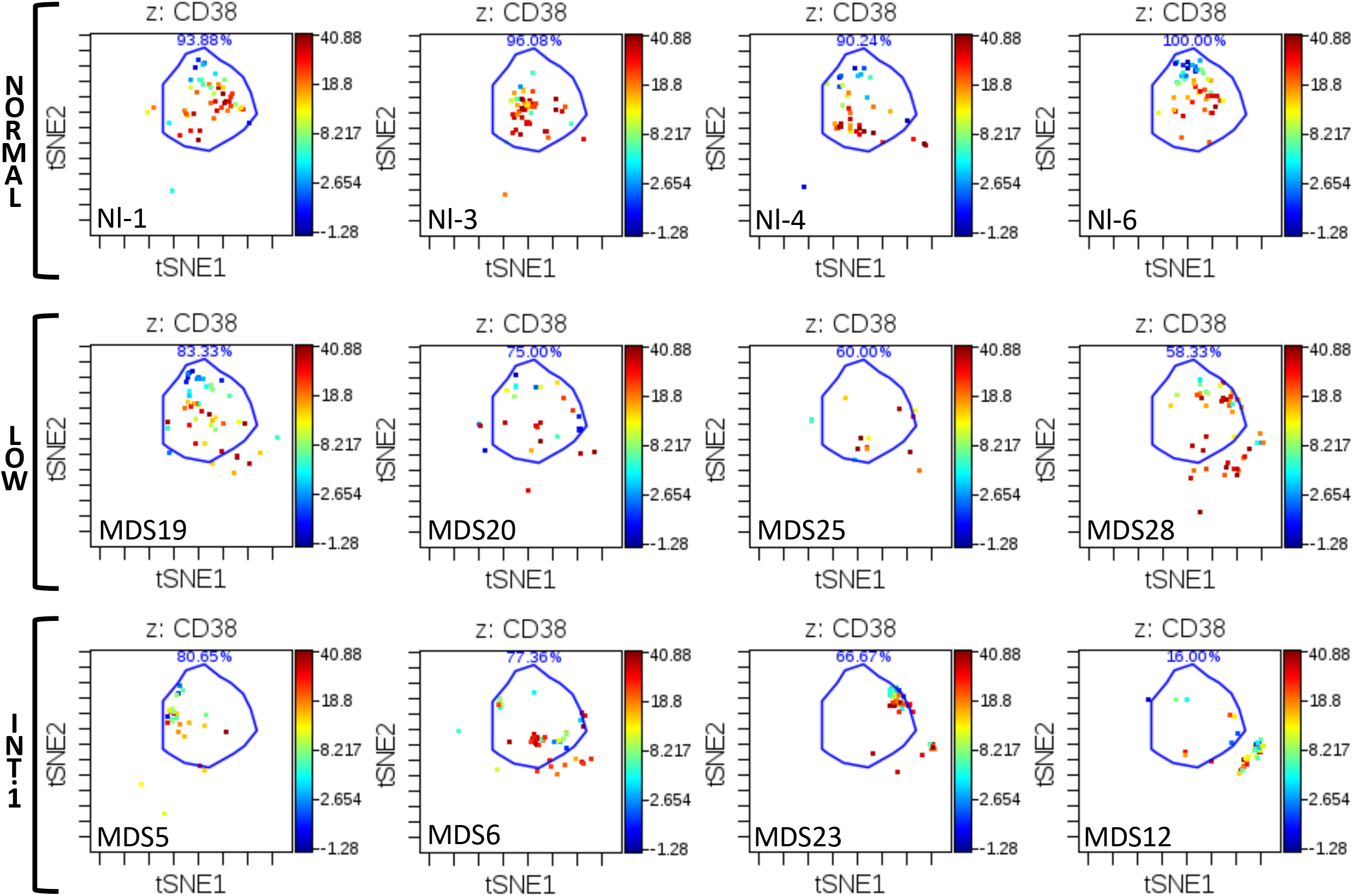
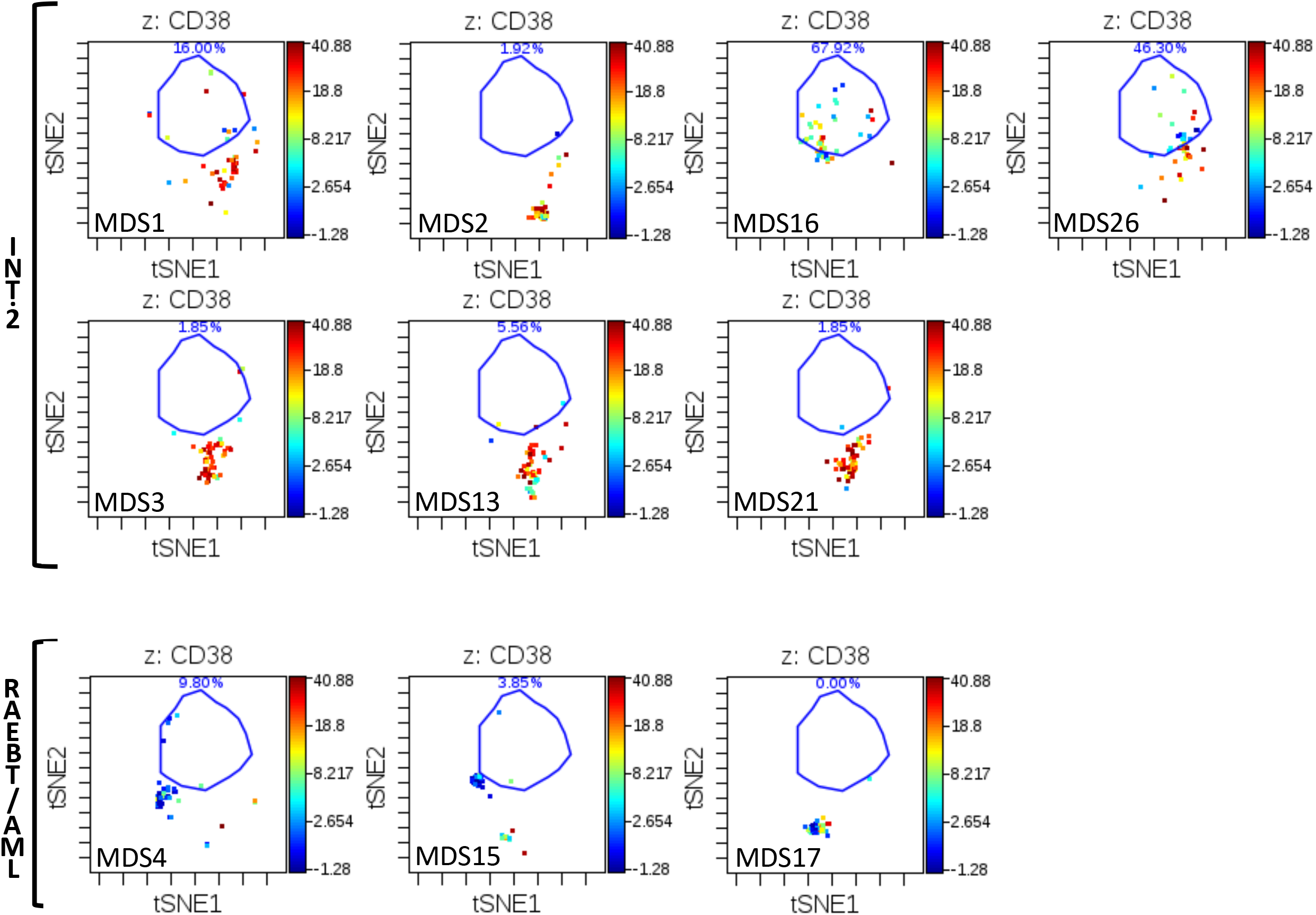
viSNE analysis of CD34^+^CD38^low^ subset reveals distinct immunophenotypic patterns in high-dimensional space. Each sample was analyzed by viSNE (up to 5,000 sampled events per individual) using 19 dimensions (Supplemental Table 1). A gate (light blue line) encompassing the vast majority of normal CD34^+^CD38^low^ events is shown for reference. The MDS subtype and sample is indicated for each viSNE map. Each cell event is colored for it’s expression level of CD38 from blue (0 ion counts) to red (approximately 40 ion counts). Red cell events still fall within the CD34^+^CD38^low^ gate and demonstrated dim CD38 expression. Note that samples MDS3, MDS13, and MDS21 come from serial biopsies of the same patent (each several months apart) and demonstrate consistent properties.

### Patterns of cell frequency distribution correlate with clinical risk and allow for automated classification

The distribution of cell frequencies across the immunophenotypic populations (by SPADE analysis or manual gating) demonstrated that the pattern of cell frequencies throughout development was correlated with the clinical risk of each patient (Figure 4). The most significant single distinguishing feature between clinical risk groups was the increased frequency (>40-fold) of immunophenotypic HSPCs in HR-MDS compared to LR-MDS (p=9×10^-7^) or normal (p=6.3×10^-6^). Furthermore, this high-parameter analysis detected a >12-fold increase in the HSPC frequency in 2 patients with IPSS Int-2 disease with blast frequencies of <5% following therapy (MDS#21 and MDS#27), similar to a previous study using fluorescent flow cytometry ^30^. The frequency of cells in each SPADE cluster was then used to perform a hierarchical clustering of the global cell distribution across development in all samples. This approach clustered patients into groups with different clinical risk as shown in Figure 5. As this analysis was based on the simultaneous analysis of all 19 surface markers utilized in the clustering and on all cells in each sample, the results were stable across a variety of perturbations. For example, removing data regarding the expression of CD34 did not reduce the ability of the clustering analysis to group similar risk sample together, and resulted in a nearly identical hierarchical tree, demonstrating the robustness of this analytic approach (Figure 5). Again, an independent binning analysis of all cell events resulted in a similar hierarchical tree (Supplemental Figure 9).

**Figure 4:**
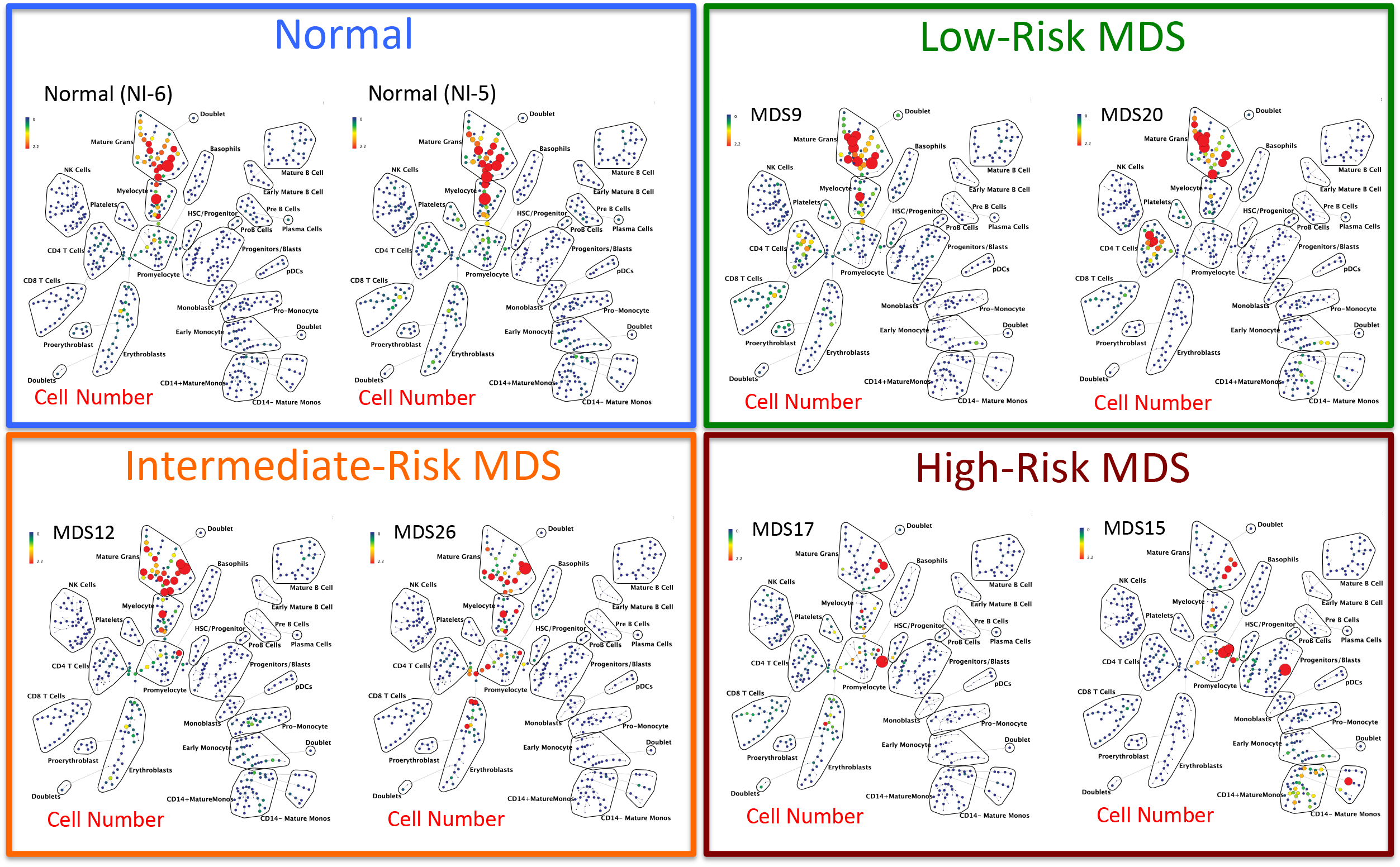
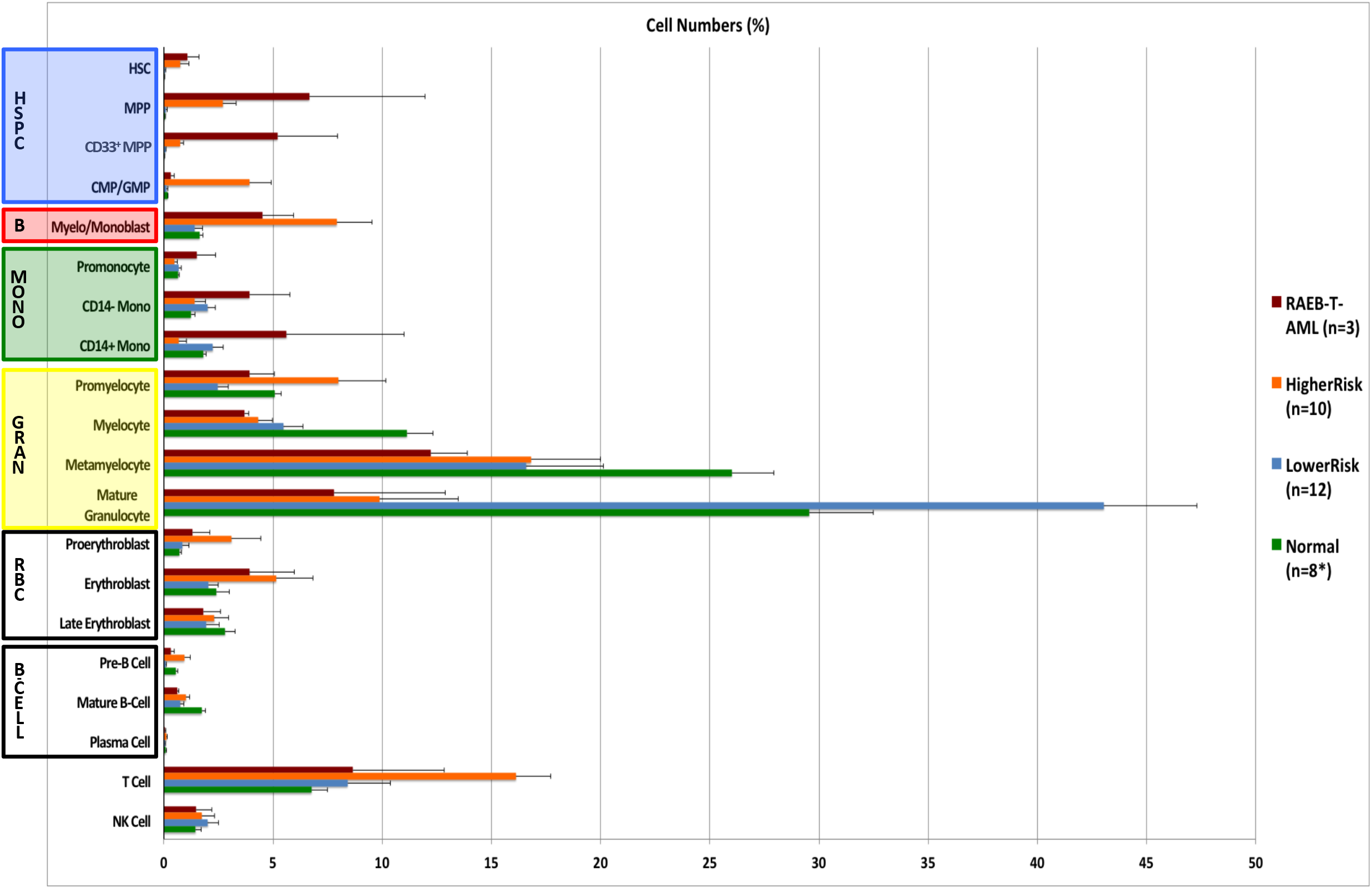
Distribution of cell frequency across hematopoietic development correlates with MDS risk. **A)** SPADE tree colored for the fraction of total cells in each node from lowest (blue) to highest (red). The size of each node is correlated to the fraction of cells mapping to the node; however, a minimum size was enforced for most nodes to allow visualization of node color. **B)** The frequency of cells in the indicated (manually-gated) stem and progenitor cell compartments for patients of each MDS risk group. Error bars indicate standard errors of the means. *The 8 replicate normal samples came from 5 donors.

### Modeling of aberrant differentiation in MDS

As these results suggested that analysis of global differentiation changes may be a diagnostically useful parameter in MDS, viSNE was used to assess cell immunophenotype across the developmental progression from HSCs to both mature granulocytes and mature monocytes by manually removing all cell events expressing antigens of non-granulocyte or non-monocyte lineages (respectively) and aligning the remaining cells in single viSNE analysis of both lineages. The approach is outlined in Supplemental Figure 10. By using gated populations of healthy donor cells at each defined immunophenotypic stage of development, this approach allows for the characterization of the high dimensional aberrations in MDS cell immunophenotype that occur during the progression of hematopoiesis. As shown in Figure 6, developmental progression is relatively normal in low-risk MDS, where a relative reduction in the frequency of committed progenitors and fully differentiated cells in the monocyte and granulocyte lineages is the most prominent finding. By contrast, in higher risk MDS samples (IPSS Int-2 and high) cell development frequently occurred along an abnormal path intermediate between granulocytes and monocytes with a relative or even absolute absence of normally differentiated granulocytes and monocytes (Figure 6). This developmental path ended in a population of immunophenotypically aberrant myeloid cells (IAMCs) that could also be identified by manual gating (Supplemental Figure 1) and were characterized by expression of CD64, CD11b, CD44, CD45, CD45RA (all at moderate levels) and the near complete absence of HLA-DR (<10% the intensity of monocyte expression), and lack of CD14 and CD15 (Supplemental Figure 11 and Supplemental Table 7). This phenotype is potentially consistent with myeloid derived suppressor cells, however, this mass cytometry assay panel was not capable of further functional characterization. These IAMCs were rare in the healthy donor samples (0.48% on average) while 10 of the MDS samples exhibited these cells at a frequency of >2.5% and 5 MDS samples had a frequency of greater than 10% (Supplemental Figure 12). Patients with higher-risk MDS (RAEB-T/AML, IPSS-High and IPSS-INT-2) demonstrated an increase in the frequency of this cell population compared to the healthy donors (p=0.012) or patients with lower-risk MDS (IPSS-Low and IPSS-Int-1; p=0.029). This type of progression modeling represents another objective and potentially automatable approach enabled by high dimensional cytometry that could further improve the recognition of abnormal hematopoietic development in patients with MDS.

## Discussion

### Mass cytometry enables high-dimensional characterization of aberrant marker expression in MDS

This first application of MCM for the analysis of MDS detected all major established aberrant expression patterns in MDS, as well as novel aberrant expression patterns of CD321, CD99 and CD47. Importantly, using high-parameter single-cell analysis and internal normal reference samples, we detected deviations from the immunophenotypic boundaries of normal hematopoiesis in every analyzed MDS sample and in at least one sample for 27 of 31 markers tested. The detected abnormalities found in this mass cytometry data were not specific to any single data analysis approach but were detected by i) manual gating with comparison to normal samples, ii) SPADE clustering with comparisons of each MDS cell cluster to the normal cells in the same cluster, iii) X-shift density maxima clustering with comparison of each density-derived cluster of MDS cells to the normal cells of the same cluster. The high degree of similarity in the results of these independent analytic approaches strongly suggests that the high sensitivity of mass cytometry for detecting abnormalities in MDS is a property of simultaneously measuring a large number of surface marker parameters and not simply the result of a specific approach used to analyze the resulting data.

The HSPC cell compartment appeared to be particularly useful for assessment of immunophenotypic aberrancies. This cell compartment was immunophenotypically normal in all three patients from our cohort with ICUS, and was abnormal (by one or more of the high dimensional analyses) in 21 of 23 samples from patients with MDS (Figures 1 and 2 and Supplemental Figures 5 and 6). The most useful markers were CD117, HLA-DR, CD34, CD45, and CD99, which were consistently high or low in MDS, while CD44 and CD47 were commonly abnormal but could be either high or low in any one MDS sample (Figure 1, Supplemental Figure 3, Supplemental Tables 3, 4, and 5). High dimensional analysis likely exhibits this additional sensitivity for detection of immunophenotypic variations because of the ability to group cells into developmentally similar immunophenotypic clusters, thereby allowing comparisons to be made across developmentally similar MDS and control cell populations. A similar increase in the sensitivity of aberrant marker detection was found in our previous mass cytometry studies of AML ^17^, and a recent report noted improved prognostic accuracy of gene expression profiling when analysis was performed based on the differential expression patterns between AML populations and developmentally similar normal cells ^31^.

### High-dimensional analysis detects altered developmental patterns characteristic of MDS

The measurement of all surface markers simultaneously on all cells enabled an extremely detailed analysis of hematopoietic development in MDS through the use of high-dimensional clustering of all bone marrow cells from each sample. This clustering demonstrated consistent distributions of normal cell frequencies across each immunophenotypic cluster in healthy donors. Changes in these frequency distributions (across immunophenotypic stages of development) could then be analyzed by comparing the cell frequency distributions of the MDS and healthy donor samples. These analyses demonstrated that every patient with MDS in this cohort exhibited an alteration in how cell frequencies were distributed across these immunophenotypic populations. This effect could be observed when clustering was performed on the basis of the SPADE analysis (Figure 5) or on the basis of a multidimensional binning approach (Supplemental Figures 7 and 9). These frequency differences were greatest in patients with the highest risk disease, and could potentially be used to classify clinical risk or disease subtype as has previously been reported for blast cell populations in patients with MDS ^4 30^.

Importantly, the utility of this holistic clustering appeared to derive from the relationship of all of the cells to one another, and appears to be independent of any one cell surface marker, such as CD34 (Figure 5). This approach is fundamentally different than most common approaches for the use of flow cytometry in the diagnosis of MDS, in which the marker expression or frequency of specific small cell subsets such as B cell progenitors or myeloblasts is utilized for diagnosis ^4–6^. In this regard, a major potential advantage of high-dimensional cytometry analysis (and clustering of the resulting data) was that every cell from each patient sample could be placed into a single cluster and is represented only once in the analysis. This made the clustering less sensitive to any one individual marker and increased the informational utility of cells that did not bind particular markers (because even cells negative for most or all markers clustered into distinct groups). This also allowed the analysis to be performed in an unsupervised manner, since the formation of clusters does not require human interpretation of the data and (if performed on a sufficiently large scale) abnormal samples could potentially be identified on an automated basis by comparing a given test sample’s frequency distribution to that of many other known MDS and normal samples. Additionally, the use of an unbiased, high-dimensional approach allowed identification of an immunophenotypically aberrant myeloid cell population with a phenotype similar to MDSCs identified by other researchers in patients with MDS ^32,33^. The observation that these cells appear to develop along an aberrant differentiation trajectory (Figure 6 and Supplemental Figure 11) is potentially consistent with other models of MDSC ontogeny ^34^ and was not an *a priori* expectation of the analysis.

**Figure 5:**
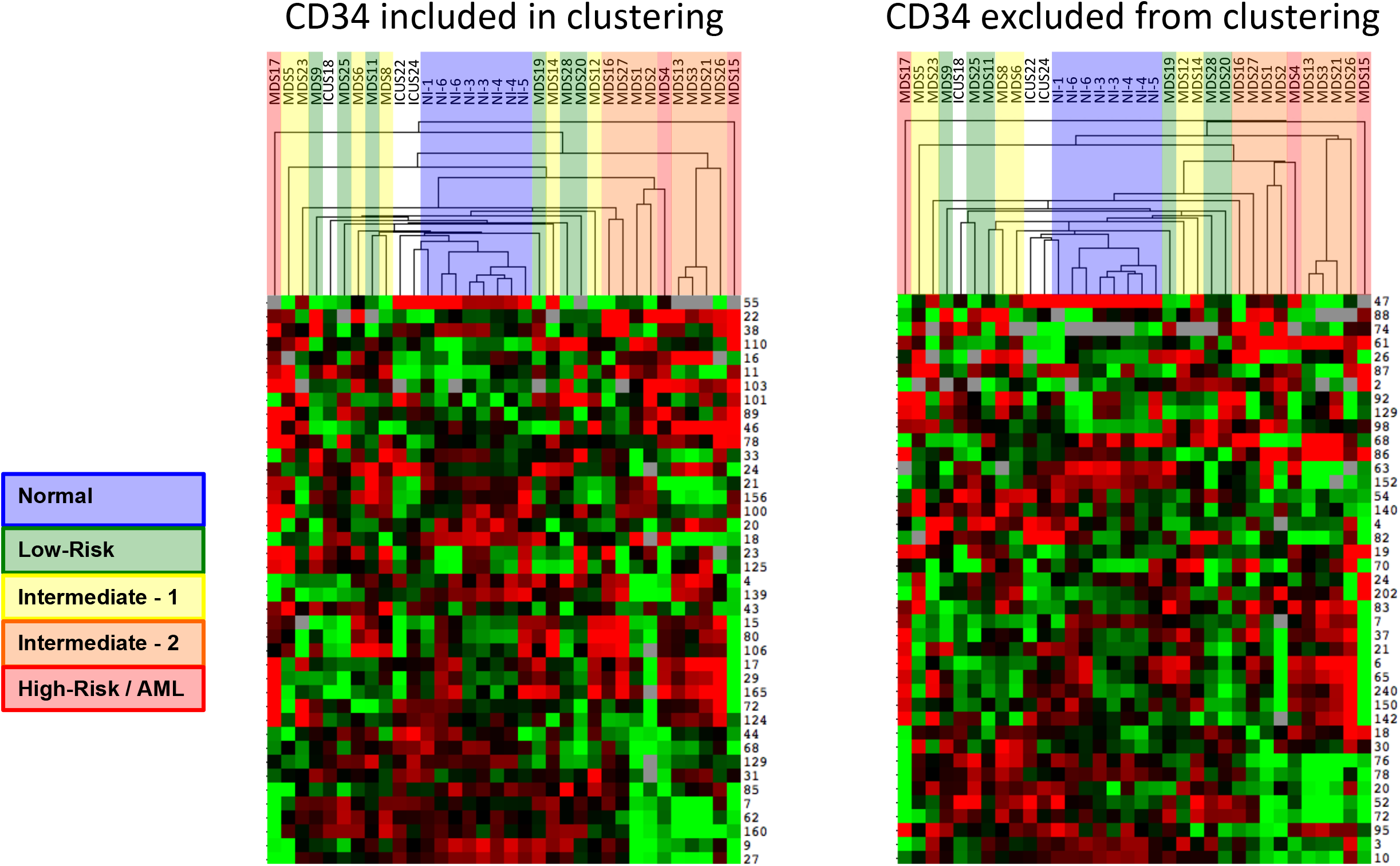
Clustering of cell frequency across SPADE nodes in MDS and control samples groups control samples and MDS samples into groups of similar clinical risk. SPADE analysis of total events from each MDS and control sample was performed, and the frequency of cell events within each cluster was extracted from the resulting SPADE trees. The cell frequency by node was then entered into a biaxial clustering analysis (similar to gene expression array analysis). A portion of the cell frequency heat map is shown, each row represents the relative cell frequency in the indicated SPADE node for each sample (columns). Red indicates higher cell fractions and green lower cell fractions. The dendrogram at the top of the figure demonstrates how the different MDS and control samples grouped together. Each sample is colored by its clinical risk as indicated. The analysis was performed once with CD34 used for generation of the SPADE tree and then again with CD34 ignored during SPADE tree generation. In both analyses, patients with higher clinical risk cluster further from the normal samples. Note that samples MDS3, MDS13, and MDS21 come from serial biopsies of the same patient (each several months apart) and demonstrate consistent properties.

**Figure 6:**
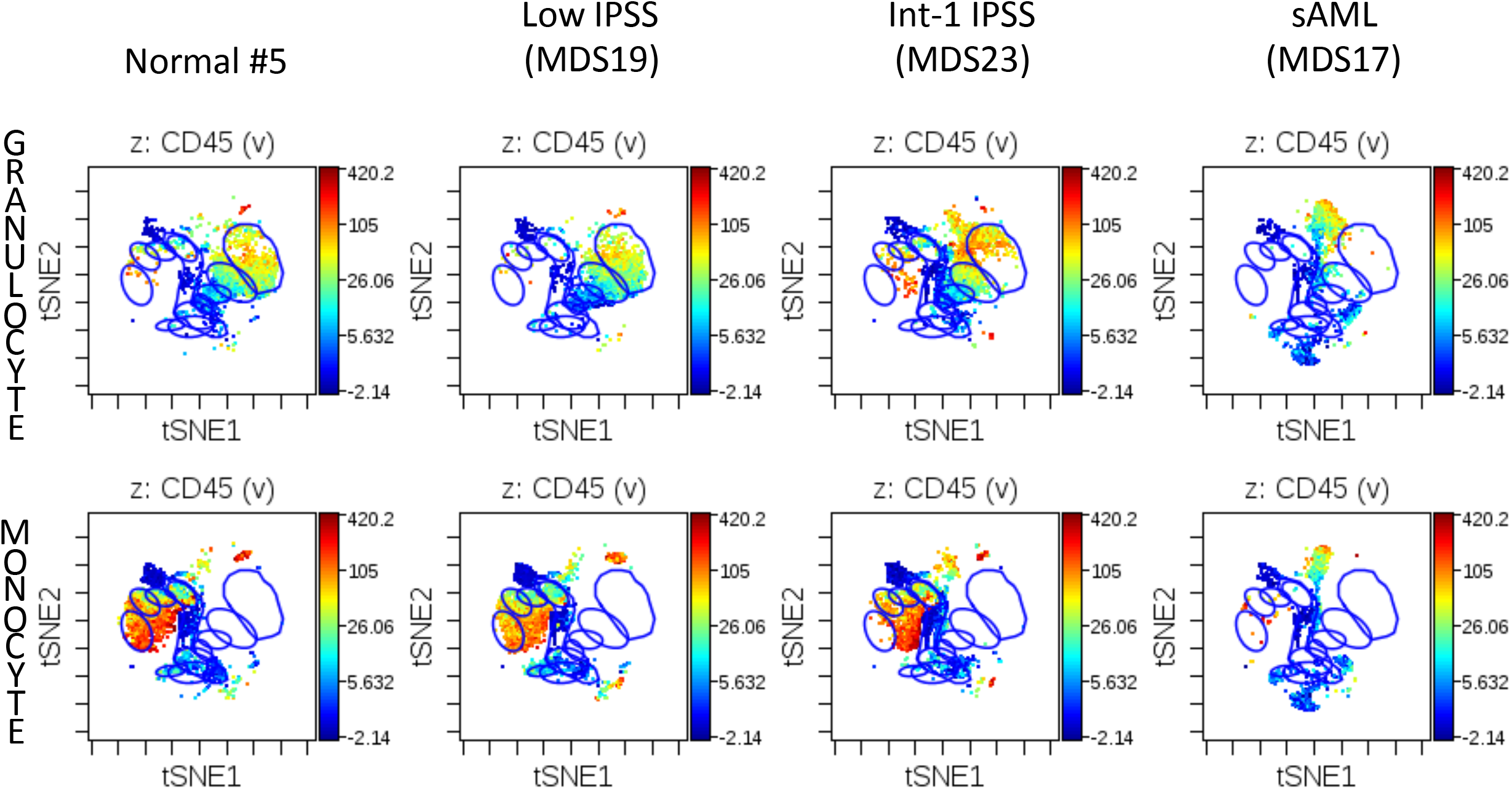
Developmental progression in control and MDS samples during granulocyte and monocyte differentiation. Blue gates on the viSNE projection show the positions of the normal immunophenotypic developmental stages as described in Supplemental Figure 10.

### Role of High-Dimensional Cytometry in MDS

While recent flow cytometry analysis approaches are sensitive and moderately specific for MDS ^5^, the results presented here suggest that the use of high-dimensional cytometry approaches could potentially enhance both the sensitivity and specificity of cytometry in MDS diagnosis. Consistent with earlier reports,^30^ our work suggests that a focus on HSPC cell marker aberrancy (enabled by the simultaneous measurement of CD34, CD38, and lineage markers) combined with the use of internal cell controls can sensitively detect MDS. Additionally, we demonstrate that use of high parameter clustering analyses can enable a global modeling of differentiation, which is likely to also be predictive of MDS subtype and outcome as had been suggested by earlier studies ^4^. Our findings strongly suggest that high parameter cytometry combined with global analysis approaches could enhance the diagnostic utility of cytometry in MDS. However, the results presented here remain preliminary and validation of this approach for MDS diagnosis and characterization will require much larger patient cohorts with age-matched healthy donors and patients with ICUS, as were previously used to characterize other flow cytometric diagnostic approaches ^5^. We believe that this preliminary report strongly suggests that such studies are warranted. The data presented here lay a foundation and a strong rationale for design of larger studies that could develop mass cytometry or high dimensional fluorescent flow cytometry as a tool for such analyses in MDS.

## Acknowledgements

The authors wish to thank Angelica Trejo and Astraea Jager for excellent technical support and the members of the Stanford University Hematology Tissue Bank for assistance with fresh sample collection. We also wish to thank the patients who contributed tissue samples, without which this work would not have been possible.

This work was supported by the following grants to the authors:

**G.K.B. -** Stanford Cancer Institute Developmental Cancer Research Award **P.L.G.** - Stanford MDS Center Research Fund, and the William E. Walsh Leukemia Research Fund **G.P.N. - NIH -** R33 CA0183692, U19 AI057229, 1U19AI100627, Subcontract 7500108142, R01CA184968, 1 R33 CA183654-01, 1R01GM10983601, 1R21CA183660, 1R01NS08953301, 5UH2AR067676, 1R01CA19665701, R01 HL120724, N01-HV-00242 HHSN26820100034C, U54-UCA149145A; **DOD -** Ovarian Cancer Teal Innovator Award 0C110674, DOD W81XWH-14-1-0180; **FDA -** BAA-15-00121, HHSF223201210194C; Bill and Melinda Gates Foundation - OPP1113682; Novartis Pharmaceuticals Corporation - CMEK162AUS06T; Pfizer, Inc. - 123214

## Authorship

G.K.B., P.L.G., W.J.F. and G.P.N. conceived and designed all experiments and interpreted the data; G.K.B. wrote the manuscript; G.K.B., P.L.G., and G.P.N. reviewed and edited the manuscript; G.K.B. performed all experiments; G.K.B., R.F., and N.S. performed data analysis; G.K.B. and K.S. collected and processed patient samples and extracted clinical data.

Reprint requests and other correspondence should be sent to either: Garry P. Nolan, Ph.D. 3220 CCSR, Baxter Laboratory, 269 Campus Dr., Stanford, CA 94305, Phone: (650) 725-7002, Fax: (650) 723-2383; or to: Gregory K. Behbehani, M.D., Ph.D., 416 Biomedical Research Tower, 460 W. 12^th^ Ave., Columbus, OH 43210, Phone: (614) 685-6015, Fax

