## Supplemental Methods for "Profiling Myelodysplastic Syndromes by Mass Cytometry Demonstrates Abnormal Progenitor Cell Phenotype and Differentiation"

### *Mass cytometry*

After completion of antibody staining, cells were washed twice with CSM and then incubated overnight or for 36 hours (samples were split in half due to the large cell numbers) in PBS with a 1:5000 dilution of the iridium intercalator pentamethylcyclopentadienyl-Ir(III)-dipyridophenazine (Fluidigm Sciences, Toronto, Canada) and 1.5% paraformaldehyde. Excess intercalator was then removed with one CSM wash and two washes in pure water. Cells were then resuspended in pure water at approximately 1 million cells per mL and mixed with mass standard beads (Fluidigm Sciences). Cell events were acquired on the CyTOF™ mass cytometer (Fluidigm Sciences) at an event rate of 100-300 events per second with instrument-calibrated dual-count detection [1]. Noise reduction was used, a cell length of 10-90, and lower convolution threshold of 200. After data acquisition, the mass bead signal was used to correct short-term signal fluctuation during the course of each experiment and bead events were removed. [2].

### *Immunophenotypic aberrancy analysis*

This study is unique in the highly parametric nature of our data and the high precision of the measurements enabled by cellular barcoding. This allowed for the creation of a new method of identifying surface marker aberrancies. To do this, the median expression level of each marker in each gated population from each patient and healthy control samples was calculated after gating the normal cell populations into developmental immunophenotypic subsets on the basis of standard surface markers (as in Supplementary Figure 1) and applying these gates (without modification) to all samples. Since the 8 replicate normal samples analyzed came from five healthy donors, an MDS sample aberrancy (for any given marker) was defined conservatively as an MDS sample median expression level (in a given gated population) greater than or less than the median of the similarly gated healthy bone marrow cell population plus or minus twice the absolute variance of the healthy control samples. For example, over-expression of CD117 in the multipotent progenitor (MPP) population of MDS sample #15 was determined as follows: MDS#23$^{MPP}$ median CD117 expression > average normal sample$^{MPP}$ median CD117 expression + 2 times the absolute variance of normal sample$^{MPP}$ CD117 expression. Specifically in this example, the median CD117 expression level from the cells of MDS sample #15 within the MPP gate was 10.6 counts, which was greater than the average median expression level of the healthy control cells within this gate, 2.3 counts, plus two times the absolute variance between the controls samples in the MPP gate which was 1.4 counts (2 * 0.7 counts = 1.4 counts; 2.3 counts + 1.4 counts = 3.7 counts; 10.6 counts > 3.7 counts). This process was repeated for each of the 30 other measured surface markers in each of 30 gated immunophenotypic cell populations from every MDS sample. The summed number of markers with aberrant expression patterns was calculated for each gated immunophenotypic population from each patient sample.

For the analysis in Supplemental Figure 8, the absolute difference between the median expression level of each marker and the average of the median expression levels from the normal samples was calculated for the CD34$^{+}$CD38$^{low}$ population from each sample. This absolute difference was then divided by the average median expression level of the normal samples. In cases where the average of normal expression was less than 1 ion count, the absolute difference between each sample median and the normal median was divided by 1 instead. These relative differences for each measured surface marker were then summed across all of the measured surface markers for each sample and compared to patient survival (from the time of sample collection if the survival outcome was known) in Supplemental Figure 8.

Immunophenotypic gates used in manual gating were defined based on the normal donor cell samples (Supplementary Figure 1). For some populations, CD33 appeared to better discriminate immunophenotypic hematopoietic stem cell (HSC) and multipotent progenitor (MPP) populations from more mature populations and was thus used instead of CD45RA, which did not stain as brightly by mass cytometry; the resulting CD33-negative HSC and MPP populations were all negative for CD45RA as well. Within the $Lin^{-}CD34^{+}CD38^{low}$ population, there was a distinct population positive for CD33 with a level of staining equal or greater than the CMP/GMP population. This CD33 positive population was gated separately as $CD33^{+}$MPP cells. While CD123 staining gave expected staining of basophils and pDCs, the CD123 signal did not allow for clear resolution of the CMP and GMP populations from within the $Lin^{-}CD34^{+}CD38^{+}$ population. Because of this, as well as controversy regarding the ability to reliably resolve these populations by flow cytometry [3], these cells were gated as a combined CMP/GMP population. CD123 was still assessed for aberrant expression, however, and was aberrantly expressed in three samples (Supplemental Tables 3, 4, and 5). The mass cytometry experimental design allowed CD123 to be simultaneously used to gate out cells with very high levels of CD123 (cells that were clearly pDCs and basophils based on multiple other markers) while still allowing us to detect several fold aberrant increases in CD123 on immature myeloid and progenitor cells. The CD123 lineage exclusion gates were carefully designed to not exclude immature cells that were likely to be part of the MDS clone. An example of this is shown in Supplemental Figure 13. The ability to use the same markers for both lineage removal and aberrant marker detection represents another advantage of high-parameter analysis.

***Barcoding***

Mass-tag barcoding was performed in groups of 20 samples using a transient partial permeabilization protocol [4]. The unique pattern of three of the six stable palladium (Pd) isotopes enabled removal of doublet events [5]. Each barcoding plate included at least three sample aliquots from one of the five healthy donors. An aliquot of the sample from donor #6 (Nl-6) was included in each barcoding plate and staining reaction as an internal reference standard to ensure that staining and antibody detection was consistent. Barcoding was performed on approximately 2 million fixed cells per sample placed into racked, 1.1-mL microtubes (BioExpress, Kaysville, UT, USA) using a multichannel pipette and a multichannel aspirator. Fixed cells were washed once in CSM and then washed once in PBS, followed by a second wash in PBS plus 0.02% saponin (Sigma-Aldrich, St. Louis, MO, USA). All pre-barcoding saponin washes were performed at 4 °C. A 100x DMSO stock of the mass tag barcoding reagent was then rapidly (<20 seconds) diluted into 1 mL ice-cold PBS plus 0.02% saponin and then quickly (<20 seconds) applied to the resuspended cells. Cells were incubated for 15 minutes (at room temperature) to allow covalent reaction of the barcode mass tags with the cells. After barcoding, cells were washed twice with CSM and then combined in a single tube. Cells were not re-exposed to saponin in subsequent manipulations or antibody staining steps. All antibody staining, methanol permeabilization, and sample measurement was performed with all cells (~40 million total) simultaneously in the same tube.

Mass-tagged barcoding reagents were prepared as described [5]. Briefly, barcoding was performed with a pattern of three of the six stable Pd isotopes (102, 104, 105, 106, 108, 110) for each sample using isotopically purified palladium nitrate (Trace Sciences International, Richmond Hill, Ontario, Canada) and isothiocyanobenzyl-EDTA (Dojindo Molecular Technologies, Rockville, MD, USA) as the chelator. Mass tag barcoding was performed at a final metal concentration of 300 nM; staining was equivalent for all Pd isotopes. The barcode signal from each cell was de-convoluted back into individual samples using a Matlab software application, which also allowed removal of doublet events.

### *Data analysis and gating*

All mass cytometry data are displayed with an arcsinh transformation and a scale argument of five (except for linear scales used for Ir intercalator and cell length parameters). During data acquisition the cell subtraction value was set to -100 (thereby adding 100 counts to each channel). After acquisition, the effect of the cell subtraction setting was negated by subtracting a value of 100 from every channel of each FCS file using the flowCore package for R (10). These manipulations were performed to better estimate the effect of background subtraction and experimental noise for cells with low signal by allowing negative values to be displayed [6].

The manual gating strategy for each population is shown in Supplemental Figure 1. The basic strategy used was to first create lineage restricted populations for the lineage negative cells, the monocyte lineage, granulocyte lineage, erythroid lineage, and B lineage cells and then distinct developmental stages within each population could be gated. Additionally, several other populations were gated as a single group: myelo/monoblasts, basophils, platelets, plasmacytoid dendritic cells (pDCs), T cells, NK cells, and plasma cells.

Lineage Gates:

**Lineage negative** cells were defined as meeting all of the following criteria: 1. CD11b$^{\mathbf{low}}$; 2. Not brightly CD38 positive (i.e. Not plasma cells); 3. Not CD321 bright and DNA$^{\mathbf{low}}$ (i.e. Not platelets); 4. CD3 negative (i.e. Not T cells); 5. Not CD45$^{\mathbf{hi}}$ and CD7$^{\mathbf{hi}}$ (i.e. Not NK cells; note that this gate was slightly adjust for a few samples to separate NK cells from myeloid cell aberrantly expressing high levels of CD7); 6. Not $CD19^+$ or $CD20^+$ (i.e. Not B cells); 7. Not $CD71^{\mathbf{hi}}$ or $CD235^{\mathbf{hi}}$ (i.e. Not committed erythroid progenitors; these cells were confirmed to be CD45low to ensure activated cells of other lineages were not excluded by the CD71 gate); 8. Not $CD123^{\mathbf{hi}}$ (i.e. Not pDCs or basophils; boundaries of this gate were established by back-gating on basophils, pDCs and myeloid blasts with aberrant CD123 expression to ensure that boundaries of this single gate excluded the former populations without excluding the latter); 9. Not CD33 bright (i.e. Not mature monocytes; gate boundaries were determined by back-gating on mature monocytes as well as normal and malignant progenitor populations to ensure the boundaries of this single gate excluded the former without excluding the latter).

**Monocyte Lineage** cells were defined as meeting all of the following criteria: 1. Not $CD34^+CD38^{low}$; 2. CD33 positive; 3. HLA-DR positive; 4. Not CD123 bright and $CD33^{low}$ (i.e. Not basophils or pDCs); 5. CD3 negative (i.e. Not T cells); 6. Not CD19+ or CD20+ (i.e. Not B cells); 7. Not CD45 high and CD7 high (i.e. Not NK cells; note that this gate was slightly adjust for a few samples to separate NK cells from myeloid cell aberrantly expressing high levels of CD7); 8. Not brightly CD38 positive (i.e. Not plasma cells); 9. Not CD71high or CD235 high (i.e. Not committed erythroid progenitors; these cells were confirmed to be CD45low to ensure activated cells of other lineages were not excluded by the CD71 gate). From this monocyte-restricted population, immature promonocytes could be gated based on low CD11b and CD14 expression, CD14 negative monocytes could be gated as $CD11b^+CD14^{low}$, and $CD14^+$ monocytes could be gated as $CD14^+CD11b^+$.

**Granulocyte Lineage** cells were defined as meeting all of the following criteria: 1. Not CD71high or CD235 high (i.e. Not committed erythroid progenitors; these cells were confirmed to be

CD45low to ensure activated cells of other lineages were not excluded by the CD71 gate); 2. CD3 negative (i.e. Not T cells); 3. Not CD19+ or CD20+ (i.e. Not B cells); 4. Not brightly CD38 positive (i.e. Not plasma cells); 5. CD15 positive. From this granulocyte-restricted population, promyelocytes were gated based on lack of CD16 and CD11b, Myelocytes were gated as $CD11b^{mid}CD16^{low}$, Metamyelocytes were gated as $CD11b^{+}$ and $CD16^{low}$, and mature granulocytes were gated as $CD11b^{+}$ and $CD16^{+}$.

**Erythroid Lineage** cells were defined as meeting all of the following criteria: 1. Not $CD34^{+}CD38^{low}$; 2. CD3 negative (i.e. Not T cells); 3. Not $CD19^{+}$ or $CD20^{+}$ (i.e. Not B cells); 4. CD15 negative (i.e. Not granulocyte lineage). From within this population, CD71 bright CD235 low cells were gated as pro-erythroblasts, CD71 bright CD235 positive cells were gated as Early erythroblasts, and late erythroblasts were defined as CD235 positive $CD71^{mid}$, CD45 negative, and CD321 negative (i.e. not platelets).

**B cell Lineage** cells were defined as meeting all of the following criteria: 1. CD33 low (i.e. not monocytes); 2. Not CD321 bright and $DNA^{low}$ (i.e. Not platelets); 3. CD3 negative (i.e. Not T cells); 4. Not CD71high or CD235 high (i.e. Not committed erythroid progenitors; these cells were confirmed to be CD45low to ensure activated cells of other lineages were not excluded by the CD71 gate); 5. $CD11b^{low}$ (i.e. Not mature myeloid). From this population, CD19 positive and $CD20^{low}$ cells that also did not express extremely high levels of CD38 (to exclude plasma cells) were defined as pre-B cells, while cells that were positive for both CD19 and CD20 were gated as mature B cells.

**Early progenitor populations** were defined from within the lineage negative population. HSCs were defined as $CD34^{+}CD38^{lo}$, CD45RA negative, CD33 negative (this was a lower threshold than used for the lineage gating), and CD90 positive. MPP cells were defined as $CD34^{+}CD38^{lo}$, CD45RA negative, CD33 negative (this was a lower threshold than used for the lineage gating), and CD90 negative. CMP/GMP cells were defined as $CD34^{+}CD38^{lo}$ and lineage negative. Additional populations from within the lineage negative gate could be identified that do not have a clear correlate in traditional gating strategies: $CD34^{neg}CD38^{+}$ Undifferentiated, $CD34^{neg}CD38^{neg}$ Undifferentiated, and $CD33^{+}$MPP cells (gated as MPP cells above except that these cells have levels of CD33 expression that are clearly positive but less than a mature myeloid cell).

**Plasma cells** were defined as being very bright for CD38, negative for CD11b, negative for HLA-DR, and completely negative for CD33.

**T cells** were defined simply by positivity for CD3.

**NK cells** were defined by all of the following: positivity for CD7 and CD45 (note that this gate was slightly adjust for a few samples to separate NK cells from myeloid cell aberrantly expressing high levels of CD7), expression of CD7 in the absence of CD3, and lack of CD34 expression. Note that almost all of these cells were positive for either CD56 and/or CD16, but these markers were not used to define the population.

**Plasmacytoid Dendritic cells (pDCs)** were defined as cells meeting all of the following criteria: $CD123^{hi}$ and $CD33^{lo}$, HLA-DR positive, and not $CD34^{+}CD38^{lo}$.

Basophils were defined as cells meeting all of the following criteria: $CD123^{hi}$ and HLA-DR negative, $CD123^{hi}$ and CD38 positive, $CD11b^{+}$, and not $CD34^{+}CD38^{lo}$.

**Platelets** were defined as meeting all of the following criteria: $CD321^{hi}$ and $CD45^{lo}$, $CD321^{hi}$ and DNA intercalator low, and negative for CD33.

**Immunophenotypically aberrant myeloid cells** were defined through removal of non-monocyte and non-granulocyte lineage cells. This population was created from cells that feel with all of the following gates: 1. CD3 negative, 2. Not $CD45^{hi}$ and $CD7^{hi}$ (i.e. Not NK cells; note that this gate was slightly adjust for a few samples to separate NK cells from myeloid cell aberrantly expressing high levels of CD7); 3. Not brightly CD38 positive (i.e. Not plasma cells); 4. Not CD321 bright and DNA$^{low}$ (i.e. Not platelets); 5. Not CD19$^{+}$ or CD20$^{+}$ (i.e. Not B cells); 6. Not CD71$^{hi}$ or CD235$^{hi}$ (i.e. Not committed erythroid progenitors; these cells were confirmed to be CD45low to ensure activated cells of other lineages were not excluded by the CD71 gate); 7. Not CD123$^{hi}$ (i.e. Not pDCs or basophils; boundaries of this gate were established by back-gating on basophils, pDCs and myeloid blasts with aberrant CD123 expression to ensure that boundaries of this single gate excluded the former populations without excluding the latter). Once these other lineages were excluded the aberrant cell population was defined by the simultaneous absence of HLA-DR, absence of CD15 and CD14, and the positive expression of CD11b. Cells meeting these criteria were very rare in normal the normal bone marrow samples and in patients with lower risk disease (see Supplemental Figure 12).

To perform comparisons to the LeukemiaNet MDS analysis method, several modifications were made to the LeukemiaNet criteria published in Della Porta, et al.. Specifically, B cell progenitor frequency was measured as the combined frequency of pre-B cells (CD10+CD19+CD20lo) and CD38+ mature B cells (CD19+CD20+CD38+); myeloblast cluster size was determined based on immunophenotypic criteria for myeloblasts (Supplemental Figure 1); and instead of using the granulocyte to lymphocyte SSC ratio, abnormal granulocyte development was assessed by the presence of immunophenotypic aberrancy in the mature granulocyte population. Values for each approximated parameter outside the absolute variance of the normal samples were considered abnormal.

SPADE analysis was performed as previously described [7], clustering markers are indicated in Supplementary Table 1. Clusters were manually grouped and annotated into immunophenotypic populations based on examination of relevant biaxial plots (e.g., CD3 vs. CD45) of the cell events in each cluster by utilizing information from previous reports [6,8, 9]. ViSNE analysis was performed in two t-SNE dimensions using the viSNE analysis tool in Cytobank as described previously. [10] Data files were sampled to ≤5,000 events each, and the surface markers used for the analysis are shown in Supplementary Table 1. K-means binning analysis of $CD34^{+}CD38^{low}$ cells (Supplemental Figure 7) was performed utilizing the same sampled cells and same surface markers employed in the viSNE analysis. Binning analysis of total cell populations (Supplemental Figure 9) was performed similarly using all cells and the same surface markers. Both analyses were performed using the K-mediods algorithm with K=100. Cluster frequencies per sample were then extracted and the ASINH Z-score was calculated and z-score vectors of all samples were clustered using average linkage hierarchical clustering with Euclidean distance (Supplemental Figures 7 and 9). X-shift clustering was performed in accordance with the previously published methods [11]; briefly, 10,000 cells from each sample were sampled from each MDS or controls sample and all sampled events were pooled into a single X-shift analysis to generate minimum spanning trees for each MDS and control sample.
