## Supplemental Figure 1 for "Profiling Myelodysplastic Syndromes by Mass Cytometry Demonstrates Abnormal Progenitor Cell Phenotype and Differentiation"

Supplemental Figure 1
Gating Hierarchy (Healthy Donor #6)

$CD34^{+}CD38^{low}$

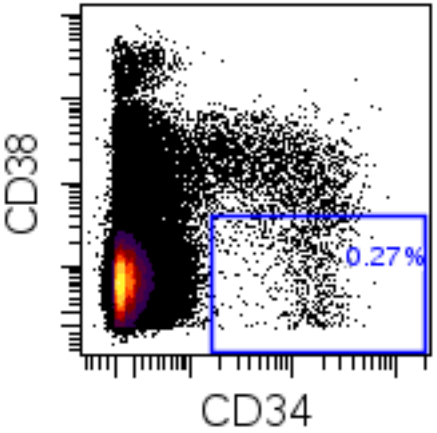


**Supplemental Figure 1:** Gating strategy is shown for each population. Cell events for normal sample #6 are shown. Gates for all samples were applied based on the gating of the normal samples, In almost all cases, the exact same gate boundaries defined by the normal samples was used for all MDS samples (without regard to the cell distribution), in rare cases, minimal adjustments were made to separate NK cells from MDS cells with bright aberrant CD7 expression. For mature cells of each lineage, a lineage population was first defined (blue box) and then each stage of maturation was sub-gated as shown. Some populations required multiple Boolean gates to achieve a pure cell population. In these cases, only events falling within all of the gates were considered to be in the population. For the CMP and GMP populations, CD123 signal (shown in brackets) did not allow for confident resolution of these populations from within the lineage-negative $CD34^{+}CD38^{+}$ cells.

Lineage negative

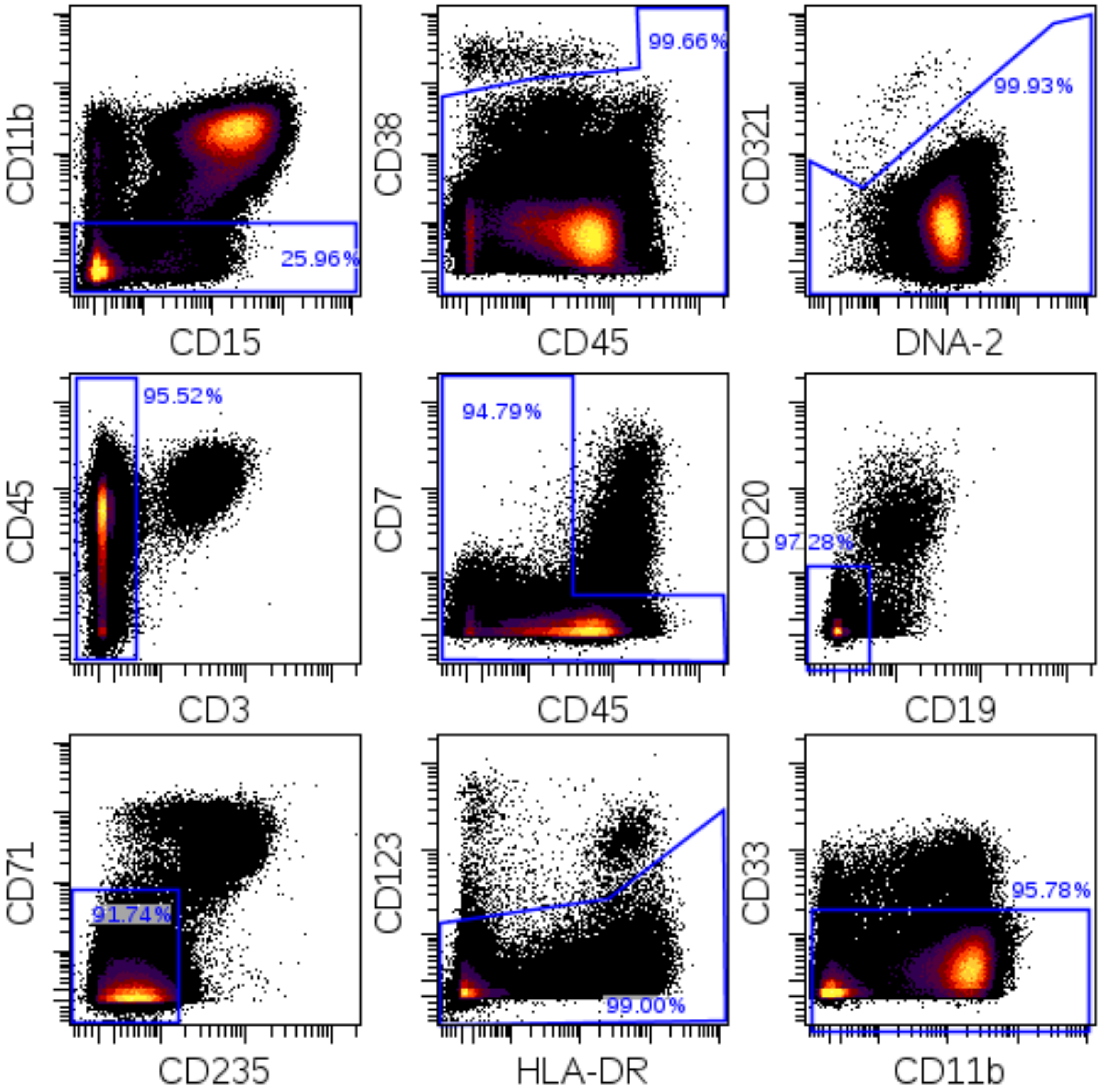

CD45RAnegCD34$^{+}$CD38$^{low}$ (also lineage negative)

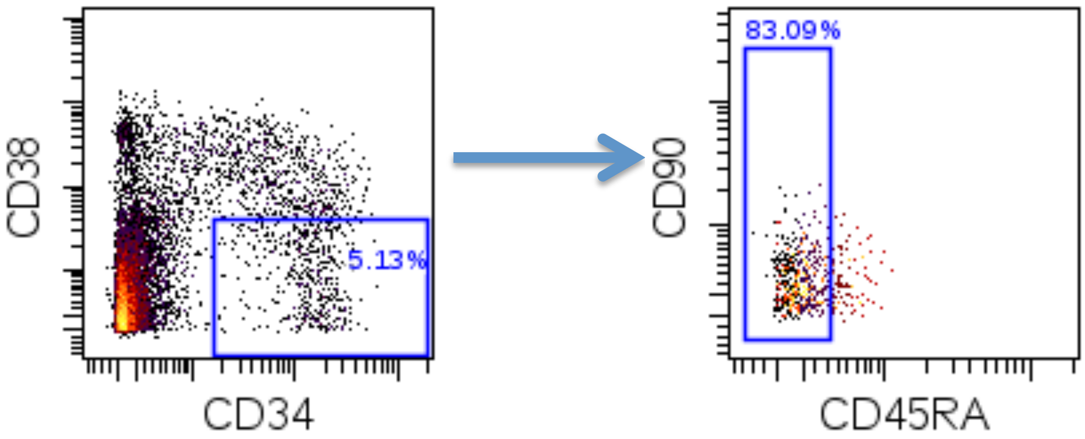


CD33negCD34$^{+}$CD38$^{low}$ (also lineage negative)

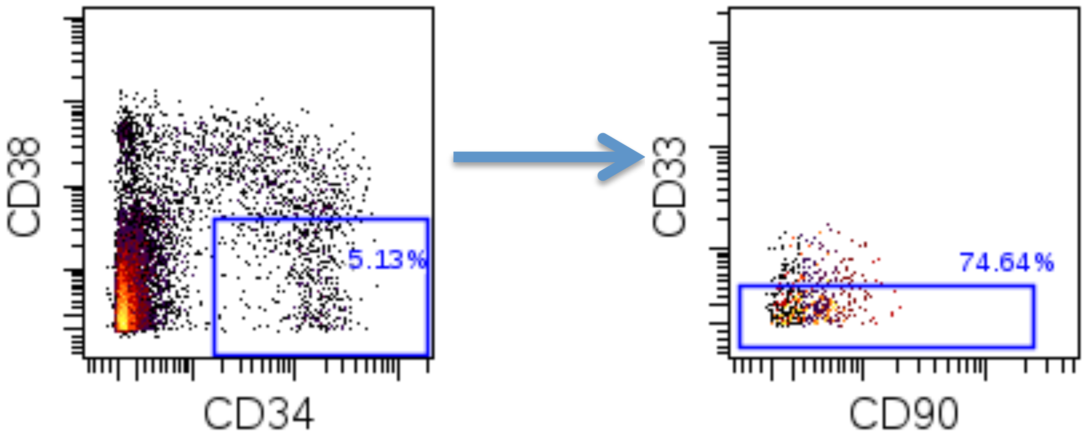

HSC-CD90hiCD33neg (also lineage negative)

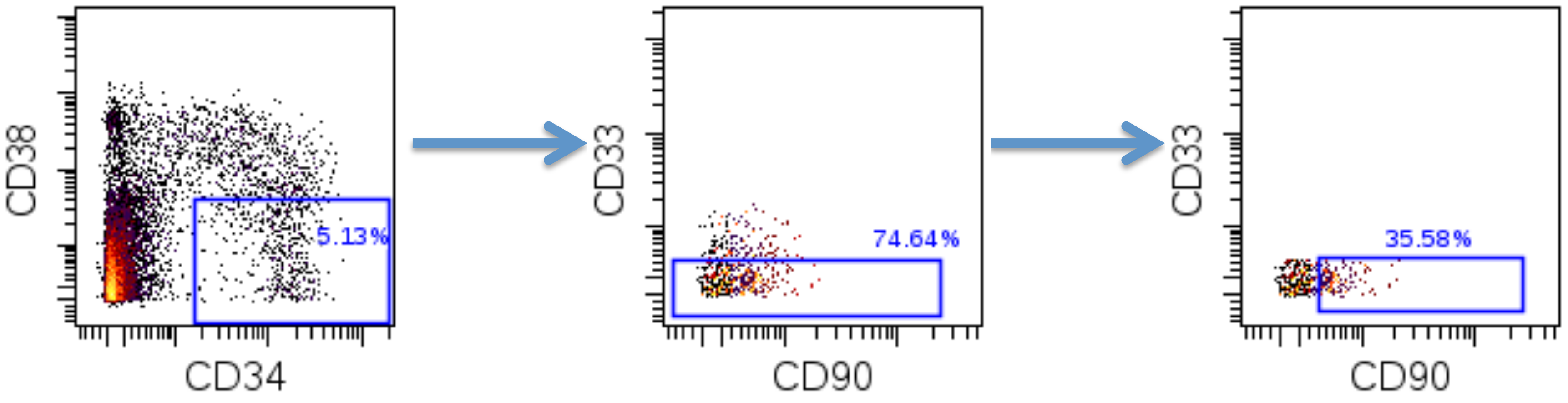


MPP-CD33negCD34$^{+}$CD38$^{low}$ (also lineage negative)

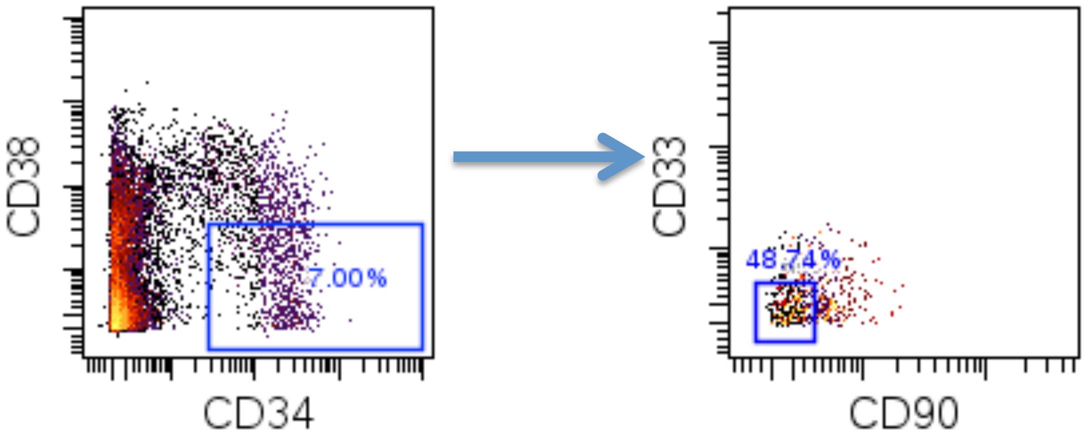

CD33+ MPP - CD34$^{+}$CD38$^{low}$ (also lineage negative)

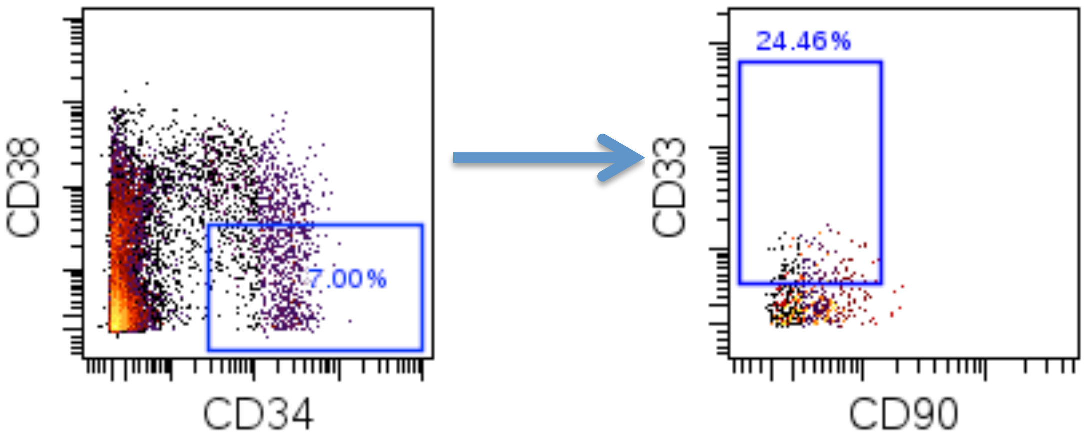


CMP/GMP-CD34$^{+}$CD38$^{+}$ (also lineage negative)

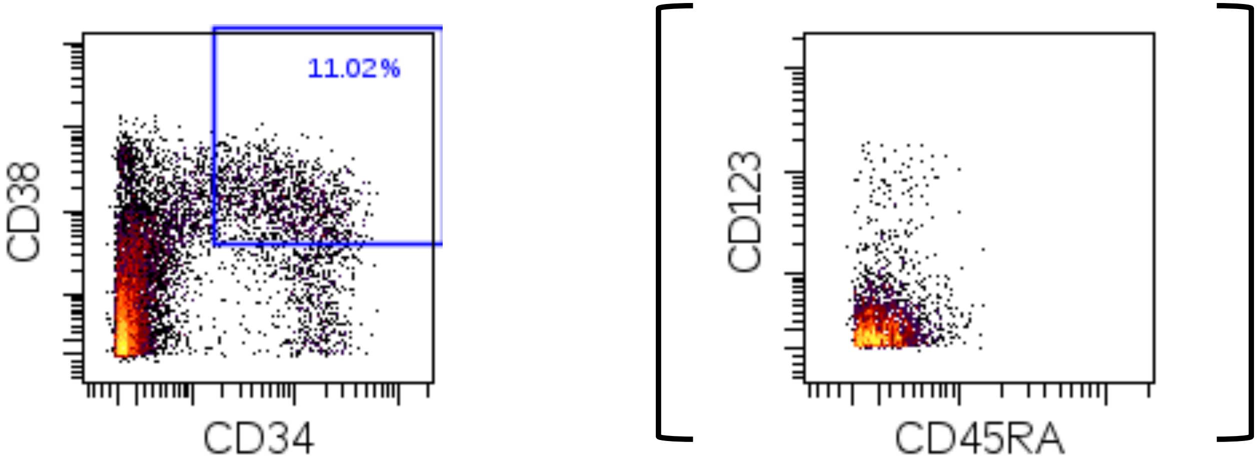

CD34$^{neg}$CD38$^{+}$ Undifferentiated (also lineage negative)

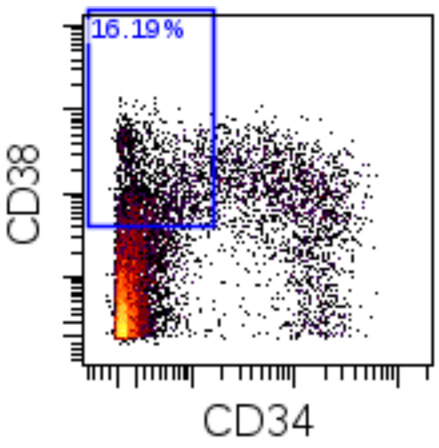


CD34$^{neg}$CD38$^{neg}$ Undifferentiated (also lineage negative)

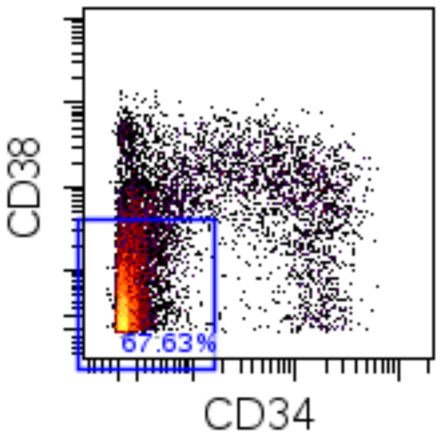

Myelo/Monoblast

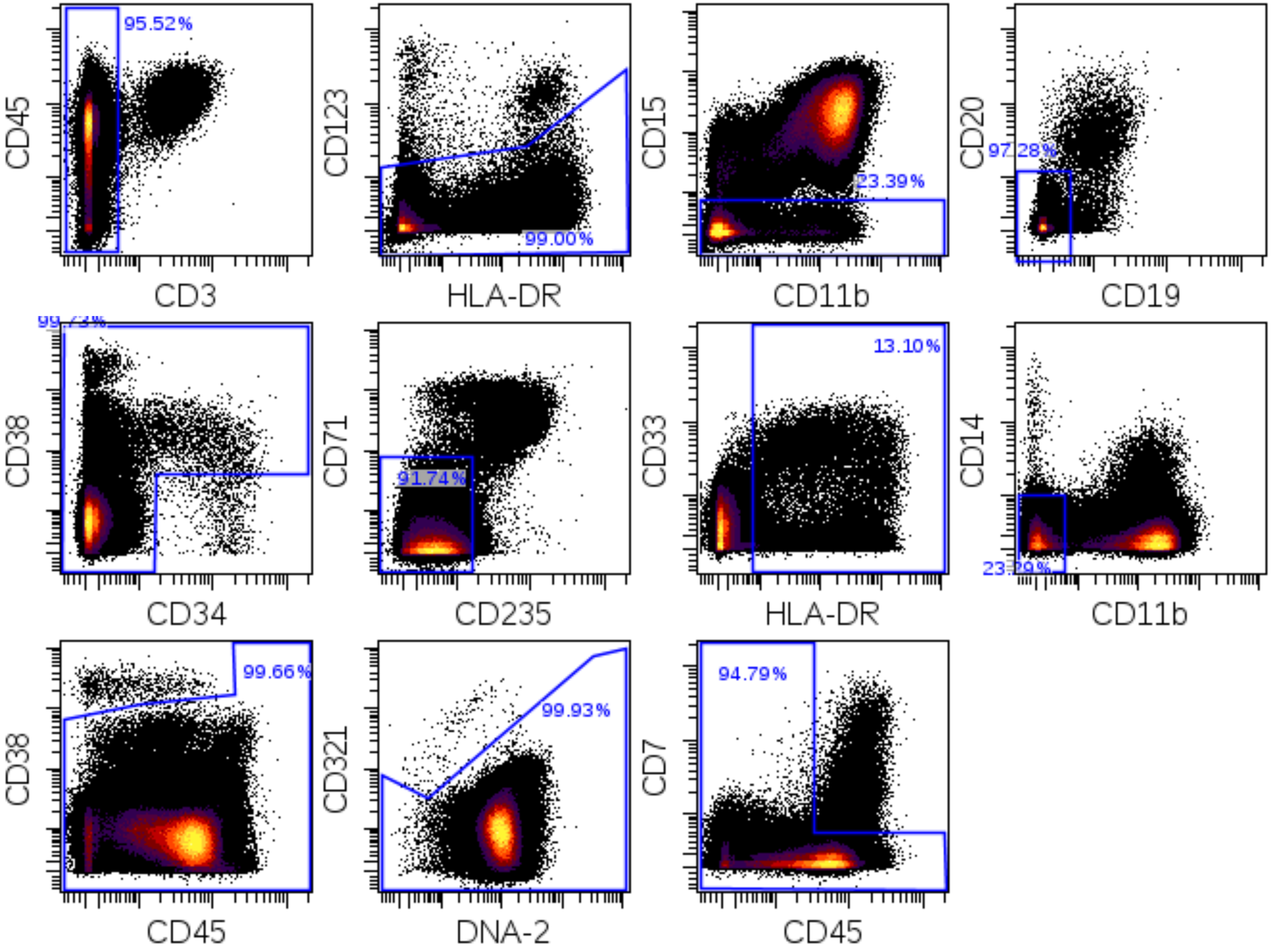

Monocyte Lineage

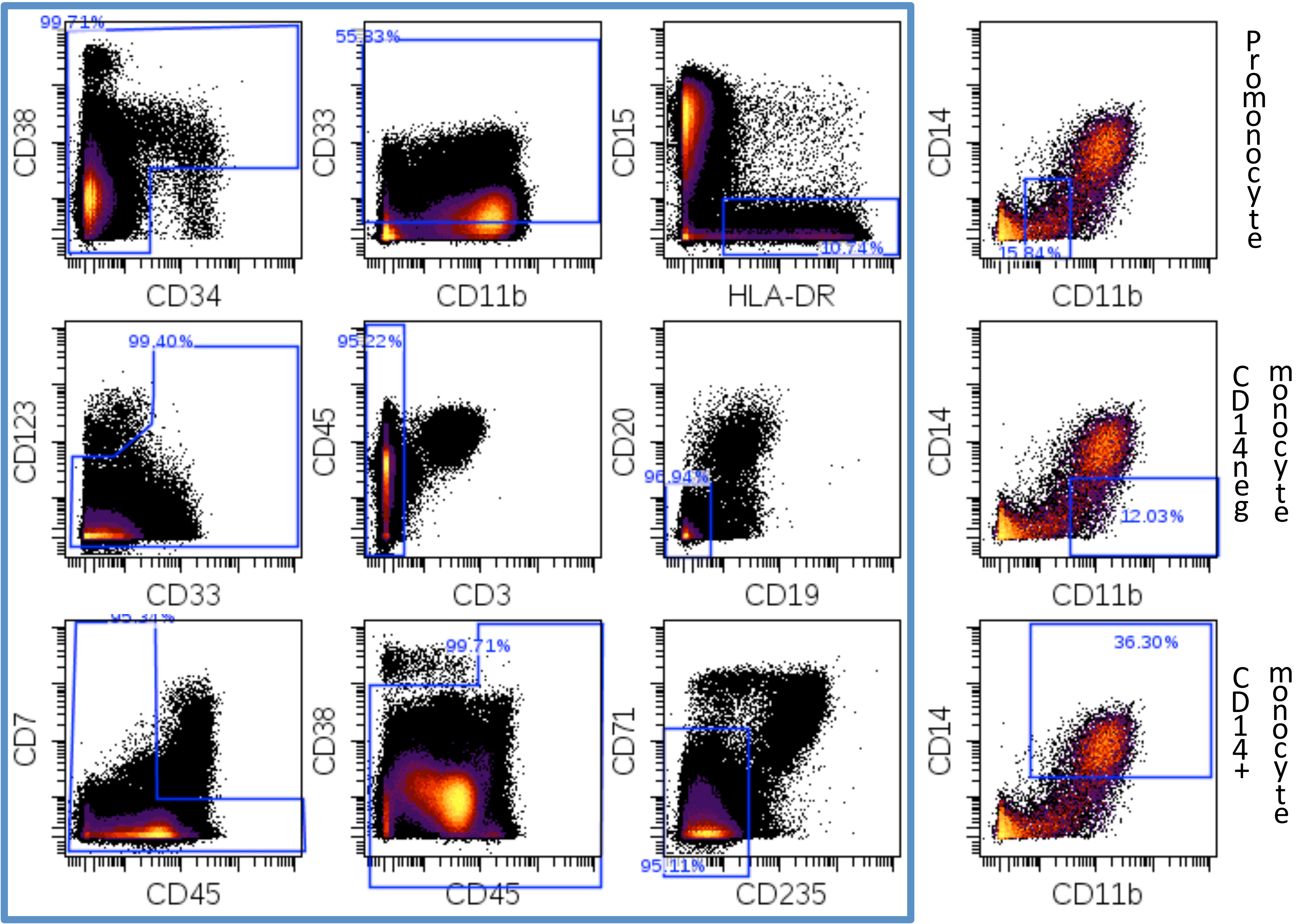

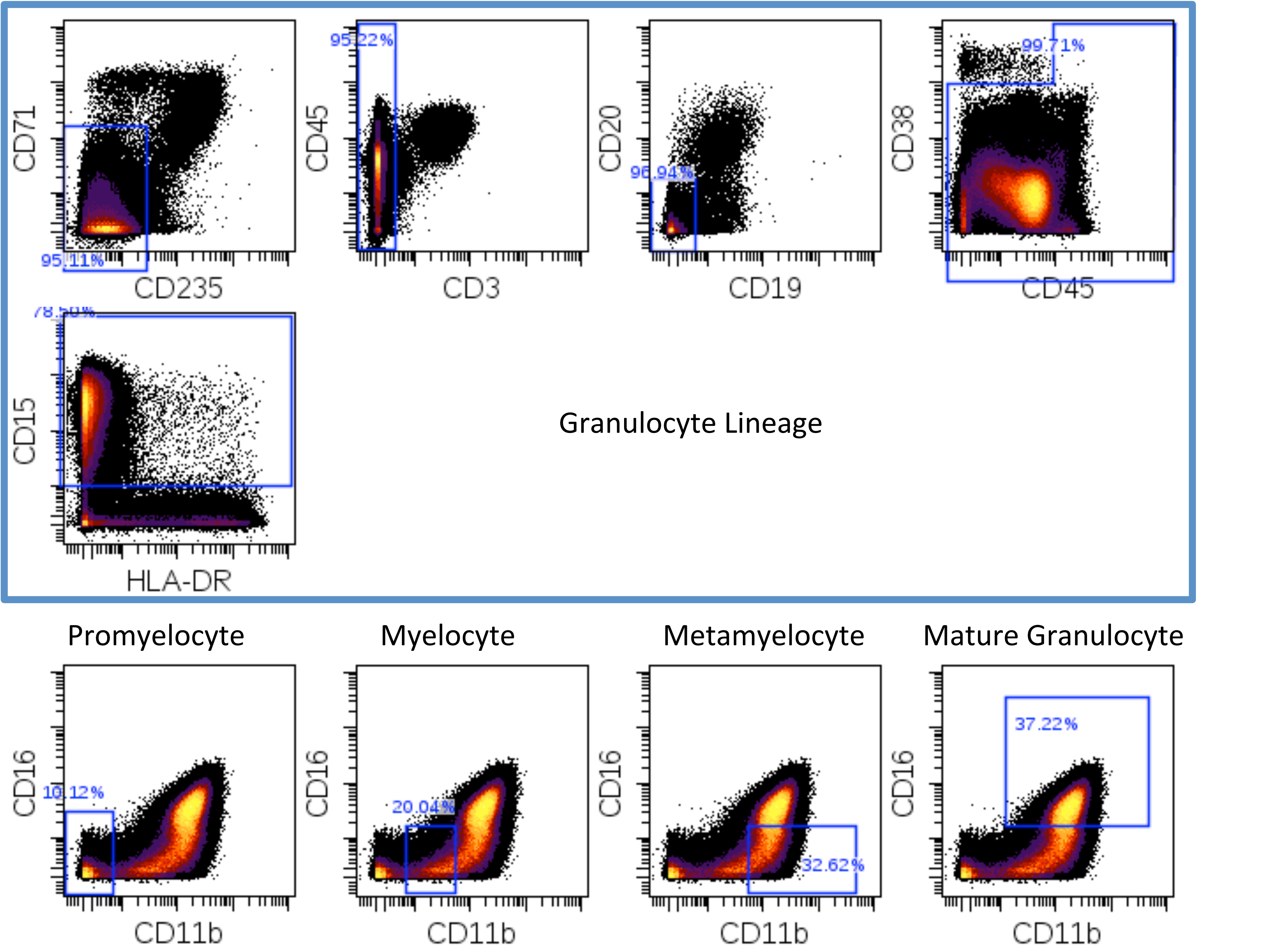
CD71
CD235
95.11%
CD45
CD3
95.22%
CD20
CD19
96.94%
CD38
CD45
99.71%
CD15
HLA-DR
Granulocyte Lineage
Promyelocyte
Myelocyte
Metamyelocyte
Mature Granulocyte
CD16
CD11b
10.12%
20.04%
32.62%
37.22%

Erythroid Lineage

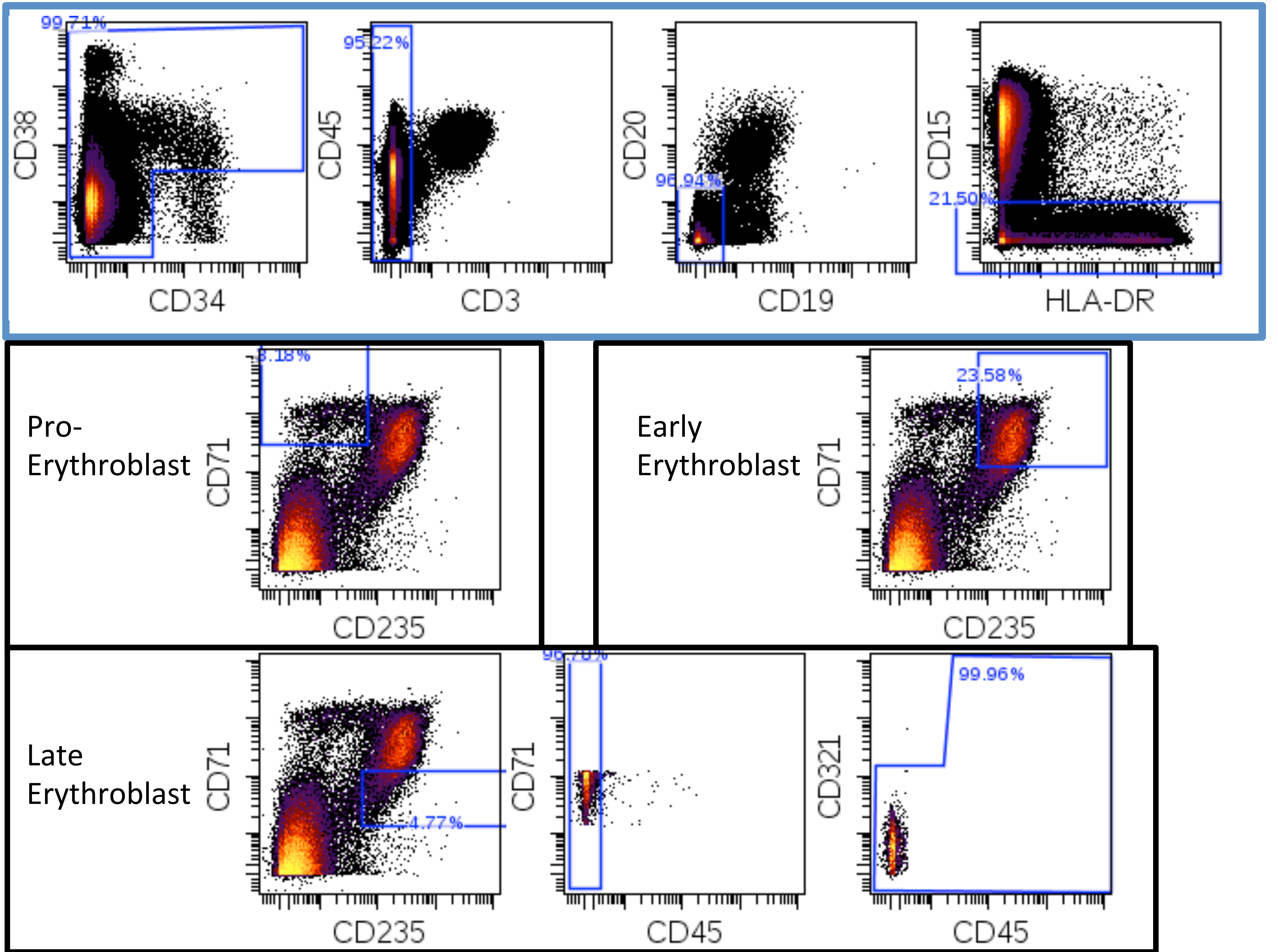

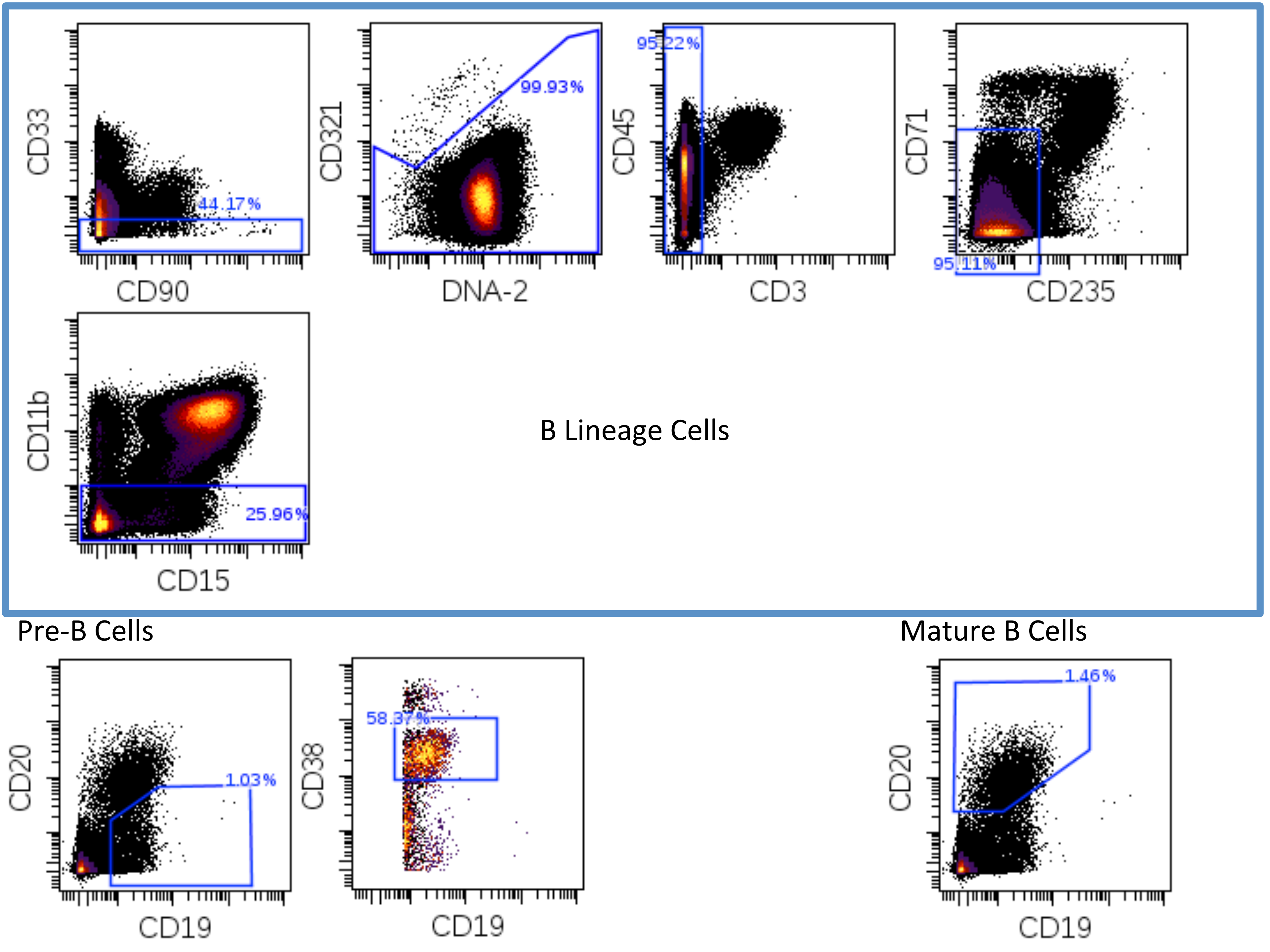
CD33
CD90
44.17%
CD321
DNA-2
99.93%
CD45
CD3
95.22%
CD71
CD235
95.11%
CD11b
CD15
25.96%
B Lineage Cells
Pre-B Cells
CD20
CD19
1.03%
CD38
CD19
58.37%
Mature B Cells
CD20
CD19
1.46%

### Plasma Cells

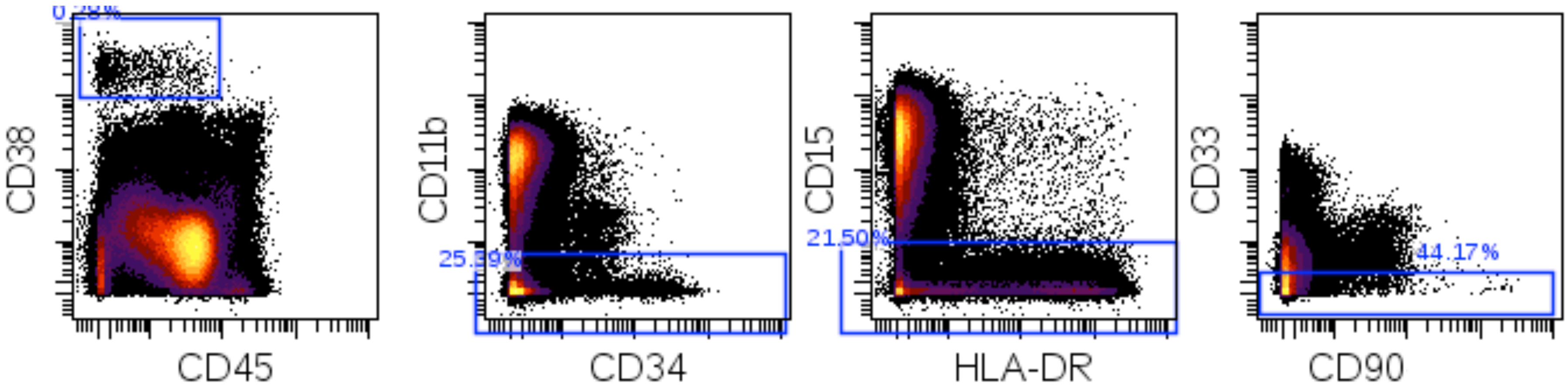


### T Cells

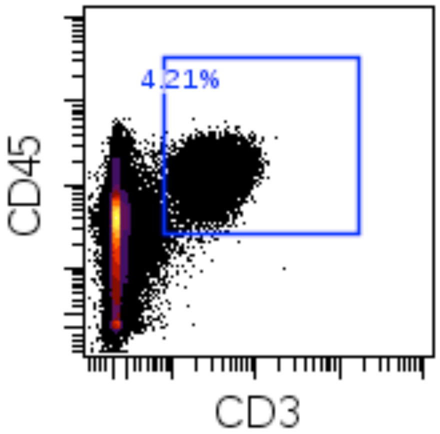


### NK Cell

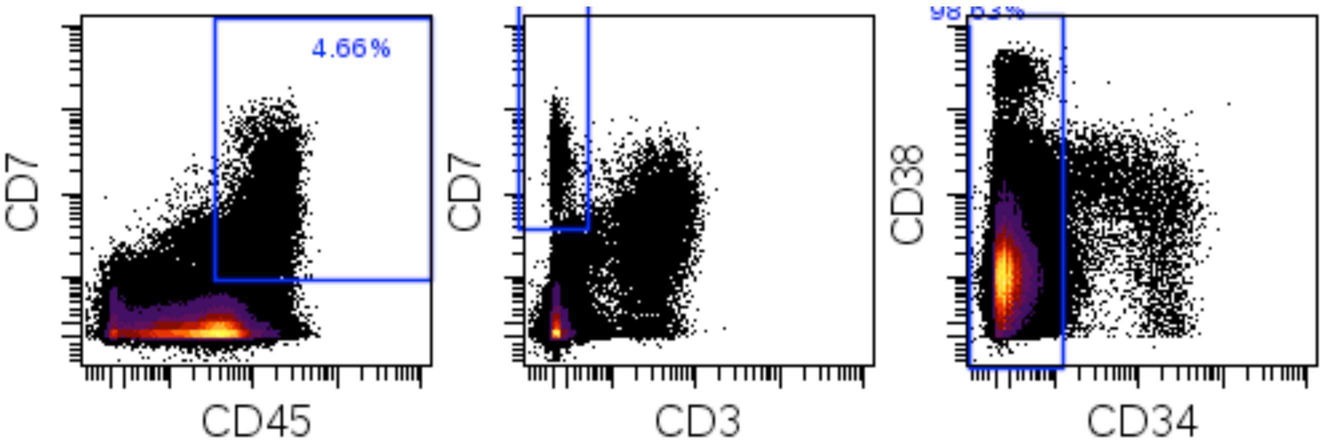

pDCs

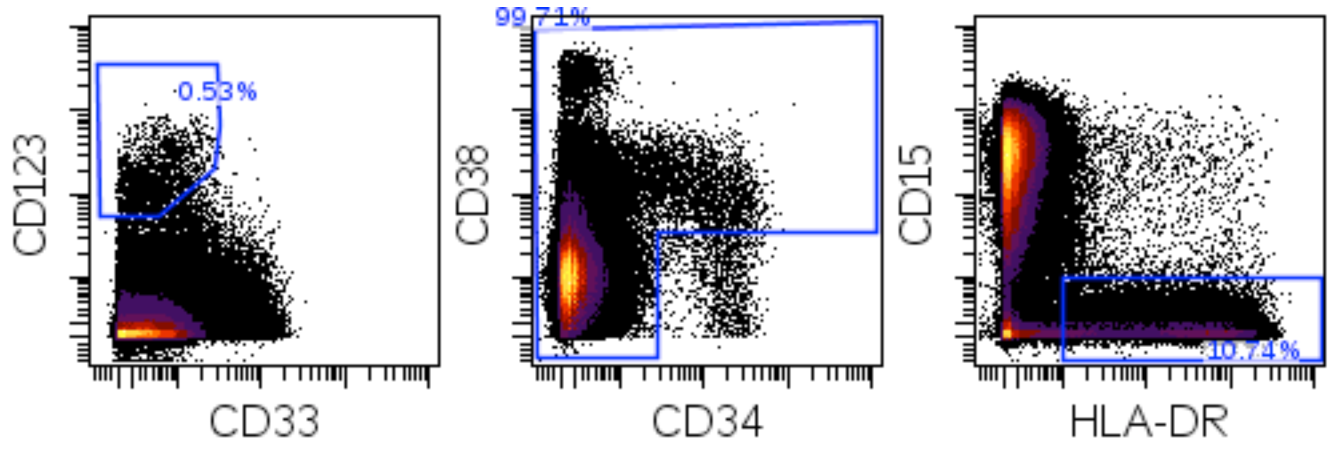


Basophils

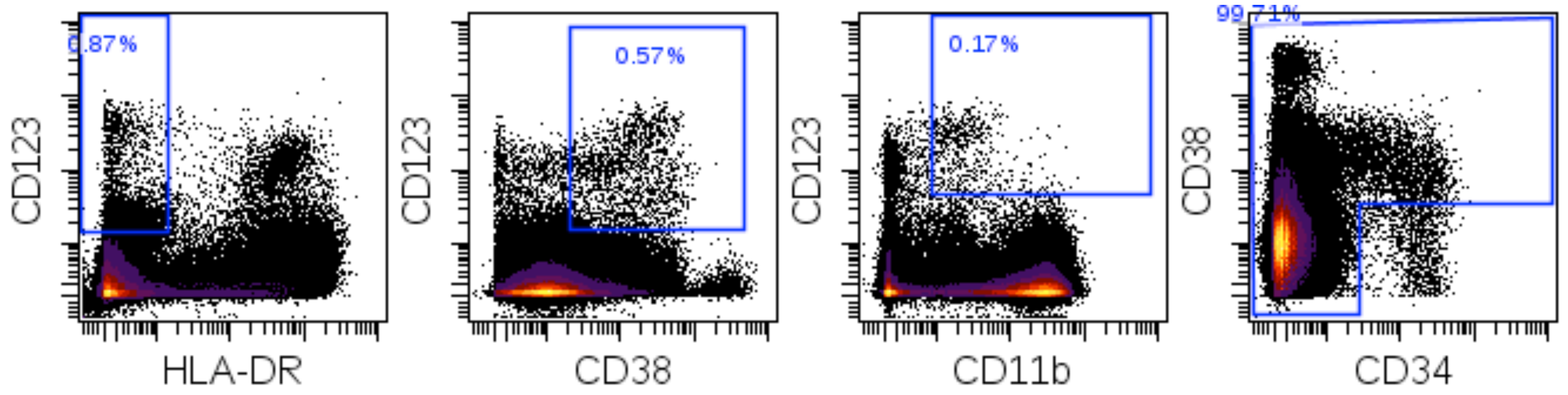

### Platelets

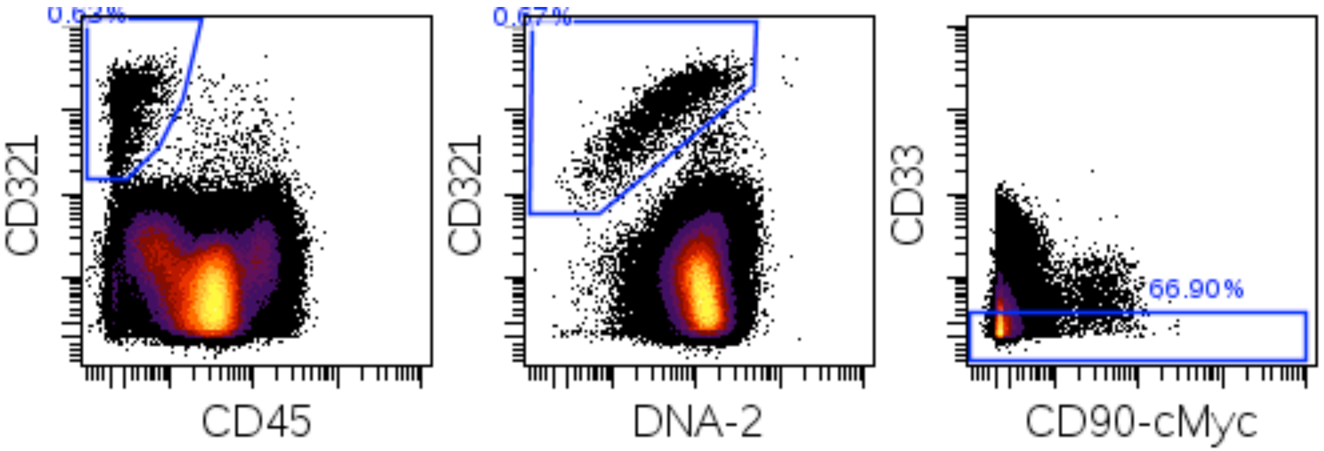

### Immunophenotypically Aberrant Myeloid Cells (IAMCs)

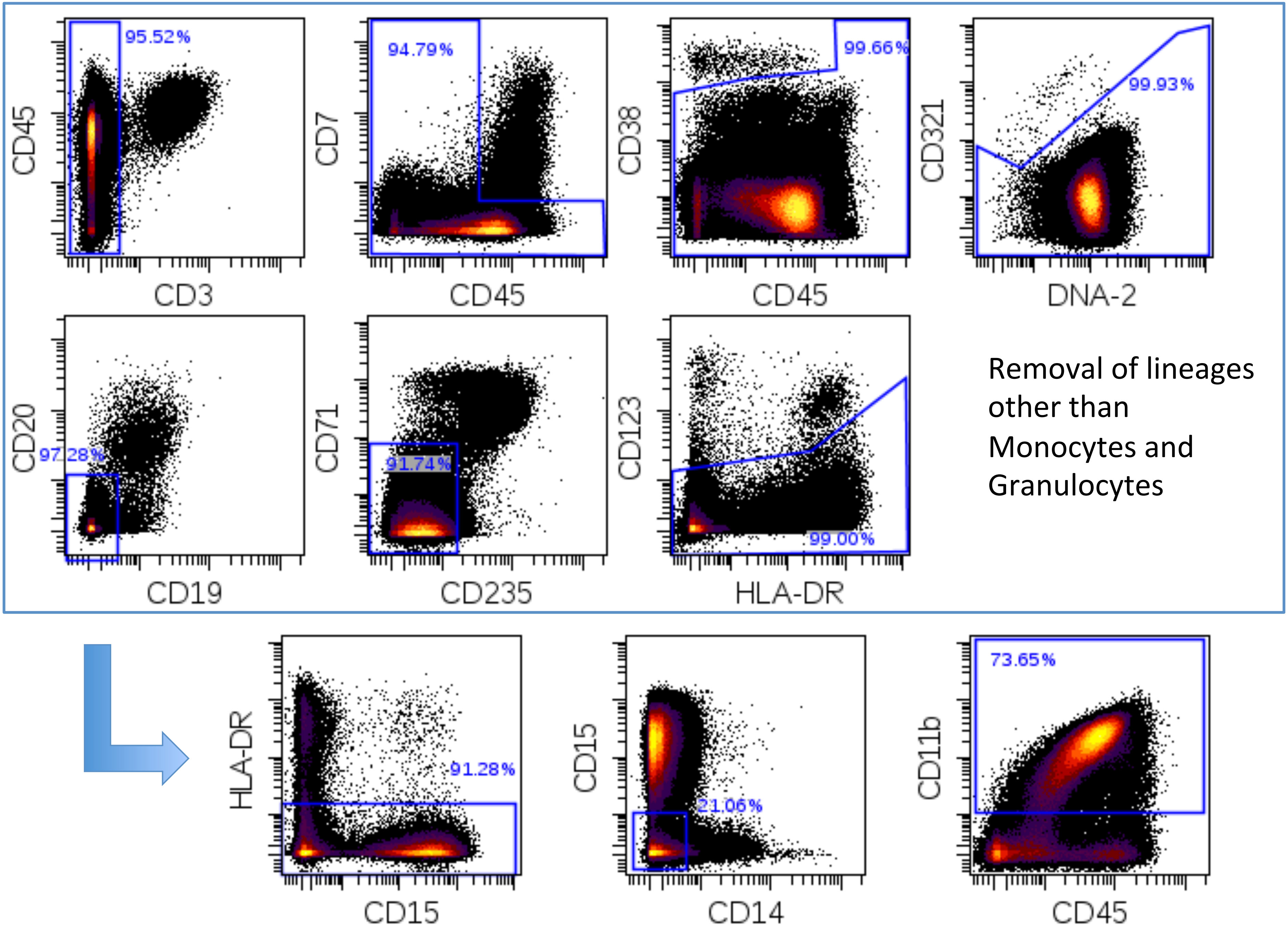
