## Supplemental Tables for "Profiling Myelodysplastic Syndromes by Mass Cytometry Demonstrates Abnormal Progenitor Cell Phenotype and Differentiation"

**Supplemental Table1:** Antibodies used in this study. The staining panel (A, B, or both) is indicated for each. The "Analyses" column indicates whether the marker was used for each dimensionality reduction analysis (S, SPADE analysis; V, viSNE analysis; B, multidimensional binning; X, X-shift clustering). Only markers measured in both panels had sufficient normal replicates for calculation of CVs. Calreticulin and cleaved-PARP did not have sufficient signal to calculate CV.

| Antigen | Conjugate | Clone | Concentration | Manufacturer | Panel | CV (%) | Analyses |
|---|---|---|---|---|---|---|---|
| CD3 | In-113 | UCHT1 | 2 µg/mL | Biolegend | A & B | 13.2 | S, V, B, X |
| CD45 | In-115 | Hl30 | 2 µg/mL | Biolegend | A & B | 11.0 | S, V, B, X |
| CD45RA | La-139 | Hl100 | 1.5 µg/mL | Biolegend | A & B | 18.8 | V, B, X |
| CD133 | Pr-141 | AC133 | 3 µg/mL | Militeney | A | N/A | |
| cleaved-Caspase3 | Pr-141 | C92-605 | 3 µg/mL | BD bioscience | B | N/A | Intra-cellular |
| CD7 | Nd-142 | M-T701 | 2 µg/mL | BD Biosciences | A & B | 7.1 | S, V, B, X |
| CD71 | Nd-143 | R17217 | 2 µg/mL | eBiosciences | A & B | 18.2 | S, V, B, X |
| CD235 | Nd-144 | HIR2 | 6 µg/mL | Biolegend | A & B | 24.4 | S, V, B, X |
| CD47 | Nd-145 | B6H12 | 1.5 µg/mL | BD Biosciences | A & B | 13.2 | V, B, X |
| CD8 | Nd-146 | RPA-T8 | 1 µg/mL | Biolegend | A & B | 4.6 | S, V, B, X |
| CD56 | Sm-147 | NCAM16.2 | 1.5 µg/mL | BD Biosciences | A & B | 4.6 | V, B, X |
| CD34 | Nd-148 | 8G12 | 3 µg/mL | BD Biosciences | A & B | 6.5 | S, V, B, X |
| CD90 | Sm-149 | 5E10 | 3 µg/mL | Biolegend | A & B | 16.7 | V, B, X |
| CD117 | Nd-150 | 104D2 | 1 µg/mL | Biolegend | A & B | 17.5 | V, B, X |
| CD123 | Eu-151 | 6H6 | 1 µL per 100 µL | DVS | A & B | 8.9 | S, V, B, X |
| CD33 | Sm-152 | P67.6 | 1.5 µg/mL | BD Biosciences | A & B | 6.3 | S, V, B, X |
| HLA-DR | Eu-153 | L243 | 2 µg/mL | Biolegend | A & B | 10.5 | S, V, B, X |
| CD64 | Sm-154 | 10.1 | 2 µg/mL | Biolegend | A & B | 4.9 | V, B, X |
| CD44 | Gd-156 | G44-26 | 1.5 µg/mL | BD Biosciences | A & B | 17.8 | V, B, X |
| Ki-67 | Gd-158 | SolA15 | 1 µg/mL | eBiosciences | A & B | 13.3 | Intra-cellular |
| CD38 | Tb-159 | HIT2 | 1 µg/mL | Biolegend | A & B | 9.0 | S, V, B, X |
| CD14 | Gd-160 | M5E2 | 2 µg/mL | Biolegend | A & B | 16.7 | S, V, B, X |
| CD16 | Dy-161 | 3G8 | 2 µg/mL | Biolegend | A & B | 15.5 | S, V, B, X |
| CD11b | Dy-162 | ICRF44 | 2 µg/mL | Biolegend | A & B | 9.7 | S, V, B, X |
| CD15 | Dy-164 | W6D3 | 3 µg/mL | Biolegend | A & B | 20.6 | S, V, B, X |
| Calreticulin | Ho-165 | FMC75 | 4 µg/mL | Thermo Fisher | A & B | N/A | |
| CD321 | Er-166 | WK9 | 3 µg/mL | eBiosciences | A & B | 10.7 | S, V, B, X |
| CD99 | Er-167 | HCD99 | 0.5 µg/mL | Biolegend | A & B | 10.8 | V, B, X |
| CD13 | Er-168 | L138 | 2 µg/mL | BD Biosciences | A & B | 6.6 | V, B, X |
| cleaved-PARP(D214) | Yb-171 | F21-852 | 1 µg/mL | BD Biosciences | A & B | N/A | Intra-cellular |
| CD10 | Yb-172 | Hl10a | 3 µg/mL | Biolegend | A & B | 7.8 | S, V, B, X |
| CD19 | Yb-173 | H1B19 | 2 µg/mL | BD Biosciences | A & B | 7.5 | S, V, B, X |
| CD20 | Yb-174 | 2H7 | 2 µg/mL | Biolegend | A & B | 14.6 | S, V, B, X |
| CXCR4 | Yb-175 | 12G5 | 1uL per test | DVS | A & B | 14.5 | V, B, X |
| CD69 | Yb-176 | FN50 | 1 µg/mL | BD Biosciences | A & B | 7.6 | V, B, X |

**Supplemental Table 2**: Clinical characteristics of MDS patients tested in this study. Survival time is from time of sample collection. "Alive" indicates patients alive at last follow-up (average follow of 53 months). Asterisks indicate serial samples from the same patient. "HSCT" denotes patients who received a hematopoietic stem cell transplant. "N/A" indicates follow not available.

| Patient | Sex | Age | Cytogenetics | FAB | WHO | IPSS | Blast % | Survival (mo.) |
|---|---|---|---|---|---|---|---|---|
| **IPSS High Risk** | | | | | | | | |
| MDS4 | M | 73 | del5q, del8q, Trisomy8 | RAEB-T | AML | High/3.5 | 26% | 14 |
| MDS15 | F | 71 | Monosomy 7 | AML (from CMML-2) | AML (from CMML-2) | High | 32% | 2.6 |
| MDS17 | M | 69 | Normal | RAEB-T | AML | High | 25% | 28.5 |
| **IPSS Intermediate-2 Risk** | | | | | | | | |
| MDS1 | M | 80 | Normal | RAEB | RAEB-2 | Int-2/2 | 16% | 12.9 |
| MDS2 | M | 80 | Normal | RAEB | RAEB-2 | INT-2/2.0 | 15% | 1.5 |
| MDS3* | M | 74 | Trisomy 8, Trisomy 9 | RAEB | RAEB-1 | INT-2/1.5 | 10-18% | 18 |
| MDS13* | M | 74 | Trisomy 8, Trisomy 9 | RAEB | RAEB-2 | INT-2/1.5 | 10.2% | 16 |
| MDS16 | M | 67 | Normal | RAEB | RAEB-2 | INT-2/2.0 | 15-20% | 5.8, HSCT |
| MDS21* | M | 75 | Trisomy 8, Trisomy 9 | RAEB | RAEB-2 | Int2/1.5 | 6% (following treatment) | 9.2 |
| MDS26 | F | 65 | del5q and del13q | MDS/MPN | MDS NOS | INT-2/1.5 | 4.70% | Alive |
| MDS27 | M | 69 | Normal (-Y) | RAEB | RAEB-2 | INT-2 | 2% (following treatment) | 10.5 |
| **IPSS Intermediate-1 Risk** | | | | | | | | |
| MDS5 | M | 67 | Normal | RA | RCMD | INT-1/1.0 | <5% | Alive; HSCT |
| MDS6 | M | 52 | Normal | RAEB | RAEB-1 | INT-1/1.0 | 5-7% | Alive |
| MDS8 | F | 56 | Normal | RARS | RCMD-RS | INT-1/0.5 | <5% | 39.5 |
| MDS12 | M | 65 | Normal | RA | RCMD | INT-1/0.5 | <5% | Alive |
| MDS14 | F | 58 | Normal (Trisomy 15 sub-clone) | RA | RCMD | INT-1/1.0 | <5% | Alive |
| MDS23 | M | 78 | Normal (-Y) | RA | RCMD | Int-1/0.5 | <5% | Alive |
| **IPSS Low Risk** | | | | | | | | |
| MDS9 | M | 69 | Normal (-Y) | RA | RCMD | LOW/0 (post-tx) | 4.4% | 40.5, HSCT |
| MDS11 | M | 29 | Normal | RA | RA | Low | 0% | Alive |
| MDS19 | M | 81 | Normal | RARS | RCMD-RS | LOW/0 | <5% | Alive |
| MDS20 | F | 83 | Normal | RA | RCMD | LOW/0 | <5% | 29.4 |
| MDS25 | F | 66 | Normal | RA | RCMD | LOW | <5% | Alive |
| MDS28 | M | 78 | Normal | CMML | CMML-1 | LOW/0 | <5% | Alive |
| **Idiopathic Cytopenia of Undetermined Significance** | | | | | | | | |
| ICUS18 | M | 71 | Normal | N/A | N/A | N/A | <5% | N/A |
| ICUS22 | M | 59 | Normal | N/A | N/A | N/A | 0% | N/A |
| ICUS24 | F | 26 | Normal | N/A | N/A | N/A | <5% | N/A |

**Supplemental Table 3**: Median expression of each surface marker on each cell population.

**Supplemental Table 3**: Median expression of each surface marker on each cell population.
The expression level of each marker is shown for each gated cell populaiton. Expression level is shown in median ion counts detected. Each cell is colored from green (lowest expression) to red (highest expression).
Samples 3, 5, and 21 were run twice due to lower numbers of cell events in the first run.
MDS samples 3, 13, and 21 come from the same patient tested on three seperate bone marrow aspirations each several months apart.
*Indicates samples from patients with CMML or an MDS/MPN overlap syndrome.

### CD45RA Median

| | | Ungated | $CD34^{+}CD38^{lo}$ | HSCs | MPP | CMP/GMP | Myelo/Mono-Blasts | ProMonocytes | CD14neg Monocytes | Mature Monocytes | ProMyelocytes | Myelocytes | Meta-Myelocytes | Mature Granulocytes |
|---|---|---|---|---|---|---|---|---|---|---|---|---|---|---|
| Run1_MDS17 | | 1.35 | 0.69 | 0.5 | 0.41 | 1.03 | 0.53 | 1.74 | 1.98 | 1.61 | 1.17 | 3 | 2.5 | 2.71 |
| Run2_MDS15* | AML/RAEB-T | 1.52 | 1.33 | 0.68 | 0.68 | 0.97 | 1.26 | 1.79 | 1.72 | 1.68 | 1.47 | 1.63 | 1.53 | 1.34 |
| Run2_MDS4 | | 1.45 | 1.06 | 0.64 | 0.43 | 0.99 | 0.95 | 1.47 | 1.95 | 3.11 | 1.68 | 1.51 | 1.52 | 1.65 |
| Run1_MDS21 | | 1.1 | 0.77 | 0.66 | 0.47 | 0.86 | 0.78 | 1.42 | 1.32 | 1.55 | 0.63 | 0.82 | 0.93 | 1.32 |
| Run1_MDS3 | | 1.19 | 0.79 | 0.59 | 0.39 | 0.88 | 0.85 | 1.8 | 1.66 | 2.58 | 0.88 | 1.17 | 1.28 | 1.77 |
| Run2_MDS13 | | 1.1 | 0.77 | 0.56 | 0.5 | 0.88 | 0.84 | 1.57 | 2.49 | 2.65 | 0.91 | 1.36 | 1.62 | 1.65 |
| Run2_MDS16 | | 0.98 | 0.82 | 0.58 | 0.55 | 0.8 | 1.55 | 1.57 | 1.48 | 1.47 | 0.79 | 0.64 | 0.69 | 0.73 |
| Run2_MDS1 | | 0.89 | 0.77 | 0.32 | 0.51 | 0.86 | 0.68 | 0.99 | 1.13 | 1.68 | 0.67 | 0.65 | 0.84 | 1.02 |
| Run2_MDS21 | Higher Risk | 0.87 | 0.65 | 0.43 | 0.44 | 0.75 | 0.67 | 0.96 | 1.09 | 1.01 | 0.81 | 0.96 | 0.94 | 0.95 |
| Run2_MDS26* | | 0.62 | 0.71 | 0.49 | 0.44 | 1.07 | 0.64 | 0.74 | 0.95 | 0.61 | 0.41 | 0.41 | 0.56 | 0.62 |
| Run2_MDS27 | | 1.28 | 0.95 | 0.66 | 0.59 | 0.92 | 1.14 | 1.62 | 1.79 | 1.68 | 0.88 | 0.83 | 0.84 | 1.01 |
| Run2_MDS2 | | 1.32 | 0.49 | 0.36 | 0.21 | 0.51 | 0.56 | 1.71 | 2.04 | 4.98 | 1.53 | 0.99 | 1.89 | 2.16 |
| Run2_MDS3 | | 1.27 | 0.68 | 0.56 | 0.47 | 0.81 | 0.8 | 1.88 | 2.35 | 1.92 | 1.48 | 2 | 2.04 | 2.35 |
| Run1_MDS23 | | 1.39 | 1.02 | 0.81 | 0.53 | 1.52 | 1.69 | 2.54 | 2.37 | 2.13 | 1.83 | 1.31 | 1.26 | 1.26 |
| Run1_MDS25 | | 0.9 | 1.82 | 0.04 | 0.39 | 0.6 | 1.41 | 1.1 | 0.83 | 0.8 | 0.86 | 0.7 | 0.74 | 0.82 |
| Run1_MDS5 | | 1.93 | 1.91 | 0.08 | 0.9 | 1.75 | 2.52 | 2.14 | 3.27 | 2.22 | 1.65 | 1.24 | 1.64 | 1.87 |
| Run1_MDS6 | | 1.87 | 1.24 | 0.47 | 0.75 | 1.34 | 1.79 | 2.06 | 2.13 | 2.17 | 1.76 | 1.48 | 1.75 | 1.72 |
| Run1_MDS8 | | 1.28 | 2.53 | 0.66 | 0.3 | 1.15 | 2.1 | 2.57 | 1.85 | 1.95 | 0.95 | 0.85 | 0.93 | 0.99 |
| Run1_MDS9 | | 1.34 | 1.2 | 0.77 | 0.65 | 1.52 | 1.64 | 2.64 | 1.7 | 1.02 | 1.06 | 0.89 | 0.98 | 1.06 |
| Run2_MDS12 | Lower Risk | 1.07 | 1.53 | 0.44 | 0.93 | 1.32 | 1.41 | 1.38 | 1.38 | 1.33 | 0.86 | 0.9 | 1.02 | 1.06 |
| Run2_MDS14 | | 1.03 | 1.94 | 0.58 | 2.13 | 0.19 | 1.71 | 1.75 | 1.73 | 1.81 | 0.94 | 0.85 | 0.91 | 0.86 |
| Run2_MDS19 | | 0.97 | 0.86 | 0.34 | 0.61 | 0.99 | 1.61 | 1.53 | 1.33 | 1.3 | 0.9 | 0.85 | 0.9 | 0.91 |
| Run2_MDS20 | | 1.71 | 1.07 | 1.07 | -0.1 | 0.88 | 1.56 | 1.7 | 1.53 | 1.71 | 1.52 | 1.56 | 1.56 | 1.68 |
| Run2_MDS28* | | 2.51 | 1.08 | 0.82 | 0.49 | 1.02 | 2.67 | 3.37 | 3.35 | 3.77 | 3.39 | 2.8 | 2.72 | 2.32 |
| Run2_MDS5 | | 1.75 | 2.98 | -0.2 | -0.15 | 1.15 | 2.21 | 1.7 | 2.98 | 2.46 | 1.24 | 1.27 | 1.37 | 1.73 |
| Run1_ICUS18 | | 1.31 | 1.54 | 0.87 | 0.85 | 1.3 | 2.18 | 1.76 | 1.57 | 1.44 | 1.22 | 0.94 | 1.02 | 1.09 |
| Run1_ICUS22 | ICUS | 1.09 | 1.04 | 0.73 | 0.63 | 1.21 | 1.7 | 1.68 | 1.25 | 1.21 | 1.21 | 0.98 | 0.96 | 0.98 |
| Run1_ICUS24 | | 1.15 | 1.09 | 0.88 | 0.57 | 1.35 | 1.6 | 1.75 | 1.45 | 1.44 | 1.19 | 0.93 | 0.95 | 0.93 |
| Run1_Nl-1 | | 1.07 | 0.94 | 0.66 | 0.58 | 1.16 | 1.85 | 1.76 | 1.59 | 1.35 | 1.07 | 0.81 | 0.82 | 0.85 |
| Run1_Nl-3 | | 1.34 | 1.2 | 0.53 | 0.65 | 1.07 | 1.98 | 4.14 | 4.79 | 3.14 | 1.04 | 0.91 | 1.18 | 1.2 |
| Run1_Nl-4 | | 1.42 | 1.65 | 0.63 | 0.77 | 1.2 | 2.06 | 4.95 | 5.27 | 3.42 | 1.06 | 0.97 | 1.22 | 1.26 |
| Run1_Nl-6 | | 1.31 | 0.98 | 0.52 | 0.53 | 1.13 | 1.92 | 4.51 | 4.78 | 3.69 | 1.09 | 0.96 | 1.19 | 1.19 |
| Run2_Nl-3 | Normal | 1.04 | 0.77 | 0.84 | 0.27 | 0.77 | 1.63 | 2.89 | 3.04 | 2.21 | 0.85 | 0.8 | 0.96 | 0.99 |
| Run2_Nl-4 | | 1.15 | 1.11 | 0.62 | 0.55 | 1.04 | 1.9 | 3.47 | 3.44 | 2.55 | 0.93 | 0.83 | 1.05 | 1.08 |
| Run2_Nl-5 | | 1.15 | 0.81 | 0.49 | 0.54 | 0.92 | 1.53 | 2.89 | 3.04 | 2.47 | 0.75 | 0.78 | 0.97 | 1.05 |
| Run2_Nl-6 | | 0.94 | 0.75 | 0.61 | 0.52 | 0.77 | 1.61 | 3.08 | 3.11 | 2.49 | 0.76 | 0.74 | 0.89 | 0.92 |

| | | ProErythro-blasts | Erythroblasts | Late Erythroblasts | Pre-B cells | Mature B cells | Plasma Cells | T cells | NK cells | pDCs | Basophils | Platelets | CD8+ T cells | CD8neg T cells |
|---|---|---|---|---|---|---|---|---|---|---|---|---|---|---|
| Run1_MDS17 | | 1.27 | 1.23 | 1.05 | 1.2 | 4.52 | 2.18 | 1.34 | 3.02 | 1.73 | 1.3 | 0.31 | 1.6 | 1.03 |
| Run2_MDS15* | AML/RAEB-T | 2.27 | 2.96 | 2.79 | 1.66 | 5.96 | 3.57 | 1.39 | 2.93 | 1.84 | 1.03 | 1.11 | 2.12 | 0.86 |
| Run2_MDS4 | | 2.11 | 3.81 | 1.19 | 1.23 | 1.81 | 3.43 | 1.05 | 1.63 | 1.27 | 1.01 | 0.34 | 1.33 | 0.86 |
| Run1_MDS21 | | 1.69 | 2.4 | 4.17 | 1.93 | 6.23 | 1.61 | 1.32 | 3.42 | 1.51 | 0.56 | 1.85 | 2.79 | 0.9 |
| Run1_MDS3 | | 1.29 | 1.7 | 2.09 | 2.53 | 6.53 | 2.72 | 1.53 | 4.12 | 2.56 | 1.01 | 0.98 | 3.09 | 1.06 |
| Run2_MDS13 | | 1.45 | 1.24 | 0.99 | 1.24 | 3.21 | 2.23 | 1.24 | 2.39 | 1.85 | 0.93 | 0.46 | 2.02 | 0.9 |
| Run2_MDS16 | | 1.28 | 2.12 | 1.21 | 1.38 | 2.53 | 1.46 | 1.37 | 3.88 | 1.81 | 0.58 | 0.87 | 2.47 | 1.17 |
| Run2_MDS1 | | 0.94 | 1.95 | 1.54 | 0.9 | 2.89 | 2.11 | 0.77 | 2.27 | 0.93 | 1.03 | 0.33 | 1.01 | 0.73 |
| Run2_MDS21 | Higher Risk | 1.01 | 1.02 | 0.92 | 1.31 | 3.55 | 0.8 | 1.03 | 1.87 | 0.77 | 0.58 | 0.87 | 1.83 | 0.77 |
| Run2_MDS26* | | 1.19 | 1.66 | 1.14 | 0.62 | 1.72 | 1.58 | 1.12 | 2.37 | 0.62 | 0.68 | 0.44 | 1.4 | 0.98 |
| Run2_MDS27 | | 1.39 | 2.53 | 0.91 | 1.04 | 3.17 | 2.44 | 0.85 | 3.23 | 3.08 | 1.26 | 1.44 | 1 | 0.67 |
| Run2_MDS2 | | 2.35 | 2.95 | 1.23 | 0.92 | 7.45 | -0.08 | 0.73 | 1.57 | 2.59 | 0.75 | 0.23 | 1.43 | 0.69 |
| Run2_MDS3 | | 0.97 | 1.01 | 0.73 | 1.58 | 3.78 | 1.1 | 1.14 | 2.57 | 1.94 | 1 | 0.69 | 2.11 | 0.84 |
| Run1_MDS23 | | 1.72 | 2.28 | 2.44 | 1.8 | 4.08 | 0.79 | 0.96 | 4.91 | 2.4 | 0.99 | 0.97 | 1.29 | 0.86 |
| Run1_MDS25 | | 2.39 | 3.33 | 2.42 | 1.51 | 6.51 | 2.83 | 1.45 | 4.6 | 2.92 | 0.74 | 0.65 | 1.95 | 1.03 |
| Run1_MDS5 | | 4.35 | 4.23 | 2.11 | 2.35 | 5.38 | 3.42 | 1.39 | 5.99 | 3.81 | 1.76 | 0.55 | 1.55 | 1.32 |
| Run1_MDS6 | | 3.65 | 4.19 | 2.81 | 2.5 | 5.72 | 3.64 | 1.73 | 5.66 | 3.08 | 1.7 | 2.13 | 1.96 | 1.54 |
| Run1_MDS8 | | 4.4 | 4.69 | 3.44 | 2.34 | 4.76 | 7.8 | 1.96 | 4.57 | 2.95 | 1.06 | 1.58 | 2.62 | 1.77 |
| Run1_MDS9 | | 3.18 | 4.34 | 2.97 | 1.96 | 4.24 | 2.27 | 1.4 | 5.44 | 2.94 | 1.53 | 0.76 | 2.15 | 1.05 |
| Run2_MDS12 | Lower Risk | 1.58 | 1.76 | 1.33 | 1.27 | 3.33 | 0.68 | 0.79 | 2.72 | 1.69 | 0.78 | 0.92 | 1.09 | 0.62 |
| Run2_MDS14 | | 1.89 | 2.36 | 1.52 | 1.21 | 2.83 | 1.71 | 0.89 | 3.97 | 1.54 | 0.79 | 0.94 | 1.48 | 0.69 |
| Run2_MDS19 | | 1.45 | 2.37 | 1.35 | 1.26 | 3.7 | 1.17 | 0.89 | 3.09 | 2.07 | 0.61 | 1.18 | 1.76 | 0.71 |
| Run2_MDS20 | | 1.85 | 2.16 | 2.24 | 1.88 | 4.65 | 2.04 | 1.83 | 2.63 | 2.32 | 1.2 | 0.69 | 2.02 | 1.81 |
| Run2_MDS28* | | 1.97 | 1.57 | 2.94 | 2.41 | 2.83 | 1.07 | 0.98 | 2.99 | 2.77 | 1.09 | 1.98 | 1.17 | 0.85 |
| Run2_MDS5 | | 4.13 | 2.9 | 2.15 | 2.62 | 2.94 | 3.35 | 1.12 | 4.02 | 1.48 | 1.36 | 0.75 | 1.25 | 1.04 |
| Run1_ICUS18 | | 2.38 | 1.94 | 1.65 | 2.09 | 4.59 | 1.61 | 1.55 | 5.1 | 4.14 | 0.58 | 0.97 | 2.05 | 1.44 |
| Run1_ICUS22 | ICUS | 2.39 | 2.71 | 1.7 | 1.55 | 4.17 | 1.19 | 1.03 | 4.2 | 2.85 | 1.31 | 0.6 | 1.32 | 0.93 |
| Run1_ICUS24 | | 2.48 | 2.35 | 2.65 | 1.47 | 3.63 | 1.15 | 1.19 | 4.49 | 2.31 | 0.92 | 2.37 | 1.53 | 0.97 |
| Run1_Nl-1 | | 2.2 | 2.35 | 1.91 | 1.45 | 3.38 | 1.53 | 1.98 | 5.89 | 3.16 | 1.8 | 3.38 | 2.82 | 1.65 |
| Run1_Nl-3 | | 2.6 | 1.91 | 1.81 | 1.78 | 2.72 | 1.55 | 1.48 | 5.43 | 2.96 | 1.01 | 0.38 | 2.14 | 1.14 |
| Run1_Nl-4 | | 2.71 | 2.11 | 1.73 | 2.22 | 4.07 | 2.46 | 1.8 | 5.71 | 2.55 | 1.01 | 0.25 | 2.56 | 1.3 |
| Run1_Nl-6 | | 2.43 | 2.01 | 1.64 | 1.52 | 2.87 | 1.88 | 1.15 | 4.17 | 2.7 | 0.88 | 0.3 | 1.55 | 1.04 |
| Run2_Nl-3 | Normal | 1.48 | 1.39 | 1.26 | 1.26 | 1.92 | 1.26 | 1.08 | 3.26 | 1.88 | 0.7 | 0.44 | 1.6 | 0.81 |
| Run2_Nl-4 | | 1.65 | 1.5 | 1.37 | 1.63 | 2.86 | 1.8 | 1.33 | 3.4 | 1.83 | 0.67 | 0.2 | 1.83 | 1 |
| Run2_Nl-5 | | 1.66 | 1.48 | 1.26 | 1.06 | 2.26 | 1.22 | 1.42 | 4.19 | 1.85 | 0.68 | 0.28 | 2.06 | 0.98 |
| Run2_Nl-6 | | 1.32 | 1.22 | 0.99 | 1.01 | 1.84 | 1.63 | 0.89 | 2.55 | 1.73 | 0.66 | 0.36 | 1.16 | 0.79 |

**Supplemental Table 3**: Median expression of each surface marker on each cell population.

### CD7 Median

| | | Ungated | CD34+CD38low | HSCs | MPP | CMP/GMP | Myelo/Mono-Blasts | ProMonocytes | CD14neg Monocytes | Mature Monocytes | ProMyelocytes | Myelocytes | MetaMyelocytes | MatureGrans | ProErythroblasts | Erythroblasts | LateErythroblasts | PreBcells | Mature Bcells | Plasma Cells | T cells | NK cells | pDCs | Basophils | Platelets | CD8+ T cells | CD8neg T cells |
|---|---|---|---|---|---|---|---|---|---|---|---|---|---|---|---|---|---|---|---|---|---|---|---|---|---|---|---|
| Run1_MDS17 | | 0.41 | 0.48 | 0.55 | 0.42 | 1.32 | 0.48 | -0.06 | -0.16 | -0.14 | -0.2 | 0 | -0.02 | 0.1 | 2.3 | 1.47 | 0.57 | 0.35 | -0.32 | 0.13 | 13.15 | 46.63 | 4.5 | 0.37 | 0.45 | 12.01 | 15.9 |
| Run2_MDS15* | AML/RAEB-T | -0.11 | 0.77 | 0.51 | 0.5 | 0.61 | 0.02 | -0.18 | -0.22 | -0.22 | -0.2 | -0.24 | -0.23 | -0.24 | 1.64 | 1.42 | 0.54 | -0.17 | -0.18 | 0.03 | 21.86 | 137.58 | 0.44 | -0.15 | 0.6 | 31.7 | 13.81 |
| Run2_MDS4 | | 0.08 | 4.2 | 1.04 | 13.09 | 1.01 | -0.11 | -0.23 | -0.2 | -0.15 | -0.25 | -0.27 | -0.27 | -0.25 | 1.02 | 0.99 | 0 | 0.41 | 0.05 | -0.17 | 19.39 | 147.26 | 21.25 | -0.1 | 0.74 | 22.35 | 15.81 |
| Run1_MDS21 | | 0.76 | 0.58 | 0.52 | 0.55 | 0.9 | 0.76 | 0.17 | -0.25 | -0.28 | -0.29 | -0.21 | -0.19 | -0.1 | 2.2 | 2.78 | 2.77 | -0.21 | -0.21 | 0.21 | 22 | 146.67 | 7.8 | -0.01 | 1.74 | 30.45 | 18.84 |
| Run1_MDS3 | | 0.45 | 0.33 | 0.16 | 0.31 | 0.57 | 0.41 | 0.13 | -0.09 | -0.28 | -0.28 | -0.18 | -0.15 | -0.02 | 1.19 | 1.62 | 1.21 | -0.22 | -0.2 | 0 | 26.05 | 154.8 | 4.96 | -0.14 | 0.84 | 36.59 | 22.01 |
| Run2_MDS13 | | 0.2 | 0.28 | 0.3 | 0.26 | 0.37 | 0.18 | -0.16 | -0.16 | 0.09 | -0.21 | -0.15 | -0.18 | -0.17 | 0.93 | 1.18 | 0.17 | -0.23 | -0.31 | 0.07 | 23.08 | 145.75 | -0.39 | -0.17 | 0.33 | 30.48 | 20.1 |
| Run2_MDS16 | | 0.23 | 0.4 | 0.32 | 0.32 | 0.27 | -0.09 | -0.17 | -0.26 | -0.22 | -0.13 | -0.29 | -0.25 | -0.24 | 1.39 | 3.44 | 0.23 | 17.56 | -0.3 | 0.12 | 38.03 | 125.16 | -0.02 | -0.28 | 0.79 | 44.21 | 36.76 |
| Run2_MDS1 | | -0.05 | -0.12 | -0.28 | -0.16 | -0.12 | -0.22 | -0.24 | -0.28 | -0.24 | -0.32 | -0.37 | -0.26 | -0.2 | 0.66 | 0.78 | 0.04 | 0.51 | -0.13 | -0.1 | 14.6 | 70.91 | -0.22 | -0.31 | 0.36 | 12.3 | 15.21 |
| Run2_MDS21 | Higher Risk | 0.32 | 0.21 | 0.39 | 0.2 | 0.55 | 0.31 | -0.22 | -0.23 | -0.16 | -0.21 | -0.18 | -0.22 | -0.18 | 0.71 | 1.04 | 0.07 | -0.23 | -0.21 | -0.02 | 17.25 | 68.03 | 6.23 | 0.23 | 0.92 | 25.52 | 14.63 |
| Run2_MDS26* | | -0.24 | 0.16 | 0.14 | 0.1 | 0.34 | -0.15 | -0.31 | -0.24 | 0.09 | -0.32 | -0.33 | -0.28 | -0.26 | 1.16 | 0.62 | -0.12 | -0.04 | -0.33 | -0.05 | 16.6 | 114.19 | 1.03 | -0.2 | -0.09 | 18.34 | 15.72 |
| Run2_MDS27 | | 1.14 | 0.44 | 0.46 | 0.34 | 0.73 | 0.4 | -0.18 | -0.24 | -0.2 | -0.12 | -0.23 | -0.22 | -0.2 | 1.46 | 3.8 | -0.07 | 11.55 | -0.24 | -0.06 | 25.25 | 107.87 | -0.04 | -0.15 | 1.39 | 29.61 | 21.02 |
| Run2_MDS2 | | -0.01 | -0.18 | -0.04 | -0.22 | -0.16 | -0.2 | -0.23 | -0.25 | -0.09 | -0.18 | -0.34 | -0.22 | -0.19 | 1.54 | 2.17 | 0.42 | 1.08 | 0.1 | -0.02 | 23.14 | 69.58 | -0.02 | -0.28 | 0.1 | 36.81 | 22.41 |
| Run2_MDS3 | | 0.37 | 0.26 | 0.4 | 0.16 | 0.43 | 0.12 | -0.26 | -0.14 | -0.15 | -0.17 | -0.06 | -0.03 | 0.03 | 1.43 | 0.74 | -0.04 | -0.23 | -0.25 | 0.03 | 22.24 | 140.38 | 1.98 | -0.19 | 0.48 | 32.15 | 18.43 |
| Run1_MDS23 | | -0.02 | 0.15 | 0.18 | 0.06 | 0.47 | -0.05 | -0.07 | -0.07 | -0.09 | -0.08 | -0.15 | -0.12 | -0.12 | 1.55 | 2.14 | 0.54 | 0.26 | -0.19 | -0.24 | 12.49 | 91.5 | 0.35 | -0.26 | 0.81 | 15.27 | 11.42 |
| Run1_MDS25 | | -0.11 | 1.66 | 1.34 | 1.77 | 0.89 | -0.1 | -0.21 | -0.22 | -0.21 | -0.17 | -0.23 | -0.24 | -0.21 | 2.86 | 3.28 | 0.78 | 0.85 | -0.22 | -0.13 | 25.32 | 74.65 | 1.76 | -0.23 | 0.67 | 24.04 | 26.59 |
| Run1_MDS5 | | 0.37 | 1.1 | 0.77 | 0.23 | 0.46 | 0.46 | 0.11 | 0.29 | 0.1 | 0.26 | -0.11 | 0.02 | 0 | 1.4 | 4.97 | 1.19 | 1.13 | -0.18 | 0.09 | 32.32 | 88.11 | 0.42 | -0.13 | 0.59 | 34.64 | 30.94 |
| Run1_MDS6 | | 0.01 | 0.62 | 0.34 | -0.13 | 0.89 | 0.25 | -0.09 | -0.14 | -0.11 | -0.13 | -0.18 | -0.15 | -0.19 | 5.12 | 5.48 | 0.77 | 0.49 | -0.26 | 0.28 | 26.56 | 170.82 | 0.59 | -0.13 | 2.48 | 35.01 | 22.04 |
| Run1_MDS8 | | 0.03 | 0.64 | 0.14 | 0.45 | 0.66 | 0.3 | 0.09 | -0.05 | -0.12 | -0.15 | -0.26 | -0.23 | -0.21 | 6.15 | 4.45 | 0.72 | 9.44 | -0.26 | 1.28 | 27.76 | 149 | 1.41 | -0.03 | 1.15 | 26.82 | 28.12 |
| Run1_MDS9 | | 0.5 | 0.52 | 0.54 | 0.3 | 0.85 | 0.37 | 0 | -0.11 | -0.07 | -0.14 | -0.18 | -0.16 | -0.13 | 4.39 | 6.12 | 1.41 | 15.63 | -0.23 | 0.26 | 19.84 | 140.33 | 0.44 | -0.03 | 0.51 | 21.56 | 18.9 |
| Run2_MDS12 | Lower Risk | -0.12 | 0.56 | 1.23 | 0.28 | 0.41 | -0.08 | -0.18 | -0.16 | -0.16 | -0.23 | -0.22 | -0.17 | -0.16 | 1.79 | 1.62 | -0.06 | -0.04 | -0.23 | -0.2 | 13.16 | 108.41 | 0.02 | -0.26 | 0.35 | 14.74 | 12.26 |
| Run2_MDS14 | | -0.1 | -0.11 | -0.56 | -0.18 | 0.43 | -0.19 | -0.18 | -0.15 | -0.22 | -0.15 | -0.22 | -0.18 | -0.2 | 2.51 | 4.59 | -0.01 | -0.27 | -0.27 | -0.18 | 13.76 | 152.83 | 0 | -0.15 | 1.78 | 17.88 | 12.22 |
| Run2_MDS19 | | -0.13 | 0.32 | -0.05 | 0.35 | 0.36 | -0.07 | -0.22 | -0.24 | -0.24 | -0.19 | -0.21 | -0.2 | -0.21 | 1.24 | 0.87 | -0.16 | 1.24 | -0.27 | -0.05 | 17.64 | 73.36 | 0.1 | -0.21 | 1.19 | 18.05 | 17.53 |
| Run2_MDS20 | | 0.48 | 0.26 | 0.14 | 0.06 | 0.91 | -0.03 | -0.11 | -0.18 | -0.16 | -0.18 | -0.19 | -0.14 | -0.16 | 3.07 | 2.36 | 0.39 | 8.22 | -0.12 | 1.29 | 18.45 | 79.52 | 0.08 | -0.22 | 0.65 | 16.79 | 18.63 |
| Run2_MDS28* | | 0.05 | 0.02 | -0.18 | -0.04 | 0.32 | 0.11 | -0.05 | -0.09 | -0.04 | -0.09 | -0.11 | -0.02 | -0.05 | 2.32 | 1.02 | 0.23 | 0.75 | -0.27 | -0.29 | 18.88 | 112.25 | 0.17 | -0.19 | 1.01 | 18.19 | 19.46 |
| Run2_MDS5 | | 0.12 | 0.65 | -0.05 | 0.28 | 0.88 | 0.11 | -0.16 | 0.12 | -0.06 | -0.05 | -0.18 | -0.11 | -0.08 | 0.74 | 3.03 | 0.75 | 1.3 | -0.26 | 0.33 | 30.62 | 96.77 | 0 | -0.08 | 0.74 | 36.12 | 28.78 |
| Run1_ICUS18 | | 0.14 | 0.75 | 0.29 | 0.41 | 0.67 | -0.06 | -0.08 | -0.19 | -0.23 | -0.09 | -0.15 | -0.13 | -0.15 | 2.21 | 1.84 | 0.18 | 10.59 | -0.29 | 0.09 | 24.1 | 129.79 | 0.54 | -0.23 | 0.99 | 25.12 | 23.91 |
| Run1_ICUS22 | ICUS | -0.06 | 0.46 | 0.54 | 0.62 | 0.45 | 0.08 | -0.06 | -0.11 | -0.14 | -0.12 | -0.16 | -0.16 | -0.19 | 2.66 | 3.09 | 0.35 | 0.04 | -0.24 | 0.03 | 14.1 | 87.72 | 0.26 | -0.19 | 0.77 | 13.45 | 14.36 |
| Run1_ICUS24 | | 0.02 | 0.37 | 0.33 | 0.16 | 0.58 | -0.02 | -0.14 | -0.22 | -0.23 | -0.11 | -0.16 | -0.15 | -0.18 | 3.32 | 2.89 | 1.28 | -0.14 | -0.25 | -0.21 | 10.16 | 85.87 | 0.51 | -0.26 | 2.14 | 14.02 | 7.17 |
| Run1_Nl-1 | | -0.04 | 0.14 | 0.11 | 0.06 | 0.22 | -0.1 | -0.14 | -0.13 | -0.2 | -0.07 | -0.19 | -0.19 | -0.19 | 2.39 | 2.86 | 0.59 | -0.22 | -0.31 | -0.08 | 20.68 | 154.96 | 0.63 | -0.09 | 3.14 | 21.73 | 20.13 |
| Run1_Nl-3 | | 0.13 | 0.56 | 0.46 | 0.23 | 0.37 | 0.06 | -0.02 | 0.11 | -0.09 | -0.07 | -0.14 | -0.08 | -0.08 | 3.16 | 1.86 | 0.29 | 0.01 | -0.34 | -0.16 | 19.71 | 125.46 | 0.17 | -0.17 | 0.08 | 21.87 | 18.27 |
| Run1_Nl-4 | | 0.14 | 0.53 | 0.78 | 0.45 | 0.74 | 0.15 | 0.17 | 0.13 | -0.08 | -0.04 | -0.1 | -0.06 | -0.07 | 3.46 | 2.05 | 0.26 | -0.01 | -0.29 | 0.31 | 22.33 | 158.7 | 0.57 | -0.24 | 0.19 | 25 | 20.26 |
| Run1_Nl-6 | | -0.01 | 0.35 | 0.21 | 0.2 | 0.42 | 0.1 | 0.1 | 0.15 | -0.02 | -0.08 | -0.14 | -0.12 | -0.15 | 2.68 | 2.14 | 0.21 | -0.21 | -0.3 | -0.07 | 18.7 | 143.11 | 0.22 | -0.21 | 0 | 22.27 | 17.64 |
| Run2_Nl-3 | Normal | -0.02 | 0.32 | 0.35 | 0.24 | 0.32 | 0 | -0.07 | -0.02 | -0.12 | -0.12 | -0.18 | -0.11 | -0.1 | 2.06 | 1.2 | 0.03 | -0.05 | -0.32 | -0.13 | 16.67 | 110.97 | 0.04 | -0.23 | 0.13 | 18.99 | 15.19 |
| Run2_Nl-4 | | 0.03 | 0.53 | 0.48 | 0.49 | 0.55 | 0.08 | -0.02 | 0.05 | -0.07 | -0.09 | -0.15 | -0.07 | -0.07 | 1.95 | 1.34 | 0.09 | -0.09 | -0.29 | -0.15 | 19.58 | 140.9 | 0.29 | -0.25 | 0.31 | 22.11 | 17.24 |
| Run2_Nl-5 | | 0.11 | 0.39 | 0.44 | 0.3 | 0.57 | -0.04 | 0.02 | 0.05 | -0.07 | -0.12 | -0.17 | -0.11 | -0.1 | 2.52 | 2.09 | 0.05 | -0.21 | -0.33 | -0.14 | 20.48 | 162.96 | 0.26 | -0.24 | 0.08 | 23.19 | 18.45 |
| Run2_Nl-6 | | -0.12 | 0.19 | 0.15 | 0.13 | 0.27 | 0.04 | -0.05 | 0.01 | -0.07 | -0.18 | -0.2 | -0.16 | -0.16 | 1.11 | 1.07 | -0.09 | -0.2 | -0.3 | -0.06 | 15.46 | 130.32 | -0.07 | -0.25 | 0.59 | 17.89 | 14.61 |

**Supplemental Table 3**: Median expression of each surface marker on each cell population.

### CD71 Median

| Sample | Group | Ungated | CD34+CD38low | HSCs | MPP | CMP/GMP | Myelo/Mono-Blasts | ProMonocytes | CD14neg Monocytes | Mature Monocytes | ProMyelocytes | Myelocytes | MetaMyelocytes | MatureGrans | ProErythroblasts | Erythroblasts | LateErythroblasts | PreBcells | Mature Bcells | Plasma Cells | T cells | NK cells | pDCs | Basophils | Platelets | CD8+ T cells | CD8neg T cells |
|---|---|---|---|---|---|---|---|---|---|---|---|---|---|---|---|---|---|---|---|---|---|---|---|---|---|---|---|
| Run1_MDS17 | | 3.32 | 10.51 | 10.78 | 10.08 | 19.57 | 10.6 | 3.73 | 1.4 | 1.7 | 4.09 | 1.88 | 0.54 | 0.7 | 196.03 | 201.9 | 44.74 | 2.76 | 1.7 | 9.34 | 1.21 | 1.96 | 4.75 | 2.7 | 3.47 | 1.07 | 1.44 |
| Run2_MDS15* | AML/RAEB-T | 0.91 | 8.34 | 4.91 | 4.58 | 6.29 | 6.6 | 2.66 | 0.16 | 0.11 | 3.74 | 0.9 | 0.1 | -0.11 | 108.2 | 221.06 | 3.03 | 0.23 | 1.93 | 45.19 | 1.46 | 3.16 | 4.08 | 0.77 | 3.64 | 1.7 | 1.17 |
| Run2_MDS4 | | 0.99 | 9.36 | 4.59 | 8.74 | 0.87 | 4.24 | 1.8 | 1.24 | 1.83 | 2.96 | 0.52 | -0.11 | -0.11 | 207.46 | 284.51 | 5 | 1.48 | 3.4 | 2.07 | 0.93 | 3.58 | 7.3 | 1.24 | 5.19 | 1.05 | 0.79 |
| Run1_MDS21 | | 3.19 | 6.8 | 7.02 | 6.35 | 9.25 | 9.63 | 6.24 | 2.73 | 2.46 | 2.13 | 1.08 | 0.19 | 0.49 | 393.38 | 482.58 | 15.07 | 1.99 | 2.72 | 7.73 | 1.5 | 3.9 | 4.8 | 1.42 | 8.49 | 1.85 | 1.33 |
| Run1_MDS3 | | 3.07 | 6.53 | 6.02 | 5.74 | 8.32 | 9.39 | 6.5 | 2.04 | 3.41 | 2.53 | 0.96 | 0.25 | 0.85 | 226.88 | 239.18 | 10.05 | 2.01 | 2.45 | 5.99 | 1.63 | 4.05 | 8.07 | 1.25 | 7.86 | 1.9 | 1.52 |
| Run2_MDS13 | | 2.37 | 4.7 | 4.83 | 4.39 | 6.06 | 6.67 | 2.29 | 1.28 | 1.04 | 5.19 | 1.35 | -0.04 | -0.06 | 147.99 | 300.66 | 25.93 | 1.4 | 1.37 | 46.54 | 1.11 | 3.4 | 11.15 | 0.75 | 6.44 | 1.4 | 0.98 |
| Run2_MDS16 | | 0.53 | 23.68 | 18.51 | 23.67 | 22.93 | 7.76 | 3.7 | 2.8 | 1.79 | 10.93 | 0.39 | -0.17 | -0.19 | 199.02 | 772.24 | 7.76 | 2.18 | 0.85 | 7.58 | 1.59 | 2.88 | 7.52 | 0.51 | 8.08 | 1.81 | 1.53 |
| Run2_MDS1 | | 0.59 | 4.92 | 4.86 | 4.57 | 5.64 | 4.24 | 1.36 | 0.54 | 0.67 | 0.29 | -0.2 | -0.14 | -0.11 | 188.05 | 219.04 | 14.26 | 1.22 | 1.83 | 2.54 | 1.04 | 2.01 | 4.94 | 0.93 | 5.55 | 0.81 | 1.12 |
| Run2_MDS21 | Higher Risk | 1.93 | 4.74 | 4.84 | 4.44 | 6.75 | 6.53 | 2.9 | 1.15 | 0.78 | 2.02 | 0.63 | -0.06 | -0.1 | 155.09 | 268.91 | 31.65 | 1.26 | 1.72 | 6.51 | 0.93 | 2.3 | 5.22 | 0.39 | 5.43 | 1.14 | 0.82 |
| Run2_MDS26* | | 0.51 | 14.75 | 14.14 | 13.94 | 30.06 | 11.89 | 5.9 | 3.25 | 2.58 | 1.36 | 0.24 | -0.08 | -0.09 | 195.68 | 163.55 | 31.2 | 1.35 | 1.11 | 6.94 | 1.09 | 2.78 | 7.15 | 1.78 | 1.66 | 1.2 | 1.02 |
| Run2_MDS27 | | 2.54 | 9.41 | 6.59 | 10.89 | 21.33 | 15.9 | 5.21 | 1.44 | 0.76 | 10.07 | 1.11 | -0.09 | -0.14 | 208.48 | 835.57 | 23.68 | 2.09 | 1.26 | 4.18 | 1.3 | 2.64 | 3.49 | 0.77 | 8.28 | 1.43 | 1.13 |
| Run2_MDS2 | | 0.3 | 1.71 | 2.93 | 1.44 | 2.27 | 1.9 | 0.43 | -0.02 | 0.23 | 1.89 | -0.03 | -0.11 | -0.07 | 148.77 | 257.59 | 4.24 | 0.91 | 1.71 | 27.82 | 0.87 | 1.27 | 0.26 | 0 | 1.59 | 1.33 | 0.84 |
| Run2_MDS3 | | 2.68 | 4.42 | 4.99 | 4.15 | 6.38 | 7.43 | 2.41 | 1.52 | 1.95 | 4.06 | 1.08 | 0.08 | 0.11 | 299.06 | 190.97 | 32.14 | 1.14 | 1.74 | 5.82 | 1.07 | 2.82 | 5.88 | 0.53 | 6.68 | 1.34 | 0.97 |
| Run1_MDS23 | | 0.79 | 6.25 | 5.29 | 5.22 | 26.33 | 11.33 | 4.89 | 3.16 | 1.93 | 1.42 | 0.35 | 0.22 | 0.27 | 248.86 | 304.76 | 4.26 | 2.31 | 1.64 | 4.36 | 1.51 | 3.53 | 19.67 | 1.77 | 4.74 | 1.82 | 1.4 |
| Run1_MDS25 | | 0.42 | 6.33 | 8.16 | 5.02 | 39.09 | 12.67 | 10.08 | 7.48 | 4.68 | 6.52 | 0.68 | 0.03 | 0.04 | 452.03 | 505.28 | 2.22 | 3.6 | 2.47 | 19.36 | 2.49 | 3.2 | 22.23 | 1.58 | 4.9 | 2.18 | 2.85 |
| Run1_MDS5 | | 1.93 | 5.51 | 4.27 | 4.18 | 10.71 | 11.79 | 4.13 | 3.55 | 3.75 | 16.9 | 2.21 | 2.15 | 1.44 | 123.19 | 662.25 | 4.68 | 2.26 | 1.06 | 37.6 | 2.06 | 3.11 | 4.39 | 1.8 | 3.67 | 2.76 | 1.85 |
| Run1_MDS6 | | 0.56 | 11.28 | 7.43 | 7.63 | 17.89 | 11.25 | 4.93 | 2.83 | 1.69 | 3.89 | 0.63 | 0.17 | 0.01 | 311.41 | 905.93 | 3.73 | 1.47 | 1.83 | 7.18 | 1.69 | 4.31 | 7.24 | 0.71 | 8.04 | 2.23 | 1.28 |
| Run1_MDS8 | | 0.55 | 4.03 | 4.34 | 2.52 | 17.08 | 10.45 | 5.27 | 3.51 | 2.42 | 4.68 | 0.61 | 0.02 | 0.05 | 1054.81 | 811.08 | 7.17 | 1.68 | 1.21 | 5.16 | 1.28 | 4.02 | 10.34 | 4.2 | 6.23 | 1.59 | 1.19 |
| Run1_MDS9 | | 1.06 | 26.5 | 24.36 | 27.67 | 37.3 | 23.11 | 6.87 | 4.22 | 2.66 | 2.49 | 0.59 | 0.31 | 0.34 | 740.39 | 1033.66 | 6.27 | 3.06 | 2.1 | 7.1 | 1.57 | 4.31 | 6.33 | 2.31 | 4.76 | 1.65 | 1.54 |
| Run2_MDS12 | Lower Risk | 0.41 | 22.63 | 17.89 | 20.41 | 26.06 | 8.28 | 4.12 | 2.66 | 1.79 | 2.05 | 0.47 | -0.03 | -0.03 | 353.76 | 410.44 | 14.45 | 1.86 | 1.13 | 2.98 | 0.8 | 2.96 | 28.47 | 1.24 | 4.97 | 1.04 | 0.65 |
| Run2_MDS14 | | 0.09 | 3.58 | 0.89 | 2.4 | 17.69 | 5.8 | 5.17 | 4.38 | 2.77 | 8.22 | 0.53 | -0.08 | -0.17 | 501.01 | 1059.69 | 0.77 | 1.16 | 0.96 | 5.63 | 1.17 | 3.35 | 14.18 | 0.68 | 9.97 | 1.32 | 1.05 |
| Run2_MDS19 | | 0.18 | 8.01 | 5.79 | 7.16 | 21.81 | 8.77 | 3.18 | 1.74 | 0.9 | 2.46 | 0.16 | -0.13 | -0.16 | 224.87 | 256.37 | 14.08 | 1.51 | 1.48 | 1.8 | 1.21 | 2.27 | 13.58 | 0.86 | 5.1 | 1.18 | 1.22 |
| Run2_MDS20 | | 0.35 | 1.47 | 4.98 | 1.01 | 27.8 | 4.23 | 2.7 | 0.96 | 0.6 | 2.64 | 0.38 | -0.08 | -0.11 | 499 | 413.63 | 1.36 | 1 | 2.47 | 10.01 | 0.85 | 2.59 | 5.07 | 0.8 | 4.85 | 1.2 | 0.82 |
| Run2_MDS28* | | 0.45 | 8.15 | 4.7 | 8.96 | 15.97 | 5.44 | 2.82 | 1.52 | 1.67 | 1.63 | 0.43 | 0.22 | 0.12 | 307.54 | 220.85 | 1.8 | 1.29 | 1.47 | 0.91 | 0.94 | 2.67 | 5.37 | 1.26 | 10.5 | 0.96 | 0.92 |
| Run2_MDS5 | | 1.11 | 4.14 | 2.38 | 4.52 | 8.01 | 10.49 | 2.35 | 2.44 | 2.15 | 11.13 | 1 | 1.15 | 0.81 | 135.48 | 535.92 | 3.14 | 2.07 | 0.56 | 33.87 | 1.61 | 2.7 | 6.69 | 1.17 | 4.1 | 2.32 | 1.35 |
| Run1_ICUS18 | | 0.54 | 6.8 | 5.8 | 4.71 | 18.62 | 8.3 | 4.82 | 2.51 | 1.48 | 4.27 | 0.93 | 0.26 | 0.14 | 317.53 | 302.12 | 0.76 | 2.41 | 1.8 | 2.67 | 1.66 | 3.58 | 18.57 | 0.79 | 4.78 | 1.74 | 1.64 |
| Run1_ICUS22 | ICUS | 0.61 | 17.17 | 12.54 | 18.98 | 29.37 | 13.01 | 8.21 | 5.21 | 2.96 | 3.35 | 1 | 0.25 | 0.15 | 405.22 | 499.64 | 1.3 | 2.08 | 2.04 | 5.5 | 1.61 | 3.57 | 12.21 | 1.04 | 6.16 | 1.42 | 1.68 |
| Run1_ICUS24 | | 0.72 | 14.56 | 9.45 | 17.64 | 29.84 | 13.34 | 8.32 | 5.03 | 2.79 | 3.8 | 0.78 | 0.18 | 0.12 | 544.75 | 510.06 | 9.3 | 1.45 | 1.64 | 7.06 | 1.14 | 3.15 | 14.91 | 1.58 | 10.5 | 1.13 | 1.15 |
| Run1_Nl-1 | | 0.69 | 12.21 | 9.82 | 11.92 | 15.1 | 13.67 | 12.28 | 10.39 | 5.65 | 8.27 | 1.21 | 0.25 | 0.11 | 374.34 | 495.25 | 2.18 | 1.61 | 1.4 | 5.6 | 1.82 | 4.17 | 23.68 | 2.22 | 14.6 | 1.7 | 1.88 |
| Run1_Nl-3 | | 0.71 | 12.62 | 10.77 | 10.37 | 22.1 | 11.56 | 5.31 | 4.12 | 2.05 | 7.16 | 0.7 | 0.24 | 0.14 | 475.28 | 301.68 | 0.9 | 1.32 | 0.9 | 3.79 | 1.26 | 3.76 | 10.76 | 1.51 | 2.88 | 1.44 | 1.15 |
| Run1_Nl-4 | | 0.73 | 11.82 | 14.24 | 10.55 | 29.93 | 13.26 | 4.84 | 2.81 | 1.39 | 7.05 | 0.87 | 0.26 | 0.16 | 530.22 | 334.24 | 0.78 | 1.34 | 1.44 | 5.12 | 1.33 | 4.13 | 13.26 | 1.05 | 3.01 | 1.5 | 1.19 |
| Run1_Nl-6 | | 0.8 | 7.09 | 6.57 | 7.07 | 19.67 | 13.36 | 6.14 | 3.64 | 2.12 | 6.03 | 0.83 | 0.19 | 0.1 | 443.19 | 359 | 1.95 | 1.4 | 1.35 | 6.08 | 1.42 | 3.9 | 9.43 | 1.58 | 3.18 | 1.53 | 1.36 |
| Run2_Nl-3 | Normal | 0.19 | 10.29 | 9.79 | 10.29 | 20.65 | 8.74 | 3.26 | 2.01 | 1.24 | 4.69 | 0.02 | -0.13 | -0.15 | 369.95 | 260.19 | 0.18 | 0.83 | 0.62 | 2.44 | 0.83 | 2.82 | 8.27 | 0.88 | 2.97 | 0.95 | 0.76 |
| Run2_Nl-4 | | 0.22 | 12.2 | 11.99 | 10.96 | 27.66 | 9.81 | 2.59 | 1.27 | 0.86 | 5 | 0.17 | -0.1 | -0.13 | 331.43 | 288.7 | 0.2 | 0.82 | 1.01 | 4.05 | 0.88 | 3.14 | 10.83 | 0.67 | 3.17 | 1 | 0.77 |
| Run2_Nl-5 | | 0.5 | 17.38 | 15.12 | 18.43 | 24.09 | 13.05 | 5.71 | 2.39 | 1.45 | 7.81 | 0.18 | -0.12 | -0.17 | 372.8 | 449.97 | 0.5 | 0.73 | 0.71 | 4.24 | 1.06 | 3.46 | 10.82 | 0.97 | 2.23 | 1.18 | 0.96 |
| Run2_Nl-6 | | 0.19 | 5.13 | 4.32 | 4.84 | 17.14 | 10.18 | 3.79 | 2.17 | 1.42 | 3.87 | 0.29 | -0.13 | -0.18 | 203.31 | 245.29 | 2.46 | 0.87 | 0.93 | 5.27 | 0.88 | 3.05 | 7.82 | 0.66 | 3.85 | 0.97 | 0.85 |

**Supplemental Table 3**: Median expression of each surface marker on each cell population.

### CD235 Median

| | | Ungated | CD34+CD38low | HSCs | MPP | CMP/GMP | Myelo/Mono-Blasts | ProMonocytes | CD14neg Monocytes | Mature Monocytes | ProMyelocytes | Myelocytes | MetaMyelocytes | MatureGrans | ProErythroblasts | Erythroblasts | LateErythroblasts | PreBcells | Mature Bcells | Plasma Cells | T cells | NK cells | pDCs | Basophils | Platelets | CD8+ T cells | CD8neg T cells |
|---|---|---|---|---|---|---|---|---|---|---|---|---|---|---|---|---|---|---|---|---|---|---|---|---|---|---|---|
| Run1_MDS17 | | 2.59 | 1.67 | 1.68 | 1.54 | 2.73 | 1.58 | 2.13 | 2.52 | 2.2 | 1.99 | 3.2 | 3.25 | 3.81 | 11.23 | 40.51 | 26.47 | 1.93 | 0.84 | 1.92 | 1.24 | 1.31 | 1.96 | 2.89 | 3.92 | 1.39 | 1.04 |
| Run2_MDS15* | AML/RAEB-T | 1.73 | 2.51 | 1.97 | 1.81 | 1.89 | 1.81 | 1.49 | 1.46 | 1.42 | 1.68 | 1.76 | 1.93 | 1.95 | 5.46 | 83.03 | 27.28 | 1.51 | 0.97 | 3.55 | 1.38 | 1.68 | 2.04 | 1.44 | 2.49 | 1.63 | 1.16 |
| Run2_MDS4 | | 1.54 | 1.81 | 1.55 | 1.44 | 0.26 | 0.99 | 1.11 | 1.39 | 1.7 | 1.24 | 1.21 | 1.44 | 1.63 | 10.1 | 78.12 | 44.94 | 0.91 | 0.91 | 1.89 | 0.97 | 1.3 | 1.07 | 1.11 | 24.82 | 1.23 | 0.76 |
| Run1_MDS21 | | 2.05 | 1.31 | 1.67 | 1.15 | 1.49 | 1.46 | 1.47 | 1.65 | 1.27 | 1.31 | 2.13 | 2.47 | 3.33 | 9.07 | 26.46 | 22.12 | 0.65 | 0.95 | 2.65 | 1.11 | 2.17 | 2.03 | 1.46 | 6.6 | 1.38 | 1 |
| Run1_MDS3 | | 2.23 | 1.31 | 1.28 | 1.09 | 1.49 | 1.6 | 1.44 | 1.92 | 2.71 | 1.66 | 2.34 | 2.83 | 3.83 | 8.67 | 26.5 | 23.74 | 0.79 | 1.02 | 2.49 | 1.36 | 2.05 | 3.76 | 1.01 | 5.49 | 1.69 | 1.23 |
| Run2_MDS13 | | 1.52 | 1.09 | 1.18 | 0.96 | 1.22 | 1.23 | 1.54 | 2.38 | 2.62 | 1.49 | 2.03 | 2.38 | 2.45 | 6.32 | 73.16 | 28.07 | 0.52 | 0.65 | 2.84 | 0.94 | 1.34 | 0.57 | 1.07 | 3.24 | 1.21 | 0.82 |
| Run2_MDS16 | | 2.54 | 2.49 | 2.35 | 2.21 | 1.98 | 1.85 | 1.33 | 1.39 | 1.56 | 2.19 | 1.87 | 2.71 | 3.1 | 5.4 | 212.13 | 33.39 | 1.34 | 0.46 | 1.49 | 1.1 | 1.41 | 1.76 | 0.99 | 5.54 | 1.54 | 1.01 |
| Run2_MDS1 | | 0.86 | 0.99 | 0.96 | 0.77 | 1.2 | 0.78 | 0.7 | 0.6 | 0.97 | 0.5 | 0.36 | 1.13 | 1.39 | 5.02 | 64.29 | 45.34 | 0.83 | 0.9 | 1.89 | 0.8 | 1.02 | 0.87 | 1.08 | 7.09 | 0.88 | 0.79 |
| Run2_MDS21 | Higher Risk | 1.57 | 0.86 | 1.09 | 0.78 | 1.09 | 1 | 0.79 | 1.31 | 1.56 | 1.37 | 1.9 | 2.1 | 2.32 | 9.43 | 54.3 | 28.25 | 0.45 | 0.59 | 2.25 | 0.77 | 0.88 | 0.35 | 1.05 | 4.97 | 1 | 0.67 |
| Run2_MDS26* | | 1.1 | 1.63 | 1.72 | 1.51 | 2.12 | 1.15 | 1.09 | 1.25 | 1.19 | 0.53 | 0.54 | 0.88 | 1.05 | 6.41 | 85.85 | 43.03 | 0.92 | 0.23 | 0.78 | 0.76 | 1.13 | 1.36 | 0.89 | 0.93 | 0.98 | 0.63 |
| Run2_MDS27 | | 3.46 | 2.16 | 2.12 | 2.03 | 2.24 | 2.06 | 1.32 | 1 | 1.11 | 2.62 | 2.61 | 3.51 | 4.38 | 5.31 | 327.88 | 58.35 | 1.46 | 0.7 | 2.22 | 1.27 | 1.11 | 1.8 | 1.15 | 4.38 | 1.49 | 1.02 |
| Run2_MDS2 | | 0.78 | 0.51 | 0.83 | 0.29 | 0.64 | 0.41 | 0.18 | 0.45 | 0.78 | 0.77 | 0.72 | 0.98 | 1.21 | 8.66 | 88.58 | 38.25 | 0.67 | 0.76 | 0.35 | 0.64 | 0.59 | 0.65 | 0.35 | 3.53 | 0.98 | 0.61 |
| Run2_MDS3 | | 2.16 | 1.05 | 1.27 | 0.96 | 1.3 | 1.27 | 1.36 | 2.15 | 2.33 | 1.99 | 3.04 | 3.32 | 4.14 | 6.95 | 74.86 | 35.21 | 0.42 | 0.81 | 1.26 | 1 | 1.25 | 1.68 | 1.04 | 5.47 | 1.26 | 0.89 |
| Run1_MDS23 | | 2.29 | 1.53 | 1.58 | 1.06 | 2.74 | 2.06 | 1.94 | 1.89 | 1.72 | 2.26 | 2.15 | 2.43 | 2.65 | 8.85 | 26.47 | 25.45 | 1.5 | 0.72 | 2.04 | 1.31 | 1.6 | 2.09 | 1.11 | 9.32 | 1.73 | 1.16 |
| Run1_MDS25 | | 2.54 | 5.51 | 2.96 | 3.87 | 2.95 | 1.86 | 1.7 | 1.39 | 1.23 | 2.5 | 2.06 | 2.32 | 2.72 | 11.89 | 34.79 | 22.18 | 1.62 | 1.21 | 2.92 | 1.65 | 1.83 | 2.51 | 1.33 | 4.39 | 1.99 | 1.29 |
| Run1_MDS5 | | 3.66 | 4.14 | 0.42 | 2.91 | 1.94 | 2.52 | 2.8 | 3.35 | 2.74 | 3.55 | 2.72 | 3.36 | 3.66 | 7.47 | 183.02 | 62.63 | 3.15 | 0.99 | 3.94 | 1.72 | 2.2 | 2.18 | 2.15 | 26.26 | 2.16 | 1.52 |
| Run1_MDS6 | | 3.63 | 2.28 | 1.7 | 1.31 | 1.93 | 2.31 | 1.94 | 1.95 | 1.96 | 2.66 | 2.8 | 3.61 | 4.02 | 9.16 | 133.76 | 22.16 | 1.35 | 0.91 | 3.24 | 1.64 | 2.26 | 2.35 | 1.61 | 11.69 | 2.15 | 1.22 |
| Run1_MDS8 | | 2.77 | 3.83 | 2.9 | 3.05 | 2.51 | 2 | 2.08 | 1.85 | 1.68 | 2.34 | 2.16 | 2.64 | 3.01 | 11.4 | 52.56 | 30.16 | 1.43 | 0.72 | 5.78 | 1.01 | 1.85 | 2.96 | 1.89 | 7.1 | 1.32 | 0.92 |
| Run1_MDS9 | | 2.83 | 2.71 | 2.68 | 2.4 | 3.35 | 2.58 | 2.1 | 1.85 | 2 | 2.62 | 2.49 | 2.84 | 3.42 | 10.3 | 33.5 | 23.03 | 1.61 | 0.83 | 3.06 | 1.27 | 2.04 | 1.64 | 1.96 | 7.02 | 1.63 | 1.07 |
| Run2_MDS12 | Lower Risk | 2.08 | 2.42 | 2.48 | 1.66 | 2.66 | 1.76 | 1.63 | 1.76 | 1.67 | 1.06 | 1.41 | 2.14 | 2.31 | 10.42 | 75.7 | 31.15 | 1.12 | 0.66 | 1.22 | 0.75 | 1.32 | 2.18 | 0.94 | 5.04 | 1.02 | 0.58 |
| Run2_MDS14 | | 3.54 | 3.32 | 1.84 | 5.52 | 0.79 | 1.26 | 1.55 | 1.58 | 1.56 | 2.72 | 2.56 | 3.43 | 3.93 | 9.51 | 104.83 | 21.57 | 0.73 | 0.67 | 1.52 | 1.08 | 1.55 | 1.22 | 1.36 | 11.35 | 1.36 | 0.92 |
| Run2_MDS19 | | 2.96 | 2.03 | 2 | 1.87 | 2.48 | 1.79 | 1.2 | 1.08 | 1.07 | 2.07 | 2.65 | 3.25 | 3.47 | 7.65 | 97.24 | 32.18 | 1.1 | 0.64 | 1.94 | 1.14 | 1.19 | 1.92 | 0.83 | 8.95 | 1.42 | 1.06 |
| Run2_MDS20 | | 2.6 | 3.17 | 1.51 | 2.11 | 2.1 | 1.54 | 1.59 | 1.35 | 1.44 | 2.5 | 2.93 | 3.38 | 4.09 | 7.77 | 144.42 | 21.3 | 1.32 | 1.14 | 3.63 | 0.87 | 0.98 | 1.81 | 1.27 | 4.41 | 1.34 | 0.83 |
| Run2_MDS28* | | 2.62 | 1.4 | 1.17 | 0.87 | 1.99 | 1.83 | 1.55 | 1.64 | 1.64 | 1.98 | 2.07 | 2.96 | 3.18 | 6.06 | 88.52 | 21.58 | 1.37 | 0.55 | 1.51 | 0.95 | 1.27 | 2.02 | 1.18 | 6.37 | 1.1 | 0.83 |
| Run2_MDS5 | | 3.39 | 2.63 | 0.71 | 1.77 | 2.31 | 2.55 | 2.58 | 2.97 | 2.13 | 2.7 | 2.72 | 3.45 | 3.67 | 5.69 | 249.27 | 64 | 2.44 | 0.61 | 2.88 | 1.34 | 1.66 | 2.65 | 1.73 | 59.58 | 1.67 | 1.17 |
| Run1_ICUS18 | | 3.98 | 2.53 | 2.05 | 1.29 | 2.82 | 1.94 | 1.86 | 1.5 | 1.45 | 3.4 | 3.5 | 4.19 | 4.77 | 11.4 | 42 | 19.31 | 1.86 | 0.85 | 2.07 | 1.46 | 1.88 | 2.66 | 1.02 | 5.31 | 1.93 | 1.35 |
| Run1_ICUS22 | ICUS | 2.78 | 2.66 | 2.44 | 2.64 | 2.73 | 2.16 | 1.83 | 1.54 | 1.41 | 2.56 | 2.66 | 3.02 | 3.17 | 9.75 | 29.29 | 19.77 | 1.03 | 0.79 | 1.34 | 1.41 | 1.83 | 2.35 | 1.29 | 5.66 | 1.72 | 1.29 |
| Run1_ICUS24 | | 2.88 | 2.88 | 2.29 | 2.72 | 2.79 | 1.97 | 1.54 | 1.51 | 1.45 | 2.86 | 2.72 | 3.12 | 3.46 | 10.93 | 41.1 | 24.5 | 0.89 | 0.8 | 2.28 | 1.24 | 1.63 | 2.47 | 1.09 | 11.59 | 1.36 | 1.15 |
| Run1_Nl-1 | | 3.29 | 2.46 | 3.46 | 1.22 | 2.32 | 1.87 | 2.11 | 1.82 | 1.74 | 3.45 | 2.99 | 3.53 | 3.85 | 10.7 | 31.76 | 19.94 | 0.81 | 0.71 | 1.36 | 1.43 | 1.91 | 2.71 | 1.66 | 12.37 | 1.77 | 1.25 |
| Run1_Nl-3 | | 5.1 | 3.01 | 2.88 | 2.3 | 2.59 | 2.56 | 3.08 | 3.94 | 2.75 | 4.06 | 4.32 | 5.83 | 6.4 | 9.53 | 65.41 | 19.68 | 1.04 | 0.58 | 2.34 | 1.29 | 1.86 | 2.25 | 1.54 | 4.35 | 1.56 | 1.11 |
| Run1_Nl-4 | | 5.94 | 3.47 | 2.97 | 3.17 | 2.9 | 2.76 | 3.65 | 4.14 | 3.1 | 4.62 | 5.17 | 6.55 | 6.83 | 9.62 | 65.54 | 19.7 | 1.14 | 0.8 | 2.89 | 1.29 | 1.89 | 2.5 | 1.51 | 3.08 | 1.53 | 1.1 |
| Run1_Nl-6 | | 4.44 | 2.17 | 1.87 | 1.99 | 2.72 | 2.54 | 3.22 | 3.7 | 2.97 | 3.52 | 3.83 | 4.69 | 4.81 | 11.13 | 55.79 | 20.08 | 0.94 | 0.65 | 2.79 | 1.22 | 1.84 | 2.08 | 1.43 | 2.64 | 1.6 | 1.12 |
| Run2_Nl-3 | Normal | 4.72 | 2.17 | 1.9 | 2.06 | 2.28 | 2.25 | 2.78 | 3.49 | 2.45 | 3.62 | 4.01 | 5.76 | 6.59 | 7.96 | 85.65 | 20.12 | 0.86 | 0.34 | 1.59 | 0.89 | 1.24 | 1.69 | 0.96 | 5.03 | 1.1 | 0.75 |
| Run2_Nl-4 | | 5.83 | 2.62 | 2.33 | 2.91 | 2.57 | 2.46 | 3.39 | 3.78 | 2.96 | 4.28 | 4.94 | 6.74 | 7.17 | 7.81 | 82.75 | 20.15 | 0.76 | 0.55 | 2.38 | 0.94 | 1.2 | 1.79 | 0.77 | 3.89 | 1.14 | 0.81 |
| Run2_Nl-5 | | 4.94 | 2.95 | 1.77 | 2.56 | 2.33 | 2.47 | 3.41 | 4.15 | 3.17 | 3.69 | 4.32 | 5.83 | 6.54 | 7.52 | 120.65 | 20.5 | 0.7 | 0.37 | 2.28 | 0.95 | 1.25 | 1.82 | 0.97 | 3.06 | 1.19 | 0.75 |
| Run2_Nl-6 | | 3.91 | 1.8 | 1.31 | 1.7 | 2.13 | 2.13 | 2.86 | 3.12 | 2.62 | 2.86 | 3.34 | 4.28 | 4.7 | 8.23 | 66.45 | 21.66 | 0.69 | 0.44 | 2.11 | 0.86 | 1.3 | 1.47 | 0.89 | 3.79 | 1.02 | 0.81 |

**Supplemental Table 3**: Median expression of each surface marker on each cell population.

### CD47 Median

| | | Ungated | CD34+CD38low | HSCs | MPP | CMP/GMP | Myelo/Mono-Blasts | ProMonocytes | CD14neg Monocytes | Mature Monocytes | ProMyelocytes | Myelocytes | MetaMyelocytes | MatureGrans | ProErythroblasts | Erythroblasts | LateErythroblasts | PreBcells | Mature Bcells | Plasma Cells | T cells | NK cells | pDCs | Basophils | Platelets | CD8+ T cells | CD8neg T cells |
|---|---|---|---|---|---|---|---|---|---|---|---|---|---|---|---|---|---|---|---|---|---|---|---|---|---|---|---|
| Run1_MDS17 | | 51.05 | 34.69 | 34.28 | 32.92 | 67.75 | 30.82 | 43.89 | 61 | 56.4 | 66.53 | 67.36 | 91.89 | 109.16 | 20.84 | 21.03 | 23.19 | 45.56 | 31.6 | 65.03 | 35.55 | 35.98 | 40.68 | 63.69 | 77.44 | 28.48 | 49.12 |
| Run2_MDS15* | AML/RAEB-T | 39.61 | 51.32 | 39.88 | 39.37 | 45.55 | 49.19 | 38.64 | 32.5 | 26.59 | 50.12 | 40.03 | 45.37 | 48.48 | 90.44 | 43.02 | 55.21 | 30.14 | 38.79 | 48.67 | 60.17 | 53.36 | 48.69 | 68.76 | 80.62 | 42.91 | 78.23 |
| Run2_MDS4 | | 35.94 | 32.46 | 32.69 | 31.84 | 69.66 | 25.65 | 30.5 | 35.79 | 36.82 | 45.09 | 42.03 | 45.87 | 43.02 | 16.1 | 16.56 | 13.38 | 35.27 | 31.7 | 30.68 | 37.94 | 33.08 | 25.23 | 73.16 | 179.71 | 33.38 | 43.15 |
| Run1_MDS21 | | 41.1 | 27.47 | 29.05 | 26.08 | 34.35 | 32.93 | 38.34 | 52.98 | 51.26 | 36.73 | 58.97 | 82.99 | 92.87 | 17.22 | 21.04 | 544.06 | 32.7 | 40.67 | 166.43 | 49.95 | 46.82 | 35.34 | 54.43 | 584.93 | 41.06 | 55.02 |
| Run1_MDS3 | | 42.56 | 26.15 | 25.02 | 24.6 | 33.03 | 32.51 | 37.28 | 53.73 | 54.75 | 40.7 | 56.99 | 78.54 | 96.83 | 22.62 | 25.67 | 118.6 | 32.14 | 40.26 | 78.53 | 59.94 | 52.19 | 78.84 | 65.19 | 352.73 | 45.15 | 67.69 |
| Run2_MDS13 | | 36.65 | 18.69 | 19.35 | 17.92 | 25.58 | 25.58 | 36.5 | 44.48 | 40.53 | 46.22 | 61.5 | 69.67 | 64.61 | 16.58 | 12.05 | 12.28 | 23.45 | 26.96 | 30.08 | 42.89 | 37.99 | 42.53 | 52.73 | 147.74 | 35.68 | 47.64 |
| Run2_MDS16 | | 41.79 | 29.59 | 29.66 | 26.85 | 36.15 | 41.68 | 40.01 | 36.94 | 34.4 | 69.18 | 43.33 | 45.11 | 41.12 | 20.72 | 9.14 | 47.18 | 39.68 | 25.18 | 54.85 | 48.62 | 55.67 | 47.1 | 66.05 | 410.73 | 39.65 | 50.94 |
| Run2_MDS1 | | 48.48 | 40.55 | 33.7 | 38.64 | 51.25 | 39.98 | 35.83 | 46.76 | 50.89 | 23.14 | 23.53 | 65.55 | 71.45 | 23.78 | 16.9 | 17.03 | 45.32 | 29.89 | 48.2 | 52.72 | 57.83 | 54.74 | 66.24 | 366.05 | 35.36 | 56.24 |
| Run2_MDS21 | Higher Risk | 35.55 | 20.85 | 22.74 | 20.61 | 27.15 | 24.63 | 33.76 | 47.52 | 42.85 | 48.21 | 59.31 | 79.41 | 71.97 | 10.23 | 9.8 | 10.81 | 25.16 | 32.23 | 70.16 | 43.69 | 41.34 | 17.12 | 49.8 | 394.03 | 37.64 | 46.58 |
| Run2_MDS26* | | 40.15 | 32.25 | 33.68 | 30.85 | 43.3 | 23.27 | 37.93 | 51.5 | 49.68 | 32.87 | 31.2 | 57.48 | 64.24 | 26.07 | 26.22 | 15.84 | 30.62 | 20.01 | 35.82 | 42.76 | 45.94 | 50.21 | 64.75 | 52.01 | 35.45 | 47.49 |
| Run2_MDS27 | | 39.79 | 41.78 | 42.11 | 38.79 | 51.33 | 40.06 | 34.84 | 30.69 | 30.25 | 86.64 | 62.05 | 57.98 | 50.7 | 25.66 | 8.79 | 10.06 | 39.58 | 29.09 | 82.7 | 48.53 | 37.43 | 69.22 | 69.88 | 165.99 | 40.27 | 58.3 |
| Run2_MDS2 | | 53.83 | 17.17 | 19.22 | 15.18 | 21.89 | 22.16 | 31.46 | 42.4 | 34.49 | 36.26 | 36.9 | 73.78 | 73.3 | 58.02 | 52.47 | 79.83 | 38.05 | 49.9 | 24.43 | 41.83 | 58.87 | 50.28 | 48.13 | 532.67 | 34.09 | 42.27 |
| Run2_MDS3 | | 41.27 | 21.46 | 24.3 | 19.89 | 28.8 | 28.72 | 37.36 | 46.81 | 39.57 | 51.32 | 64.49 | 79.46 | 79.69 | 14.51 | 15.14 | 10.34 | 27 | 35.32 | 49.83 | 53.77 | 43.56 | 50.77 | 50.72 | 278.22 | 41.48 | 60.1 |
| Run1_MDS23 | | 83.78 | 49.91 | 48.91 | 45.76 | 68.01 | 48.42 | 52.53 | 59.36 | 57.23 | 65 | 69.44 | 102.02 | 109.89 | 40.89 | 47.35 | 120.4 | 55.76 | 38.04 | 58.95 | 67.64 | 52.2 | 74.89 | 92.77 | 181.77 | 50.71 | 74.16 |
| Run1_MDS25 | | 73.65 | 67.18 | 35.49 | 89.09 | 29.88 | 51.31 | 39.9 | 39.71 | 39.58 | 77.33 | 73.06 | 73.87 | 78.48 | 31.64 | 45.51 | 103.07 | 45.22 | 49.19 | 87.52 | 57.57 | 74.84 | 71.77 | 99.45 | 400.58 | 51.88 | 64.31 |
| Run1_MDS5 | | 91.19 | 86.43 | 71.5 | 61.3 | 84.51 | 69.51 | 60.76 | 78.66 | 57.45 | 146.8 | 78.79 | 104.83 | 94.41 | 116.91 | 28.72 | 167.51 | 78.72 | 38.64 | 67.2 | 59.63 | 76.71 | 137.44 | 117.25 | 130.57 | 47.75 | 64.6 |
| Run1_MDS6 | | 50.04 | 38.76 | 29.25 | 26.64 | 40.29 | 46.25 | 42.66 | 44.17 | 46.37 | 84.81 | 62.2 | 62.64 | 46.39 | 39.58 | 15.46 | 81.78 | 41.35 | 36.7 | 89.46 | 59.35 | 54.67 | 81.31 | 94.02 | 723.54 | 57.45 | 60.58 |
| Run1_MDS8 | | 62.34 | 48.42 | 56.22 | 46.84 | 50.17 | 52.44 | 61.51 | 53.99 | 49.58 | 69.43 | 60.9 | 66.85 | 67.52 | 23.53 | 25.38 | 68.85 | 41.43 | 32.81 | 100.44 | 38.57 | 60.59 | 112.4 | 110.33 | 550.51 | 35.99 | 39.56 |
| Run1_MDS9 | | 76.84 | 50.7 | 54.05 | 38.67 | 59.23 | 56.18 | 76.38 | 78.61 | 68.85 | 77.4 | 70.34 | 89.61 | 90.17 | 19.26 | 17.38 | 204.31 | 60.3 | 38.96 | 129.4 | 59.37 | 75.98 | 77.36 | 127.72 | 216.58 | 52.11 | 62.97 |
| Run2_MDS12 | Lower Risk | 69.93 | 44.8 | 34.59 | 39.57 | 42.9 | 37.4 | 36.32 | 45.94 | 47.3 | 43.7 | 57.19 | 86.38 | 85.54 | 11.25 | 12.59 | 32.64 | 35.09 | 39.27 | 54.45 | 36.78 | 35.37 | 52.29 | 56.5 | 321.37 | 35.15 | 37.95 |
| Run2_MDS14 | | 46.17 | 40.72 | 32.18 | 64.83 | 47.18 | 46.68 | 52.1 | 35.39 | 34.48 | 60.59 | 46.39 | 56.92 | 46.88 | 8.45 | 6.18 | 55.11 | 36.29 | 36.07 | 63.51 | 59.22 | 70.97 | 55.24 | 105 | 391.36 | 45.46 | 68.51 |
| Run2_MDS19 | | 55.39 | 43.42 | 42.19 | 39.02 | 57.56 | 51.3 | 40.46 | 40.85 | 38.9 | 56.04 | 58.56 | 65.79 | 54.46 | 12.04 | 10.26 | 10.05 | 48.2 | 34.47 | 109.36 | 69.81 | 56.8 | 68.76 | 63.45 | 562.04 | 42.5 | 78.95 |
| Run2_MDS20 | | 54.15 | 39.64 | 39.56 | 32.29 | 53.56 | 56.02 | 52.38 | 44.67 | 40.44 | 65.49 | 56.74 | 67.94 | 57.4 | 26.63 | 16.17 | 76.74 | 46.68 | 44.38 | 72.73 | 46.91 | 40.82 | 77.75 | 93.25 | 371.3 | 42.56 | 47.63 |
| Run2_MDS28* | | 92.3 | 42.42 | 36.38 | 39.84 | 50.85 | 44.82 | 41.66 | 49.75 | 47.26 | 72.11 | 70.61 | 108.88 | 110.56 | 18.59 | 15.23 | 106.06 | 45.6 | 32.04 | 41.67 | 52.49 | 55.49 | 86.05 | 88.32 | 603.77 | 42.99 | 62.72 |
| Run2_MDS5 | | 72.69 | 60.12 | 43.6 | 44.13 | 58.29 | 62.54 | 50.19 | 67.43 | 45.62 | 108.48 | 63.45 | 85.2 | 74.51 | 79.11 | 21.78 | 118.14 | 55.31 | 29.63 | 49.77 | 47.3 | 66.1 | 119.81 | 104.7 | 170.71 | 37.18 | 52.43 |
| Run1_ICUS18 | | 71.44 | 40.7 | 39.41 | 27.83 | 47.12 | 59.34 | 46.6 | 42.35 | 39.97 | 84.08 | 79.49 | 86.06 | 73.03 | 39.73 | 35.74 | 99.43 | 58.75 | 35.17 | 102.84 | 73 | 71.08 | 77.54 | 66.15 | 414.91 | 61.02 | 75.23 |
| Run1_ICUS22 | ICUS | 74.3 | 47 | 48.67 | 38.97 | 55.19 | 65.74 | 69.3 | 57.6 | 52.01 | 87.67 | 81.56 | 90.44 | 75.01 | 27.7 | 36.48 | 106.36 | 54.04 | 42.54 | 65.8 | 75.72 | 77.84 | 103.11 | 103.17 | 347.19 | 60.06 | 81.78 |
| Run1_ICUS24 | | 68.44 | 50.05 | 44.2 | 45.04 | 61.36 | 49.19 | 40.33 | 43.39 | 44.73 | 87.83 | 75.56 | 83.37 | 72.11 | 20.4 | 22.74 | 183.59 | 45.52 | 41.05 | 44.7 | 59.22 | 52.6 | 68.47 | 81.35 | 788.17 | 44.32 | 75.28 |
| Run1_Nl-1 | | 69.07 | 40.39 | 38.59 | 31.6 | 63.24 | 48.87 | 52.57 | 46.86 | 43.73 | 92.44 | 72.11 | 78.18 | 68.88 | 30.96 | 33.52 | 116.59 | 41.62 | 38.48 | 51.43 | 68.58 | 74.89 | 73.3 | 125.66 | 1085.1 | 55.31 | 76.54 |
| Run1_Nl-3 | | 54.92 | 43.03 | 40.94 | 38.84 | 48.97 | 41.83 | 33.32 | 36.97 | 35.91 | 77.67 | 55.08 | 61.83 | 54.42 | 44.6 | 32.65 | 76.67 | 39.57 | 26.9 | 57.63 | 47.9 | 49.06 | 56.8 | 83.49 | 146.11 | 35.13 | 57.55 |
| Run1_Nl-4 | | 64.25 | 54.81 | 46.99 | 39.82 | 49.95 | 41.35 | 30.11 | 29.55 | 28 | 82.18 | 68.25 | 73.56 | 65.04 | 49.33 | 30.94 | 90.21 | 36.28 | 33.19 | 81.11 | 44.48 | 48.81 | 56.01 | 83.85 | 140.06 | 37.28 | 52.15 |
| Run1_Nl-6 | | 58.33 | 46.04 | 45.35 | 40.73 | 58.98 | 49.07 | 36.41 | 38.71 | 36.76 | 76.28 | 57.67 | 67.37 | 61.01 | 36.37 | 26.93 | 74.77 | 43.56 | 29.71 | 73.29 | 53.94 | 55.02 | 67.58 | 92.3 | 117.04 | 38.48 | 60.05 |
| Run2_Nl-3 | Normal | 37.64 | 36.09 | 36.82 | 32.48 | 41.74 | 35.66 | 27.97 | 30.2 | 30.19 | 58.47 | 37.28 | 40.87 | 34.44 | 26.59 | 22.51 | 43.23 | 30.96 | 22.35 | 45.33 | 40.01 | 39.21 | 41.6 | 63.76 | 180.89 | 28.6 | 48.95 |
| Run2_Nl-4 | | 44.04 | 41.07 | 46.35 | 36.93 | 44.51 | 35.41 | 23.45 | 22.96 | 22.13 | 63.48 | 45.91 | 49.19 | 41 | 29.5 | 20.74 | 55.39 | 30.59 | 27.64 | 74.22 | 39.12 | 39.06 | 43.47 | 61.49 | 179.55 | 32.74 | 45.64 |
| Run2_Nl-5 | | 34.79 | 34.56 | 35.87 | 30.6 | 41.46 | 30.44 | 23.15 | 22.84 | 22.59 | 60.31 | 42.56 | 41.27 | 32.19 | 18.95 | 10.08 | 38.23 | 30.73 | 22.67 | 57.92 | 35.59 | 36.69 | 37.75 | 54.33 | 193.38 | 28.82 | 42.87 |
| Run2_Nl-6 | | 42.52 | 40.6 | 39.61 | 36.63 | 49.92 | 42.3 | 31.25 | 33.38 | 32.04 | 50.12 | 41.6 | 49.79 | 41.02 | 19.95 | 15.86 | 38.42 | 34.98 | 24.41 | 59.67 | 46.72 | 46.76 | 55.13 | 73.01 | 255.89 | 33.76 | 52.59 |

**Supplemental Table 3**: Median expression of each surface marker on each cell population.

### CD8 Median

| | | Ungated | CD34+CD38low | HSCs | MPP | CMP/GMP | Myelo/Mono-Blasts | ProMonocytes | CD14neg Monocytes | Mature Monocytes | ProMyelocytes | Myelocytes | MetaMyelocytes | MatureGrans | ProErythroblasts | Erythroblasts | LateErythroblasts | PreBcells | Mature Bcells | Plasma Cells | T cells | NK cells | pDCs | Basophils | Platelets | CD8+ T cells | CD8neg T cells |
|---|---|---|---|---|---|---|---|---|---|---|---|---|---|---|---|---|---|---|---|---|---|---|---|---|---|---|---|
| Run1_MDS17 | | 3.72 | 1.87 | 1.99 | 1.72 | 3.03 | 1.49 | 3.27 | 4.39 | 3.92 | 3.99 | 7.95 | 7.36 | 8.13 | 1 | 1.05 | 1.85 | 3.61 | 1.34 | 2.31 | 79.69 | 10.47 | 4.79 | 1.06 | 3.54 | 173.28 | 2.19 |
| Run2_MDS15* | AML/RAEB-T | 1.97 | 3.04 | 2.63 | 2.27 | 2.21 | 2.11 | 1.96 | 1.73 | 1.59 | 2.09 | 1.81 | 1.93 | 2 | 2.34 | 2.06 | 3.19 | 1.83 | 1.42 | 2.75 | 10.86 | 16.57 | 2.13 | 2.71 | 3.81 | 271.61 | 2.61 |
| Run2_MDS4 | | 1.73 | 1.61 | 1.72 | 1.22 | 0.77 | 0.86 | 1.16 | 1.56 | 2.09 | 1.65 | 1.61 | 1.71 | 1.8 | 0.35 | 0.54 | 0.78 | 1.92 | 1.07 | 1.06 | 14.34 | 10.2 | 1.03 | 2.23 | 4.48 | 186.36 | 1.54 |
| Run1_MDS21 | | 2.42 | 1.57 | 1.63 | 1.35 | 1.82 | 1.66 | 2.38 | 2.81 | 3.28 | 2 | 3.04 | 3.42 | 4.25 | 0.82 | 1.52 | 9.72 | 1.71 | 1.84 | 3.74 | 4.48 | 5.35 | 1.98 | 2.71 | 8.93 | 76.33 | 2.42 |
| Run1_MDS3 | | 2.72 | 1.58 | 1.69 | 1.37 | 1.94 | 1.83 | 2.6 | 3.32 | 3.39 | 2.61 | 3.7 | 4.16 | 5.31 | 0.99 | 1.42 | 4.75 | 1.45 | 1.84 | 3.3 | 4.94 | 6.78 | 5.19 | 2.85 | 7.14 | 158.47 | 2.84 |
| Run2_MDS13 | | 2.1 | 1.06 | 1.25 | 0.98 | 1.36 | 1.23 | 2.21 | 3.03 | 3.11 | 2.5 | 3.46 | 3.21 | 3.36 | 0.72 | 0.51 | 1.09 | 0.92 | 0.93 | 1.66 | 4.44 | 4.12 | 2 | 1.69 | 4.43 | 114.99 | 2.03 |
| Run2_MDS16 | | 1.83 | 1.64 | 1.89 | 1.3 | 1.16 | 1.43 | 1.52 | 1.59 | 1.57 | 2.25 | 1.57 | 1.74 | 1.66 | 0.88 | 0.4 | 2.03 | 2.33 | 0.84 | 2.75 | 2.46 | 5.34 | 2.12 | 2.41 | 7.29 | 208.58 | 1.74 |
| Run2_MDS1 | | 1.79 | 1.7 | 2.55 | 1.32 | 1.85 | 1.22 | 1.02 | 1.22 | 1.8 | 0.89 | 0.82 | 2.26 | 2.49 | 0.65 | 0.74 | 0.9 | 2.46 | 1.45 | 1.65 | 2.51 | 4.41 | 1.76 | 3.95 | 5.01 | 171.01 | 1.85 |
| Run2_MDS21 | Higher Risk | 1.89 | 1.03 | 1.13 | 0.93 | 1.29 | 1.06 | 1.34 | 2.12 | 2.38 | 2.75 | 3.22 | 2.86 | 2.98 | 0.35 | 0.4 | 0.85 | 1.04 | 1.32 | 3.19 | 3.84 | 3.6 | 1.06 | 1.81 | 6.45 | 72.66 | 1.94 |
| Run2_MDS26* | | 1.42 | 1.5 | 1.53 | 1.44 | 1.73 | 0.93 | 1.58 | 2.06 | 2.3 | 1.24 | 1.17 | 2.01 | 2.35 | 0.99 | 0.88 | 0.7 | 1.48 | 0.68 | 1.03 | 4.25 | 2.08 | 0.97 | 2.3 | 2.08 | 142.22 | 1.75 |
| Run2_MDS27 | | 1.96 | 2.12 | 2.36 | 1.76 | 1.99 | 1.8 | 1.76 | 1.51 | 1.57 | 2.61 | 2.31 | 1.96 | 1.93 | 1.05 | 0.32 | 0.67 | 4.52 | 1.16 | 2.81 | 62.15 | 2.99 | 2.45 | 2.08 | 5.39 | 201.93 | 1.96 |
| Run2_MDS2 | | 1.72 | 0.69 | 1.14 | 0.52 | 0.73 | 0.59 | 0.43 | 1.08 | 1.26 | 1.38 | 1.21 | 2.35 | 2.61 | 2.14 | 1.96 | 3.31 | 1.55 | 2.58 | 0.03 | 1.44 | 2.8 | 0.25 | 1.37 | 5.18 | 223.22 | 1.23 |
| Run2_MDS3 | | 2.56 | 1.18 | 1.3 | 1.06 | 1.49 | 1.31 | 2.01 | 3.41 | 4.1 | 3.35 | 4.79 | 4.7 | 5.65 | 0.6 | 0.68 | 0.63 | 1.2 | 1.31 | 1.84 | 4.15 | 3.72 | 2.32 | 1.88 | 5.33 | 146.49 | 2.25 |
| Run1_MDS23 | | 3.89 | 2.71 | 2.48 | 2.39 | 3.16 | 2.35 | 3.33 | 3.4 | 3.28 | 4.69 | 4.27 | 4.39 | 4.38 | 1.52 | 1.79 | 4.73 | 3.51 | 1.82 | 3.12 | 4.49 | 2.87 | 3.05 | 3.65 | 6.79 | 157.56 | 2.89 |
| Run1_MDS25 | | 3.14 | 6.22 | 3.63 | 4.27 | 0.79 | 2.07 | 1.72 | 1.7 | 1.89 | 3.04 | 2.87 | 2.91 | 3.1 | 1.14 | 2.2 | 5.68 | 3.18 | 2.31 | 2.85 | 18.77 | 4.09 | 2.89 | 3.92 | 7.5 | 268.11 | 2.73 |
| Run1_MDS5 | | 3.66 | 3.9 | 1.99 | 3.11 | 2.31 | 3.31 | 2.29 | 3.61 | 4 | 4.57 | 3.01 | 3.66 | 3.83 | 4.42 | 1.63 | 5.32 | 4.1 | 1.76 | 3.37 | 3.91 | 3.27 | 5.5 | 4.16 | 4.37 | 246.1 | 2.57 |
| Run1_MDS6 | | 2.85 | 2.37 | 1.47 | 1.36 | 2 | 2.4 | 2.7 | 2.9 | 3.05 | 3.44 | 3.04 | 3.02 | 2.57 | 1.69 | 0.72 | 4.54 | 2.63 | 1.64 | 4.27 | 7.99 | 59.04 | 3.33 | 3.66 | 9.94 | 206.12 | 2.72 |
| Run1_MDS8 | | 2.94 | 3.01 | 3.72 | 2.15 | 2.27 | 2.18 | 2.68 | 2.63 | 2.56 | 2.87 | 2.88 | 2.84 | 2.95 | 0.86 | 1.09 | 3.89 | 2.33 | 1.37 | 1.49 | 2.75 | 5.87 | 3.95 | 4.13 | 8.97 | 70.59 | 1.7 |
| Run1_MDS9 | | 3.64 | 2.64 | 2.85 | 2.06 | 3.18 | 2.64 | 3.24 | 2.95 | 3.27 | 3.35 | 3.15 | 3.49 | 3.55 | 0.85 | 0.89 | 7.34 | 5.88 | 1.76 | 6.18 | 5.57 | 19.34 | 3.3 | 4.34 | 6.11 | 126.53 | 2.68 |
| Run2_MDS12 | Lower Risk | 2.62 | 2.01 | 2.26 | 1.59 | 2.07 | 1.83 | 2.09 | 2.41 | 2.41 | 1.97 | 2.45 | 3.13 | 3.19 | 0.44 | 0.48 | 1.37 | 2.28 | 1.57 | 1.31 | 4.15 | 1.68 | 1.95 | 1.9 | 5.9 | 118.91 | 1.49 |
| Run2_MDS14 | | 2.23 | 2.49 | 0.86 | 2.09 | 2.23 | 1.83 | 2.03 | 2.29 | 2.42 | 2.54 | 2.21 | 2.71 | 2.22 | 0.49 | 0.2 | 2.51 | 1.68 | 1.26 | 2.06 | 4.44 | 3.52 | 1.76 | 3.33 | 4.41 | 218.96 | 2.28 |
| Run2_MDS19 | | 2.18 | 2.29 | 2.54 | 1.98 | 2.17 | 2.01 | 1.76 | 1.76 | 1.73 | 2.29 | 2.37 | 2.4 | 2.13 | 0.56 | 0.39 | 0.59 | 2.95 | 1.39 | 3.55 | 3.66 | 2.93 | 2.5 | 2.02 | 8.93 | 210.2 | 2.55 |
| Run2_MDS20 | | 2.23 | 1.92 | 2.39 | 2.02 | 1.96 | 1.9 | 1.98 | 1.74 | 1.96 | 2.42 | 2.24 | 2.4 | 2.28 | 0.66 | 0.86 | 2.81 | 2.37 | 1.95 | 4.58 | 1.97 | 1.8 | 2.67 | 2.97 | 6.95 | 181.67 | 1.66 |
| Run2_MDS28* | | 3.75 | 2.32 | 2.18 | 2.12 | 1.95 | 2.26 | 2.22 | 2.57 | 2.53 | 3.1 | 2.99 | 4.04 | 4 | 0.83 | 0.75 | 3.33 | 3.3 | 1.24 | 2.25 | 9.1 | 4.25 | 3.12 | 2.63 | 8.53 | 130.65 | 2.4 |
| Run2_MDS5 | | 2.51 | 3.66 | 1.94 | 1.72 | 2.68 | 2.34 | 1.47 | 2.9 | 2.84 | 3.09 | 1.89 | 2.45 | 2.66 | 2.89 | 1.06 | 3.48 | 2.7 | 1.24 | 1.79 | 3.48 | 2.27 | 1.96 | 3.4 | 3.54 | 212.24 | 1.91 |
| Run1_ICUS18 | | 3.36 | 2.92 | 1.47 | 2.92 | 3.33 | 2.47 | 2.48 | 2.07 | 2.42 | 3.76 | 3.66 | 3.52 | 3.22 | 1.49 | 1.67 | 3.81 | 3.7 | 1.48 | 3.42 | 3.77 | 5.93 | 3.03 | 2.66 | 8.95 | 210.9 | 2.97 |
| Run1_ICUS22 | ICUS | 3.17 | 2.55 | 2.49 | 2.3 | 2.53 | 2.63 | 2.85 | 2.25 | 2.36 | 3.7 | 3.41 | 3.51 | 3.04 | 1.13 | 1.5 | 3.98 | 2.93 | 1.82 | 2.99 | 4.95 | 4.64 | 3.79 | 3.65 | 7.71 | 130.19 | 3.26 |
| Run1_ICUS24 | | 3.26 | 2.74 | 3.14 | 2.53 | 3.04 | 2.4 | 2.36 | 2.26 | 2.25 | 3.58 | 3.46 | 3.42 | 3.09 | 0.79 | 0.95 | 6.44 | 2.3 | 1.83 | 2.21 | 8.51 | 3.34 | 2.74 | 3.43 | 10.74 | 113.17 | 3.21 |
| Run1_Nl-1 | | 3.01 | 2.08 | 1.93 | 1.85 | 2.54 | 2.24 | 2.56 | 2.35 | 2.21 | 3.58 | 3.09 | 3.16 | 2.88 | 1.01 | 1.46 | 4.61 | 2.02 | 1.66 | 2.71 | 5.24 | 3.77 | 2.96 | 4.51 | 12.49 | 178.21 | 2.94 |
| Run1_Nl-3 | | 3.23 | 2.96 | 2.57 | 2.72 | 2.54 | 2.79 | 4.6 | 5.92 | 4.06 | 3.46 | 2.99 | 3.3 | 2.98 | 1.95 | 1.56 | 3.97 | 2.69 | 1.08 | 3.18 | 5.52 | 5.01 | 2.5 | 3.44 | 4.92 | 189.51 | 2.39 |
| Run1_Nl-4 | | 3.59 | 3.19 | 2.95 | 2.96 | 2.86 | 2.93 | 5.63 | 6.42 | 4.43 | 3.62 | 3.41 | 3.63 | 3.42 | 1.71 | 1.37 | 4.2 | 2.52 | 1.39 | 3.66 | 6.72 | 4.59 | 2.53 | 3.01 | 4.75 | 141.59 | 2.35 |
| Run1_Nl-6 | | 3.12 | 2.78 | 3.04 | 2.08 | 2.98 | 3.03 | 4.64 | 5.81 | 4.5 | 3.42 | 3.17 | 3.54 | 3.23 | 1.28 | 1.18 | 3.5 | 2.18 | 1.19 | 3.44 | 3.94 | 4.22 | 2.82 | 3.63 | 4.75 | 170.86 | 2.47 |
| Run2_Nl-3 | Normal | 2.16 | 2 | 1.81 | 1.76 | 2.15 | 2.15 | 3.21 | 4.12 | 3.04 | 2.32 | 1.94 | 2.16 | 1.99 | 0.95 | 0.8 | 2.24 | 1.9 | 0.78 | 1.75 | 4.4 | 3.3 | 1.42 | 2.12 | 4.79 | 171.13 | 1.77 |
| Run2_Nl-4 | | 2.53 | 2.65 | 3.02 | 2.09 | 1.98 | 2.36 | 4.08 | 4.74 | 3.32 | 2.5 | 2.21 | 2.52 | 2.4 | 0.97 | 0.83 | 2.63 | 1.94 | 1.04 | 2.63 | 6.29 | 3.57 | 1.57 | 2.33 | 4.1 | 125.13 | 1.81 |
| Run2_Nl-5 | | 2.38 | 2.28 | 2.26 | 1.9 | 1.85 | 2 | 3.64 | 4.66 | 3.67 | 2.34 | 2.07 | 2.45 | 2.29 | 0.74 | 0.4 | 2.39 | 1.7 | 0.84 | 2.57 | 7.61 | 6.82 | 1.62 | 1.99 | 5.37 | 195.22 | 1.6 |
| Run2_Nl-6 | | 2.08 | 2 | 2.62 | 1.76 | 2.19 | 2.28 | 3.66 | 4.15 | 3.48 | 2.08 | 1.97 | 2.34 | 2.09 | 0.81 | 0.6 | 1.57 | 1.51 | 0.92 | 2.55 | 3.12 | 2.97 | 1.99 | 2.5 | 4.6 | 152.97 | 1.84 |

**Supplemental Table 3**: Median expression of each surface marker on each cell population.

### CD34 Median

| | | Ungated | CD34+CD38low | HSCs | MPP | CMP/GMP | Myelo/Mono-Blasts | ProMonocytes | CD14neg Monocytes | Mature Monocytes | ProMyelocytes | Myelocytes | MetaMyelocytes | MatureGrans | ProErythroblasts | Erythroblasts | LateErythroblasts | PreBcells | Mature Bcells | Plasma Cells | T cells | NK cells | pDCs | Basophils | Platelets | CD8+ T cells | CD8neg T cells |
|---|---|---|---|---|---|---|---|---|---|---|---|---|---|---|---|---|---|---|---|---|---|---|---|---|---|---|---|
| Run1_MDS17 | | 1.31 | 50.67 | 61.19 | 48.38 | 64.27 | 13.48 | 2.03 | 1.2 | 0.95 | 0.95 | 0.95 | 0.65 | 0.9 | 0.52 | -0.07 | 0.32 | 2.2 | 0.1 | 1.5 | 0.34 | 1.08 | 3.82 | 1.53 | 2.2 | 0.68 | -0.1 |
| Run2_MDS15* | AML/RAEB-T | 1.08 | 293.81 | 263.47 | 237.16 | 75.4 | 7.55 | 1.44 | 0.6 | 0.62 | 1.9 | 0.72 | 0.49 | 0.35 | 31.31 | 0.83 | 1.63 | 0.91 | 0.68 | 3.61 | 0.41 | 1.82 | 31.15 | 1.76 | 18.53 | 0.67 | 0.14 |
| Run2_MDS4 | | 0.1 | 109.73 | 187.52 | 97.36 | 23.89 | 1.21 | 0.28 | 0.27 | 0.28 | -0.01 | -0.16 | -0.08 | 0.01 | -0.01 | -0.13 | -0.07 | 0.44 | 0.35 | 0.19 | -0.04 | 0.55 | 0.22 | -0.05 | 0.95 | 0.3 | -0.21 |
| Run1_MDS21 | | 0.56 | 49.33 | 52.65 | 47.41 | 63.49 | 54.87 | 5.19 | 1.73 | 2.34 | 0.04 | 0.21 | 0.21 | 0.6 | -0.07 | 0.26 | 1.58 | -0.03 | -0.02 | 0.13 | -0.03 | 1.68 | 14.13 | 0.17 | 1.13 | 0.59 | -0.19 |
| Run1_MDS3 | | 0.29 | 70.65 | 80.36 | 65.21 | 86.03 | 70.51 | 6.91 | 1.34 | 2.91 | -0.08 | 0 | 0.14 | 0.61 | -0.04 | 0.09 | 0.58 | -0.14 | 0 | 0.93 | -0.03 | 1.5 | 1.69 | 0.11 | 0.87 | 0.6 | -0.19 |
| Run2_MDS13 | | 0.34 | 91.9 | 105.28 | 83.01 | 82.81 | 64.29 | 0.62 | 0.62 | 0.72 | -0.01 | 0.04 | 0.01 | 0.16 | 0.44 | -0.16 | -0.01 | -0.07 | -0.11 | 0.95 | -0.03 | 0.86 | 3.41 | 0.39 | 0.39 | 0.42 | -0.17 |
| Run2_MDS16 | | -0.05 | 134.03 | 145.94 | 129.64 | 130.97 | 0.85 | 0.19 | 0.28 | 0.33 | -0.12 | -0.2 | -0.17 | -0.11 | 36.38 | -0.23 | -0.02 | 0.25 | -0.16 | 0.59 | -0.1 | 0.84 | 0.84 | 0.01 | 0.95 | 0.42 | -0.17 |
| Run2_MDS1 | | 0 | 30.62 | 52.43 | 28.37 | 27.4 | 1.74 | 0.24 | 0.1 | 0.29 | -0.22 | -0.21 | -0.05 | 0.15 | -0.12 | -0.14 | -0.19 | 0.28 | -0.09 | 0.63 | -0.18 | 1.28 | 0 | -0.05 | 0.25 | 0.28 | -0.24 |
| Run2_MDS21 | Higher Risk | 0.3 | 54.42 | 59.94 | 51.4 | 70.7 | 53.08 | 0.37 | 0.45 | 0.67 | -0.06 | -0.05 | -0.01 | 0.12 | 0.22 | -0.18 | -0.07 | 0.08 | -0.04 | 0.92 | -0.02 | 0.82 | 12.6 | 0.07 | 0.8 | 0.45 | -0.14 |
| Run2_MDS26* | | -0.18 | 38.59 | 42.95 | 37.24 | 35.94 | 3.92 | 0.44 | 0.44 | 1.14 | -0.22 | -0.22 | -0.18 | -0.15 | 2.54 | -0.22 | -0.23 | 0 | -0.23 | 0.79 | -0.08 | 0.37 | 0.62 | -0.07 | -0.18 | 0.29 | -0.21 |
| Run2_MDS27 | | 0.36 | 170.12 | 197.48 | 146.59 | 76.36 | 9.47 | 0.62 | 0.48 | 0.6 | 0.16 | -0.08 | 0 | 0.26 | 14.21 | -0.21 | -0.15 | 0.87 | 0.18 | 1.93 | 0.18 | 0.64 | 0.36 | 0.49 | 5.94 | 0.45 | -0.1 |
| Run2_MDS2 | | 0.03 | 93.84 | 104.13 | 93.19 | 84.93 | 34.98 | 0.33 | -0.03 | 0.59 | 1.19 | -0.22 | -0.11 | -0.05 | 0.35 | 0.41 | 0.55 | 0.22 | 1.18 | -0.39 | -0.25 | 0.23 | -0.24 | -0.13 | 0.37 | 0.24 | -0.28 |
| Run2_MDS3 | | 0.24 | 90.74 | 109.1 | 80.17 | 106.77 | 71.22 | 0.39 | 0.42 | 1.01 | 0 | 0.14 | 0.14 | 0.39 | 0.58 | -0.13 | -0.15 | -0.08 | -0.09 | 0.54 | -0.04 | 0.89 | 1.71 | 0.5 | 0.72 | 0.49 | -0.17 |
| Run1_MDS23 | | 0.26 | 65.77 | 99.92 | 57.14 | 47.25 | 0.55 | 0.25 | 0.23 | 0.27 | 0.4 | 0.31 | 0.28 | 0.37 | -0.1 | -0.05 | 0.37 | 0.31 | -0.24 | 0.41 | -0.06 | 0.92 | 0.34 | -0.08 | 0.46 | 0.55 | -0.18 |
| Run1_MDS25 | | 0.36 | 23.34 | 154.42 | 30.95 | 54.29 | 0.48 | 0.39 | 0.38 | 0.44 | 0.34 | 0.05 | 0.26 | 0.39 | -0.04 | 0.09 | 1.15 | 0.27 | -0.1 | 0.72 | 0.13 | 0.48 | 0.49 | -0.05 | 0.39 | 0.58 | -0.19 |
| Run1_MDS5 | | 0.49 | 58.42 | 136.51 | 37.32 | 80.16 | 3.07 | 0.74 | 0.84 | 0.93 | 0.87 | 0.27 | 0.39 | 0.5 | 0.84 | 0 | 0.47 | 0.75 | -0.19 | 0.57 | -0.06 | 3.72 | 0.77 | 0.88 | 0 | 0.57 | -0.19 |
| Run1_MDS6 | | 0.25 | 91.69 | 69.78 | 46.94 | 61.42 | 4.75 | 1.28 | 1.39 | 1.08 | 0.67 | 0.41 | 0.3 | 0.16 | 1.45 | -0.2 | 0.74 | 0.26 | -0.13 | 1.01 | 0.03 | 1.24 | 1.02 | 0.17 | 2.02 | 0.59 | -0.22 |
| Run1_MDS8 | | -0.03 | 132.23 | 135.77 | 122.6 | 98.69 | 2.12 | 0.42 | 0.31 | 0.38 | -0.02 | -0.14 | -0.11 | -0.05 | 0.07 | -0.1 | 0.38 | -0.11 | -0.2 | 0.47 | -0.22 | 0.71 | 1.6 | 0.59 | 1.19 | 0.17 | -0.3 |
| Run1_MDS9 | | 0.17 | 84.33 | 117.16 | 49.15 | 80.26 | 15.65 | 0.31 | 0.22 | 0.4 | -0.02 | -0.07 | 0.09 | 0.19 | 0.06 | -0.11 | 0.59 | 0.41 | -0.16 | 0.72 | -0.06 | 0.83 | 0.5 | 0.51 | 0.27 | 0.46 | -0.24 |
| Run2_MDS12 | Lower Risk | -0.07 | 52.28 | 90.71 | 64.2 | 76.07 | 0.94 | 0.64 | 0.56 | 0.7 | -0.14 | -0.15 | -0.1 | -0.07 | 0.07 | -0.22 | -0.13 | 0.39 | -0.13 | 0.32 | -0.1 | 1.14 | 0.8 | 0.04 | 1.1 | 0.33 | -0.22 |
| Run2_MDS14 | | -0.03 | 48.79 | 46.51 | 57.62 | 39.22 | 0.31 | 0.09 | 0.26 | 0.18 | -0.08 | -0.13 | -0.08 | -0.02 | -0.02 | -0.22 | 0.06 | -0.03 | -0.17 | 0.23 | -0.08 | 1.17 | 0.77 | 0.14 | 0.95 | 0.33 | -0.22 |
| Run2_MDS19 | | -0.05 | 164.54 | 178.84 | 156.45 | 102.49 | 1.33 | 0.25 | 0.07 | 0.12 | -0.12 | -0.14 | -0.1 | -0.01 | 0.29 | -0.23 | -0.2 | 0.25 | -0.16 | 0.35 | -0.11 | 0.89 | 0.9 | 0.52 | 3.59 | 0.44 | -0.2 |
| Run2_MDS20 | | -0.07 | 88.27 | 161.28 | 66.97 | 121.49 | 0.33 | 0.12 | 0.06 | 0.16 | -0.08 | -0.12 | -0.09 | -0.03 | 0.51 | -0.2 | 0 | 0.04 | -0.05 | 1.59 | -0.19 | 0.33 | 0.29 | 0.19 | 0.5 | 0.33 | -0.22 |
| Run2_MDS28* | | 0.07 | 88.11 | 127.52 | 75.24 | 88.59 | 1.14 | 0.51 | 0.33 | 0.6 | 0.05 | -0.03 | 0 | 0 | 1.3 | -0.22 | 0.01 | 0.28 | -0.3 | -0.09 | -0.04 | 0.67 | 0.67 | 0.14 | 0.67 | 0.36 | -0.2 |
| Run2_MDS5 | | 0.43 | 88.55 | 207.49 | 65.5 | 61.26 | 2.28 | 0.61 | 0.64 | 1.05 | 0.35 | 0.14 | 0.32 | 0.47 | 0.77 | -0.13 | 0.4 | 0.67 | -0.19 | 0.33 | -0.1 | 3.11 | 0.7 | 0.72 | -0.01 | 0.4 | -0.21 |
| Run1_ICUS18 | | 0.3 | 123.49 | 192.06 | 85.63 | 140.05 | 0.61 | 0.32 | 0.39 | 0.37 | 0.18 | -0.03 | 0.16 | 0.35 | -0.05 | -0.16 | 0.54 | 0.14 | -0.22 | 0.46 | -0.13 | 0.61 | 0.45 | -0.07 | 2.66 | 0.62 | -0.21 |
| Run1_ICUS22 | ICUS | 0.03 | 109.25 | 131.63 | 86.13 | 68.17 | 0.76 | 0.34 | 0.16 | 0.18 | -0.06 | -0.13 | -0.07 | 0.06 | -0.07 | -0.08 | 0.25 | 0.12 | -0.19 | 0.02 | -0.08 | 0.28 | 0.39 | 0.21 | 1.22 | 0.42 | -0.18 |
| Run1_ICUS24 | | 0.05 | 131.23 | 151.83 | 102.25 | 59.7 | 2.25 | 0.48 | 0.29 | 0.3 | 0.04 | -0.08 | -0.06 | 0.09 | -0.05 | -0.19 | 0.77 | 0.01 | -0.19 | 0.39 | 0.01 | 0.4 | 1.64 | 0.34 | 1.67 | 0.42 | -0.17 |
| Run1_Nl-1 | | -0.02 | 119.58 | 140.16 | 103.09 | 39.69 | 0.97 | 0.3 | 0.26 | 0.3 | -0.07 | -0.14 | -0.07 | 0 | -0.04 | -0.12 | 0.48 | -0.06 | -0.24 | 0.4 | -0.06 | 0.57 | 0.87 | 0.18 | 1.95 | 0.53 | -0.22 |
| Run1_Nl-3 | | 0.21 | 182.12 | 182.03 | 156.53 | 95.03 | 3.14 | 0.83 | 0.71 | 0.51 | 0.26 | -0.03 | 0.15 | 0.23 | 0.77 | -0.1 | 0.47 | 0.22 | -0.24 | 0.56 | 0.05 | 0.91 | 1.07 | 0.2 | 2.1 | 0.63 | -0.2 |
| Run1_Nl-4 | | 0.27 | 182.95 | 220.43 | 162.18 | 105.12 | 2.24 | 0.67 | 0.54 | 0.47 | 0.57 | 0.1 | 0.22 | 0.28 | 0.67 | -0.11 | 0.48 | -0.06 | -0.18 | 0.9 | 0.07 | 0.66 | 1.29 | 0.48 | 1.9 | 0.55 | -0.18 |
| Run1_Nl-6 | | 0.26 | 142.1 | 156.47 | 124.4 | 67.6 | 3.48 | 0.81 | 0.6 | 0.57 | 0.52 | 0.18 | 0.22 | 0.27 | 0.26 | -0.14 | 0.42 | 0.25 | -0.2 | 0.78 | -0.07 | 0.65 | 0.84 | 0.68 | 4 | 0.54 | -0.19 |
| Run2_Nl-3 | Normal | 0.07 | 161.86 | 194.12 | 160.74 | 80.98 | 2.12 | 0.53 | 0.39 | 0.39 | -0.04 | -0.09 | 0.05 | 0.18 | 0.61 | -0.16 | 0.25 | 0.1 | -0.24 | 0.47 | -0.05 | 0.65 | 0.8 | 0.02 | 1.97 | 0.39 | -0.2 |
| Run2_Nl-4 | | 0.17 | 185.08 | 278.46 | 152.75 | 93.38 | 1.6 | 0.52 | 0.44 | 0.41 | 0.02 | -0.03 | 0.17 | 0.28 | 0.62 | -0.17 | 0.27 | 0.1 | -0.16 | 0.78 | -0.02 | 0.5 | 1.15 | 0.31 | 2.57 | 0.31 | -0.17 |
| Run2_Nl-5 | | 0.16 | 157.97 | 205.4 | 123.43 | 64.56 | 1.21 | 0.48 | 0.44 | 0.44 | -0.05 | -0.09 | 0.12 | 0.29 | 0.61 | -0.21 | 0.25 | 0.05 | -0.18 | 0.68 | -0.03 | 0.61 | 1.09 | 0.25 | 2.04 | 0.4 | -0.23 |
| Run2_Nl-6 | | 0.12 | 128.71 | 148.83 | 116.94 | 81.68 | 2.34 | 0.51 | 0.5 | 0.49 | -0.06 | -0.08 | 0.02 | 0.34 | 0.36 | -0.18 | 0.02 | 0.13 | -0.19 | 0.6 | -0.08 | 0.48 | 0.67 | 0.25 | 4.47 | 0.33 | -0.17 |

**Supplemental Table 3**: Median expression of each surface marker on each cell population.

### CD117 Median

| | | Ungated | CD34+CD38low | HSCs | MPP | CMP/GMP | Myelo/Mono-Blasts | ProMonocytes | CD14neg Monocytes | Mature Monocytes | ProMyelocytes | Myelocytes | MetaMyelocytes | MatureGrans | ProErythroblasts | Erythroblasts | LateErythroblasts | PreBcells | Mature Bcells | Plasma Cells | T cells | NK cells | pDCs | Basophils | Platelets | CD8+ T cells | CD8neg T cells |
|---|---|---|---|---|---|---|---|---|---|---|---|---|---|---|---|---|---|---|---|---|---|---|---|---|---|---|---|
| Run1_MDS17 | | 0.33 | 7.38 | 7.9 | 7.01 | 13.81 | 4.64 | -0.04 | -0.01 | -0.02 | -0.19 | -0.03 | -0.02 | 0.22 | 0.27 | -0.17 | -0.16 | 0.78 | -0.15 | -0.07 | -0.3 | -0.27 | 0.6 | -0.56 | -0.11 | -0.29 | -0.32 |
| Run2_MDS15* | AML/RAEB-T | 0.09 | 13.41 | 11.24 | 10.61 | 21.21 | 3.17 | -0.07 | -0.09 | -0.04 | 0.04 | -0.17 | -0.17 | -0.17 | 12.6 | 0.31 | 0.73 | -0.04 | -0.08 | 0.14 | -0.25 | -0.13 | 6.77 | 1.03 | 4.91 | -0.26 | -0.24 |
| Run2_MDS4 | | -0.13 | 8.87 | 6.72 | 3.65 | -0.32 | 0.63 | -0.1 | -0.09 | 0.02 | -0.13 | -0.23 | -0.22 | -0.18 | 0.24 | -0.07 | -0.21 | -0.06 | -0.11 | -0.13 | -0.28 | -0.09 | -0.15 | -0.14 | 0.21 | -0.29 | -0.28 |
| Run1_MDS21 | | -0.03 | 7.33 | 8.03 | 7.07 | 8.69 | 7.69 | 0.88 | -0.1 | 0.16 | -0.27 | -0.23 | -0.19 | 0.06 | -0.01 | 0.13 | 0.32 | -0.28 | -0.17 | -0.04 | -0.3 | -0.1 | 0.68 | 0.35 | 0.15 | -0.27 | -0.31 |
| Run1_MDS3 | | -0.09 | 5.46 | 6.01 | 5.18 | 7.06 | 6.24 | 1.5 | -0.08 | 1.08 | -0.24 | -0.2 | -0.14 | 0.28 | -0.05 | -0.01 | 0.09 | -0.25 | -0.13 | 0.73 | -0.31 | -0.2 | 0.83 | -0.01 | -0.02 | -0.3 | -0.31 |
| Run2_MDS13 | | -0.06 | 3.43 | 3.78 | 3.22 | 6.37 | 5.09 | -0.04 | -0.01 | 0.17 | -0.21 | -0.17 | -0.14 | -0.07 | 0.61 | -0.19 | -0.21 | -0.24 | -0.3 | -0.03 | -0.28 | -0.16 | 0.18 | 0.24 | -0.09 | -0.26 | -0.29 |
| Run2_MDS16 | | -0.2 | 5.17 | 5.29 | 4.23 | 4.36 | 0.3 | -0.17 | -0.09 | -0.08 | -0.21 | -0.28 | -0.25 | -0.22 | 2.1 | -0.19 | -0.16 | -0.23 | -0.25 | -0.02 | -0.28 | -0.19 | 0.15 | -0.17 | -0.01 | -0.26 | -0.28 |
| Run2_MDS1 | | -0.19 | 8.76 | 5.98 | 9.66 | 15.14 | 3.46 | -0.1 | -0.22 | -0.11 | -0.21 | -0.35 | -0.24 | -0.2 | 1.06 | -0.1 | -0.23 | -0.14 | -0.24 | -0.03 | -0.29 | -0.22 | -0.22 | 0.06 | -0.12 | -0.3 | -0.29 |
| Run2_MDS21 | Higher Risk | -0.07 | 7.66 | 8.2 | 7.58 | 9.66 | 7.45 | -0.06 | -0.1 | -0.02 | -0.21 | -0.2 | -0.17 | -0.13 | 0.76 | -0.19 | -0.23 | -0.26 | -0.26 | -0.1 | -0.29 | -0.28 | 7.24 | 0.25 | -0.01 | -0.27 | -0.3 |
| Run2_MDS26* | | -0.25 | 9.73 | 10.01 | 8.76 | 19.84 | 4.22 | 0.07 | -0.16 | -0.06 | -0.3 | -0.32 | -0.27 | -0.22 | 2.82 | -0.22 | -0.27 | -0.08 | -0.28 | -0.07 | -0.3 | -0.19 | 0.16 | 0.37 | -0.27 | -0.3 | -0.3 |
| Run2_MDS27 | | -0.12 | 2.42 | 2.66 | 1.96 | 2.06 | 0.9 | 0.12 | -0.12 | -0.12 | -0.16 | -0.23 | -0.21 | -0.17 | 0.4 | -0.22 | -0.25 | -0.02 | -0.22 | 0.12 | -0.27 | -0.24 | -0.08 | -0.03 | 0.43 | -0.27 | -0.27 |
| Run2_MDS2 | | -0.1 | 2.86 | 4.06 | 2.27 | 3.82 | 2.07 | -0.18 | -0.22 | 0 | 0.46 | -0.24 | -0.17 | -0.12 | 1.69 | 0.31 | 0 | -0.13 | 0.2 | -0.66 | -0.28 | -0.23 | 0.57 | -0.07 | -0.1 | -0.18 | -0.28 |
| Run2_MDS3 | | -0.07 | 5.73 | 6.46 | 5.5 | 8.32 | 6.54 | -0.09 | -0.14 | -0.14 | -0.18 | -0.09 | -0.06 | 0.09 | 1.8 | -0.19 | -0.26 | -0.26 | -0.13 | -0.02 | -0.29 | -0.14 | 0.35 | 0.15 | -0.06 | -0.28 | -0.29 |
| Run1_MDS23 | | -0.16 | 5.41 | 6.88 | 5.18 | 10.03 | 0.36 | -0.05 | -0.01 | 0.09 | -0.17 | -0.21 | -0.18 | -0.16 | 0.33 | -0.01 | 0.06 | -0.11 | -0.23 | -0.09 | -0.3 | -0.25 | -0.05 | -0.18 | -0.04 | -0.29 | -0.3 |
| Run1_MDS25 | | -0.2 | 3.34 | 3.31 | 4.86 | 4.9 | -0.05 | -0.17 | -0.17 | -0.14 | -0.28 | -0.28 | -0.25 | -0.2 | 0 | 0.18 | 0.11 | -0.17 | -0.09 | 0.33 | -0.3 | -0.15 | 0.1 | -0.15 | -0.12 | -0.3 | -0.3 |
| Run1_MDS5 | | -0.05 | 2.03 | 1.4 | 1.92 | 1.4 | 0.71 | -0.02 | 0.34 | 0.4 | 0.03 | -0.17 | -0.06 | 0.02 | 0.4 | -0.02 | 0.07 | 0.15 | -0.28 | 0.38 | -0.26 | -0.09 | 0.49 | -0.13 | -0.17 | -0.22 | -0.27 |
| Run1_MDS6 | | -0.15 | 5.5 | 4.1 | 1.79 | 4.67 | 1.73 | 0.14 | 0.12 | 0.13 | -0.14 | -0.18 | -0.16 | -0.16 | 0.99 | -0.11 | 0.07 | -0.2 | -0.25 | 0.58 | -0.29 | -0.19 | 0.44 | 0.35 | 0.28 | -0.27 | -0.3 |
| Run1_MDS8 | | -0.2 | 5.97 | 6.73 | 3.45 | 5.4 | 0.96 | -0.02 | -0.07 | -0.09 | -0.23 | -0.26 | -0.24 | -0.2 | 0.83 | 0.05 | 0.04 | -0.22 | -0.28 | -0.04 | -0.34 | -0.15 | 0.38 | -0.07 | 0.03 | -0.31 | -0.35 |
| Run1_MDS9 | | -0.18 | 4.13 | 4.79 | 2.8 | 4.53 | 1.85 | -0.04 | -0.02 | -0.06 | -0.22 | -0.24 | -0.21 | -0.16 | 0.32 | 0.01 | 0.02 | -0.1 | -0.27 | -0.15 | -0.33 | -0.2 | 0.04 | 0.09 | -0.15 | -0.3 | -0.34 |
| Run2_MDS12 | Lower Risk | -0.18 | 12.03 | 16.05 | 11.57 | 28.74 | 0.31 | -0.12 | -0.09 | -0.05 | -0.2 | -0.25 | -0.18 | -0.15 | 0.49 | -0.19 | -0.19 | -0.13 | -0.19 | -0.04 | -0.31 | -0.18 | 0.15 | -0.09 | -0.05 | -0.29 | -0.32 |
| Run2_MDS14 | | -0.18 | 3.19 | 2.4 | 0.88 | 1.36 | -0.15 | -0.13 | -0.07 | -0.05 | -0.16 | -0.23 | -0.17 | -0.19 | 0.22 | -0.18 | -0.09 | -0.25 | -0.21 | 0.17 | -0.26 | -0.15 | 0.16 | -0.19 | -0.18 | -0.25 | -0.26 |
| Run2_MDS19 | | -0.22 | 4.49 | 4.34 | 4.01 | 7.5 | 0.51 | -0.14 | -0.18 | -0.13 | -0.23 | -0.25 | -0.22 | -0.21 | 0.3 | -0.19 | -0.28 | -0.19 | -0.24 | -0.35 | -0.28 | -0.18 | 0.06 | -0.23 | 0 | -0.3 | -0.27 |
| Run2_MDS20 | | -0.19 | 1.96 | 2.36 | 1.96 | 5.11 | 0.08 | -0.03 | -0.05 | 0.01 | -0.18 | -0.21 | -0.2 | -0.16 | 1.83 | -0.11 | -0.14 | -0.17 | -0.13 | 0.43 | -0.29 | -0.18 | -0.12 | 0 | -0.02 | -0.28 | -0.3 |
| Run2_MDS28* | | -0.07 | 7.63 | 8.54 | 7.31 | 10.38 | 0.82 | 0.12 | 0.06 | 0.22 | -0.11 | -0.15 | -0.08 | -0.05 | 10.87 | -0.23 | -0.09 | -0.06 | -0.19 | 0.01 | -0.29 | -0.21 | 0.24 | 0.09 | 0 | -0.29 | -0.29 |
| Run2_MDS5 | | -0.1 | 3.14 | 0.69 | 0.9 | 2.03 | 0.55 | -0.09 | 0.14 | 0.24 | -0.18 | -0.2 | -0.09 | -0.05 | 0.48 | -0.08 | -0.01 | -0.06 | -0.22 | 0.43 | -0.27 | -0.1 | -0.1 | -0.08 | -0.17 | -0.27 | -0.26 |
| Run1_ICUS18 | | -0.17 | 3.52 | 4.09 | 3.83 | 3.43 | 0.15 | -0.06 | -0.05 | -0.06 | -0.18 | -0.18 | -0.16 | -0.16 | 0.23 | -0.08 | -0.02 | -0.16 | -0.24 | 0.02 | -0.29 | -0.21 | -0.02 | -0.17 | 0.19 | -0.26 | -0.3 |
| Run1_ICUS22 | ICUS | -0.16 | 2.68 | 3.16 | 1.99 | 3.25 | 0.38 | -0.03 | -0.04 | 0.02 | -0.23 | -0.21 | -0.18 | -0.16 | 0.13 | 0.03 | -0.06 | -0.18 | -0.19 | -0.2 | -0.27 | -0.16 | -0.03 | -0.05 | -0.15 | -0.27 | -0.26 |
| Run1_ICUS24 | | -0.2 | 4.07 | 3.93 | 4.04 | 4.25 | 0.56 | -0.13 | -0.16 | -0.13 | -0.21 | -0.24 | -0.21 | -0.2 | 0.29 | -0.13 | -0.03 | -0.21 | -0.2 | -0.1 | -0.3 | -0.23 | 0.14 | -0.21 | 0.12 | -0.31 | -0.28 |
| Run1_Nl-1 | | -0.2 | 3.3 | 3.34 | 2.09 | 1.22 | 0.2 | -0.07 | -0.1 | -0.04 | -0.23 | -0.26 | -0.22 | -0.19 | 0.13 | -0.02 | -0.07 | -0.23 | -0.25 | 0 | -0.3 | -0.19 | 0.07 | 0.3 | 0.19 | -0.28 | -0.31 |
| Run1_Nl-3 | | -0.17 | 3.1 | 2.95 | 3.13 | 3.16 | 0.76 | 0.32 | 0.35 | 0.08 | -0.18 | -0.22 | -0.17 | -0.18 | 1.09 | -0.1 | -0.05 | -0.16 | -0.3 | -0.07 | -0.31 | -0.2 | -0.05 | -0.16 | -0.19 | -0.31 | -0.32 |
| Run1_Nl-4 | | -0.15 | 3.06 | 2.5 | 2.89 | 3.67 | 0.84 | 0.37 | 0.37 | 0.23 | -0.17 | -0.21 | -0.15 | -0.15 | 1.23 | -0.1 | -0.06 | -0.18 | -0.23 | -0.07 | -0.3 | -0.21 | -0.03 | -0.09 | -0.22 | -0.3 | -0.3 |
| Run1_Nl-6 | | -0.16 | 2.52 | 2.66 | 1.83 | 1.95 | 0.9 | 0.27 | 0.49 | 0.22 | -0.17 | -0.21 | -0.17 | -0.17 | 0.49 | -0.13 | -0.12 | -0.15 | -0.25 | 0.16 | -0.3 | -0.24 | 0 | -0.07 | -0.12 | -0.29 | -0.3 |
| Run2_Nl-3 | Normal | -0.19 | 2.76 | 2.84 | 2.32 | 3.32 | 0.63 | -0.01 | 0.1 | -0.01 | -0.22 | -0.25 | -0.19 | -0.17 | 0.81 | -0.17 | -0.17 | -0.18 | -0.3 | -0.11 | -0.29 | -0.25 | -0.06 | -0.16 | -0.13 | -0.28 | -0.31 |
| Run2_Nl-4 | | -0.17 | 3.54 | 3.63 | 2.71 | 4.09 | 0.69 | 0.19 | 0.18 | 0.03 | -0.19 | -0.23 | -0.17 | -0.14 | 1.13 | -0.15 | -0.13 | -0.2 | -0.2 | 0.09 | -0.3 | -0.21 | -0.03 | -0.21 | -0.12 | -0.29 | -0.31 |
| Run2_Nl-5 | | -0.18 | 2.73 | 3.82 | 1.74 | 2.05 | 0.42 | 0.25 | 0.27 | 0.13 | -0.2 | -0.24 | -0.17 | -0.16 | 0.53 | -0.2 | -0.14 | -0.24 | -0.29 | -0.03 | -0.3 | -0.24 | -0.14 | -0.17 | -0.11 | -0.29 | -0.31 |
| Run2_Nl-6 | | -0.18 | 2.01 | 2.79 | 1.44 | 2.17 | 0.83 | 0.23 | 0.27 | 0.15 | -0.21 | -0.23 | -0.19 | -0.19 | 0.42 | -0.18 | -0.21 | -0.16 | -0.22 | 0.13 | -0.29 | -0.24 | -0.07 | -0.1 | -0.17 | -0.27 | -0.29 |

**Supplemental Table 3**: Median expression of each surface marker on each cell population.

### CD56 Median

| Sample | Group | Ungated | CD34+CD38low | HSCs | MPP | CMP/GMP | Myelo/Mono-Blasts | ProMonocytes | CD14neg Monocytes | Mature Monocytes | ProMyelocytes | Myelocytes | MetaMyelocytes | MatureGrans | ProErythroblasts | Erythroblasts | LateErythroblasts | PreBcells | Mature Bcells | Plasma Cells | T cells | NK cells | pDCs | Basophils | Platelets | CD8+ T cells | CD8neg T cells |
|---|---|---|---|---|---|---|---|---|---|---|---|---|---|---|---|---|---|---|---|---|---|---|---|---|---|---|---|
| Run1_MDS17 | | 1.79 | 2.35 | 2.43 | 2.28 | 3.11 | 2.33 | 3.02 | 3.35 | 3.9 | 1.62 | 1.76 | 1.45 | 1.81 | 1.23 | 0.68 | 1.25 | 2.16 | 2.02 | 4.1 | 2.41 | 18.78 | 2.26 | 1.43 | 2.77 | 3.43 | 1.31 |
| Run2_MDS15* | AML/RAEB-T | 1.7 | 5.95 | 2.51 | 2.12 | 1.11 | 2.64 | 1.94 | 0.6 | 0.77 | 6.3 | 3.36 | 1.17 | 0.55 | 2.19 | 1.39 | 1.87 | 1.06 | 0.71 | 2.87 | 1.56 | 45.82 | 1.18 | 0.83 | 0.9 | 3.44 | 0.59 |
| Run2_MDS4 | | 1.02 | 2.8 | 2.28 | 2.08 | 1.35 | 1.89 | 1.28 | 1.28 | 1.72 | 1.14 | 0.72 | 0.72 | 0.89 | 0.42 | 0.03 | 0.57 | 1.67 | 1.88 | 2.09 | 1.11 | 13.81 | 1 | 0.5 | 1.4 | 2.2 | 0.44 |
| Run1_MDS21 | | 1.22 | 2.01 | 2.24 | 1.93 | 2.27 | 2.3 | 2.71 | 2.26 | 2.49 | 0.35 | 0.69 | 1.01 | 1.54 | 0.51 | 0.75 | 2.27 | 1.26 | 1.44 | 3.38 | 1.51 | 24.86 | 3.16 | 0.34 | 1.67 | 4.16 | 0.88 |
| Run1_MDS3 | | 1.25 | 2.37 | 2.12 | 2.29 | 2.7 | 2.7 | 2.85 | 3.87 | 4.94 | 0.53 | 0.86 | 1.17 | 1.91 | 1.02 | 1.13 | 1.11 | 1.03 | 1.17 | 3.15 | 1.49 | 21.94 | 3.04 | 0.7 | 0.9 | 4.47 | 0.84 |
| Run2_MDS13 | | 1.75 | 2.8 | 2.86 | 2.67 | 3.2 | 3.43 | 3.38 | 4.55 | 4.38 | 1.16 | 1.43 | 1.16 | 1.39 | 3.22 | 0.41 | 0.98 | 1.19 | 1.28 | 4.02 | 1.62 | 21.49 | 1.78 | 0.51 | 0.59 | 3.78 | 0.92 |
| Run2_MDS16 | | 0.88 | 1.54 | 1.52 | 1.37 | 1.75 | 1.94 | 1.79 | 1.68 | 1.81 | 0.81 | 0.62 | 0.7 | 0.84 | 1.08 | -0.14 | 1.07 | 1.23 | 0.6 | 3.72 | 0.78 | 20.29 | 2.14 | 0.32 | 2.16 | 2.37 | 0.55 |
| Run2_MDS1 | | 1.03 | 1.95 | 1.68 | 1.53 | 1.65 | 1.47 | 3.11 | 4.8 | 4.38 | 0.89 | 0.58 | 0.66 | 0.68 | 0.5 | 0.08 | 0.48 | 2.06 | 0.95 | 3.22 | 0.51 | 34.94 | 0.86 | 0.26 | 1.14 | 1.92 | 0.32 |
| Run2_MDS21 | Higher Risk | 1.09 | 1.77 | 1.91 | 1.69 | 2.09 | 2.14 | 2.27 | 2.02 | 2.08 | 0.37 | 0.6 | 0.85 | 1.14 | 1.42 | 0.28 | 0.73 | 1 | 1.14 | 2.85 | 1.32 | 17.23 | 0.84 | 0.45 | 1.15 | 3.38 | 0.81 |
| Run2_MDS26* | | 0.75 | 2.38 | 2.55 | 2.28 | 3.06 | 2.59 | 2.57 | 2.8 | 2.22 | 0.71 | 0.96 | 0.74 | 0.71 | 1.69 | 0.25 | 0.48 | 1.15 | 0.83 | 3.03 | 1.13 | 9.85 | 1.61 | 0.39 | 0.03 | 2.13 | 0.72 |
| Run2_MDS27 | | 0.97 | 2.26 | 2.31 | 2.03 | 2.51 | 2.23 | 1.86 | 1.31 | 0.96 | 0.91 | 0.83 | 0.8 | 1.14 | 1.57 | -0.25 | -0.08 | 2.12 | 0.57 | 3.88 | 1.23 | 12.7 | 0.88 | 0.21 | 2.34 | 2.14 | 0.52 |
| Run2_MDS2 | | 0.7 | 1.09 | 2.18 | 0.79 | 1.18 | 0.84 | 0.73 | 0.66 | 1.68 | 2.03 | 0.82 | 0.62 | 0.74 | 3.44 | 1.16 | 0.63 | 0.82 | 0.95 | 0.27 | 0.4 | 9.92 | 0.31 | 0.14 | 0.66 | 2.25 | 0.34 |
| Run2_MDS3 | | 1.19 | 2.05 | 2.08 | 1.99 | 2.44 | 2.54 | 2.4 | 3.01 | 3.75 | 0.74 | 1.02 | 1.1 | 1.63 | 1.52 | 0.56 | 0.62 | 0.84 | 0.91 | 2.08 | 1.29 | 20.19 | 1.09 | 0.49 | 0.61 | 3.94 | 0.71 |
| Run1_MDS23 | | 0.84 | 1.86 | 2.14 | 1.44 | 2.71 | 2.19 | 2.24 | 1.98 | 1.94 | 0.42 | 0.57 | 0.62 | 0.58 | 0.73 | 0.64 | 1.03 | 2.13 | 1.38 | 3.48 | 1.8 | 17.84 | 1.77 | 0.85 | 1.27 | 2.84 | 1.43 |
| Run1_MDS25 | | 1.18 | 3.87 | 3.73 | 2.71 | 3.25 | 2.58 | 3.04 | 2.77 | 2.69 | 1.1 | 1.05 | 0.92 | 1.03 | 0.82 | 0.91 | 2.03 | 2.37 | 1.32 | 4.22 | 1.86 | 9.36 | 2.63 | 0.78 | 1.78 | 2.8 | 1.06 |
| Run1_MDS5 | | 2.11 | 3.66 | 4.98 | 3.07 | 4.56 | 4.48 | 4.73 | 3.38 | 5.2 | 4.19 | 1.94 | 2.19 | 2.13 | 4.27 | 0.49 | 1.7 | 3.15 | 1.84 | 4.76 | 1.38 | 133.6 | 1.71 | 0.61 | 1.06 | 2.86 | 0.95 |
| Run1_MDS6 | | 2.17 | 5.58 | 4.18 | 2.35 | 4.96 | 5.75 | 5.81 | 14.55 | 6.34 | 7.12 | 6.51 | 3.89 | 1.85 | 2.34 | -0.1 | 1.91 | 2.11 | 1.64 | 4.07 | 1.63 | 21.51 | 2.67 | 0.67 | 1.54 | 3.08 | 0.82 |
| Run1_MDS8 | | 0.85 | 2.9 | 2.6 | 2.59 | 3.77 | 2.52 | 1.59 | 1.32 | 1.37 | 1.28 | 0.65 | 0.65 | 0.76 | 0.77 | 0.44 | 0.75 | 1.6 | 1.75 | 2.84 | 0.89 | 12.37 | 1.89 | 0.59 | 1.44 | 1.99 | 0.65 |
| Run1_MDS9 | | 0.92 | 2.19 | 2.14 | 2.07 | 1.94 | 1.68 | 1.41 | 1.43 | 1.49 | 0.86 | 0.87 | 0.65 | 0.72 | 0.17 | -0.02 | 1.7 | 2.07 | 0.84 | 2.39 | 1.2 | 13.69 | 1.13 | 0.71 | 1.25 | 2.87 | 0.61 |
| Run2_MDS12 | Lower Risk | 0.84 | 2.92 | 2.14 | 2.36 | 2.4 | 7.75 | 9.48 | 7.95 | 7.98 | 0.43 | 0.55 | 0.69 | 0.77 | 0.85 | 0 | 0.7 | 3.41 | 0.91 | 2.43 | 1.1 | 33.16 | 2.85 | 0.55 | 1.37 | 2.29 | 0.65 |
| Run2_MDS14 | | 0.87 | 1.69 | 1.58 | 1.8 | 1.79 | 2.01 | 1.54 | 1.93 | 1.9 | 0.8 | 0.63 | 0.79 | 0.82 | 0.46 | -0.21 | 0.98 | 1.03 | 0.9 | 2.36 | 1.19 | 38.76 | 3.1 | 0.19 | 0.75 | 2.55 | 0.74 |
| Run2_MDS19 | | 1.01 | 1.55 | 1.34 | 1.37 | 2.39 | 2.25 | 1.53 | 1.25 | 1.22 | 0.64 | 0.81 | 0.99 | 1.05 | 1.47 | -0.02 | 0.46 | 1.45 | 0.63 | 2.86 | 1.03 | 21.91 | 1.89 | 0.25 | 0.74 | 2.88 | 0.71 |
| Run2_MDS20 | | 0.91 | 2.52 | 2.88 | 1.47 | 2.3 | 1.39 | 1.32 | 1.12 | 1.33 | 0.86 | 0.67 | 0.71 | 0.91 | 1.5 | 0.23 | 1.06 | 1.14 | 1.13 | 4.1 | 0.72 | 3.87 | 1.31 | 0.25 | 1.86 | 2.13 | 0.63 |
| Run2_MDS28* | | 0.93 | 1.78 | 2.15 | 1.63 | 2.59 | 3.46 | 5.41 | 3.32 | 4.91 | 0.87 | 0.81 | 0.72 | 0.66 | 1.81 | 0.07 | 0.68 | 2.78 | 0.97 | 4.1 | 1.68 | 17.69 | 1.82 | 0.59 | 1.4 | 3.12 | 0.85 |
| Run2_MDS5 | | 1.71 | 3.02 | 0.95 | 3.23 | 3.39 | 2.65 | 2.48 | 2.59 | 4.07 | 3.7 | 1.44 | 1.76 | 1.73 | 3.18 | 0.05 | 1.36 | 1.93 | 0.5 | 3.28 | 0.85 | 132.45 | 2.02 | 0.34 | 0.74 | 2.55 | 0.53 |
| Run1_ICUS18 | | 1.16 | 1.99 | 2.04 | 1.99 | 3.02 | 2.1 | 2.01 | 1.94 | 1.75 | 0.88 | 0.91 | 0.9 | 0.96 | 0.57 | 0.59 | 1.22 | 2.17 | 0.88 | 1.93 | 1.12 | 11.81 | 2.07 | 0.56 | 1.96 | 3.2 | 0.85 |
| Run1_ICUS22 | ICUS | 0.74 | 1.81 | 2.13 | 1.67 | 1.82 | 1.82 | 1.87 | 1.71 | 1.75 | 0.36 | 0.56 | 0.64 | 0.58 | 0.14 | 0.21 | 0.94 | 1.3 | 0.88 | 3.25 | 1.28 | 6.18 | 1.68 | 0.59 | 1.15 | 2.44 | 0.93 |
| Run1_ICUS24 | | 0.94 | 2.3 | 2.3 | 2 | 2.76 | 2.67 | 2.16 | 2.04 | 2.16 | 0.55 | 0.73 | 0.73 | 0.74 | 0.29 | -0.02 | 1.08 | 1.52 | 1 | 3.11 | 1.86 | 8.32 | 1.64 | 0.75 | 1.47 | 2.72 | 1.27 |
| Run1_Nl-1 | | 0.86 | 2.18 | 2.79 | 2.05 | 1.88 | 1.61 | 1.48 | 1.4 | 1.32 | 0.57 | 0.71 | 0.78 | 0.76 | 0.34 | 0.24 | 1.17 | 1.16 | 0.87 | 3.15 | 1.19 | 12.3 | 1.51 | 0.42 | 1.59 | 2.56 | 0.71 |
| Run1_Nl-3 | | 2.01 | 3.72 | 3.01 | 3.77 | 4.84 | 4.43 | 3.55 | 3.27 | 2.7 | 1.81 | 1.46 | 1.87 | 1.98 | 2.21 | 1.01 | 2.5 | 2.58 | 1.6 | 5.47 | 2.24 | 17.01 | 4.35 | 0.85 | 3.45 | 3.43 | 1.6 |
| Run1_Nl-4 | | 2.18 | 4.54 | 4.34 | 4.31 | 5.26 | 4.9 | 3.92 | 3.45 | 3.09 | 2.22 | 1.71 | 2.03 | 2.11 | 2.23 | 1.13 | 2.5 | 2.86 | 2.27 | 6.44 | 2.4 | 13.56 | 4.03 | 0.72 | 3.56 | 3.17 | 1.85 |
| Run1_Nl-6 | | 1.7 | 2.8 | 2.72 | 2.46 | 3.68 | 4.28 | 3.69 | 3.74 | 3.24 | 2.02 | 1.39 | 1.42 | 1.6 | 1.54 | 1.03 | 1.95 | 2.46 | 1.87 | 6.11 | 2.08 | 13.31 | 3.11 | 0.77 | 1.81 | 3.38 | 1.68 |
| Run2_Nl-3 | Normal | 1.7 | 3.42 | 3.8 | 2.94 | 3.78 | 4.13 | 3.08 | 2.93 | 2.59 | 1.6 | 1.19 | 1.61 | 1.81 | 1.92 | 0.57 | 1.94 | 2.04 | 1 | 4.59 | 1.67 | 14.96 | 3.82 | 0.57 | 2.96 | 2.64 | 1.11 |
| Run2_Nl-4 | | 2.01 | 4.38 | 4.89 | 3.72 | 4.94 | 5.05 | 3.7 | 3.23 | 3.14 | 1.86 | 1.43 | 1.94 | 2.04 | 2.45 | 0.73 | 2.39 | 2.14 | 1.47 | 6.35 | 1.94 | 12.71 | 3.78 | 0.49 | 3.25 | 2.61 | 1.43 |
| Run2_Nl-5 | | 1.73 | 2.97 | 2.96 | 2.7 | 3.47 | 4.27 | 3.33 | 2.9 | 2.66 | 1.4 | 1.07 | 1.52 | 1.73 | 1.51 | 0.26 | 1.98 | 1.97 | 1.42 | 5.82 | 1.81 | 14.68 | 3.44 | 0.62 | 3.21 | 2.74 | 1.12 |
| Run2_Nl-6 | | 1.5 | 2.52 | 2.1 | 2.39 | 3.58 | 4.45 | 3.39 | 2.94 | 2.76 | 1.47 | 1.09 | 1.25 | 1.5 | 2.31 | 0.8 | 1.72 | 1.89 | 1.25 | 5.94 | 1.66 | 11.69 | 2.69 | 0.39 | 1.71 | 2.7 | 1.36 |

**Supplemental Table 3**: Median expression of each surface marker on each cell population.

**CD90 Median**

| | | Ungated | CD34+CD38low | HSCs | MPP | CMP/GMP | Myelo/Mono-Blasts | ProMonocytes | CD14neg Monocytes | Mature Monocytes | ProMyelocytes | Myelocytes | MetaMyelocytes | MatureGrans | ProErythroblasts | Erythroblasts | LateErythroblasts | PreBcells | Mature Bcells | Plasma Cells | T cells | NK cells | pDCs | Basophils | Platelets | CD8+ T cells | CD8neg T cells |
|---|---|---|---|---|---|---|---|---|---|---|---|---|---|---|---|---|---|---|---|---|---|---|---|---|---|---|---|
| Run1_MDS17 | | -0.12 | 0.55 | 4.15 | 0.32 | 0.72 | -0.06 | -0.14 | -0.02 | 0.06 | -0.3 | 0.24 | 0.05 | 0.16 | -0.24 | -0.34 | -0.22 | -0.08 | -0.41 | 0.42 | -0.44 | -0.27 | -0.12 | -0.85 | -0.11 | -0.43 | -0.45 |
| Run2_MDS15* | AML/RAEB-T | -0.2 | 2.02 | 4.58 | 0.91 | 0.69 | -0.03 | -0.25 | -0.28 | -0.24 | -0.26 | -0.29 | -0.32 | -0.34 | 0.12 | -0.21 | -0.05 | -0.21 | -0.4 | -0.12 | -0.39 | -0.11 | 0.33 | -0.27 | 0.21 | -0.39 | -0.38 |
| Run2_MDS4 | | -0.33 | 1.25 | 5.12 | 0.47 | 1.43 | -0.29 | -0.29 | -0.26 | -0.23 | -0.38 | -0.38 | -0.37 | -0.33 | -0.38 | -0.38 | -0.28 | -0.14 | -0.2 | 0.07 | -0.41 | -0.26 | -0.25 | -0.31 | -0.29 | -0.41 | -0.41 |
| Run1_MDS21 | | -0.32 | 0.63 | 4.41 | 0.31 | 0.71 | 0.46 | 0.01 | -0.23 | -0.13 | -0.4 | -0.37 | -0.36 | -0.32 | -0.34 | -0.21 | -0.23 | -0.44 | -0.41 | 0.41 | -0.46 | -0.26 | -0.68 | -0.25 | -0.25 | -0.41 | -0.47 |
| Run1_MDS3 | | -0.29 | 0.86 | 4.39 | 0.48 | 0.9 | 0.67 | 0.14 | -0.13 | -0.01 | -0.36 | -0.29 | -0.28 | -0.18 | -0.22 | -0.23 | -0.19 | -0.38 | -0.35 | 0.34 | -0.45 | -0.23 | -0.04 | -0.38 | -0.34 | -0.39 | -0.47 |
| Run2_MDS13 | | -0.25 | 0.98 | 4.21 | 0.6 | 0.84 | 0.48 | -0.22 | -0.15 | -0.19 | -0.28 | -0.22 | -0.27 | -0.24 | -0.25 | -0.33 | -0.25 | -0.36 | -0.37 | -0.21 | -0.4 | -0.21 | -0.44 | -0.4 | -0.33 | -0.38 | -0.41 |
| Run2_MDS16 | | -0.33 | 1.58 | 4.59 | 0.77 | 1.48 | -0.21 | -0.33 | -0.32 | -0.31 | -0.36 | -0.38 | -0.35 | -0.33 | 0.08 | -0.39 | -0.33 | -0.31 | -0.41 | -0.19 | -0.4 | -0.23 | -0.29 | -0.4 | -0.32 | -0.39 | -0.41 |
| Run2_MDS1 | | -0.36 | 0.14 | 4.98 | 0.01 | 0.22 | -0.28 | -0.26 | -0.31 | -0.29 | -0.41 | -0.41 | -0.38 | -0.36 | -0.23 | -0.3 | -0.4 | -0.22 | -0.31 | 0.72 | -0.42 | -0.15 | -0.4 | -0.34 | -0.36 | -0.4 | -0.42 |
| Run2_MDS21 | Higher Risk | -0.28 | 0.57 | 4.33 | 0.33 | 0.66 | 0.31 | -0.3 | -0.25 | -0.25 | -0.28 | -0.23 | -0.32 | -0.3 | -0.31 | -0.34 | -0.32 | -0.32 | -0.33 | 0.8 | -0.41 | -0.29 | -0.1 | -0.44 | -0.26 | -0.39 | -0.42 |
| Run2_MDS26* | | -0.38 | 0.45 | 4.36 | 0.23 | 0.23 | -0.24 | -0.33 | -0.25 | -0.22 | -0.39 | -0.39 | -0.38 | -0.37 | -0.27 | -0.39 | -0.37 | -0.35 | -0.41 | 0.28 | -0.42 | -0.3 | -0.28 | -0.4 | -0.17 | -0.41 | -0.43 |
| Run2_MDS27 | | -0.27 | 2.36 | 5.69 | 0.96 | 0.97 | 0.18 | -0.2 | -0.3 | -0.25 | -0.35 | -0.36 | -0.35 | -0.29 | -0.01 | -0.4 | -0.37 | -0.25 | -0.37 | 0.18 | -0.4 | -0.29 | -0.27 | -0.32 | 0.74 | -0.39 | -0.4 |
| Run2_MDS2 | | -0.34 | 0.43 | 4.38 | 0.15 | 0.41 | -0.11 | -0.3 | -0.33 | -0.2 | -0.15 | -0.4 | -0.37 | -0.35 | 0.2 | -0.41 | -0.21 | -0.24 | -0.37 | -0.35 | -0.44 | -0.36 | -0.31 | -0.48 | -0.35 | -0.35 | -0.45 |
| Run2_MDS3 | | -0.22 | 0.84 | 4.43 | 0.51 | 0.99 | 0.45 | -0.11 | -0.16 | -0.07 | -0.22 | -0.04 | -0.08 | 0.17 | -0.2 | -0.29 | -0.28 | -0.34 | -0.33 | 0.2 | -0.39 | -0.26 | -0.06 | -0.29 | -0.3 | -0.36 | -0.41 |
| Run1_MDS23 | | -0.29 | 1.69 | 4.66 | 0.67 | 0.64 | -0.28 | -0.21 | -0.2 | -0.2 | -0.04 | -0.17 | -0.29 | -0.32 | -0.31 | -0.31 | -0.21 | -0.31 | -0.37 | 0.48 | -0.46 | -0.26 | -0.36 | -0.44 | -0.17 | -0.44 | -0.46 |
| Run1_MDS25 | | -0.39 | 3.39 | 5.96 | -0.29 | 0.67 | -0.28 | -0.38 | -0.36 | -0.35 | -0.39 | -0.46 | -0.42 | -0.39 | -0.4 | -0.36 | -0.02 | -0.44 | -0.37 | 0 | -0.44 | -0.3 | -0.32 | -0.3 | -0.34 | -0.42 | -0.47 |
| Run1_MDS5 | | -0.29 | 1.52 | 4.5 | 0.65 | 0.96 | -0.11 | -0.22 | -0.13 | 0.07 | -0.34 | -0.38 | -0.33 | -0.3 | -0.18 | -0.32 | -0.26 | -0.23 | -0.4 | -0.24 | -0.43 | 0.43 | -0.1 | -0.38 | -0.38 | -0.41 | -0.43 |
| Run1_MDS6 | | -0.27 | 7.33 | 7.49 | 1.08 | 4.77 | 0.62 | -0.03 | 0.16 | 0.11 | -0.31 | -0.28 | -0.25 | -0.25 | 0.03 | -0.4 | 0.04 | -0.31 | -0.43 | 1.6 | -0.44 | -0.26 | -0.16 | -0.34 | 1.47 | -0.42 | -0.46 |
| Run1_MDS8 | | -0.39 | 2.89 | 6.39 | 1.18 | 1.16 | -0.09 | -0.21 | -0.28 | -0.31 | -0.44 | -0.42 | -0.42 | -0.38 | -0.38 | -0.31 | -0.33 | -0.41 | -0.47 | -0.53 | -0.47 | -0.28 | 0.24 | -0.43 | -0.35 | -0.46 | -0.48 |
| Run1_MDS9 | | -0.39 | 2.17 | 5.8 | 0.67 | 1.57 | 0.21 | -0.38 | -0.41 | -0.36 | -0.38 | -0.39 | -0.4 | -0.38 | -0.37 | -0.4 | -0.26 | -0.43 | -0.43 | -0.12 | -0.47 | -0.33 | -0.4 | -0.4 | -0.41 | -0.44 | -0.49 |
| Run2_MDS12 | Lower Risk | -0.3 | 0.65 | 4.65 | 0.44 | 0.53 | -0.15 | -0.14 | -0.13 | -0.13 | -0.3 | -0.32 | -0.3 | -0.29 | -0.33 | -0.4 | -0.33 | -0.3 | -0.37 | 0.5 | -0.39 | -0.24 | -0.2 | -0.4 | -0.3 | -0.37 | -0.41 |
| Run2_MDS14 | | -0.26 | 5.08 | 53.7 | 1.35 | 0.84 | -0.27 | -0.35 | -0.18 | -0.16 | -0.29 | -0.31 | -0.23 | -0.24 | -0.31 | -0.36 | -0.19 | -0.34 | -0.4 | 1.19 | -0.39 | -0.17 | -0.12 | -0.43 | -0.42 | -0.38 | -0.39 |
| Run2_MDS19 | | -0.34 | 1.59 | 4.36 | 0.86 | 0.97 | -0.2 | -0.32 | -0.32 | -0.32 | -0.33 | -0.36 | -0.34 | -0.33 | -0.32 | -0.39 | -0.37 | -0.36 | -0.4 | 0.19 | -0.4 | -0.26 | -0.28 | -0.34 | -0.39 | -0.39 | -0.41 |
| Run2_MDS20 | | -0.33 | 2.1 | 5.67 | 0.44 | 1.25 | -0.26 | -0.32 | -0.26 | -0.26 | -0.34 | -0.34 | -0.31 | -0.29 | -0.23 | -0.31 | -0.3 | -0.34 | -0.37 | 0 | -0.41 | -0.28 | -0.39 | -0.4 | -0.33 | -0.39 | -0.41 |
| Run2_MDS28* | | -0.24 | 1.56 | 5.66 | 0.92 | 1.07 | -0.05 | -0.2 | -0.17 | -0.12 | -0.25 | -0.29 | -0.22 | -0.24 | -0.25 | -0.38 | -0.23 | -0.29 | -0.35 | -0.05 | -0.4 | -0.26 | -0.18 | -0.38 | -0.11 | -0.39 | -0.42 |
| Run2_MDS5 | | -0.28 | 1.78 | 4.43 | 0.47 | 0.36 | -0.12 | -0.23 | -0.15 | 0 | -0.28 | -0.35 | -0.31 | -0.28 | -0.12 | -0.38 | -0.28 | -0.12 | -0.39 | -0.23 | -0.4 | 0.23 | -0.33 | -0.39 | -0.36 | -0.42 | -0.4 |
| Run1_ICUS18 | | -0.34 | 1.44 | 5.68 | 0.73 | 1.43 | -0.34 | -0.31 | -0.26 | -0.27 | -0.33 | -0.34 | -0.35 | -0.32 | -0.38 | -0.4 | -0.27 | -0.35 | -0.49 | -0.22 | -0.46 | -0.33 | -0.38 | -0.46 | -0.33 | -0.42 | -0.47 |
| Run1_ICUS22 | ICUS | -0.37 | 1.5 | 5.28 | 0.6 | 0.57 | -0.28 | -0.34 | -0.33 | -0.3 | -0.34 | -0.38 | -0.38 | -0.37 | -0.34 | -0.34 | -0.3 | -0.36 | -0.44 | -0.18 | -0.45 | -0.39 | -0.34 | -0.41 | -0.44 | -0.43 | -0.46 |
| Run1_ICUS24 | | -0.36 | 2.28 | 5.48 | 1.03 | 0.6 | -0.2 | -0.29 | -0.32 | -0.3 | -0.37 | -0.36 | -0.35 | -0.35 | -0.32 | -0.35 | -0.31 | -0.39 | -0.43 | 1.1 | -0.46 | -0.35 | -0.26 | -0.38 | -0.35 | -0.45 | -0.46 |
| Run1_Nl-1 | | -0.38 | 2.04 | 4.02 | 1.11 | 0.33 | -0.26 | -0.28 | -0.27 | -0.26 | -0.39 | -0.39 | -0.37 | -0.37 | -0.28 | -0.25 | -0.3 | -0.38 | -0.45 | 1.92 | -0.47 | -0.34 | -0.35 | -0.43 | -0.33 | -0.45 | -0.47 |
| Run1_Nl-3 | | -0.27 | 2.18 | 4.97 | 0.98 | 1.23 | -0.02 | 0.22 | 0.51 | 0.1 | -0.34 | -0.32 | -0.24 | -0.23 | -0.28 | -0.38 | -0.14 | -0.36 | -0.47 | 0.19 | -0.45 | -0.25 | -0.31 | -0.43 | -0.35 | -0.44 | -0.46 |
| Run1_Nl-4 | | -0.24 | 2.1 | 4.92 | 1.24 | 1.16 | 0.03 | 0.48 | 0.73 | 0.27 | -0.3 | -0.31 | -0.23 | -0.2 | -0.24 | -0.35 | -0.14 | -0.32 | -0.44 | 0.54 | -0.43 | -0.29 | -0.25 | -0.39 | -0.33 | -0.43 | -0.44 |
| Run1_Nl-6 | | -0.25 | 1.96 | 4.89 | 0.79 | 0.74 | 0.1 | 0.37 | 0.53 | 0.24 | -0.31 | -0.28 | -0.23 | -0.22 | -0.27 | -0.37 | -0.17 | -0.34 | -0.45 | 0.96 | -0.44 | -0.32 | -0.28 | -0.39 | -0.27 | -0.43 | -0.45 |
| Run2_Nl-3 | Normal | -0.24 | 1.9 | 4.68 | 0.95 | 0.84 | -0.06 | 0.01 | 0.27 | -0.03 | -0.31 | -0.28 | -0.19 | -0.16 | -0.31 | -0.35 | -0.16 | -0.3 | -0.4 | -0.27 | -0.41 | -0.29 | -0.31 | -0.41 | -0.32 | -0.4 | -0.41 |
| Run2_Nl-4 | | -0.19 | 1.96 | 4.98 | 0.83 | 0.99 | -0.04 | 0.35 | 0.49 | 0.2 | -0.28 | -0.27 | -0.15 | -0.09 | -0.23 | -0.33 | -0.09 | -0.29 | -0.4 | 0.37 | -0.39 | -0.27 | -0.23 | -0.37 | -0.33 | -0.39 | -0.39 |
| Run2_Nl-5 | | -0.2 | 1.85 | 5.07 | 0.74 | 0.8 | -0.08 | 0.19 | 0.48 | 0.19 | -0.32 | -0.28 | -0.15 | -0.06 | -0.26 | -0.35 | -0.12 | -0.3 | -0.39 | 0.44 | -0.41 | -0.27 | -0.29 | -0.42 | -0.3 | -0.4 | -0.42 |
| Run2_Nl-6 | | -0.25 | 1.65 | 5.05 | 0.88 | 0.8 | -0.04 | 0.17 | 0.28 | 0.17 | -0.3 | -0.29 | -0.24 | -0.21 | -0.25 | -0.34 | -0.27 | -0.34 | -0.4 | 0.73 | -0.4 | -0.3 | -0.32 | -0.36 | -0.27 | -0.38 | -0.41 |

**Supplemental Table 3**: Median expression of each surface marker on each cell population.

### CD33 Median

| | | Ungated | CD34+CD38low | HSCs | MPP | CMP/GMP | Myelo/Mono-Blasts | ProMonocytes | CD14neg Monocytes | Mature Monocytes | ProMyelocytes | Myelocytes | MetaMyelocytes | MatureGrans | ProErythroblasts | Erythroblasts | LateErythroblasts | PreBcells | Mature Bcells | Plasma Cells | T cells | NK cells | pDCs | Basophils | Platelets | CD8+ T cells | CD8neg T cells |
|---|---|---|---|---|---|---|---|---|---|---|---|---|---|---|---|---|---|---|---|---|---|---|---|---|---|---|---|
| Run1_MDS17 | | 0.95 | -0.08 | -0.06 | -0.16 | 0.32 | -0.1 | 13.79 | 32.58 | 40.96 | 2.92 | 4.54 | 6.59 | 7.59 | -0.39 | -0.44 | -0.23 | 0.66 | -0.31 | -0.44 | -0.44 | -0.37 | 2.22 | 12.15 | -0.26 | -0.44 | -0.43 |
| Run2_MDS15* | AML/RAEB-T | 12.74 | 9.19 | 1.67 | 1.45 | 1.61 | 10.57 | 17.17 | 38.24 | 51.85 | 6.31 | 7.5 | 10.82 | 10.06 | 6.68 | 6.51 | 15.05 | 24.08 | -0.21 | 0.03 | -0.39 | -0.3 | 12.99 | 13.78 | 1.32 | -0.39 | -0.39 |
| Run2_MDS4 | | 3.96 | 7.75 | 1.48 | 1.53 | 0.06 | 22.5 | 55.32 | 70.77 | 69.28 | 5.79 | 5.56 | 6.27 | 7.36 | -0.23 | -0.35 | -0.21 | 0.47 | 0.46 | -0.29 | -0.39 | -0.25 | 2.05 | 7.88 | 0.04 | -0.39 | -0.4 |
| Run1_MDS21 | | 0.22 | 0.66 | 0.56 | 0.35 | 0.81 | 1.24 | 8.38 | 51.54 | 48.47 | 0.7 | 2.38 | 5.51 | 7.19 | -0.35 | -0.23 | 0.48 | -0.4 | -0.39 | -0.26 | -0.46 | -0.31 | 3.57 | 15.04 | -0.35 | -0.44 | -0.46 |
| Run1_MDS3 | | 0.47 | 1.83 | 0.64 | 0.73 | 1.17 | 2.36 | 6.1 | 40.93 | 49.38 | 0.9 | 2.22 | 5.44 | 6.89 | -0.29 | -0.25 | 0.28 | -0.36 | -0.29 | 0.4 | -0.44 | -0.27 | 4.07 | 19.02 | -0.33 | -0.43 | -0.45 |
| Run2_MDS13 | | 0.98 | 0.69 | 0.52 | 0.36 | 0.79 | 1.6 | 25.61 | 45.99 | 50.31 | 2.35 | 3.98 | 6.34 | 6.53 | -0.19 | -0.29 | -0.21 | -0.32 | -0.32 | -0.25 | -0.41 | -0.38 | 8.99 | 11.97 | -0.21 | -0.39 | -0.41 |
| Run2_MDS16 | | 1.98 | 2.72 | 1 | 1.07 | 1.15 | 11.78 | 29.49 | 53.1 | 55.89 | 2.01 | 3.23 | 3.65 | 4.46 | 1.22 | -0.37 | 0.9 | -0.18 | -0.34 | -0.3 | -0.41 | -0.35 | 2.96 | 7.17 | -0.29 | -0.41 | -0.41 |
| Run2_MDS1 | | -0.15 | -0.01 | -0.07 | -0.2 | -0.1 | 0.28 | 2.95 | 7.87 | 11.67 | -0.2 | -0.28 | -0.06 | 0.17 | -0.39 | -0.4 | -0.32 | -0.07 | -0.28 | -0.24 | -0.41 | -0.29 | 0.86 | 0.94 | -0.33 | -0.4 | -0.41 |
| Run2_MDS21 | Higher Risk | 0.14 | 0.42 | 0.38 | 0.19 | 0.73 | 1.04 | 17.36 | 49.95 | 51.68 | 0.97 | 2.35 | 5.47 | 6.26 | -0.32 | -0.35 | -0.32 | -0.3 | -0.34 | -0.21 | -0.41 | -0.4 | 6.64 | 14.1 | -0.3 | -0.41 | -0.41 |
| Run2_MDS26* | | 0.37 | 0.7 | 0.57 | 0.42 | 0.96 | 0.55 | 4.31 | 25.26 | 33.01 | 0 | 0.36 | 1.19 | 1.55 | -0.12 | -0.4 | -0.38 | -0.09 | -0.37 | -0.63 | -0.4 | -0.39 | 1.85 | 4.19 | 0.4 | -0.4 | -0.41 |
| Run2_MDS27 | | -0.16 | 0.51 | 0.31 | 0.3 | 0.97 | 1.14 | 4.65 | 4.09 | 5.12 | 0.14 | 0.54 | 0.49 | 0.51 | 0.44 | -0.41 | -0.35 | -0.28 | -0.27 | -0.19 | -0.4 | -0.35 | 1.98 | 2.96 | -0.21 | -0.4 | -0.39 |
| Run2_MDS2 | | 1.22 | 0.25 | 0.56 | 0 | 0.42 | 0.7 | 26.73 | 27.26 | 37.46 | 0.84 | 0.88 | 2.48 | 3.43 | 1.41 | 1.45 | -0.05 | -0.06 | 0.31 | -0.15 | -0.41 | -0.37 | 3.79 | 4.6 | -0.36 | -0.4 | -0.41 |
| Run2_MDS3 | | 0.72 | 0.75 | 0.54 | 0.5 | 1.08 | 2.36 | 16.33 | 34.73 | 37.35 | 2.07 | 3.76 | 6.36 | 7.04 | -0.21 | -0.31 | -0.3 | -0.33 | -0.3 | -0.23 | -0.4 | -0.27 | 4.3 | 15.7 | -0.24 | -0.4 | -0.4 |
| Run1_MDS23 | | 0.48 | -0.16 | -0.23 | -0.28 | 0.62 | 2.55 | 14.96 | 24.39 | 31.25 | -0.18 | 0.12 | 0.58 | 0.75 | -0.29 | -0.24 | 1.02 | 0.68 | -0.31 | -0.17 | -0.47 | -0.39 | 1.02 | 4 | -0.25 | -0.46 | -0.47 |
| Run1_MDS25 | | 2.82 | 2.75 | -0.07 | 2.16 | 0.52 | 9.47 | 20.25 | 24.09 | 23.2 | 0.96 | 1.75 | 2.4 | 3.09 | -0.21 | 0.27 | 2.72 | 1.2 | -0.29 | -0.14 | -0.44 | -0.35 | 3.43 | 8.66 | -0.44 | -0.46 | -0.43 |
| Run1_MDS5 | | 2.23 | 3.98 | 1.6 | 1.43 | 2.28 | 5.29 | 30.42 | 25.1 | 41.38 | 2.01 | 2.02 | 2.62 | 3.16 | 0.73 | -0.22 | 0.67 | 1.21 | -0.35 | -0.15 | -0.42 | -0.15 | 10.22 | 16.04 | -0.29 | -0.41 | -0.43 |
| Run1_MDS6 | | 5.3 | 7.1 | 1.79 | 1.33 | 2.68 | 11.54 | 31.58 | 47.76 | 52.26 | 3.36 | 5.52 | 7.23 | 6.28 | 2.68 | -0.39 | 3.47 | -0.22 | -0.32 | -0.05 | -0.45 | -0.34 | 8.06 | 9.22 | -0.11 | -0.44 | -0.46 |
| Run1_MDS8 | | 0.36 | 0.46 | 0.04 | 0.56 | 0.02 | 1.5 | 5.23 | 8.02 | 8.61 | 0.02 | 0.54 | 0.49 | 0.66 | -0.38 | -0.34 | -0.15 | -0.25 | -0.4 | -0.15 | -0.48 | -0.35 | 4.22 | 2.88 | -0.36 | -0.46 | -0.49 |
| Run1_MDS9 | | -0.15 | 0.45 | 0.37 | 0.11 | 0.51 | 0.81 | 2.5 | 5.15 | 5.51 | -0.14 | 0.11 | -0.03 | 0.08 | -0.34 | -0.4 | -0.02 | -0.29 | -0.31 | -0.42 | -0.47 | -0.4 | 1.74 | 2.47 | -0.35 | -0.46 | -0.47 |
| Run2_MDS12 | Lower Risk | 1.95 | 2.16 | 0.65 | 0.65 | 1.01 | 14.71 | 22.49 | 28.47 | 34.06 | 0.29 | 1.55 | 2.22 | 2.11 | -0.33 | -0.4 | 0 | 11.03 | -0.25 | -0.38 | -0.41 | -0.37 | 2.33 | 6.61 | -0.01 | -0.41 | -0.41 |
| Run2_MDS14 | | 1.97 | 1.28 | 0.42 | 1.28 | 0.78 | 6.72 | 14.57 | 20.87 | 27.09 | 0.89 | 1.68 | 2.42 | 2.26 | -0.27 | -0.36 | 2.17 | -0.2 | -0.32 | -0.28 | -0.4 | -0.27 | 2.52 | 4.04 | -0.41 | -0.41 | -0.4 |
| Run2_MDS19 | | 1.79 | 0.79 | 0.18 | 0.35 | 1.39 | 14.59 | 28.38 | 32.84 | 37.73 | 0.97 | 1.93 | 2.27 | 2.24 | -0.33 | -0.36 | -0.22 | 0.25 | -0.34 | -0.37 | -0.41 | -0.33 | 2.43 | 5.62 | -0.39 | -0.39 | -0.41 |
| Run2_MDS20 | | 1.33 | 0.65 | 0.65 | 0.41 | 1.63 | 10.04 | 21.19 | 33.91 | 33.52 | 0.65 | 1.69 | 2.32 | 2.7 | 0.27 | -0.25 | 1.81 | -0.15 | -0.18 | 0.09 | -0.41 | -0.28 | 4.04 | 4.31 | -0.32 | -0.4 | -0.41 |
| Run2_MDS28* | | 2.7 | 0.8 | 0.24 | 0.56 | 0.9 | 11.27 | 21 | 28.03 | 40.2 | 0.61 | 1.34 | 2.57 | 3.32 | 0.08 | -0.32 | 2.12 | 6.28 | -0.31 | -0.2 | -0.4 | -0.36 | 15.82 | 8.16 | -0.24 | -0.41 | -0.4 |
| Run2_MDS5 | | 1.82 | 5.4 | 0.26 | 0.95 | 1.69 | 4.73 | 22.83 | 22.27 | 42.33 | 1.6 | 1.43 | 1.99 | 2.54 | 0.12 | -0.19 | 1.05 | 0.33 | -0.35 | -0.22 | -0.38 | -0.17 | 11.61 | 13.98 | -0.26 | -0.37 | -0.38 |
| Run1_ICUS18 | | 4.43 | 1.64 | 1.38 | 0.81 | 1.29 | 15.18 | 36.2 | 68.97 | 67.55 | 2.28 | 4.52 | 5.46 | 5.43 | -0.21 | -0.24 | 6.42 | -0.15 | -0.37 | -0.06 | -0.46 | -0.39 | 3.61 | 7.24 | -0.37 | -0.46 | -0.46 |
| Run1_ICUS22 | ICUS | 2.39 | 1.09 | 0.75 | 0.59 | 1.18 | 8.06 | 17.19 | 29.93 | 33.1 | 0.48 | 1.51 | 2.37 | 2.75 | -0.31 | -0.25 | 2.85 | -0.1 | -0.31 | -0.34 | -0.45 | -0.37 | 5.12 | 5.67 | -0.41 | -0.45 | -0.45 |
| Run1_ICUS24 | | 1.29 | 0.06 | -0.08 | -0.01 | 0.53 | 4.3 | 15.01 | 19.68 | 24.77 | 0.33 | 1.14 | 1.93 | 2.25 | -0.38 | -0.42 | 0.19 | -0.26 | -0.27 | -0.34 | -0.46 | -0.44 | 2.32 | 4.23 | -0.35 | -0.46 | -0.46 |
| Run1_Nl-1 | | 5.66 | 1.68 | 0.95 | 1.12 | 0.76 | 12.87 | 40.24 | 66.9 | 90.29 | 2.09 | 4.3 | 6.52 | 7.56 | -0.26 | -0.26 | 6.39 | -0.26 | -0.34 | -0.09 | -0.45 | -0.39 | 3.41 | 9.19 | -0.34 | -0.45 | -0.46 |
| Run1_Nl-3 | | 1.33 | 1.46 | 1.19 | 0.72 | 1.4 | 9.78 | 19.63 | 26.26 | 32.35 | 0.91 | 1.76 | 1.78 | 1.57 | -0.15 | -0.27 | 2.03 | -0.13 | -0.37 | -0.15 | -0.46 | -0.34 | 2.43 | 4.58 | -0.38 | -0.46 | -0.47 |
| Run1_Nl-4 | | 1.44 | 1.33 | 0.5 | 0.73 | 1.54 | 8.82 | 21.22 | 27.35 | 32.85 | 0.89 | 1.76 | 1.79 | 1.62 | -0.04 | -0.14 | 2.15 | -0.16 | -0.32 | 0.47 | -0.46 | -0.36 | 2.81 | 5.52 | -0.31 | -0.46 | -0.45 |
| Run1_Nl-6 | | 2.43 | 1.38 | 0.79 | 0.84 | 1.04 | 11.38 | 37.04 | 51.06 | 59.73 | 1.28 | 2.8 | 3.35 | 3.01 | -0.2 | -0.31 | 3 | -0.22 | -0.18 | -0.04 | -0.46 | -0.38 | 4.09 | 10.91 | -0.46 | -0.47 | -0.45 |
| Run2_Nl-3 | Normal | 1.15 | 1.09 | 0.72 | 0.67 | 1.36 | 8.63 | 18.39 | 23.78 | 29.25 | 0.93 | 1.41 | 1.47 | 1.41 | -0.25 | -0.3 | 1.41 | -0.14 | -0.35 | -0.3 | -0.41 | -0.35 | 1.81 | 3.44 | -0.32 | -0.41 | -0.41 |
| Run2_Nl-4 | | 1.36 | 0.98 | 0.24 | 0.65 | 1.1 | 9.38 | 20.04 | 24.9 | 28.35 | 0.93 | 1.52 | 1.64 | 1.54 | -0.19 | -0.21 | 1.75 | -0.13 | -0.29 | 0.18 | -0.4 | -0.35 | 2.21 | 3.91 | -0.32 | -0.41 | -0.4 |
| Run2_Nl-5 | | 1.23 | 1.02 | 0.7 | 0.7 | 0.83 | 9.78 | 20.99 | 30.26 | 33.04 | 0.83 | 1.82 | 2 | 1.84 | -0.1 | -0.31 | 1.76 | -0.2 | -0.34 | -0.07 | -0.41 | -0.35 | 1.87 | 5.6 | -0.36 | -0.42 | -0.41 |
| Run2_Nl-6 | | 2.4 | 0.94 | 0.61 | 0.46 | 0.96 | 11.49 | 35.76 | 47.23 | 52.68 | 1.27 | 2.51 | 3.08 | 2.78 | -0.18 | -0.33 | 1.42 | -0.21 | -0.21 | -0.15 | -0.39 | -0.34 | 3.08 | 7.8 | -0.34 | -0.41 | -0.39 |

**Supplemental Table 3**: Median expression of each surface marker on each cell population.

### CD64 Medians

| Sample | Group | Ungated | CD34+CD38low | HSCs | MPP | CMP/GMP | Myelo/Mono-Blasts | ProMonocytes | CD14neg Monocytes | Mature Monocytes | ProMyelocytes | Myelocytes | MetaMyelocytes | MatureGrans | ProErythroblasts | Erythroblasts | LateErythroblasts | PreBcells | Mature Bcells | Plasma Cells | T cells | NK cells | pDCs | Basophils | Platelets | CD8+ T cells | CD8neg T cells |
|---|---|---|---|---|---|---|---|---|---|---|---|---|---|---|---|---|---|---|---|---|---|---|---|---|---|---|---|
| Run1_MDS17 | | 1.53 | 0.37 | 0.46 | 0.28 | 0.59 | 0.34 | 26.2 | 60.94 | 82.28 | 1.79 | 9.88 | 12.86 | 13.37 | -0.16 | -0.29 | -0.07 | 1.37 | 2.86 | 0.84 | -0.24 | -0.22 | 2.65 | 1.75 | -0.06 | -0.26 | -0.19 |
| Run2_MDS15* | AML/RAEB-T | 3.6 | 2.16 | 1.32 | 0.93 | 0.73 | 2.8 | 11.82 | 6.91 | 18.89 | 3.66 | 3.34 | 2.45 | 1.59 | 2.04 | 1.93 | 3.65 | 5.72 | 3.02 | 2.1 | -0.19 | -0.09 | 3.33 | 0.28 | 0.54 | -0.17 | -0.2 |
| Run2_MDS4 | | 10.55 | 3.11 | 2.41 | 1.79 | 1.97 | 13.5 | 99.7 | 152.64 | 152.41 | 29.96 | 26.6 | 30.03 | 55.57 | 0 | -0.17 | 0.25 | 2.85 | 3.07 | -0.14 | -0.13 | 0.02 | 1.92 | -0.19 | 0.43 | -0.12 | -0.14 |
| Run1_MDS21 | | 0.7 | 1.63 | 1.63 | 1.49 | 1.76 | 1.58 | 8.11 | 139.72 | 97.78 | 1.36 | 7.1 | 4.02 | 4.11 | -0.21 | -0.11 | 1.02 | 1.21 | 2.97 | 0.23 | -0.3 | 0.11 | 2.47 | -0.05 | -0.21 | -0.28 | -0.31 |
| Run1_MDS3 | | 0.8 | 1.12 | 0.91 | 1.14 | 1.38 | 1.23 | 5.97 | 160.45 | 165.91 | 1.59 | 6.53 | 3.95 | 5.05 | -0.16 | -0.02 | 1.14 | 1.47 | 3.24 | 0.67 | -0.28 | -0.04 | 3.13 | 0 | -0.22 | -0.28 | -0.27 |
| Run2_MDS13 | | 1.12 | 2.26 | 2.39 | 2.07 | 2.36 | 2.32 | 61.09 | 170.55 | 167.68 | 1.74 | 9.44 | 3.85 | 3.28 | 0.23 | -0.21 | -0.11 | 0.95 | 2.39 | 1.35 | -0.27 | -0.24 | 3.85 | 0.22 | -0.13 | -0.22 | -0.29 |
| Run2_MDS16 | | 1.1 | 1.61 | 1.69 | 1.3 | 1.51 | 5.34 | 45.53 | 124.06 | 97.85 | 39 | 35.25 | 6.29 | 1.81 | 1.08 | -0.28 | 0.69 | -0.07 | 2.2 | 0.29 | -0.28 | -0.23 | 2.81 | -0.26 | -0.15 | -0.3 | -0.28 |
| Run2_MDS1 | | 1.05 | 2.74 | 2.25 | 2.03 | 2.5 | 3.3 | 48.6 | 176.54 | 168.85 | 4.27 | 2.29 | 2.35 | 2.19 | 0.37 | -0.12 | -0.11 | 0.89 | 1.95 | 0.35 | -0.25 | -0.21 | 1.45 | -0.04 | 0.06 | -0.22 | -0.26 |
| Run2_MDS21 | Higher Risk | 0.67 | 1.93 | 2.06 | 1.78 | 2.19 | 1.98 | 44.7 | 127.47 | 98.74 | 1.42 | 8.28 | 4.63 | 3.7 | -0.11 | -0.27 | -0.24 | 1.02 | 2.52 | 0.26 | -0.28 | -0.25 | 3.23 | -0.18 | -0.19 | -0.25 | -0.3 |
| Run2_MDS26* | | 0.72 | 1.52 | 1.62 | 1.34 | 1.83 | 0.83 | 2.95 | 25.92 | 58.39 | 2.48 | 2.83 | 1.36 | 1.26 | -0.07 | -0.3 | -0.31 | 0.37 | 1.91 | -0.51 | -0.29 | -0.25 | 2.26 | -0.31 | 2.06 | -0.29 | -0.29 |
| Run2_MDS27 | | 1.16 | 2.67 | 2.87 | 2.28 | 3.01 | 3.17 | 69.32 | 120.54 | 99.76 | 29.17 | 44.93 | 16.03 | 6.04 | 1.89 | -0.25 | -0.21 | 0.15 | 2.91 | 0.39 | -0.21 | -0.21 | 3.45 | -0.28 | -0.26 | -0.22 | -0.19 |
| Run2_MDS2 | | 10.02 | 2.66 | 4.74 | 2.09 | 2.76 | 2.75 | 294.6 | 352.81 | 415.89 | 9.22 | 12.21 | 27.16 | 30.06 | 3.67 | 6.46 | 1.08 | 1.62 | 4.82 | 2.76 | -0.21 | -0.24 | 6.57 | -0.03 | -0.09 | -0.16 | -0.22 |
| Run2_MDS3 | | 0.95 | 1.97 | 1.78 | 1.82 | 2.24 | 2.19 | 44.69 | 131.15 | 127.95 | 2 | 9.09 | 5.54 | 4.5 | 0.21 | -0.17 | -0.19 | 1.14 | 2.58 | 0.38 | -0.26 | -0.21 | 7.37 | 0.01 | -0.13 | -0.25 | -0.27 |
| Run1_MDS23 | | 13.41 | 2.63 | 1.88 | 2.78 | 3.62 | 8.59 | 83.09 | 154.04 | 183.27 | 17.97 | 20 | 14.22 | 13.82 | 0.5 | 0.74 | 14.56 | 5.33 | 3.14 | 0.57 | -0.25 | -0.16 | 3.65 | -0.22 | 0.96 | -0.15 | -0.27 |
| Run1_MDS25 | | 2.93 | 2.29 | 1.2 | 1.57 | 2.76 | 12.09 | 76.53 | 94.62 | 87.47 | 14.67 | 15.66 | 2.35 | 2.96 | 0.92 | 1.87 | 3.93 | 4.2 | 3.72 | 1.77 | -0.11 | -0.08 | 3.73 | -0.18 | -0.18 | -0.09 | -0.14 |
| Run1_MDS5 | | 284.42 | 6.43 | 0.95 | 6.34 | 2.33 | 3.98 | 354.97 | 365.99 | 540.94 | 254.2 | 259.34 | 333.53 | 350.11 | 14.78 | 0.84 | 5.85 | 6.89 | 2.57 | 1.4 | -0.1 | 0.54 | 4.84 | 0.79 | 0.83 | 0.3 | -0.17 |
| Run1_MDS6 | | 10.26 | 3.18 | 3.03 | 2.79 | 3.11 | 6.89 | 62.27 | 127.31 | 129.62 | 38.82 | 43.26 | 27.23 | 10.27 | 2.31 | -0.09 | 5.47 | 1.41 | 2.98 | 1.31 | -0.23 | -0.16 | 3.16 | -0.07 | 0.13 | -0.19 | -0.26 |
| Run1_MDS8 | | 3.58 | 2.1 | 1.45 | 1.71 | 1.89 | 4.26 | 33.45 | 119.89 | 118.41 | 13.98 | 19.96 | 5.35 | 4.43 | -0.1 | 0.12 | 0.82 | 0.22 | 2.42 | 0.99 | -0.35 | -0.16 | 2.79 | -0.3 | -0.08 | -0.31 | -0.36 |
| Run1_MDS9 | | 4.46 | 2.75 | 3.01 | 2.33 | 2.94 | 3.28 | 11.01 | 59.82 | 72.19 | 20.24 | 23.76 | 9.75 | 7.01 | -0.02 | -0.06 | 2.86 | -0.06 | 2.86 | 0.82 | -0.32 | -0.28 | 3.73 | -0.25 | -0.02 | -0.3 | -0.34 |
| Run2_MDS12 | Lower Risk | 16.66 | 4.26 | 3.16 | 3.59 | 3 | 34.89 | 137.78 | 169.06 | 175.79 | 17.38 | 38.43 | 22.84 | 7.44 | -0.15 | -0.3 | 1.24 | 34.86 | 2.99 | -0.04 | -0.27 | -0.12 | 2.3 | -0.16 | 3.82 | -0.24 | -0.29 |
| Run2_MDS14 | | 1.96 | 1.6 | 0.4 | 3.18 | 0.76 | 6.41 | 23 | 150.6 | 140.63 | 26.21 | 29.89 | 18.68 | 1.72 | -0.17 | -0.3 | 2.16 | 1.96 | 2.57 | -0.1 | -0.23 | -0.2 | 1.8 | -0.22 | -0.32 | -0.25 | -0.22 |
| Run2_MDS19 | | 1.74 | 1.76 | 2.21 | 1.15 | 2.65 | 7.24 | 32.41 | 44.69 | 38.35 | 11.2 | 11.08 | 3.32 | 0.9 | -0.01 | -0.23 | -0.11 | 0.86 | 2.44 | 0.17 | -0.25 | -0.15 | 2.62 | -0.29 | -0.24 | -0.26 | -0.25 |
| Run2_MDS20 | | 3.13 | 1.92 | 1.78 | 1.49 | 2.82 | 10.43 | 55.12 | 201.84 | 194.69 | 23.24 | 33.36 | 14.13 | 5.23 | 2.67 | 1.16 | 3.92 | 0.35 | 3.2 | 1.82 | -0.26 | -0.04 | 3.9 | -0.23 | -0.17 | -0.2 | -0.27 |
| Run2_MDS28* | | 14.68 | 2.47 | 2.24 | 1.73 | 2.31 | 22.63 | 81.45 | 117.31 | 163.87 | 26.61 | 31.45 | 21.94 | 8.81 | 0.7 | -0.13 | 9.24 | 11.3 | 2.65 | -0.01 | -0.26 | -0.21 | 11.76 | -0.23 | 0.62 | -0.24 | -0.28 |
| Run2_MDS5 | | 298.94 | 5.49 | 0.59 | 3.01 | 2.26 | 4.67 | 307.22 | 382.89 | 562.18 | 290.28 | 257.41 | 341.85 | 354.91 | 3.61 | 0.98 | 176.3 | 3.04 | 1.82 | 1.38 | -0.15 | 0.36 | 3.15 | 0.55 | 0.79 | 0.01 | -0.21 |
| Run1_ICUS18 | | 2.83 | 2.16 | 2.84 | 1.5 | 1.86 | 5.47 | 18.67 | 63 | 57.85 | 25.46 | 34.11 | 12.72 | 3.24 | 0.35 | 0.11 | 4.25 | 0.72 | 2.98 | 0.62 | -0.26 | -0.23 | 3.16 | -0.26 | -0.24 | -0.27 | -0.26 |
| Run1_ICUS22 | ICUS | 6.57 | 2.22 | 2.24 | 2.07 | 2.62 | 8.84 | 45.67 | 139.7 | 134.27 | 30.12 | 37.27 | 19.61 | 5.48 | 0.73 | 0.91 | 7.32 | 3.58 | 3.43 | 0.67 | -0.24 | -0.27 | 6.31 | -0.27 | -0.08 | -0.2 | -0.25 |
| Run1_ICUS24 | | 9.52 | 1.69 | 1.16 | 1.74 | 2.42 | 6.49 | 99.29 | 136.16 | 119.49 | 36.01 | 51 | 29.52 | 8.18 | -0.1 | -0.24 | 1.93 | 2.94 | 3.53 | -0.03 | -0.27 | -0.28 | 3.12 | -0.2 | -0.15 | -0.27 | -0.26 |
| Run1_Nl-1 | | 4.53 | 1.06 | 0.49 | 1.1 | 1.68 | 5.17 | 51.45 | 97.03 | 88.65 | 30.81 | 37.1 | 18.74 | 3.36 | -0.12 | -0.19 | 4.17 | 2.19 | 2.59 | 0.23 | -0.33 | -0.26 | 2.21 | -0.15 | -0.15 | -0.34 | -0.33 |
| Run1_Nl-3 | | 2.51 | 1.42 | 1.13 | 1.41 | 1.86 | 9.2 | 62.9 | 69.06 | 52.57 | 16.27 | 17.5 | 4.99 | 1.02 | 0.62 | 0.2 | 2.73 | 2.4 | 2.36 | 0.66 | -0.3 | -0.22 | 2.5 | -0.18 | -0.14 | -0.3 | -0.3 |
| Run1_Nl-4 | | 4.98 | 3.08 | 1.59 | 3.04 | 2.71 | 9.09 | 79.55 | 75.28 | 53.74 | 19.55 | 22.29 | 8.93 | 1.97 | 0.77 | 0.28 | 6.43 | 2.61 | 3.29 | 1.13 | -0.29 | -0.23 | 2.89 | -0.18 | -0.08 | -0.3 | -0.27 |
| Run1_Nl-6 | | 10.22 | 1.34 | 1.21 | 0.86 | 1.71 | 8.3 | 140.23 | 163.41 | 143.18 | 50.24 | 58.52 | 31.31 | 6.16 | 0.29 | -0.09 | 10.27 | 3.21 | 3.32 | 0.64 | -0.26 | -0.24 | 3.67 | -0.2 | -0.03 | -0.21 | -0.27 |
| Run2_Nl-3 | Normal | 1.96 | 1.29 | 1.08 | 1.02 | 1.87 | 8.68 | 60.7 | 62.65 | 50.27 | 14.6 | 12.21 | 2.3 | 0.71 | 0.14 | 0.06 | 1.17 | 2.01 | 2.04 | 0.3 | -0.29 | -0.25 | 2.05 | -0.21 | -0.16 | -0.29 | -0.29 |
| Run2_Nl-4 | | 4.25 | 2.39 | 2.14 | 1.89 | 2.18 | 11.72 | 71.99 | 65.56 | 49 | 18.81 | 17.4 | 6.02 | 1.39 | 0.41 | 0.09 | 3.7 | 2.29 | 2.91 | 0.99 | -0.27 | -0.27 | 2.85 | -0.19 | -0.07 | -0.29 | -0.27 |
| Run2_Nl-5 | | 2.09 | 1.08 | 1.19 | 0.87 | 1.98 | 11.91 | 91.31 | 78.89 | 62.34 | 14.74 | 19.59 | 5.69 | 1.28 | 0.31 | -0.23 | 1.85 | 2 | 1.68 | 0.21 | -0.3 | -0.26 | 1.34 | -0.15 | -0.23 | -0.31 | -0.3 |
| Run2_Nl-6 | | 9.92 | 1.22 | 0.99 | 0.96 | 1.59 | 9.48 | 143.8 | 178.25 | 158.49 | 49.81 | 58.59 | 27.91 | 4.88 | 0.27 | -0.19 | 3.26 | 2.56 | 2.89 | 0.58 | -0.25 | -0.26 | 2.95 | -0.24 | -0.08 | -0.25 | -0.26 |

**Supplemental Table 3**: Median expression of each surface marker on each cell population.

### CD16 Median

| | | Ungated | CD34+CD38low | HSCs | MPP | CMP/GMP | Myelo/Mono-Blasts | ProMonocytes | CD14neg Monocytes | Mature Monocytes | ProMyelocytes | Myelocytes | MetaMyelocytes | MatureGrans | ProErythroblasts | Erythroblasts | LateErythroblasts | PreBcells | Mature Bcells | Plasma Cells | T cells | NK cells | pDCs | Basophils | Platelets | CD8+ T cells | CD8neg T cells |
|---|---|---|---|---|---|---|---|---|---|---|---|---|---|---|---|---|---|---|---|---|---|---|---|---|---|---|---|
| Run1_MDS17 | | 1 | 0.11 | 0.14 | 0.03 | 0.55 | 0.04 | 0.41 | 1.07 | 1.63 | 1.05 | 1.52 | 3.08 | 12.61 | 0.08 | 0.2 | 0.6 | 0.87 | -0.06 | 0.1 | 0.17 | 4.55 | 0.07 | 0.5 | 1.63 | 0.04 | 0.35 |
| Run2_MDS15* | AML/RAEB-T | 0.92 | -0.22 | -0.21 | -0.25 | -0.09 | -0.04 | 0.12 | 4.98 | 2.01 | 0.36 | 0.74 | 2.06 | 24.68 | -0.14 | 1.04 | 1.49 | 4.69 | -0.15 | -0.15 | 0.31 | 2.78 | 0.2 | 0.49 | 1.08 | 0.13 | 0.46 |
| Run2_MDS4 | | 0.84 | -0.19 | -0.18 | -0.15 | -0.38 | -0.26 | -0.21 | -0.03 | 0.46 | 0.52 | 1.18 | 4.8 | 19.88 | -0.13 | 0.07 | 0.32 | 0.08 | -0.12 | -0.26 | -0.04 | 0.14 | -0.13 | 1.21 | 3.09 | -0.1 | 0.05 |
| Run1_MDS21 | | 0.85 | 0.16 | 0.16 | 0.09 | 0.17 | 0.18 | 0.33 | 0.7 | 2.04 | 0.82 | 1.62 | 4.27 | 14.48 | 0.28 | 0.75 | 8.31 | 0.07 | -0.02 | -0.45 | 0.53 | 2.06 | 0.91 | 0.99 | 6.63 | 0.37 | 0.6 |
| Run1_MDS3 | | 0.83 | 0.07 | -0.07 | -0.01 | 0.1 | 0.17 | 0.2 | 0.61 | 1.56 | 0.9 | 1.86 | 3.38 | 13.97 | 0.27 | 0.66 | 3.4 | 0.14 | -0.01 | 0.08 | 0.55 | 3.47 | 1.13 | 1.22 | 4.63 | 0.29 | 0.67 |
| Run2_MDS13 | | 0.46 | -0.24 | -0.22 | -0.25 | -0.17 | -0.15 | 0.11 | 0.33 | 1 | 0.75 | 1.47 | 2.91 | 13.65 | -0.11 | 0.07 | 0.3 | -0.1 | -0.21 | -0.21 | 0.19 | 1.82 | 1 | 0.64 | 2.31 | 0.04 | 0.27 |
| Run2_MDS16 | | 6.44 | -0.1 | -0.12 | -0.16 | -0.03 | 0.65 | 2.68 | 0.42 | 0.5 | 1.29 | 2.46 | 5.58 | 25.06 | -0.1 | 0.12 | 5.92 | 0.49 | -0.16 | -0.17 | 0.25 | 84.39 | 0.66 | 0.61 | 5.36 | 0.11 | 0.28 |
| Run2_MDS1 | | 0.81 | -0.02 | -0.09 | -0.08 | -0.03 | -0.12 | -0.07 | 0.22 | 0.7 | 0.27 | 0.91 | 3.89 | 16.07 | -0.01 | 0.31 | 0.22 | 0.76 | 0.14 | -0.13 | 0.25 | 2.65 | 0.21 | 0.2 | 3.84 | -0.01 | 0.3 |
| Run2_MDS21 | Higher Risk | 0.5 | -0.16 | -0.09 | -0.17 | -0.14 | -0.13 | 0.15 | 0.4 | 1.29 | 0.87 | 1.59 | 3.44 | 14.97 | -0.08 | -0.08 | 0.09 | -0.11 | -0.15 | -0.08 | 0.24 | 0.7 | 1.77 | 0.51 | 4.6 | 0.12 | 0.31 |
| Run2_MDS26* | | 1.77 | 0 | -0.02 | -0.04 | 0.03 | 0.02 | 0.74 | 0.64 | 1.4 | 0.39 | 0.79 | 3.61 | 13.46 | 0.24 | 0.33 | -0.03 | 0.59 | -0.16 | -0.13 | 0.24 | 17.96 | 0.81 | 0.86 | 1.5 | 0.05 | 0.34 |
| Run2_MDS27 | | 0.96 | -0.14 | -0.14 | -0.18 | 0.15 | -0.02 | 0.88 | 0.37 | 0.72 | 1.22 | 1.78 | 5.15 | 26.92 | -0.03 | -0.03 | -0.02 | 0.47 | -0.17 | -0.21 | 0.11 | 36.73 | 25.67 | 0.3 | 3.92 | 0 | 0.24 |
| Run2_MDS2 | | 2.14 | -0.29 | -0.29 | -0.32 | -0.25 | -0.17 | -0.06 | 0.34 | 0.31 | 0.61 | 0.97 | 4.4 | 16.36 | 0.35 | 1.96 | 2.51 | 0.99 | -0.22 | 0.17 | 0.13 | 9.68 | 1.71 | 0.45 | 4.28 | 0.01 | 0.13 |
| Run2_MDS3 | | 0.51 | -0.15 | -0.2 | -0.13 | -0.18 | -0.15 | 0.16 | 0.29 | 1.36 | 0.92 | 1.78 | 2.71 | 13.34 | -0.13 | -0.04 | -0.09 | -0.1 | -0.17 | -0.31 | 0.25 | 2.15 | 0.44 | 0.89 | 3.47 | 0.1 | 0.32 |
| Run1_MDS23 | | 3.37 | 0.09 | 0.06 | 0.17 | 0.01 | -0.1 | 0.18 | 0.44 | 0.91 | 1.17 | 1.76 | 4.64 | 16.71 | 0.51 | 0.75 | 2.84 | 0.87 | -0.15 | 0.03 | 0.62 | 1.28 | 0.09 | 1.05 | 4.2 | 0.36 | 0.72 |
| Run1_MDS25 | | 21.72 | 6.18 | -0.08 | 8.92 | 0.11 | 0.72 | 0.51 | 0.63 | 1.42 | 1.37 | 2.68 | 6.65 | 30.26 | 1.05 | 3.02 | 19.02 | 0.8 | -0.06 | -0.17 | 0.52 | 22.21 | 0.24 | 0.94 | 5.44 | 0.41 | 0.66 |
| Run1_MDS5 | | 19.56 | 3.69 | -0.9 | 1.41 | -0.03 | -0.09 | 0.65 | 1.47 | 1.55 | 1.3 | 7.26 | 6.19 | 36.25 | 1.42 | 1.42 | 6.64 | 5.83 | 0.35 | -0.14 | 0.56 | 9.43 | 1.33 | 1.31 | 3.1 | 0.34 | 0.64 |
| Run1_MDS6 | | 16.49 | -0.07 | -0.17 | -0.2 | -0.15 | -0.08 | 0.26 | 0.43 | 1.07 | 1.71 | 2.37 | 4.99 | 32.47 | 0.13 | 0.36 | 15.15 | 0.45 | -0.06 | -0.08 | 0.54 | 3.82 | 1.05 | 1.51 | 8 | 0.47 | 0.6 |
| Run1_MDS8 | | 14.19 | 1.53 | 1.53 | 0.84 | -0.03 | 0.88 | 2.1 | 0.61 | 1.16 | 1.63 | 2.67 | 6.74 | 23.27 | 0.46 | 0.99 | 5.29 | 1.13 | 0.15 | -0.19 | 0.4 | 12.88 | 1.76 | 1.32 | 6.71 | 0.31 | 0.43 |
| Run1_MDS9 | | 5.98 | 0.16 | 0.06 | 0.13 | 0.17 | 0.69 | 10.53 | 1.13 | 1.52 | 1.68 | 2.41 | 5.97 | 20.87 | 0.37 | 0.7 | 11.19 | 1.33 | 0.09 | 0.05 | 0.66 | 20.94 | 16.43 | 1.33 | 4 | 0.55 | 0.72 |
| Run2_MDS12 | Lower Risk | 3.21 | -0.32 | -0.27 | -0.3 | -0.13 | -0.29 | -0.04 | 0.28 | 0.72 | 0.62 | 1.29 | 4.12 | 15.31 | -0.16 | -0.03 | 0.61 | 0.23 | -0.1 | -0.16 | 0.05 | 0.56 | -0.16 | 0.29 | 4.5 | -0.02 | 0.12 |
| Run2_MDS14 | | 30.99 | 2.85 | 0.62 | 2.79 | 0.2 | 0.26 | 0.62 | 0.2 | 0.64 | 1.52 | 2.48 | 4.64 | 50.15 | -0.14 | -0.08 | 37.39 | 0.28 | -0.1 | -0.14 | 0.26 | 2.12 | -0.08 | 0.91 | 3.34 | 0.06 | 0.38 |
| Run2_MDS19 | | 6.5 | 0.08 | 0.38 | 0 | -0.16 | -0.13 | 0.18 | 0.3 | 0.67 | 1.08 | 2.21 | 4.89 | 24.2 | -0.14 | -0.06 | 0.14 | 0.73 | -0.13 | 0.07 | 0.4 | 7.73 | 0.01 | 0.5 | 7.24 | 0.09 | 0.51 |
| Run2_MDS20 | | 11.2 | -0.12 | -0.32 | -0.3 | 0.03 | 1.31 | 3.46 | 0.22 | 0.58 | 1.12 | 1.9 | 5.29 | 44.31 | -0.04 | 0.61 | 24.94 | 0.68 | -0.14 | -0.14 | 0.38 | 1.76 | 36.05 | 0.64 | 5.11 | 0.15 | 0.4 |
| Run2_MDS28* | | 6.6 | -0.01 | -0.07 | -0.03 | -0.07 | -0.25 | -0.15 | 0.23 | 0.5 | 1.34 | 1.93 | 4.97 | 23.89 | 0.12 | 0.19 | 5.91 | 0.64 | -0.14 | -0.15 | 0.24 | 1.88 | 1.15 | 0.8 | 4.61 | 0.11 | 0.37 |
| Run2_MDS5 | | 16.72 | 2.74 | -0.04 | 0.31 | -0.09 | -0.04 | 0.5 | 0.88 | 0.87 | 0.88 | 6.58 | 5.72 | 32.49 | -0.23 | 0.83 | 7.54 | 4.57 | -0.15 | -0.18 | 0.22 | 8.12 | 0.54 | 0.64 | 2.92 | -0.02 | 0.3 |
| Run1_ICUS18 | | 27.18 | 0.82 | -0.31 | 0.25 | 0.32 | 1.95 | 1.62 | 0.47 | 0.98 | 1.59 | 2.56 | 6.05 | 36.28 | 0.56 | 0.84 | 42.41 | 1.33 | -0.03 | -0.18 | 0.77 | 29.78 | 0.4 | 0.99 | 6.07 | 0.59 | 0.81 |
| Run1_ICUS22 | ICUS | 18.61 | 0.35 | 0.7 | -0.06 | 0.3 | 0.01 | 0.57 | 0.46 | 1.1 | 1.93 | 2.56 | 5.5 | 40.84 | 0.36 | 0.9 | 27.38 | 0.74 | -0.08 | 0.05 | 0.72 | 3.63 | 1.94 | 1.18 | 5.66 | 0.48 | 0.81 |
| Run1_ICUS24 | | 5.96 | 0.42 | 0.47 | 0.36 | 0.17 | -0.04 | 0.21 | 0.53 | 1.13 | 1.84 | 2.14 | 4.97 | 23.51 | 0.17 | 0.46 | 9.98 | 0.35 | -0.1 | -0.11 | 0.56 | 4.38 | 0.07 | 1.2 | 8.84 | 0.33 | 0.77 |
| Run1_Nl-1 | | 10.18 | 0.36 | 0.45 | 0.2 | 0.16 | 0.33 | 0.89 | 0.52 | 1.1 | 1.82 | 2.05 | 4.91 | 35.21 | 0.39 | 0.73 | 18.78 | 0.5 | 0.09 | -0.08 | 0.72 | 5.88 | 0.52 | 1.56 | 11.18 | 0.5 | 0.84 |
| Run1_Nl-3 | | 5.15 | 0.46 | 0.41 | 0.23 | -0.02 | -0.03 | 0.25 | 0.64 | 1.18 | 1.4 | 2.25 | 4.94 | 19.01 | 0.67 | 0.53 | 11.87 | 0.39 | -0.08 | -0.16 | 0.38 | 1.77 | 0.3 | 1.05 | 2.73 | 0.14 | 0.54 |
| Run1_Nl-4 | | 5.35 | 0.17 | 0.36 | 0.14 | 0.15 | -0.05 | 0.21 | 0.52 | 1 | 1.42 | 2.1 | 4.85 | 18.37 | 0.7 | 0.71 | 10.09 | 0.4 | -0.11 | -0.07 | 0.4 | 6.07 | -0.03 | 1.38 | 2.51 | 0.19 | 0.55 |
| Run1_Nl-6 | | 5.24 | 0.39 | 0.38 | 0.24 | 0.24 | -0.01 | 0.24 | 0.43 | 0.82 | 1.21 | 1.86 | 4.77 | 24.18 | 0.27 | 0.41 | 6.91 | 0.33 | -0.11 | -0.15 | 0.48 | 2.65 | 0.41 | 0.87 | 2.19 | 0.15 | 0.57 |
| Run2_Nl-3 | Normal | 3.59 | -0.04 | 0.16 | -0.16 | -0.06 | -0.17 | 0.03 | 0.23 | 0.57 | 0.89 | 1.94 | 4.56 | 16.32 | -0.01 | 0.18 | 7.94 | 0.1 | -0.17 | -0.23 | 0.05 | 1.29 | -0.06 | 0.59 | 2.87 | -0.1 | 0.19 |
| Run2_Nl-4 | | 3.8 | 0.06 | 0.38 | -0.07 | -0.13 | -0.21 | -0.02 | 0.1 | 0.47 | 0.91 | 1.78 | 4.26 | 16.34 | 0.1 | 0.32 | 6.75 | 0.01 | -0.18 | -0.23 | 0.06 | 5.4 | -0.17 | 0.47 | 3.23 | -0.05 | 0.18 |
| Run2_Nl-5 | | 3.61 | -0.08 | -0.09 | -0.1 | -0.02 | -0.13 | -0.01 | 0.15 | 0.52 | 0.9 | 1.95 | 4.86 | 19.83 | -0.06 | -0.04 | 8.79 | 0.11 | -0.14 | -0.23 | 0.01 | 1.6 | -0.02 | 0.67 | 2.97 | -0.08 | 0.14 |
| Run2_Nl-6 | | 4.61 | 0.05 | 0.22 | 0.01 | -0.07 | -0.19 | -0.13 | 0.01 | 0.45 | 0.64 | 1.37 | 4.34 | 20.23 | -0.13 | -0.01 | 2.91 | -0.04 | -0.2 | -0.24 | 0.13 | 2.52 | 0.09 | 0.52 | 3.03 | -0.04 | 0.2 |

**Supplemental Table 3**: Median expression of each surface marker on each cell population.

### CD11b Median

| Sample | Group | Ungated | CD34+CD38low | HSCs | MPP | CMP/GMP | Myelo/Mono-Blasts | ProMonocytes | CD14neg Monocytes | Mature Monocytes | ProMyelocytes | Myelocytes | MetaMyelocytes | MatureGrans | ProErythroblasts |
|---|---|---|---|---|---|---|---|---|---|---|---|---|---|---|---|
| Run1_MDS17 | | 3.02 | -0.48 | -0.53 | -0.51 | -0.39 | -0.45 | 11.69 | 105.53 | 114.71 | 0.79 | 22.36 | 251.8 | 386.32 | -0.48 |
| Run2_MDS15* | AML/RAEB-T | 64.26 | -0.38 | -0.4 | -0.37 | -0.32 | -0.15 | 12.52 | 230.23 | 209.51 | 0.92 | 26.88 | 166.97 | 332.63 | -0.3 |
| Run2_MDS4 | | 8.97 | -0.34 | -0.35 | -0.35 | -0.29 | -0.03 | 14.62 | 74.43 | 86.67 | 0.97 | 24.61 | 206.87 | 325.35 | -0.31 |
| Run1_MDS21 | | 0.93 | -0.48 | -0.58 | -0.55 | -0.58 | -0.55 | 9.22 | 123.08 | 149.93 | 0.9 | 16.93 | 255.86 | 429.5 | -0.46 |
| Run1_MDS3 | | 1.06 | -0.48 | -0.66 | -0.53 | -0.59 | -0.57 | 8.24 | 69.25 | 122.84 | 1.1 | 16.95 | 189.8 | 321.13 | -0.48 |
| Run2_MDS13 | | 0.62 | -0.35 | -0.3 | -0.36 | -0.35 | -0.34 | 13.07 | 94.99 | 104.66 | 0.57 | 18.58 | 246.85 | 357.39 | -0.32 |
| Run2_MDS16 | | 87 | -0.28 | -0.29 | -0.3 | -0.28 | 0.32 | 12.52 | 79.9 | 113.3 | 1.27 | 30.96 | 146.2 | 212.8 | -0.29 |
| Run2_MDS1 | | 14.17 | -0.27 | -0.59 | -0.25 | -0.36 | -0.27 | 11.71 | 153.04 | 222.77 | 1.47 | 30.19 | 193.83 | 298.99 | -0.31 |
| Run2_MDS21 | Higher Risk | 0.5 | -0.34 | -0.32 | -0.34 | -0.34 | -0.33 | 14.16 | 151.92 | 180.75 | 0.9 | 18.5 | 298.8 | 457.66 | -0.28 |
| Run2_MDS26* | | 85.45 | -0.24 | -0.19 | -0.27 | -0.31 | -0.25 | 8.96 | 151.44 | 209.18 | 1.75 | 25.47 | 253.11 | 413.86 | -0.22 |
| Run2_MDS27 | | 1.29 | -0.31 | -0.31 | -0.34 | -0.26 | -0.22 | 13.23 | 110.75 | 193.36 | 0.76 | 25.32 | 184.36 | 318.13 | -0.33 |
| Run2_MDS2 | | 109.5 | -0.41 | -0.37 | -0.44 | -0.41 | -0.3 | 13.55 | 130.71 | 142.76 | 0.84 | 30.03 | 319.84 | 369.39 | 0.13 |
| Run2_MDS3 | | 0.6 | -0.3 | -0.26 | -0.3 | -0.35 | -0.33 | 13.46 | 103.15 | 87.99 | 0.87 | 18.85 | 191.22 | 316.29 | -0.29 |
| Run1_MDS23 | | 127.3 | -0.61 | -0.46 | -0.59 | -0.82 | -0.41 | 13.7 | 102.1 | 171.14 | 1.93 | 26.93 | 206.29 | 348.67 | -0.47 |
| Run1_MDS25 | | 215.56 | 119.05 | -0.92 | 125.57 | -0.8 | 0.09 | 14.71 | 105.72 | 165.25 | 1.25 | 30.1 | 167.24 | 255.55 | -0.34 |
| Run1_MDS5 | | 96.51 | 2.65 | -1.76 | 1.11 | -0.77 | -0.38 | 16.39 | 145.9 | 161.17 | 1.54 | 37.35 | 111.6 | 131.84 | 0.57 |
| Run1_MDS6 | | 166.45 | -0.6 | -0.58 | -0.63 | -0.87 | -0.45 | 13.2 | 100.94 | 180.1 | 1.25 | 28.26 | 141.82 | 248.89 | -0.33 |
| Run1_MDS8 | | 182.92 | 0.68 | -0.46 | 0.53 | -0.63 | -0.1 | 14.38 | 119 | 181.85 | 1.37 | 28.91 | 169.23 | 251.69 | -0.46 |
| Run1_MDS9 | | 82.73 | -0.56 | -0.67 | -0.62 | -0.65 | -0.29 | 12.29 | 84.65 | 135.33 | 1.72 | 24.22 | 160.99 | 274.6 | -0.46 |
| Run2_MDS12 | Lower Risk | 198.4 | -0.35 | -0.29 | -0.36 | -0.35 | -0.17 | 14.12 | 91.94 | 126.55 | 1.85 | 29.21 | 287.08 | 467.96 | -0.34 |
| Run2_MDS14 | | 233.89 | 0.31 | -0.43 | -0.18 | -0.58 | 0.26 | 13.23 | 103.53 | 189.69 | 2.21 | 29.28 | 152.68 | 307.61 | -0.34 |
| Run2_MDS19 | | 122.31 | -0.25 | -0.2 | -0.27 | -0.39 | -0.12 | 13.72 | 89.34 | 130.82 | 1.56 | 25.65 | 143.02 | 238.19 | -0.33 |
| Run2_MDS20 | | 173.59 | -0.09 | -0.39 | 0.28 | -0.31 | 0.6 | 13.54 | 114.38 | 170.29 | 1.37 | 25.86 | 185.43 | 360.41 | -0.22 |
| Run2_MDS28* | | 262.32 | -0.29 | -0.28 | -0.26 | -0.33 | -0.07 | 13.98 | 176.22 | 215.3 | 1.67 | 24.84 | 240.5 | 479.16 | -0.22 |
| Run2_MDS5 | | 101 | 6.71 | 0.05 | 0.35 | -0.12 | 0.04 | 15.85 | 152.69 | 166.3 | 1.47 | 35.65 | 120.85 | 141.81 | -0.34 |
| Run1_ICUS18 | | 256.33 | -0.27 | -0.76 | -0.77 | -0.72 | 0.09 | 13.3 | 93.9 | 176.28 | 1.5 | 28.26 | 180.25 | 320.6 | -0.4 |
| Run1_ICUS22 | ICUS | 337.3 | -0.58 | -0.72 | -0.7 | -0.76 | -0.19 | 13.61 | 123.05 | 224.3 | 1.91 | 23.61 | 220.46 | 460.41 | -0.47 |
| Run1_ICUS24 | | 191.74 | -0.32 | -0.22 | -0.52 | -0.71 | -0.41 | 14.04 | 101.56 | 194.34 | 1.72 | 24.55 | 207.08 | 395.68 | -0.52 |
| Run1_Nl-1 | | 228.64 | 0.1 | 0.2 | -0.13 | -0.43 | -0.28 | 14 | 71.39 | 173.28 | 1.34 | 26.03 | 184.55 | 370.35 | -0.45 |
| Run1_Nl-3 | | 92.37 | -0.36 | -0.56 | -0.7 | -0.51 | -0.4 | 14.15 | 75.3 | 160.54 | 1.27 | 32.13 | 115.85 | 171.35 | -0.39 |
| Run1_Nl-4 | | 117.91 | -0.31 | -0.72 | -0.73 | -0.75 | -0.39 | 14.5 | 95.19 | 158.61 | 1.2 | 29.2 | 139.46 | 204.71 | -0.3 |
| Run1_Nl-6 | | 114.93 | -0.42 | -0.48 | -0.46 | -0.73 | -0.43 | 13.9 | 71.24 | 126.28 | 1.25 | 27.38 | 140.17 | 232.36 | -0.51 |
| Run2_Nl-3 | Normal | 73.26 | -0.31 | -0.32 | -0.4 | -0.32 | -0.17 | 13.42 | 92.88 | 148.6 | 1.33 | 31.75 | 112.19 | 151.2 | -0.33 |
| Run2_Nl-4 | | 95.21 | -0.31 | -0.23 | -0.3 | -0.3 | -0.14 | 14.33 | 99.55 | 127.53 | 1.26 | 30.63 | 132.54 | 170.99 | -0.27 |
| Run2_Nl-5 | | 84.02 | -0.22 | -0.28 | -0.25 | -0.32 | -0.13 | 13.67 | 107.55 | 157.07 | 0.85 | 28.43 | 144.69 | 186.26 | -0.27 |
| Run2_Nl-6 | | 125.12 | -0.29 | -0.34 | -0.26 | -0.31 | -0.22 | 13.94 | 81.14 | 124.72 | 1.25 | 27.64 | 163.64 | 231.23 | -0.31 |

| Sample | Erythroblasts | LateErythroblasts | PreBcells | Mature Bcells | Plasma Cells | T cells | NK cells | pDCs | Basophils | Platelets | CD8+ T cells | CD8neg T cells |
|---|---|---|---|---|---|---|---|---|---|---|---|---|
| Run1_MDS17 | -0.45 | 0.3 | 2.2 | -0.5 | -1.19 | 1.7 | 18.97 | -0.02 | 67.04 | 0.76 | 3.53 | -0.22 |
| Run2_MDS15* | 7.65 | 117.2 | 182.92 | -0.25 | -0.53 | 0.57 | 11.69 | 0.35 | 10.37 | -0.1 | 2.32 | -0.23 |
| Run2_MDS4 | -0.25 | 0.72 | 0.03 | -0.28 | -0.35 | 0.39 | 8.56 | -0.23 | 50.56 | 1.53 | 1.89 | -0.29 |
| Run1_MDS21 | -0.23 | 2.98 | -0.41 | -0.57 | -0.61 | 0.14 | 15.52 | -0.1 | 54.82 | 0.41 | 2.22 | -0.39 |
| Run1_MDS3 | -0.23 | 2.03 | -0.43 | -0.62 | -0.99 | 0.16 | 18.22 | 6.46 | 31.7 | 0.19 | 2.57 | -0.43 |
| Run2_MDS13 | -0.3 | -0.08 | -0.27 | -0.27 | -0.49 | 0.16 | 10.01 | 1.26 | 30.63 | 0.1 | 2.3 | -0.23 |
| Run2_MDS16 | -0.3 | 32.39 | 0.17 | -0.34 | -0.56 | -0.15 | 30.55 | 0.06 | 14.1 | 0.34 | 2.06 | -0.3 |
| Run2_MDS1 | -0.21 | -0.06 | 2.85 | 0.02 | -0.42 | -0.12 | 46.36 | -0.21 | 13.19 | 0.35 | 2.03 | -0.24 |
| Run2_MDS21 | -0.33 | -0.15 | -0.31 | -0.25 | -0.7 | 0 | 3.41 | -0.11 | 46.27 | 0.48 | 1.87 | -0.24 |
| Run2_MDS26* | -0.29 | -0.26 | 1 | -0.31 | -0.49 | 0.15 | 34.55 | 0.36 | 32.12 | 2.24 | 1.8 | -0.24 |
| Run2_MDS27 | -0.33 | -0.15 | 0.81 | -0.34 | -0.43 | 0.61 | 17.81 | 1.52 | 6.51 | 0.41 | 1.96 | -0.26 |
| Run2_MDS2 | 5.85 | 1.67 | 0.53 | 0.35 | -0.38 | -0.23 | 19.57 | 3.82 | 6.38 | -0.05 | 1.91 | -0.27 |
| Run2_MDS3 | -0.33 | -0.23 | -0.3 | -0.25 | -0.4 | 0 | 13.99 | 0.2 | 39.66 | 0.21 | 2.18 | -0.26 |
| Run1_MDS23 | -0.37 | 47.23 | 6.55 | -0.46 | -1.23 | -0.08 | 23.35 | -0.76 | 25.72 | 2.23 | 1.95 | -0.44 |
| Run1_MDS25 | 0.51 | 244.3 | 6.98 | -0.81 | -0.93 | 0.95 | 13.34 | -0.59 | 16.87 | -0.06 | 2.71 | -0.33 |
| Run1_MDS5 | 0.25 | 15.55 | 44.83 | -0.25 | -1.28 | -0.05 | 9.05 | 25.04 | 24.53 | 2.36 | 1.36 | -0.37 |
| Run1_MDS6 | -0.52 | 106.13 | 0.42 | -0.55 | -1.21 | 0.47 | 26.58 | 1.6 | 49.18 | 0.8 | 2.14 | -0.43 |
| Run1_MDS8 | -0.36 | 2.42 | 2.09 | -0.58 | -0.24 | -0.12 | 20.63 | 9.21 | 20.47 | 0.6 | 1.18 | -0.39 |
| Run1_MDS9 | -0.48 | 13.91 | 2.78 | -0.51 | -1.4 | 0.28 | 23.08 | 2.01 | 15.01 | 1.08 | 2.29 | -0.39 |
| Run2_MDS12 | -0.31 | 1.52 | 7.59 | -0.3 | -0.43 | -0.01 | 10.31 | -0.27 | 16.96 | 1.13 | 1.22 | -0.28 |
| Run2_MDS14 | -0.31 | 248.44 | -0.16 | -0.33 | -0.41 | 0.15 | 25.66 | -0.12 | 9.05 | 0.21 | 2.03 | -0.23 |
| Run2_MDS19 | -0.29 | -0.04 | 4.29 | -0.34 | -0.43 | -0.08 | 15.01 | -0.32 | 23.76 | 0.67 | 2.29 | -0.26 |
| Run2_MDS20 | -0.2 | 259.81 | 1.08 | -0.11 | -0.28 | -0.08 | 7.83 | 1.69 | 12.23 | 0.29 | 2.19 | -0.15 |
| Run2_MDS28* | -0.22 | 238.77 | 36.01 | -0.27 | -0.35 | 0.4 | 38.26 | 13.64 | 21.46 | 0.41 | 2 | -0.24 |
| Run2_MDS5 | 0.03 | 82.63 | 22.61 | -0.19 | -0.55 | 0.07 | 6.25 | 24.6 | 22.67 | 1.2 | 1.27 | -0.17 |
| Run1_ICUS18 | -0.3 | 353.01 | 3.42 | -0.49 | -1.35 | -0.17 | 44.39 | -0.58 | 21.17 | 0.42 | 2.71 | -0.44 |
| Run1_ICUS22 | -0.16 | 393.24 | 0.09 | -0.75 | -0.8 | 0.01 | 18.86 | 0.61 | 22.2 | 0.34 | 1.98 | -0.39 |
| Run1_ICUS24 | -0.4 | 4.54 | -0.44 | -0.67 | -1 | 0.55 | 20.54 | -0.67 | 19.43 | 0.74 | 1.75 | -0.2 |
| Run1_Nl-1 | -0.4 | 267.85 | -0.54 | -0.62 | -1.4 | 0.24 | 39.4 | -0.38 | 16.97 | 0.66 | 2.03 | -0.36 |
| Run1_Nl-3 | -0.4 | 170.51 | -0.05 | -0.57 | -0.98 | 0.35 | 21.92 | -0.44 | 12.93 | 0.48 | 2.26 | -0.44 |
| Run1_Nl-4 | -0.37 | 205.55 | -0.15 | -0.63 | -1.17 | 0.55 | 25.65 | -0.76 | 12.89 | 0.09 | 1.94 | -0.26 |
| Run1_Nl-6 | -0.45 | 172 | -0.59 | -0.7 | -1.18 | -0.08 | 33.72 | -0.31 | 12.34 | -0.28 | 1.89 | -0.47 |
| Run2_Nl-3 | -0.26 | 133.45 | -0.07 | -0.35 | -0.43 | 0.13 | 17.55 | -0.28 | 8.4 | 0.29 | 1.85 | -0.29 |
| Run2_Nl-4 | -0.26 | 161.76 | -0.11 | -0.35 | -0.42 | 0.42 | 20.67 | -0.3 | 8 | 0.14 | 1.55 | -0.18 |
| Run2_Nl-5 | -0.31 | 163.52 | -0.23 | -0.32 | -0.44 | 0.36 | 13.86 | -0.34 | 12.81 | 0.13 | 1.99 | -0.28 |
| Run2_Nl-6 | -0.27 | 101.18 | -0.27 | -0.37 | -0.44 | -0.09 | 35.51 | -0.19 | 8.08 | 0.16 | 1.55 | -0.29 |

**Supplemental Table 3**: Median expression of each surface marker on each cell population.

**CD15 Median**

| | | Ungated | CD34+CD38low | HSCs | MPP | CMP/GMP | Myelo/Mono-Blasts | ProMonocytes | CD14neg Monocytes | Mature Monocytes | ProMyelocytes | Myelocytes | MetaMyelocytes | MatureGrans | ProErythroblasts | Erythroblasts | LateErythroblasts | PreBcells | Mature Bcells | Plasma Cells | T cells | NK cells | pDCs | Basophils | Platelets | CD8+ T cells | CD8neg T cells |
|---|---|---|---|---|---|---|---|---|---|---|---|---|---|---|---|---|---|---|---|---|---|---|---|---|---|---|---|
| Run1_MDS17 | | 2.09 | 0.78 | 0.9 | 0.69 | 0.92 | 0.45 | 0.65 | 0.87 | 0.81 | 23.61 | 20.9 | 15.77 | 18.54 | -0.1 | -0.18 | 0.5 | 1.19 | -0.3 | -0.52 | -0.22 | 0.09 | 0.65 | 2.15 | 1.53 | -0.21 | -0.24 |
| Run2_MDS15* | AML/RAEB-T | 1.48 | 1.04 | 1.17 | 0.84 | 0.34 | 0.57 | 0.55 | 0.18 | -0.03 | 11.7 | 12.53 | 16.96 | 23.39 | -0.3 | 0.82 | 1.85 | 0.9 | -0.33 | -0.33 | -0.26 | -0.15 | 0.41 | 0.22 | 0.89 | -0.25 | -0.26 |
| Run2_MDS4 | | 1.48 | 0.61 | 0.93 | 0.55 | -0.62 | -0.19 | -0.23 | -0.14 | 0.4 | 23.18 | 46.34 | 156.69 | 194.36 | -0.22 | -0.12 | 1.01 | -0.17 | -0.26 | -0.43 | -0.3 | -0.23 | -0.28 | 0.16 | 2.87 | -0.3 | -0.3 |
| Run1_MDS21 | | 1.35 | 0.8 | 0.67 | 0.72 | 0.63 | 0.57 | 1.27 | 0.51 | 0.91 | 41.07 | 78.24 | 107.87 | 155.14 | -0.16 | 0.02 | 3.93 | -0.2 | -0.31 | -0.7 | -0.23 | 0.14 | 1.08 | 0.6 | 1.15 | -0.2 | -0.24 |
| Run1_MDS3 | | 2.21 | 0.99 | 0.96 | 0.87 | 0.87 | 0.69 | 1.03 | 0.12 | 0.67 | 50.46 | 102.68 | 118.15 | 184.26 | -0.2 | 0.03 | 3.33 | -0.17 | -0.3 | -0.67 | -0.27 | 0.08 | 2.8 | 0.68 | 0.72 | -0.27 | -0.27 |
| Run2_MDS13 | | 1.69 | 0.52 | 0.68 | 0.45 | 0.3 | 0.28 | 0.17 | 0.1 | 0.49 | 41.68 | 71.16 | 76.66 | 83.1 | -0.25 | -0.24 | 0.23 | -0.23 | -0.29 | -0.39 | -0.26 | -0.2 | 0.78 | 0.15 | 0.91 | -0.25 | -0.27 |
| Run2_MDS16 | | 169.58 | 1.28 | 1.51 | 1 | 0.94 | -0.18 | -0.14 | -0.11 | -0.05 | 130.42 | 150.41 | 347.39 | 372.46 | 0.4 | -0.26 | 6.2 | -0.09 | -0.31 | -0.48 | -0.26 | 0 | -0.09 | -0.13 | 0.71 | -0.24 | -0.26 |
| Run2_MDS1 | | 0.58 | 0.12 | 0.42 | -0.08 | -0.12 | -0.25 | -0.18 | -0.18 | 0.04 | 12.54 | 16.81 | 37.34 | 67.79 | -0.17 | -0.13 | -0.06 | -0.05 | -0.2 | -0.43 | -0.29 | -0.07 | -0.26 | 0.03 | 0.8 | -0.32 | -0.29 |
| Run2_MDS21 | Higher Risk | 0.83 | 0.39 | 0.47 | 0.37 | 0.36 | 0.28 | 0.62 | 0.27 | 0.66 | 74.86 | 105.68 | 149.68 | 188.04 | -0.23 | -0.24 | -0.07 | -0.26 | -0.31 | -0.29 | -0.26 | -0.23 | 0.04 | -0.15 | 0.98 | -0.25 | -0.26 |
| Run2_MDS26* | | 24.66 | 0.44 | 0.55 | 0.31 | 0.29 | 0.13 | 0.66 | 0.43 | 0.15 | 17.46 | 24.03 | 79.29 | 102.73 | 0 | -0.19 | -0.16 | 0.19 | -0.26 | -0.41 | -0.25 | -0.05 | -0.12 | -0.05 | 5.88 | -0.25 | -0.25 |
| Run2_MDS27 | | 0.79 | 0.94 | 1.3 | 0.65 | 0.25 | -0.06 | -0.09 | -0.02 | 0.05 | 72.82 | 124.46 | 376.59 | 476.38 | -0.08 | -0.3 | -0.13 | -0.07 | -0.28 | -0.42 | -0.26 | -0.05 | -0.27 | -0.2 | 1.8 | -0.25 | -0.27 |
| Run2_MDS2 | | 3.08 | 0.01 | 0.57 | -0.05 | -0.1 | -0.22 | -0.26 | 0.04 | -0.15 | 16.95 | 19.34 | 33.76 | 70.12 | -0.13 | 2.01 | 1.09 | 0.05 | 0.21 | -0.13 | -0.29 | -0.15 | -0.16 | -0.34 | -0.15 | -0.28 | -0.29 |
| Run2_MDS3 | | 1.55 | 0.62 | 1.02 | 0.53 | 0.46 | 0.41 | 0.68 | 0.45 | 1.39 | 69.86 | 144.56 | 173.91 | 240.35 | -0.21 | -0.25 | -0.13 | -0.26 | -0.24 | -0.4 | -0.29 | -0.13 | 1.26 | 0.44 | 0.89 | -0.26 | -0.3 |
| Run1_MDS23 | | 37.92 | 0.47 | 0.92 | 0.5 | -0.13 | -0.27 | -0.14 | -0.08 | 0.07 | 17.92 | 32.75 | 96.84 | 107.17 | -0.2 | -0.07 | 3.7 | 0.07 | -0.27 | -0.62 | -0.25 | -0.14 | -0.24 | -0.01 | 2.15 | -0.26 | -0.25 |
| Run1_MDS25 | | 201.19 | 22.96 | 1.73 | 164.3 | -0.63 | -0.44 | -0.29 | 0.01 | 0.48 | 60.28 | 91.98 | 223.39 | 250.57 | -0.15 | 0.31 | 185.5 | -0.02 | -0.43 | -0.7 | -0.24 | 0.05 | -0.33 | -0.16 | 0.53 | -0.23 | -0.25 |
| Run1_MDS5 | | 12.9 | 4.42 | -0.24 | 0.72 | -0.21 | -0.27 | -0.11 | 0.63 | 0.43 | 28.38 | 20.61 | 26.54 | 24.4 | 2.62 | 0.33 | 4.8 | 1.05 | -0.25 | -0.57 | -0.23 | -0.05 | 0.02 | -0.04 | 2.23 | -0.3 | -0.21 |
| Run1_MDS6 | | 162.34 | 0.65 | 0.58 | 0.56 | 0.01 | -0.1 | -0.11 | 0.01 | 0.19 | 126.02 | 179.51 | 312.58 | 208.68 | -0.09 | -0.22 | 103.39 | -0.15 | -0.3 | -0.59 | -0.23 | 0.03 | 0.01 | 0.19 | 1.77 | -0.22 | -0.24 |
| Run1_MDS8 | | 247.68 | 1.95 | 1.19 | 3.01 | 0.52 | -0.1 | -0.17 | -0.13 | 0.2 | 86.43 | 168.41 | 310.93 | 361.7 | -0.21 | -0.03 | 2.48 | -0.04 | -0.26 | -0.68 | -0.19 | 0.03 | 0.08 | -0.07 | 1.52 | -0.18 | -0.19 |
| Run1_MDS9 | | 133.35 | 0.86 | 1.1 | 0.6 | 0.53 | 0.05 | -0.18 | -0.02 | 0.39 | 71.02 | 118.64 | 295.63 | 369.15 | -0.08 | -0.13 | 11.09 | -0.06 | -0.3 | -0.62 | -0.22 | 0.13 | -0.01 | -0.1 | 1.64 | -0.21 | -0.23 |
| Run2_MDS12 | Lower Risk | 64.74 | -0.15 | 0.13 | 0.02 | -0.01 | -0.3 | -0.11 | 0.09 | 0.19 | 24.47 | 29.64 | 122.12 | 164.81 | -0.24 | -0.26 | 0.73 | 0.01 | -0.28 | -0.57 | -0.27 | -0.18 | -0.26 | -0.19 | 1.24 | -0.26 | -0.28 |
| Run2_MDS14 | | 223.59 | 4.73 | 0.89 | 0.19 | 0.3 | -0.27 | -0.25 | -0.12 | -0.04 | 349.72 | 235.12 | 382.84 | 274 | -0.16 | -0.21 | 242.32 | -0.27 | -0.31 | -0.43 | -0.28 | -0.22 | -0.09 | -0.06 | 0.66 | -0.28 | -0.28 |
| Run2_MDS19 | | 383.05 | 1.38 | 2.25 | 1.11 | 0.45 | -0.19 | -0.14 | -0.04 | 0 | 141.12 | 262.89 | 549.41 | 572.28 | -0.18 | -0.18 | 0.34 | 0 | -0.29 | -0.53 | -0.25 | -0.16 | -0.31 | -0.1 | 0.89 | -0.23 | -0.26 |
| Run2_MDS20 | | 128.3 | 1.53 | 1.69 | 0.99 | 0.45 | -0.28 | -0.27 | -0.23 | -0.06 | 81.38 | 115.6 | 301.14 | 335.36 | -0.23 | -0.13 | 227.26 | -0.18 | -0.35 | -0.51 | -0.25 | -0.17 | -0.17 | -0.12 | 0.59 | -0.24 | -0.25 |
| Run2_MDS28* | | 286.11 | 0.86 | 1.19 | 1.04 | 0.32 | -0.15 | -0.1 | -0.04 | -0.09 | 78.33 | 134.66 | 412.76 | 468.07 | -0.08 | -0.09 | 274.52 | 0.16 | -0.3 | -0.54 | -0.26 | -0.1 | -0.23 | -0.09 | 1.01 | -0.23 | -0.28 |
| Run2_MDS5 | | 16.55 | 2.64 | 0.47 | 0.82 | -0.28 | -0.22 | -0.2 | 0.22 | 0.28 | 28.26 | 21.33 | 34.9 | 30.81 | -0.14 | -0.03 | 6.14 | 0.71 | -0.26 | -0.58 | -0.25 | -0.22 | -0.07 | -0.02 | 1.74 | -0.31 | -0.22 |
| Run1_ICUS18 | | 338.68 | 2.48 | 3.66 | 1.58 | 1.35 | -0.3 | -0.24 | -0.14 | 0.14 | 91.34 | 166.49 | 434.51 | 430.85 | 0 | 0.08 | 481.86 | -0.02 | -0.27 | -0.46 | -0.24 | 0.28 | -0.26 | 0.14 | 0.83 | -0.23 | -0.24 |
| Run1_ICUS22 | ICUS | 425.18 | 1.51 | 2.05 | 1.22 | 0.39 | -0.34 | -0.2 | -0.1 | 0.15 | 146.67 | 227.73 | 540.41 | 559.38 | 0.05 | 0.44 | 499.71 | -0.2 | -0.34 | -0.63 | -0.2 | -0.12 | -0.24 | 0.27 | 1.11 | -0.2 | -0.19 |
| Run1_ICUS24 | | 115.05 | 1.49 | 2.01 | 1.04 | 0.25 | -0.17 | -0.08 | 0.04 | 0.21 | 107.05 | 106.57 | 198.34 | 185.68 | -0.09 | -0.06 | 4.36 | -0.29 | -0.37 | -0.64 | -0.21 | -0.08 | -0.26 | 0.09 | 1.56 | -0.21 | -0.21 |
| Run1_Nl-1 | | 175.19 | 1.66 | 1.45 | 1.28 | 0.13 | -0.28 | -0.07 | -0.09 | 0.14 | 46.38 | 80.9 | 275.05 | 264.87 | -0.13 | -0.08 | 212.44 | -0.37 | -0.35 | -0.71 | -0.24 | 0.05 | -0.17 | 0.16 | 1.22 | -0.23 | -0.26 |
| Run1_Nl-3 | | 153.56 | 2.28 | 2.31 | 1.79 | 0.75 | -0.01 | 0.15 | 0.25 | 0.31 | 34.68 | 87.95 | 279.46 | 253.98 | 0.29 | -0.11 | 287.64 | -0.15 | -0.29 | -0.56 | -0.23 | 0.09 | -0.11 | -0.09 | 1 | -0.23 | -0.23 |
| Run1_Nl-4 | | 116.38 | 2.63 | 2.14 | 2.52 | 0.64 | -0.09 | 0.21 | 0.32 | 0.35 | 35.9 | 57.69 | 222.75 | 174.2 | 0.17 | -0.06 | 230.38 | -0.22 | -0.33 | -0.68 | -0.22 | 0.02 | -0.24 | 0.04 | 1.09 | -0.2 | -0.24 |
| Run1_Nl-6 | | 111.01 | 1.94 | 2.44 | 1.43 | 0.37 | -0.11 | -0.04 | 0.03 | 0.15 | 28.48 | 77.1 | 255.23 | 180.52 | -0.12 | -0.17 | 174.58 | -0.33 | -0.35 | -0.55 | -0.22 | 0.14 | -0.17 | -0.18 | 0.53 | -0.2 | -0.23 |
| Run2_Nl-3 | Normal | 199.67 | 1.73 | 2.28 | 1.6 | 0.31 | -0.13 | 0.01 | 0.02 | 0.1 | 41.85 | 124.95 | 359.35 | 368.06 | -0.12 | -0.18 | 322.06 | -0.2 | -0.31 | -0.44 | -0.27 | -0.12 | -0.18 | -0.2 | 0.79 | -0.26 | -0.28 |
| Run2_Nl-4 | | 167.6 | 1.6 | 1.7 | 1.06 | 0.2 | -0.14 | 0.01 | 0.02 | 0.13 | 46.42 | 82.86 | 295.65 | 269.18 | -0.14 | -0.14 | 289.69 | -0.27 | -0.32 | -0.43 | -0.26 | -0.13 | -0.29 | -0.15 | 0.92 | -0.25 | -0.27 |
| Run2_Nl-5 | | 156.93 | 1.27 | 1.26 | 1.17 | 0.17 | -0.1 | -0.03 | 0 | 0.18 | 46.99 | 138.53 | 426.41 | 325.58 | -0.04 | -0.21 | 288.97 | -0.31 | -0.27 | -0.41 | -0.27 | -0.11 | -0.19 | -0.14 | 0.58 | -0.26 | -0.27 |
| Run2_Nl-6 | | 190.45 | 1.31 | 1.71 | 1.15 | 0.28 | -0.17 | -0.18 | -0.13 | 0.02 | 33.67 | 93.6 | 319.58 | 314.48 | -0.17 | -0.22 | 146.68 | -0.29 | -0.34 | -0.44 | -0.28 | -0.06 | -0.27 | -0.19 | 0.8 | -0.28 | -0.28 |

**Supplemental Table 3**: Median expression of each surface marker on each cell population.

### CD123 Median

| | | Ungated | CD34+CD38low | HSCs | MPP | CMP/GMP | Myelo/Mono-Blasts | ProMonocytes | CD14neg Monocytes | Mature Monocytes | ProMyelocytes | Myelocytes | MetaMyelocytes | MatureGrans | ProErythroblasts | Erythroblasts | LateErythroblasts | PreBcells | Mature Bcells | Plasma Cells | T cells | NK cells | pDCs | Basophils | Platelets | CD8+ T cells | CD8neg T cells |
|---|---|---|---|---|---|---|---|---|---|---|---|---|---|---|---|---|---|---|---|---|---|---|---|---|---|---|---|
| Run1_MDS17 | | -0.18 | 0.94 | 1.05 | 0.89 | 1.06 | 0.48 | 0.52 | -0.11 | -0.26 | -0.42 | -0.23 | -0.13 | -0.05 | -0.39 | -0.46 | -0.41 | -0.05 | -0.09 | 0.22 | -0.46 | -0.46 | 37.73 | 20.83 | -0.41 | -0.46 | -0.45 |
| Run2_MDS15* | AML/RAEB-T | 0.85 | 2.31 | 1.62 | 1.53 | 4.28 | 1.64 | 0.6 | -0.11 | -0.01 | 0.75 | 2.28 | 1.75 | -0.14 | 1.56 | 0.14 | 1.5 | 0.19 | 0.1 | 0.85 | -0.4 | -0.35 | 31.95 | 38.23 | 1.09 | -0.4 | -0.4 |
| Run2_MDS4 | | -0.26 | 1.88 | 1.28 | 1.29 | 0.35 | 0.91 | 1.06 | 0.38 | 0.23 | -0.38 | -0.39 | -0.37 | -0.36 | -0.32 | -0.4 | -0.35 | 0.56 | 0.91 | 0.23 | -0.42 | -0.29 | 130.56 | 43.29 | -0.13 | -0.41 | -0.43 |
| Run1_MDS21 | | -0.33 | 1.08 | 1 | 1.02 | 1.36 | 1.14 | 0.66 | -0.3 | -0.25 | -0.42 | -0.37 | -0.38 | -0.29 | -0.46 | -0.41 | -0.28 | -0.31 | -0.14 | 0.15 | -0.48 | -0.42 | 65.34 | 46.06 | -0.41 | -0.45 | -0.49 |
| Run1_MDS3 | | -0.33 | 1.24 | 1.19 | 1.07 | 1.45 | 1.27 | 0.92 | -0.06 | 0.56 | -0.39 | -0.33 | -0.32 | -0.2 | -0.45 | -0.44 | -0.29 | -0.34 | -0.08 | 0.51 | -0.49 | -0.34 | 46.13 | 41.98 | -0.36 | -0.49 | -0.49 |
| Run2_MDS13 | | -0.21 | 1.56 | 1.77 | 1.3 | 1.39 | 1.15 | 0.14 | -0.08 | -0.1 | -0.2 | -0.04 | -0.28 | -0.31 | -0.17 | -0.42 | -0.34 | -0.28 | -0.21 | 0.85 | -0.42 | -0.42 | 43.78 | 31.57 | -0.34 | -0.41 | -0.42 |
| Run2_MDS16 | | -0.33 | 1.53 | 1.42 | 1.25 | 1.85 | 2.88 | 2.37 | 0.65 | 0.45 | -0.26 | -0.32 | -0.34 | -0.34 | 0.75 | -0.42 | -0.33 | -0.2 | -0.21 | -0.04 | -0.42 | -0.37 | 111.7 | 59.77 | -0.36 | -0.42 | -0.42 |
| Run2_MDS1 | | -0.24 | 2.49 | 3.05 | 2.37 | 2.51 | 2.72 | 1.36 | 0.74 | 0.74 | -0.3 | -0.3 | -0.33 | -0.36 | -0.16 | -0.41 | -0.39 | -0.11 | -0.2 | 0.13 | -0.43 | -0.39 | 83.2 | 41.21 | -0.29 | -0.44 | -0.43 |
| Run2_MDS21 | Higher Risk | -0.3 | 0.81 | 1.11 | 0.74 | 1.08 | 0.76 | 0.49 | -0.18 | -0.12 | -0.35 | -0.3 | -0.34 | -0.32 | -0.34 | -0.43 | -0.39 | -0.33 | -0.21 | 0.18 | -0.42 | -0.41 | 66.49 | 50.84 | -0.36 | -0.41 | -0.43 |
| Run2_MDS26* | | -0.37 | 0.62 | 0.7 | 0.56 | 0.77 | -0.12 | -0.2 | -0.15 | 0.06 | -0.39 | -0.36 | -0.36 | -0.36 | -0.21 | -0.42 | -0.4 | -0.28 | -0.14 | -0.25 | -0.42 | -0.41 | 38.92 | 48.44 | -0.4 | -0.43 | -0.42 |
| Run2_MDS27 | | -0.26 | 0.71 | 0.68 | 0.61 | 0.9 | 0.96 | 1.96 | 0.71 | 0.49 | -0.27 | -0.08 | -0.12 | -0.24 | 0.65 | -0.42 | -0.39 | -0.22 | 0.2 | 0.3 | -0.38 | -0.4 | 84.07 | 249.82 | -0.2 | -0.36 | -0.39 |
| Run2_MDS2 | | -0.23 | 1.19 | 1.89 | 0.91 | 1.34 | 1.54 | 1.45 | 0.2 | 0.42 | 0.12 | -0.36 | -0.33 | -0.32 | 0.87 | -0.28 | -0.32 | -0.17 | 0.8 | 1.52 | -0.43 | -0.41 | 44.36 | 54.88 | -0.34 | -0.41 | -0.43 |
| Run2_MDS3 | | -0.26 | 1.02 | 1.31 | 0.83 | 1.17 | 0.95 | 0.33 | -0.02 | 0.33 | -0.26 | -0.18 | -0.23 | -0.21 | -0.29 | -0.4 | -0.4 | -0.3 | -0.16 | 0.53 | -0.42 | -0.37 | 91.6 | 31.36 | -0.33 | -0.43 | -0.42 |
| Run1_MDS23 | | -0.32 | 0.83 | 0.79 | 0.7 | 2.95 | 2.67 | 2.71 | 1.49 | 1.11 | -0.36 | -0.35 | -0.39 | -0.4 | -0.41 | -0.4 | -0.16 | -0.06 | -0.08 | 0.18 | -0.48 | -0.43 | 74.98 | 38.4 | -0.38 | -0.46 | -0.49 |
| Run1_MDS25 | | -0.36 | 2.05 | 3.35 | 0.32 | 4.24 | 3.63 | 2.31 | 0.72 | 0.35 | -0.36 | -0.38 | -0.42 | -0.4 | -0.38 | -0.38 | -0.14 | -0.01 | 0.52 | 0.27 | -0.44 | -0.39 | 241.64 | 34.53 | -0.45 | -0.44 | -0.44 |
| Run1_MDS5 | | -0.2 | 2.13 | 1.59 | 0.46 | 2.41 | 1.67 | 1.14 | 1.04 | 0.27 | 0.31 | -0.09 | -0.01 | -0.18 | -0.06 | -0.38 | -0.26 | -0.01 | -0.23 | 0.45 | -0.44 | -0.42 | 70.62 | 54.51 | -0.43 | -0.43 | -0.44 |
| Run1_MDS6 | | -0.33 | 2.28 | 1.66 | 1.72 | 2.29 | 3.46 | 4.05 | 1.24 | 0.68 | -0.34 | -0.3 | -0.32 | -0.36 | -0.06 | -0.48 | -0.31 | -0.15 | -0.06 | 0.75 | -0.46 | -0.47 | 88.49 | 82.5 | -0.38 | -0.46 | -0.47 |
| Run1_MDS8 | | -0.37 | 0.74 | 0.51 | 0.27 | 0.91 | 1.06 | 1.29 | 0.46 | 0.01 | -0.4 | -0.41 | -0.38 | -0.37 | -0.4 | -0.42 | -0.36 | -0.33 | -0.31 | 0.26 | -0.5 | -0.44 | 68.34 | 144.27 | -0.41 | -0.49 | -0.5 |
| Run1_MDS9 | | -0.34 | 1.39 | 1.53 | 0.94 | 1.92 | 1.89 | 4.5 | 0.76 | 0.43 | -0.38 | -0.37 | -0.36 | -0.33 | -0.35 | -0.45 | -0.24 | -0.29 | 0.23 | -0.02 | -0.48 | -0.43 | 48.15 | 116.85 | -0.41 | -0.46 | -0.49 |
| Run2_MDS12 | Lower Risk | -0.26 | 6.14 | 3.24 | 5 | 4.69 | 3.79 | 1.73 | 0.87 | 0.81 | -0.35 | -0.28 | -0.29 | -0.3 | -0.33 | -0.42 | -0.34 | 0.32 | -0.01 | -0.18 | -0.39 | -0.39 | 160.43 | 55.23 | -0.26 | -0.38 | -0.4 |
| Run2_MDS14 | | -0.3 | 0.71 | 0.61 | -0.29 | 0.6 | 2.66 | 3.11 | 1.02 | 0.67 | -0.26 | -0.26 | -0.28 | -0.34 | -0.35 | -0.41 | -0.26 | 0.14 | 0.08 | -0.15 | -0.42 | -0.38 | 129.2 | 50.41 | -0.29 | -0.41 | -0.43 |
| Run2_MDS19 | | -0.31 | 0.92 | 1.02 | 0.8 | 1.27 | 3.69 | 2.69 | 0.9 | 0.54 | -0.33 | -0.31 | -0.33 | -0.33 | -0.34 | -0.43 | -0.39 | -0.13 | -0.19 | 0.2 | -0.41 | -0.38 | 182.18 | 57.36 | -0.41 | -0.4 | -0.42 |
| Run2_MDS20 | | -0.31 | 0.78 | 0.27 | 1.09 | 1.62 | 4.98 | 4.32 | 1.32 | 0.91 | -0.32 | -0.31 | -0.33 | -0.32 | -0.13 | -0.36 | -0.29 | -0.25 | 0.07 | 1.13 | -0.42 | -0.38 | 59.07 | 153.65 | -0.37 | -0.41 | -0.42 |
| Run2_MDS28* | | -0.16 | 1.39 | 0.81 | 1.3 | 1.42 | 3.54 | 3.68 | 1.83 | 2.12 | -0.27 | -0.22 | -0.17 | -0.22 | -0.04 | -0.39 | -0.25 | 0.35 | 0.29 | 0.06 | -0.42 | -0.35 | 60.26 | 60.57 | -0.34 | -0.43 | -0.42 |
| Run2_MDS5 | | -0.22 | 2.02 | 0.76 | 1.39 | 1.88 | 1.34 | 0.16 | 0.84 | 0.25 | -0.05 | -0.18 | -0.14 | -0.22 | 0.06 | -0.37 | -0.25 | -0.06 | -0.24 | 0.65 | -0.39 | -0.4 | 62.38 | 50.96 | -0.4 | -0.38 | -0.39 |
| Run1_ICUS18 | | -0.33 | 1.23 | 1.67 | 1.39 | 1.84 | 5.24 | 4.08 | 1.13 | 0.46 | -0.28 | -0.26 | -0.31 | -0.34 | -0.39 | -0.43 | -0.28 | -0.26 | 0 | -0.08 | -0.47 | -0.44 | 214.64 | 51.07 | -0.41 | -0.46 | -0.47 |
| Run1_ICUS22 | ICUS | -0.25 | 1.59 | 1.56 | 1.26 | 2 | 4.81 | 5.27 | 1.58 | 0.99 | -0.32 | -0.25 | -0.23 | -0.31 | -0.38 | -0.39 | -0.23 | 0.13 | 0.2 | -0.14 | -0.45 | -0.4 | 72.46 | 268.83 | -0.29 | -0.43 | -0.46 |
| Run1_ICUS24 | | -0.33 | 1.32 | 1.1 | 1.36 | 2.09 | 3.52 | 3.54 | 1.25 | 0.64 | -0.32 | -0.29 | -0.36 | -0.38 | -0.39 | -0.46 | -0.34 | 0.83 | 0.89 | -0.09 | -0.47 | -0.45 | 135.65 | 33.64 | -0.39 | -0.48 | -0.46 |
| Run1_Nl-1 | | -0.33 | 0.75 | 1.03 | 0.53 | 1.03 | 3.49 | 4.57 | 2.31 | 0.92 | -0.37 | -0.35 | -0.34 | -0.37 | -0.38 | -0.4 | -0.31 | 0.3 | 0.37 | 0.52 | -0.48 | -0.43 | 143.79 | 59.29 | -0.37 | -0.47 | -0.48 |
| Run1_Nl-3 | | -0.28 | 0.71 | 0.56 | 0.62 | 1.35 | 4.17 | 3.08 | 1.09 | 0.55 | -0.28 | -0.26 | -0.27 | -0.32 | -0.34 | -0.43 | -0.24 | 0.17 | 0.3 | 0.2 | -0.48 | -0.43 | 117.66 | 90.76 | -0.41 | -0.48 | -0.47 |
| Run1_Nl-4 | | -0.31 | 1.39 | 1.88 | 1.22 | 1.89 | 4.58 | 4.39 | 1.69 | 0.7 | -0.33 | -0.31 | -0.32 | -0.33 | -0.27 | -0.43 | -0.25 | 0.25 | 0.23 | 0.69 | -0.48 | -0.42 | 140.95 | 58.02 | -0.46 | -0.47 | -0.49 |
| Run1_Nl-6 | | -0.22 | 0.87 | 0.86 | 0.76 | 1.2 | 2.15 | 2.24 | 1.18 | 1.09 | -0.34 | -0.22 | -0.21 | -0.22 | -0.36 | -0.45 | -0.24 | 0.79 | 3.92 | 0.36 | -0.47 | -0.44 | 112.56 | 103.66 | -0.41 | -0.49 | -0.46 |
| Run2_Nl-3 | Normal | -0.26 | 0.88 | 0.77 | 0.78 | 1.15 | 3.06 | 1.75 | 0.5 | 0.29 | -0.27 | -0.27 | -0.27 | -0.29 | -0.31 | -0.41 | -0.26 | 0.07 | 0.17 | 0.26 | -0.43 | -0.39 | 99.95 | 77.94 | -0.36 | -0.43 | -0.43 |
| Run2_Nl-4 | | -0.27 | 1.1 | 0.65 | 0.9 | 1.45 | 3.53 | 2.53 | 0.64 | 0.44 | -0.29 | -0.31 | -0.29 | -0.28 | -0.3 | -0.39 | -0.26 | 0.11 | 0.07 | 0.76 | -0.43 | -0.41 | 125.34 | 52.12 | -0.38 | -0.42 | -0.43 |
| Run2_Nl-5 | | -0.27 | 0.72 | 0.83 | 0.76 | 0.82 | 1.83 | 1.12 | 0.19 | 0.03 | -0.28 | -0.25 | -0.25 | -0.27 | -0.31 | -0.4 | -0.29 | -0.03 | -0.08 | 0.04 | -0.42 | -0.41 | 98.78 | 76.69 | -0.39 | -0.42 | -0.43 |
| Run2_Nl-6 | | -0.21 | 0.69 | 0.71 | 0.57 | 0.99 | 1.74 | 1.39 | 0.68 | 0.77 | -0.3 | -0.23 | -0.22 | -0.22 | -0.31 | -0.41 | -0.31 | 0.44 | 2.96 | -0.02 | -0.4 | -0.42 | 102.72 | 125.15 | -0.25 | -0.42 | -0.39 |

**Supplemental Table 3**: Median expression of each surface marker on each cell population.

### CD3 Median

| | | Ungated | CD34+CD38low | HSCs | MPP | CMP/GMP | Myelo/Mono-Blasts | ProMonocytes | CD14neg Monocytes | Mature Monocytes | ProMyelocytes | Myelocytes | MetaMyelocytes | MatureGrans | ProErythroblasts | Erythroblasts | LateErythroblasts | PreBcells | Mature Bcells | Plasma Cells | T cells | NK cells | pDCs | Basophils | Platelets | CD8+ T cells | CD8neg T cells |
|---|---|---|---|---|---|---|---|---|---|---|---|---|---|---|---|---|---|---|---|---|---|---|---|---|---|---|---|
| Run1_MDS17 | | -0.41 | -0.49 | -0.5 | -0.49 | -0.48 | -0.51 | -0.45 | -0.46 | -0.4 | -0.47 | -0.47 | -0.44 | -0.41 | -0.5 | -0.5 | -0.47 | -0.39 | -0.48 | -0.46 | 35.58 | -0.41 | -0.42 | -0.45 | -0.47 | 34.44 | 37.48 |
| Run2_MDS15* | AML/RAEB-T | -0.4 | -0.43 | -0.45 | -0.42 | -0.46 | -0.44 | -0.43 | -0.41 | -0.4 | -0.42 | -0.43 | -0.41 | -0.39 | -0.41 | -0.34 | -0.34 | -0.4 | -0.39 | -0.56 | 33.57 | -0.4 | -0.39 | -0.45 | -0.44 | 34.15 | 32.61 |
| Run2_MDS4 | | -0.3 | -0.43 | -0.43 | -0.41 | -0.42 | -0.44 | -0.45 | -0.43 | -0.47 | -0.45 | -0.46 | -0.44 | -0.42 | -0.48 | -0.44 | -0.45 | -0.31 | -0.43 | -0.4 | 14.81 | -0.13 | -0.45 | -0.45 | -0.38 | 14.24 | 15.48 |
| Run1_MDS21 | | -0.28 | -0.49 | -0.52 | -0.49 | -0.5 | -0.5 | -0.49 | -0.43 | -0.38 | -0.46 | -0.43 | -0.45 | -0.43 | -0.49 | -0.44 | -0.37 | -0.38 | -0.41 | -0.45 | 23.25 | -0.37 | -0.47 | -0.52 | -0.46 | 24.78 | 22.51 |
| Run1_MDS3 | | -0.31 | -0.51 | -0.52 | -0.54 | -0.51 | -0.52 | -0.47 | -0.45 | -0.52 | -0.47 | -0.48 | -0.45 | -0.44 | -0.51 | -0.45 | -0.4 | -0.43 | -0.45 | -0.32 | 31.68 | -0.49 | 0.34 | -0.57 | -0.49 | 30.79 | 32.23 |
| Run2_MDS13 | | -0.29 | -0.46 | -0.45 | -0.45 | -0.44 | -0.44 | -0.45 | -0.41 | -0.37 | -0.42 | -0.41 | -0.42 | -0.42 | -0.41 | -0.44 | -0.4 | -0.32 | -0.4 | -0.4 | 20.04 | -0.4 | -0.32 | -0.41 | -0.43 | 20.37 | 19.86 |
| Run2_MDS16 | | -0.28 | -0.44 | -0.45 | -0.45 | -0.36 | -0.41 | -0.52 | -0.46 | -0.42 | -0.43 | -0.45 | -0.43 | -0.43 | -0.44 | -0.44 | -0.39 | 18.59 | -0.43 | -0.35 | 25.96 | -0.39 | -0.4 | -0.42 | -0.44 | 22.78 | 26.85 |
| Run2_MDS1 | | -0.29 | -0.38 | -0.41 | -0.45 | -0.43 | -0.46 | -0.41 | -0.45 | -0.44 | -0.45 | -0.46 | -0.43 | -0.44 | -0.45 | -0.45 | -0.43 | -0.09 | -0.43 | -0.39 | 17.77 | -0.43 | -0.48 | -0.38 | -0.38 | 15.93 | 18.17 |
| Run2_MDS21 | Higher Risk | -0.28 | -0.45 | -0.44 | -0.45 | -0.43 | -0.43 | -0.49 | -0.43 | -0.46 | -0.41 | -0.42 | -0.42 | -0.42 | -0.41 | -0.43 | -0.43 | -0.35 | -0.43 | -0.44 | 14.85 | -0.07 | -0.56 | -0.46 | -0.43 | 15.95 | 14.24 |
| Run2_MDS26* | | -0.41 | -0.4 | -0.41 | -0.42 | -0.43 | -0.43 | -0.38 | -0.41 | -0.31 | -0.43 | -0.43 | -0.42 | -0.41 | -0.44 | -0.44 | -0.43 | -0.21 | -0.4 | -0.4 | 20.13 | -0.36 | -0.54 | -0.43 | -0.42 | 17.17 | 22.33 |
| Run2_MDS27 | | -0.28 | -0.44 | -0.45 | -0.43 | -0.46 | -0.43 | -0.43 | -0.4 | -0.4 | -0.44 | -0.42 | -0.44 | -0.42 | -0.44 | -0.44 | -0.43 | 8.73 | -0.42 | -0.45 | 17.31 | -0.38 | -0.42 | -0.4 | -0.31 | 15.66 | 19.75 |
| Run2_MDS2 | | -0.3 | -0.47 | -0.51 | -0.48 | -0.48 | -0.43 | -0.43 | -0.42 | -0.44 | -0.34 | -0.42 | -0.44 | -0.42 | -0.22 | -0.28 | -0.44 | -0.03 | -0.25 | -0.39 | 11.84 | -0.3 | -0.46 | -0.46 | -0.46 | 11.06 | 11.87 |
| Run2_MDS3 | | -0.28 | -0.44 | -0.45 | -0.45 | -0.45 | -0.44 | -0.5 | -0.42 | -0.41 | -0.42 | -0.4 | -0.41 | -0.35 | -0.45 | -0.45 | -0.43 | -0.33 | -0.42 | -0.48 | 20.41 | -0.38 | -0.58 | -0.39 | -0.47 | 19.86 | 20.71 |
| Run1_MDS23 | | -0.39 | -0.48 | -0.4 | -0.46 | -0.46 | -0.47 | -0.45 | -0.42 | -0.4 | -0.42 | -0.45 | -0.44 | -0.43 | -0.49 | -0.48 | -0.41 | -0.24 | -0.43 | -0.44 | 46.71 | -0.39 | -0.48 | -0.46 | -0.44 | 40.62 | 49.42 |
| Run1_MDS25 | | -0.41 | -0.21 | -0.73 | -0.65 | -0.29 | -0.48 | -0.46 | -0.47 | -0.44 | -0.48 | -0.48 | -0.49 | -0.47 | -0.5 | -0.49 | -0.33 | -0.06 | -0.47 | -0.46 | 30.43 | -0.46 | -0.51 | -0.51 | -0.48 | 28.32 | 32.87 |
| Run1_MDS5 | | -0.38 | -0.56 | -0.75 | -0.47 | -0.71 | -0.57 | -0.55 | -0.43 | -0.4 | -0.53 | -0.45 | -0.43 | -0.42 | -0.32 | -0.44 | -0.38 | -0.22 | -0.44 | -0.49 | 36.12 | -0.43 | -0.43 | -0.47 | -0.46 | 34.78 | 36.57 |
| Run1_MDS6 | | -0.36 | -0.47 | -0.46 | -0.46 | -0.51 | -0.44 | -0.4 | -0.44 | -0.42 | -0.48 | -0.48 | -0.44 | -0.43 | -0.41 | -0.47 | -0.41 | -0.11 | -0.41 | -0.44 | 40.31 | -0.43 | -0.42 | -0.47 | -0.46 | 39.03 | 41.52 |
| Run1_MDS8 | | -0.34 | -0.28 | -0.11 | -0.36 | -0.45 | -0.45 | -0.41 | -0.38 | -0.38 | -0.47 | -0.5 | -0.46 | -0.43 | -0.48 | -0.45 | -0.43 | 1.93 | -0.38 | -0.38 | 24.66 | -0.42 | -0.32 | -0.39 | -0.5 | 25.74 | 24.25 |
| Run1_MDS9 | | -0.27 | -0.46 | -0.46 | -0.46 | -0.47 | -0.48 | -0.44 | -0.4 | -0.33 | -0.46 | -0.48 | -0.45 | -0.43 | -0.45 | -0.5 | -0.39 | 16.52 | -0.4 | -0.39 | 28.28 | -0.41 | -0.37 | -0.45 | -0.47 | 26.41 | 29.68 |
| Run2_MDS12 | Lower Risk | -0.4 | -0.39 | -0.47 | -0.35 | -0.45 | -0.41 | -0.41 | -0.41 | -0.37 | -0.41 | -0.43 | -0.41 | -0.4 | -0.44 | -0.45 | -0.43 | -0.3 | -0.41 | -0.6 | 20.23 | -0.35 | -0.41 | -0.44 | -0.42 | 20.83 | 19.91 |
| Run2_MDS14 | | -0.39 | -0.29 | -0.2 | -0.19 | -0.46 | -0.42 | -0.41 | -0.44 | -0.41 | -0.42 | -0.45 | -0.42 | -0.43 | -0.47 | -0.45 | -0.43 | -0.32 | -0.43 | -0.55 | 20.33 | -0.4 | -0.36 | -0.46 | -0.46 | 18.76 | 21.08 |
| Run2_MDS19 | | -0.37 | -0.46 | -0.51 | -0.45 | -0.45 | -0.43 | -0.42 | -0.42 | -0.41 | -0.43 | -0.43 | -0.42 | -0.41 | -0.45 | -0.44 | -0.42 | 0.63 | -0.39 | -0.41 | 23.58 | -0.38 | -0.44 | -0.41 | -0.4 | 22.59 | 23.97 |
| Run2_MDS20 | | -0.2 | -0.33 | -0.43 | -0.32 | -0.61 | -0.43 | -0.44 | -0.43 | -0.41 | -0.44 | -0.45 | -0.44 | -0.42 | -0.41 | -0.4 | -0.43 | 24.8 | -0.43 | -0.53 | 34.54 | -0.42 | -0.47 | -0.43 | -0.45 | 28.42 | 35.29 |
| Run2_MDS28* | | -0.35 | -0.41 | -0.4 | -0.38 | -0.42 | -0.42 | -0.4 | -0.4 | -0.38 | -0.42 | -0.43 | -0.4 | -0.38 | -0.4 | -0.43 | -0.4 | -0.11 | -0.41 | -0.42 | 29.69 | -0.38 | -0.39 | -0.39 | -0.39 | 26.6 | 33.15 |
| Run2_MDS5 | | -0.36 | -0.5 | -0.87 | -0.55 | -0.75 | -0.4 | -0.39 | -0.4 | -0.38 | -0.41 | -0.42 | -0.41 | -0.39 | -0.33 | -0.4 | -0.35 | -0.14 | -0.38 | -0.48 | 28.46 | -0.37 | -0.43 | -0.33 | -0.42 | 28.01 | 28.81 |
| Run1_ICUS18 | | -0.34 | -0.37 | -0.3 | -0.37 | -0.46 | -0.5 | -0.45 | -0.45 | -0.43 | -0.46 | -0.47 | -0.46 | -0.44 | -0.48 | -0.45 | -0.39 | 0.97 | -0.49 | -0.42 | 39.16 | -0.44 | -0.42 | -0.45 | -0.45 | 31.46 | 40.97 |
| Run1_ICUS22 | ICUS | -0.36 | -0.4 | -0.3 | -0.45 | -0.51 | -0.45 | -0.4 | -0.44 | -0.4 | -0.45 | -0.47 | -0.45 | -0.42 | -0.46 | -0.46 | -0.35 | -0.31 | -0.45 | -0.34 | 34.68 | -0.42 | -0.41 | -0.48 | -0.4 | 32.61 | 35.4 |
| Run1_ICUS24 | | -0.34 | -0.5 | -0.49 | -0.56 | -0.48 | -0.47 | -0.41 | -0.43 | -0.39 | -0.45 | -0.47 | -0.45 | -0.43 | -0.48 | -0.47 | -0.41 | -0.39 | -0.45 | -0.44 | 36.41 | -0.42 | -0.43 | -0.43 | -0.45 | 35.33 | 37.31 |
| Run1_Nl-1 | | -0.39 | -0.44 | -0.42 | -0.37 | -0.4 | -0.48 | -0.42 | -0.45 | -0.43 | -0.48 | -0.5 | -0.47 | -0.44 | -0.5 | -0.48 | -0.36 | -0.43 | -0.46 | -0.51 | 46.33 | -0.41 | -0.43 | -0.45 | -0.44 | 43.35 | 48.14 |
| Run1_Nl-3 | | -0.36 | -0.51 | -0.66 | -0.43 | -0.49 | -0.48 | -0.43 | -0.38 | -0.37 | -0.47 | -0.46 | -0.44 | -0.42 | -0.45 | -0.49 | -0.39 | -0.27 | -0.46 | -0.33 | 38.82 | -0.42 | -0.49 | -0.5 | -0.47 | 33.67 | 42.8 |
| Run1_Nl-4 | | -0.39 | -0.45 | -0.35 | -0.59 | -0.46 | -0.46 | -0.44 | -0.41 | -0.41 | -0.49 | -0.48 | -0.45 | -0.44 | -0.51 | -0.51 | -0.41 | -0.33 | -0.46 | -0.54 | 35.34 | -0.44 | -0.49 | -0.51 | -0.43 | 33.54 | 36.94 |
| Run1_Nl-6 | | -0.4 | -0.46 | -0.49 | -0.46 | -0.51 | -0.46 | -0.39 | -0.39 | -0.41 | -0.45 | -0.46 | -0.43 | -0.41 | -0.46 | -0.48 | -0.4 | -0.41 | -0.44 | -0.48 | 34.97 | -0.41 | -0.43 | -0.49 | -0.35 | 31.02 | 36.74 |
| Run2_Nl-3 | Normal | -0.36 | -0.41 | -0.46 | -0.41 | -0.44 | -0.43 | -0.41 | -0.37 | -0.38 | -0.43 | -0.43 | -0.41 | -0.39 | -0.45 | -0.45 | -0.4 | -0.27 | -0.42 | -0.46 | 31.31 | -0.41 | -0.45 | -0.43 | -0.44 | 26.76 | 34.69 |
| Run2_Nl-4 | | -0.35 | -0.41 | -0.46 | -0.3 | -0.41 | -0.42 | -0.39 | -0.38 | -0.37 | -0.42 | -0.43 | -0.4 | -0.39 | -0.42 | -0.43 | -0.4 | -0.31 | -0.43 | -0.44 | 28.14 | -0.38 | -0.41 | -0.44 | -0.4 | 26.68 | 29.57 |
| Run2_Nl-5 | | -0.35 | -0.43 | -0.47 | -0.35 | -0.44 | -0.42 | -0.39 | -0.39 | -0.37 | -0.43 | -0.42 | -0.41 | -0.4 | -0.42 | -0.44 | -0.4 | -0.39 | -0.42 | -0.41 | 28.76 | -0.38 | -0.45 | -0.47 | -0.43 | 25.75 | 31.82 |
| Run2_Nl-6 | | -0.38 | -0.4 | -0.34 | -0.43 | -0.42 | -0.42 | -0.41 | -0.4 | -0.38 | -0.41 | -0.42 | -0.4 | -0.39 | -0.41 | -0.42 | -0.41 | -0.39 | -0.42 | -0.5 | 25.88 | -0.38 | -0.39 | -0.45 | -0.47 | 23.07 | 27.02 |

**Supplemental Table 3**: Median expression of each surface marker on each cell population.

### CD45 Median

| | | Ungated | CD34+CD38low | HSCs | MPP | CMP/GMP | Myelo/Mono-Blasts | ProMonocytes | CD14neg Monocytes | Mature Monocytes | ProMyelocytes | Myelocytes | MetaMyelocytes | MatureGrans | ProErythroblasts | Erythroblasts | LateErythroblasts | PreBcells | Mature Bcells | Plasma Cells | T cells | NK cells | pDCs | Basophils | Platelets | CD8+ T cells | CD8neg T cells |
|---|---|---|---|---|---|---|---|---|---|---|---|---|---|---|---|---|---|---|---|---|---|---|---|---|---|---|---|
| Run1_MDS17 | | 10.44 | 2.62 | 2.79 | 2.45 | 3.21 | 2.44 | 28.28 | 92.13 | 108.63 | 6.59 | 10.7 | 40.57 | 56.4 | -0.19 | -0.46 | -0.07 | 24.94 | 81.86 | 8.33 | 138.42 | 139.49 | 3.05 | 27.17 | 3.3 | 154.11 | 119.71 |
| Run2_MDS15* | AML/RAEB-T | 26.54 | 6.27 | 3.34 | 3.28 | 5.2 | 6.16 | 12.87 | 64.92 | 82.56 | 5.5 | 10.92 | 31.54 | 54.08 | 11.36 | 12.44 | 46.32 | 55.12 | 86.8 | 29.93 | 137.12 | 132.2 | 13.16 | 19.14 | 4.18 | 165.64 | 114.11 |
| Run2_MDS4 | | 31.47 | 5.98 | 8.54 | 4.77 | 1.86 | 6.56 | 28.59 | 47.59 | 54.98 | 4.63 | 8.87 | 32.31 | 47.21 | -0.23 | -0.42 | -0.13 | 11.38 | 10.25 | 1.59 | 85.19 | 75.24 | 10.73 | 19.69 | 3.44 | 104.97 | 71.23 |
| Run1_MDS21 | | 9.27 | 2.21 | 2.19 | 2.03 | 2.25 | 2.26 | 11.33 | 91.7 | 101.34 | 3.13 | 10.27 | 53.48 | 78.91 | -0.24 | -0.04 | 10.33 | 58.87 | 79.07 | 1.14 | 90.08 | 106.15 | 11.79 | 27.6 | 3.29 | 122.4 | 81.73 |
| Run1_MDS3 | | 9.57 | 3.15 | 2.97 | 2.77 | 3.44 | 3.64 | 11.91 | 83.57 | 112.65 | 4.16 | 10.64 | 47.46 | 68.93 | -0.12 | 0 | 6.78 | 68.56 | 92.05 | 3.24 | 117.48 | 137.09 | 89.34 | 27.26 | 2.53 | 143.57 | 108.69 |
| Run2_MDS13 | | 10.34 | 1.74 | 1.73 | 1.53 | 1.94 | 2.23 | 32.45 | 67.05 | 69.25 | 5.28 | 9.51 | 31.05 | 39.44 | 0.69 | -0.23 | -0.06 | 41.66 | 54.28 | 20.36 | 87.1 | 87.81 | 24.73 | 15.98 | 2.05 | 111.42 | 78.01 |
| Run2_MDS16 | | 29.58 | 12.52 | 12.72 | 11.24 | 8.03 | 32.74 | 78.09 | 70.7 | 85.64 | 4.13 | 7.96 | 19.67 | 29.62 | 6.55 | -0.28 | 17.04 | 69.82 | 56.22 | 4.27 | 77.37 | 122.85 | 18.03 | 20.4 | 0.7 | 83.5 | 76.08 |
| Run2_MDS1 | | 32.84 | 4.52 | 6.56 | 2.69 | 3.55 | 5.86 | 22.83 | 51.94 | 72.13 | 2.51 | 7.09 | 28.8 | 37.99 | 0.53 | -0.26 | -0.19 | 55.39 | 48.1 | 3.22 | 66.03 | 91.44 | 15.09 | 11.89 | 1.16 | 86.79 | 62.79 |
| Run2_MDS21 | Higher Risk | 6.28 | 0.83 | 1.06 | 0.78 | 0.94 | 0.88 | 14.62 | 55.58 | 59.19 | 3.48 | 5.9 | 24.08 | 33.1 | -0.28 | -0.39 | -0.25 | 39.19 | 54.48 | 1.07 | 68.51 | 70.07 | 2.28 | 14.78 | 1.42 | 94 | 61.62 |
| Run2_MDS26* | | 11.88 | 5.98 | 6.35 | 5.7 | 4.12 | 1.68 | 12.22 | 74.42 | 85.35 | 2.01 | 4.84 | 21.93 | 33.18 | 1.24 | -0.4 | -0.39 | 12.22 | 36.93 | 0.23 | 79.7 | 87.14 | 19.54 | 19.33 | 2.18 | 99.23 | 70.9 |
| Run2_MDS27 | | 23.81 | 11.09 | 13.2 | 9.17 | 7.68 | 8.44 | 37.88 | 57.5 | 85.41 | 3.79 | 6.02 | 20.51 | 36.6 | 6.25 | -0.28 | -0.26 | 74.84 | 54.51 | 3.59 | 98.6 | 106.87 | 129.4 | 22.43 | 2.25 | 106.97 | 89.87 |
| Run2_MDS2 | | 30.33 | 0.45 | 0.73 | 0.27 | 0.44 | 1.1 | 26.64 | 40.96 | 61.16 | 2.38 | 6.65 | 29.31 | 34.94 | 32.39 | 23.4 | 3.17 | 34.48 | 86.21 | 3.64 | 48.96 | 63.79 | 24.8 | 7.23 | -0.22 | 71.67 | 47.92 |
| Run2_MDS3 | | 9.23 | 1.51 | 1.54 | 1.35 | 1.61 | 1.94 | 21.96 | 57.46 | 62.47 | 6.01 | 9.16 | 24.66 | 34.56 | 0.18 | -0.29 | -0.27 | 49.57 | 66.62 | 2.89 | 90.64 | 99.58 | 11.81 | 15.95 | 1.87 | 114.14 | 82.91 |
| Run1_MDS23 | | 41.94 | 11.02 | 12.6 | 9.61 | 7.89 | 13.58 | 62.89 | 130.62 | 169.73 | 4.54 | 8.79 | 36.71 | 64.14 | 0.38 | 0.32 | 43.35 | 87.52 | 67.09 | 1.28 | 122.84 | 176.88 | 27.65 | 37.12 | 2.58 | 177.91 | 109.02 |
| Run1_MDS25 | | 69.42 | 51.49 | 4.68 | 56.9 | 6.25 | 63.98 | 74.3 | 101.83 | 118.93 | 6.46 | 14.63 | 52.19 | 68.31 | 1.08 | 2.16 | 79.63 | 104.49 | 92.13 | 10.14 | 128.59 | 141.6 | 34.11 | 35.58 | 1.52 | 148.31 | 113.19 |
| Run1_MDS5 | | 110.26 | 42.09 | 24.82 | 44.67 | 21.25 | 26.67 | 101.66 | 132.04 | 156.91 | 16.51 | 54.08 | 96.65 | 117.62 | 52.7 | 0.92 | 17.87 | 126.03 | 84.03 | 23.07 | 148.58 | 193.7 | 62.19 | 36.26 | 1.53 | 203.43 | 133.87 |
| Run1_MDS6 | | 49.93 | 14.27 | 12.05 | 11.49 | 11.78 | 16.47 | 78.86 | 105.28 | 136.31 | 7.08 | 13.57 | 34.56 | 53.45 | 6.1 | -0.16 | 35.51 | 93.96 | 112.73 | 10.91 | 137.21 | 175.5 | 49 | 43.79 | 3.96 | 158.89 | 123.54 |
| Run1_MDS8 | | 61.94 | 20.38 | 16.81 | 19.58 | 10.56 | 22.68 | 200.77 | 170.22 | 174.34 | 7.02 | 15.5 | 50.07 | 63.75 | 1.2 | 0.39 | 4.25 | 83.17 | 82.7 | 7.89 | 82.2 | 124.46 | 36.81 | 38.92 | 1.54 | 92.56 | 79.52 |
| Run1_MDS9 | | 57.61 | 11.81 | 14.71 | 9.25 | 12.19 | 14.69 | 172.61 | 150.44 | 139.07 | 5.94 | 12.55 | 42.02 | 67.27 | 1.3 | 0.53 | 23.76 | 95.23 | 79.27 | 5.28 | 97.31 | 152.59 | 134.29 | 34.88 | 1.88 | 132.31 | 86 |
| Run2_MDS12 | Lower Risk | 36 | 9.61 | 9.67 | 8.88 | 9 | 24.72 | 42.32 | 69.42 | 95.43 | 3.52 | 7.99 | 36.51 | 57.81 | -0.15 | -0.44 | 2.3 | 48.47 | 80.16 | 1.21 | 82.87 | 121.28 | 23.84 | 29.21 | 2.42 | 120.53 | 67.83 |
| Run2_MDS14 | | 28.14 | 12.1 | 0.17 | 35.33 | 4.69 | 48.4 | 139.61 | 74.39 | 103.8 | 3.46 | 5.83 | 14.57 | 28.46 | 0.5 | -0.32 | 27.1 | 17.62 | 59.05 | 1.08 | 94.55 | 126.74 | 16.3 | 15.06 | 2.62 | 107.05 | 89.37 |
| Run2_MDS19 | | 25.82 | 12.47 | 14.9 | 11.86 | 6.64 | 21.52 | 48.21 | 62.31 | 87.09 | 3.58 | 7.7 | 21.8 | 32.45 | -0.05 | -0.39 | -0.18 | 76.14 | 69.99 | 0.49 | 97.77 | 101.95 | 18.54 | 19.34 | 1.28 | 122.92 | 91.74 |
| Run2_MDS20 | | 51.48 | 15.59 | 17.71 | 17.68 | 8.16 | 81.17 | 133.51 | 115.52 | 110.88 | 4.47 | 7.78 | 24.3 | 38.98 | 3.41 | 0.34 | 36.22 | 104.98 | 84.51 | 8.16 | 112.37 | 108.33 | 101.12 | 23.51 | 1.61 | 130.84 | 110.49 |
| Run2_MDS28* | | 46.51 | 6.51 | 11.8 | 5.55 | 6.06 | 13.19 | 35.94 | 84.53 | 128.95 | 4.17 | 6.73 | 30.3 | 59.55 | 1.69 | -0.27 | 38.36 | 92.15 | 59.16 | -0.24 | 102.29 | 127.16 | 82.24 | 24.22 | 2.15 | 123.55 | 87.64 |
| Run2_MDS5 | | 78.43 | 47.41 | 23.74 | 22.7 | 15.21 | 27.62 | 90.43 | 112.04 | 109.46 | 13.66 | 41.21 | 67.46 | 80.43 | 22.54 | 0.1 | 66.79 | 97.93 | 68.16 | 19.8 | 124.31 | 158.91 | 44.96 | 29.34 | 1.37 | 171.38 | 112.38 |
| Run1_ICUS18 | | 63.94 | 20.66 | 23.19 | 15.04 | 11.56 | 79.14 | 118.26 | 129.29 | 142.58 | 5.1 | 9.73 | 34.04 | 59.71 | 0.94 | 0.29 | 70.8 | 108.42 | 78.52 | 0.94 | 107.91 | 161.31 | 44.33 | 32.15 | 2.6 | 133.91 | 103.77 |
| Run1_ICUS22 | ICUS | 74.42 | 17.04 | 21.54 | 15.78 | 10.34 | 45.14 | 143.7 | 150.25 | 178.21 | 5.76 | 10.13 | 38.9 | 77.62 | 0.97 | 0.8 | 81.31 | 39.43 | 77.52 | 1.06 | 121.8 | 137.77 | 172.65 | 29.25 | 2.6 | 155.35 | 112.02 |
| Run1_ICUS24 | | 55.1 | 15.99 | 21.05 | 14.07 | 10.12 | 21.51 | 56.81 | 102.93 | 153.42 | 6.26 | 10.72 | 35.17 | 65.25 | 0.56 | -0.2 | 16.29 | 22.22 | 77.22 | 0.43 | 114.2 | 136.91 | 23.66 | 24.98 | 6.4 | 123.65 | 108.2 |
| Run1_Nl-1 | | 66.44 | 18.95 | 19.41 | 17.18 | 10.86 | 26.62 | 109.28 | 116.03 | 181.78 | 8.05 | 13.02 | 38.76 | 77.91 | 0.78 | 0.28 | 67.9 | 21.09 | 69.05 | 2.58 | 111.92 | 173.51 | 39.93 | 31.23 | 6.81 | 118.64 | 108.56 |
| Run1_Nl-3 | | 42.6 | 18.36 | 19.19 | 19.03 | 11.15 | 22.74 | 57.11 | 94.87 | 153.67 | 7.61 | 13.82 | 34.72 | 53.65 | 2.88 | 0.39 | 51.84 | 42.9 | 71.94 | 2.4 | 115.15 | 158.93 | 33.36 | 36.36 | 1.67 | 130.29 | 106.91 |
| Run1_Nl-4 | | 45.32 | 16.67 | 16.32 | 16.24 | 10.98 | 20.87 | 50.53 | 93.45 | 144.99 | 7.74 | 14.49 | 39.52 | 56.62 | 4.4 | 0.65 | 54.26 | 47.21 | 84.37 | 6.68 | 125.76 | 154.39 | 30.08 | 30.78 | 2.37 | 132.42 | 121.5 |
| Run1_Nl-6 | | 40.01 | 17.31 | 19.48 | 15.19 | 8.8 | 15.61 | 51.54 | 94.2 | 137.68 | 6.12 | 11.95 | 34.44 | 61.29 | 1.2 | -0.13 | 46.78 | 21.16 | 59.58 | 3.37 | 113.24 | 159.99 | 32.54 | 26.8 | 1.97 | 123.38 | 110.06 |
| Run2_Nl-3 | Normal | 27.69 | 13.82 | 16.04 | 12.28 | 7.94 | 17.99 | 46.37 | 80.62 | 115.98 | 5.89 | 10.61 | 25.91 | 35.85 | 1.01 | 0.1 | 32.03 | 30.6 | 58.92 | 1.6 | 97.11 | 130.8 | 25.64 | 25.33 | 1.24 | 110.15 | 90.6 |
| Run2_Nl-4 | | 27.24 | 13.67 | 16.16 | 14.51 | 7.72 | 16.46 | 39.89 | 72.38 | 94.69 | 6.04 | 10.48 | 25.92 | 33.91 | 2.11 | 0.16 | 30.97 | 32.1 | 68.05 | 6.04 | 108.47 | 122.84 | 21.35 | 21.41 | 1.78 | 113.92 | 104.66 |
| Run2_Nl-5 | | 32.14 | 12.45 | 13.17 | 11.13 | 6.93 | 17.36 | 44.18 | 88.47 | 110.09 | 4.87 | 9.42 | 27.83 | 39.93 | 2.55 | -0.11 | 35.05 | 16.28 | 58.7 | 1.78 | 103.59 | 149.72 | 23.64 | 23.4 | 1.34 | 106.18 | 100.81 |
| Run2_Nl-6 | | 24.49 | 11.57 | 13.55 | 11.45 | 6.1 | 11.22 | 39.22 | 71.58 | 92.94 | 4.12 | 7.89 | 22.91 | 34.18 | 0.33 | -0.29 | 17.99 | 14.57 | 45.61 | 2.89 | 94.18 | 129.14 | 22.21 | 18.19 | 2.05 | 104.52 | 90.85 |

**Supplemental Table 3**: Median expression of each surface marker on each cell population.

### CD133 Median

| | | Ungated | CD34+CD38low | HSCs | MPP | CMP/GMP | Myelo/Mono-Blasts | ProMonocytes | CD14neg Monocytes | Mature Monocytes | ProMyelocytes | Myelocytes | MetaMyelocytes | MatureGrans | ProErythroblasts | Erythroblasts | LateErythroblasts | PreBcells | Mature Bcells | Plasma Cells | T cells | NK cells | pDCs | Basophils | Platelets | CD8+ T cells | CD8neg T cells |
|---|---|---|---|---|---|---|---|---|---|---|---|---|---|---|---|---|---|---|---|---|---|---|---|---|---|---|---|
| Run1_MDS17 | | -0.36 | -0.22 | -0.2 | -0.24 | -0.07 | -0.35 | -0.41 | -0.43 | -0.42 | -0.43 | -0.37 | -0.37 | -0.33 | -0.3 | -0.33 | -0.37 | -0.32 | -0.34 | -0.37 | -0.45 | -0.41 | 1.12 | -0.6 | -0.42 | -0.45 | -0.44 |
| Run2_MDS15* | AML/RAEB-T | -0.23 | 1.47 | 0.93 | 0.8 | -0.07 | -0.26 | -0.33 | -0.32 | -0.33 | -0.33 | -0.33 | -0.24 | -0.15 | 0.13 | -0.06 | -0.06 | -0.28 | -0.29 | 0.01 | -0.36 | -0.26 | -0.2 | -0.38 | -0.05 | -0.35 | -0.37 |
| Run2_MDS4 | | -0.33 | 0.13 | 0.3 | -0.08 | -0.47 | -0.33 | -0.32 | -0.32 | -0.33 | -0.38 | -0.36 | -0.34 | -0.32 | -0.27 | -0.27 | -0.33 | -0.33 | -0.29 | -0.41 | -0.39 | -0.25 | -0.27 | -0.35 | -0.33 | -0.39 | -0.4 |
| Run1_MDS21 | | -0.37 | -0.24 | -0.21 | -0.27 | -0.21 | -0.25 | -0.33 | -0.39 | -0.39 | -0.44 | -0.41 | -0.4 | -0.35 | -0.27 | -0.13 | -0.05 | -0.42 | -0.36 | -0.24 | -0.43 | -0.33 | -0.02 | -0.39 | -0.23 | -0.39 | -0.45 |
| Run1_MDS3 | | -0.39 | -0.24 | -0.3 | -0.24 | -0.15 | -0.22 | -0.32 | -0.45 | -0.38 | -0.45 | -0.39 | -0.39 | -0.31 | -0.35 | -0.28 | -0.24 | -0.43 | -0.37 | -0.27 | -0.44 | -0.28 | 0.08 | -0.44 | -0.35 | -0.42 | -0.45 |
| Run2_MDS13 | | -0.34 | -0.19 | -0.2 | -0.21 | -0.18 | -0.24 | -0.34 | -0.34 | -0.31 | -0.37 | -0.36 | -0.33 | -0.33 | -0.26 | -0.27 | -0.3 | -0.4 | -0.36 | -0.32 | -0.38 | -0.33 | -0.28 | -0.37 | -0.34 | -0.37 | -0.39 |
| Run2_MDS16 | | -0.3 | 0.91 | 1.06 | 0.78 | 0.59 | -0.29 | -0.3 | -0.37 | -0.37 | -0.35 | -0.36 | -0.33 | -0.31 | 0.19 | -0.05 | -0.27 | -0.34 | -0.38 | -0.31 | -0.37 | -0.29 | -0.29 | -0.37 | -0.29 | -0.36 | -0.37 |
| Run2_MDS1 | | -0.38 | -0.2 | -0.07 | -0.23 | -0.21 | -0.36 | -0.38 | -0.36 | -0.37 | -0.36 | -0.42 | -0.35 | -0.34 | -0.36 | -0.24 | -0.32 | -0.35 | -0.35 | -0.26 | -0.4 | -0.35 | -0.36 | -0.31 | -0.31 | -0.39 | -0.4 |
| Run2_MDS21 | Higher Risk | -0.34 | -0.24 | -0.24 | -0.24 | -0.2 | -0.25 | -0.35 | -0.32 | -0.35 | -0.37 | -0.34 | -0.35 | -0.32 | -0.31 | -0.3 | -0.32 | -0.37 | -0.31 | -0.14 | -0.39 | -0.35 | 0.77 | -0.45 | -0.26 | -0.38 | -0.39 |
| Run2_MDS26* | | -0.4 | -0.19 | -0.18 | -0.2 | -0.28 | -0.36 | -0.22 | -0.38 | -0.48 | -0.41 | -0.41 | -0.4 | -0.39 | -0.29 | -0.31 | -0.37 | -0.38 | -0.36 | -0.51 | -0.39 | -0.34 | -0.24 | -0.42 | -0.48 | -0.38 | -0.39 |
| Run2_MDS27 | | -0.21 | 1.14 | 1.33 | 0.92 | 0.43 | -0.16 | -0.3 | -0.37 | -0.36 | -0.36 | -0.32 | -0.31 | -0.25 | -0.14 | -0.01 | -0.34 | -0.32 | -0.37 | -0.29 | -0.38 | -0.32 | -0.39 | -0.44 | -0.1 | -0.37 | -0.38 |
| Run2_MDS2 | | -0.36 | -0.28 | -0.28 | -0.32 | -0.24 | -0.31 | -0.28 | -0.37 | -0.3 | -0.3 | -0.36 | -0.36 | -0.34 | -0.12 | -0.12 | -0.35 | -0.48 | -0.46 | -0.34 | -0.4 | -0.37 | -0.47 | -0.43 | -0.39 | -0.33 | -0.41 |
| Run2_MDS3 | | -0.34 | -0.21 | -0.24 | -0.21 | -0.16 | -0.22 | -0.29 | -0.36 | -0.65 | -0.36 | -0.29 | -0.29 | -0.31 | -0.27 | -0.32 | -0.36 | -0.36 | -0.37 | -0.28 | -0.39 | -0.26 | 0.14 | -0.42 | -0.31 | -0.37 | -0.4 |
| Run1_MDS23 | | -0.38 | 0.31 | 0.55 | 0.24 | 0.38 | -0.24 | -0.3 | -0.31 | -0.32 | -0.38 | -0.41 | -0.39 | -0.38 | -0.27 | -0.17 | -0.24 | -0.37 | -0.38 | -0.46 | -0.46 | -0.36 | -0.24 | -0.5 | -0.32 | -0.43 | -0.47 |
| Run1_MDS25 | | -0.36 | 0.56 | 3.03 | -0.68 | -0.21 | -0.24 | -0.42 | -0.41 | -0.43 | -0.44 | -0.44 | -0.38 | -0.35 | -0.08 | -0.07 | -0.05 | -0.37 | -0.23 | -0.4 | -0.42 | -0.35 | -0.32 | -0.39 | -0.37 | -0.42 | -0.42 |
| Run1_MDS5 | | -0.31 | 1.93 | 2.44 | 0.62 | 0.43 | 0 | -0.29 | -0.25 | -0.2 | -0.35 | -0.36 | -0.33 | -0.3 | -0.12 | 0.3 | -0.21 | -0.29 | -0.35 | -0.26 | -0.38 | -0.32 | -0.07 | -0.36 | -0.38 | -0.35 | -0.39 |
| Run1_MDS6 | | -0.3 | 1.63 | 1.07 | 0.17 | 0.7 | -0.06 | -0.33 | -0.32 | -0.34 | -0.37 | -0.38 | -0.32 | -0.29 | 0.27 | 0.23 | -0.12 | -0.42 | -0.34 | -0.21 | -0.42 | -0.26 | -0.3 | -0.4 | -0.17 | -0.39 | -0.44 |
| Run1_MDS8 | | -0.25 | 1.16 | 1.28 | 1.07 | 0.52 | -0.07 | -0.25 | -0.38 | -0.38 | -0.4 | -0.36 | -0.22 | -0.19 | 0.21 | 0.14 | -0.16 | -0.38 | -0.41 | -0.3 | -0.44 | -0.3 | -0.02 | -0.44 | -0.26 | -0.44 | -0.44 |
| Run1_MDS9 | | -0.18 | 1.97 | 3.19 | 1.04 | 1.9 | 0.4 | -0.26 | -0.37 | -0.34 | -0.38 | -0.31 | -0.08 | 0.06 | 0.19 | 0.28 | 0.23 | -0.35 | -0.38 | -0.4 | -0.44 | -0.28 | -0.29 | -0.41 | -0.33 | -0.43 | -0.45 |
| Run2_MDS12 | Lower Risk | -0.34 | 0.2 | 0.19 | 0 | 0.16 | -0.25 | -0.34 | -0.35 | -0.35 | -0.39 | -0.37 | -0.34 | -0.31 | -0.19 | -0.24 | -0.32 | -0.35 | -0.34 | -0.27 | -0.41 | -0.29 | -0.25 | -0.38 | -0.24 | -0.38 | -0.42 |
| Run2_MDS14 | | -0.3 | -0.16 | -0.36 | -0.41 | -0.45 | -0.36 | -0.36 | -0.34 | -0.32 | -0.33 | -0.35 | -0.32 | -0.3 | -0.17 | 0.03 | -0.21 | -0.39 | -0.35 | -0.25 | -0.4 | -0.25 | -0.37 | -0.39 | -0.16 | -0.39 | -0.4 |
| Run2_MDS19 | | -0.31 | 0.41 | 0.48 | 0.48 | 0.44 | -0.18 | -0.31 | -0.37 | -0.36 | -0.37 | -0.34 | -0.3 | -0.27 | -0.18 | -0.26 | -0.35 | -0.37 | -0.37 | -0.34 | -0.38 | -0.32 | -0.2 | -0.41 | -0.13 | -0.37 | -0.38 |
| Run2_MDS20 | | -0.29 | 0.49 | 0.49 | 0.97 | 0.59 | -0.26 | -0.28 | -0.31 | -0.31 | -0.33 | -0.29 | -0.25 | -0.23 | -0.11 | -0.11 | -0.16 | -0.37 | -0.29 | -0.12 | -0.38 | -0.35 | -0.34 | -0.38 | -0.3 | -0.37 | -0.39 |
| Run2_MDS28* | | -0.25 | 0.45 | 0.75 | 0.57 | 0.46 | -0.06 | -0.23 | -0.27 | -0.24 | -0.27 | -0.28 | -0.23 | -0.23 | -0.22 | -0.25 | -0.2 | -0.29 | -0.28 | -0.29 | -0.39 | -0.31 | -0.28 | -0.39 | -0.14 | -0.38 | -0.4 |
| Run2_MDS5 | | -0.29 | 1.5 | 1.5 | 1.81 | 1.69 | -0.03 | -0.41 | -0.3 | -0.24 | -0.33 | -0.33 | -0.31 | -0.29 | -0.09 | -0.02 | -0.17 | -0.27 | -0.35 | -0.21 | -0.35 | -0.29 | -0.07 | -0.38 | -0.23 | -0.35 | -0.35 |
| Run1_ICUS18 | | -0.3 | 1.2 | 2.78 | 0.67 | 1 | -0.27 | -0.24 | -0.35 | -0.37 | -0.37 | -0.36 | -0.31 | -0.27 | -0.2 | -0.29 | -0.16 | -0.38 | -0.39 | -0.29 | -0.43 | -0.32 | -0.37 | -0.46 | -0.39 | -0.41 | -0.44 |
| Run1_ICUS22 | ICUS | -0.34 | 0.77 | 0.98 | 0.44 | 0.34 | -0.19 | -0.26 | -0.32 | -0.35 | -0.4 | -0.39 | -0.35 | -0.32 | -0.08 | -0.11 | -0.22 | -0.36 | -0.34 | -0.49 | -0.43 | -0.37 | -0.28 | -0.43 | -0.37 | -0.43 | -0.43 |
| Run1_ICUS24 | | -0.32 | 0.78 | 1.16 | 0.7 | 0.19 | -0.23 | -0.36 | -0.41 | -0.4 | -0.38 | -0.38 | -0.3 | -0.26 | -0.06 | -0.14 | -0.03 | -0.38 | -0.33 | -0.49 | -0.46 | -0.35 | -0.29 | -0.45 | -0.13 | -0.46 | -0.45 |
| Run1_Nl-1 | | -0.31 | 0.9 | 1.14 | 0.56 | 0 | -0.28 | -0.36 | -0.38 | -0.39 | -0.4 | -0.38 | -0.31 | -0.25 | -0.15 | -0.17 | -0.06 | -0.44 | -0.39 | -0.55 | -0.44 | -0.31 | -0.32 | -0.39 | -0.02 | -0.44 | -0.43 |
| Run1_Nl-3 | | -0.28 | 0.4 | 0.41 | 0.14 | -0.04 | -0.25 | -0.32 | -0.34 | -0.36 | -0.36 | -0.34 | -0.24 | -0.19 | -0.06 | -0.23 | -0.09 | -0.4 | -0.4 | -0.47 | -0.44 | -0.35 | -0.34 | -0.42 | -0.43 | -0.43 | -0.44 |
| Run1_Nl-4 | | -0.28 | 0.52 | 1.22 | 0.09 | -0.01 | -0.22 | -0.31 | -0.29 | -0.34 | -0.35 | -0.34 | -0.26 | -0.23 | 0.03 | -0.2 | -0.12 | -0.37 | -0.36 | -0.33 | -0.44 | -0.3 | -0.3 | -0.47 | -0.4 | -0.44 | -0.45 |
| Run1_Nl-6 | | -0.31 | 0.65 | 0.92 | 0.31 | -0.01 | -0.21 | -0.31 | -0.26 | -0.34 | -0.37 | -0.37 | -0.3 | -0.28 | -0.11 | -0.22 | -0.22 | -0.39 | -0.36 | -0.34 | -0.43 | -0.33 | -0.29 | -0.41 | -0.46 | -0.44 | -0.43 |
| Run2_Nl-3 | Normal | -0.26 | 0.23 | 0.31 | 0.3 | -0.11 | -0.25 | -0.31 | -0.3 | -0.32 | -0.32 | -0.3 | -0.22 | -0.18 | -0.21 | -0.25 | -0.17 | -0.38 | -0.38 | -0.49 | -0.4 | -0.33 | -0.3 | -0.41 | -0.34 | -0.4 | -0.4 |
| Run2_Nl-4 | | -0.25 | 0.45 | 1.06 | 0.07 | -0.01 | -0.19 | -0.3 | -0.29 | -0.32 | -0.3 | -0.29 | -0.21 | -0.19 | -0.14 | -0.24 | -0.17 | -0.34 | -0.34 | -0.25 | -0.4 | -0.29 | -0.35 | -0.44 | -0.37 | -0.39 | -0.4 |
| Run2_Nl-5 | | -0.26 | 0.5 | 0.63 | 0.47 | -0.01 | -0.25 | -0.28 | -0.27 | -0.3 | -0.32 | -0.29 | -0.23 | -0.18 | -0.11 | -0.19 | -0.21 | -0.38 | -0.39 | -0.33 | -0.4 | -0.29 | -0.36 | -0.39 | -0.36 | -0.4 | -0.4 |
| Run2_Nl-6 | | -0.29 | 0.44 | 0.58 | 0.24 | 0.09 | -0.2 | -0.28 | -0.28 | -0.31 | -0.34 | -0.33 | -0.27 | -0.25 | -0.22 | -0.29 | -0.28 | -0.36 | -0.36 | -0.3 | -0.4 | -0.31 | -0.34 | -0.4 | -0.34 | -0.39 | -0.4 |

**Supplemental Table 3**: Median expression of each surface marker on each cell population.

### HLADR Median

| Sample | Group | Ungated | CD34+CD38low | HSCs | MPP | CMP/GMP | Myelo/Mono-Blasts | ProMonocytes | CD14neg Monocytes | Mature Monocytes | ProMyelocytes | Myelocytes | MetaMyelocytes | MatureGrans | ProErythroblasts | Erythroblasts | LateErythroblasts | PreBcells | Mature Bcells | Plasma Cells | T cells | NK cells | pDCs | Basophils | Platelets | CD8+ T cells | CD8neg T cells |
|---|---|---|---|---|---|---|---|---|---|---|---|---|---|---|---|---|---|---|---|---|---|---|---|---|---|---|---|
| Run1_MDS17 | | 0.81 | 20.39 | 23.83 | 18.93 | 36.12 | 18.89 | 20.15 | 19.04 | 25.01 | -0.23 | -0.14 | -0.11 | 0.03 | 4.71 | 1.8 | 0.5 | 2.68 | 354 | 8.84 | 0.5 | -0.11 | 54.37 | 0.58 | 0.62 | 0.89 | -0.03 |
| Run2_MDS15* | AML/RAEB-T | 5.54 | 650.24 | 163.92 | 156.34 | 91.62 | 82.86 | 67.15 | 62.2 | 161.9 | 2.91 | 0.68 | 0.56 | 0.51 | 889.03 | 5.29 | 9.51 | 23.36 | 565.57 | 257.71 | 0.4 | 0.88 | 126.79 | 1.08 | 17.77 | 1.32 | -0.09 |
| Run2_MDS4 | | 1.54 | 752.93 | 632.84 | 442.64 | 510.02 | 491.92 | 560.78 | 341.38 | 208.91 | 0.78 | 0.62 | 0.43 | 0.42 | 2.78 | 0.8 | 2.48 | 261.69 | 494.8 | 3.21 | 1.05 | 1.82 | 271.97 | 0.98 | 5.97 | 1.93 | 0.49 |
| Run1_MDS21 | | 0.72 | 118.14 | 124.88 | 115.8 | 168.56 | 136 | 40.91 | 15 | 16.51 | -0.23 | -0.18 | -0.13 | 0.08 | 2.67 | 3.82 | 1.02 | 89.91 | 333.43 | 3.81 | -0.23 | 0.83 | 124.92 | 1.22 | 0.15 | 0.36 | -0.35 |
| Run1_MDS3 | | 0.57 | 62.64 | 82.62 | 72.96 | 114.26 | 82.18 | 37.18 | 13.1 | 25.78 | -0.21 | -0.17 | -0.09 | 0.39 | 4.86 | 4.76 | 0.53 | 108.03 | 378.47 | 11.26 | -0.23 | 1.04 | 216.16 | 1.44 | 0.16 | 0.33 | -0.33 |
| Run2_MDS13 | | 0.82 | 346.51 | 341.79 | 319.39 | 386.9 | 336.54 | 67.76 | 43.91 | 64.28 | 0.23 | -0.01 | -0.02 | 0.09 | 9.36 | 3.29 | 0.69 | 74.21 | 295.2 | 79.8 | -0.15 | 0.7 | 266.05 | 0.58 | 0.09 | 0.79 | -0.28 |
| Run2_MDS16 | | -0.01 | 190.04 | 172.89 | 177.8 | 164.21 | 221.76 | 112.46 | 39.92 | 48.51 | 0.06 | -0.1 | -0.03 | 0.01 | 138.11 | 0.7 | 0.13 | 0.03 | 216.37 | 3.84 | -0.33 | -0.16 | 167.35 | 0.34 | -0.12 | -0.27 | -0.35 |
| Run2_MDS1 | | 0.41 | 467.1 | 323.71 | 496.71 | 435.7 | 337.03 | 165.52 | 168.66 | 192.18 | -0.03 | -0.29 | -0.24 | -0.21 | 22.54 | 2.18 | -0.04 | 4.73 | 266.15 | 13.1 | -0.12 | 0.11 | 119.55 | 1.89 | 0.41 | 0.86 | -0.18 |
| Run2_MDS21 | Higher Risk | 0.83 | 231.74 | 244.26 | 218.77 | 343.36 | 286.14 | 204.65 | 86.55 | 82.39 | 0.04 | -0.02 | -0.06 | 0.03 | 3.8 | 2.34 | 0.46 | 91.52 | 307.58 | 16.33 | -0.2 | -0.04 | 301.7 | 1.21 | 0.47 | 0.42 | -0.3 |
| Run2_MDS26* | | -0.24 | 170.17 | 176.81 | 159.57 | 254.34 | 54.87 | 38.03 | 41.23 | 39.88 | -0.29 | -0.31 | -0.27 | -0.23 | 6.1 | 0 | -0.23 | 0.37 | 240.16 | 0.5 | -0.22 | -0.34 | 111.01 | 0.47 | -0.26 | 0.1 | -0.33 |
| Run2_MDS27 | | 0.84 | 613.58 | 611.39 | 577.95 | 631.44 | 523.32 | 389.89 | 165.1 | 163.66 | 0.22 | 0 | 0.01 | 0.08 | 414.09 | 2.27 | -0.25 | 2.94 | 433.27 | 12.78 | 0.1 | -0.1 | 308.58 | 1.36 | 19.87 | 0.53 | -0.12 |
| Run2_MDS2 | | 0.93 | 798.8 | 726.34 | 799.34 | 871.35 | 703.58 | 636.6 | 256.89 | 315.74 | 47.14 | 0.57 | 0.27 | 1.01 | 65.3 | 3.99 | 1.62 | 3.12 | 564.94 | 187.7 | -0.14 | -0.3 | 171.15 | 0.84 | 1.95 | -0.03 | -0.14 |
| Run2_MDS3 | | 0.99 | 274.99 | 303.18 | 236.7 | 414.98 | 347.71 | 80.93 | 53.78 | 67.92 | 0.49 | 0.35 | 0.21 | 0.38 | 15.07 | 5.4 | 1.11 | 108.04 | 352.75 | 10.69 | -0.2 | 0.69 | 412.26 | 1.43 | 0.67 | 0.33 | -0.28 |
| Run1_MDS23 | | 0.16 | 323.95 | 317.36 | 282.43 | 964.25 | 552.1 | 383.72 | 335.47 | 418.48 | -0.12 | -0.2 | -0.21 | -0.07 | 34.66 | 33.25 | 7.51 | 96.69 | 465.9 | 16.42 | -0.11 | 0.79 | 689.89 | 0.45 | 7 | 2.6 | -0.25 |
| Run1_MDS25 | | -0.04 | 24.6 | 522.17 | 15.42 | 753.97 | 664.23 | 201.74 | 64.3 | 58.47 | -0.16 | -0.2 | -0.22 | -0.17 | 27.41 | 26.1 | 1.45 | 39.59 | 842.55 | 39.48 | 1.23 | -0.09 | 832.6 | 1.27 | 0.02 | 4.54 | 0.09 |
| Run1_MDS5 | | 0.48 | 141.49 | 466.65 | 141.93 | 355.22 | 442.05 | 118.47 | 37.48 | 54.36 | -0.22 | -0.12 | 0.11 | 0.22 | 11.79 | 2.41 | 1.1 | 4.67 | 224.19 | 25.53 | 0.19 | 12.2 | 132.8 | 4.67 | 0.68 | 11.43 | -0.13 |
| Run1_MDS6 | | 0.45 | 629.95 | 417.67 | 347.76 | 759.88 | 786.81 | 453.9 | 163.24 | 146.89 | 0.19 | 0.08 | 0.23 | 0.28 | 149.04 | 3.25 | 1.2 | 82.48 | 345.45 | 19.83 | -0.16 | 0.71 | 190.18 | 1.94 | 4.81 | 0.03 | -0.27 |
| Run1_MDS8 | | -0.1 | 160.25 | 157.96 | 143.64 | 229.53 | 273.58 | 388.64 | 157.65 | 79 | -0.19 | -0.26 | -0.22 | -0.18 | 5.36 | 8.56 | 0.11 | 1.73 | 202.89 | 11.3 | -0.34 | -0.02 | 77.59 | 1.07 | 0.44 | -0.18 | -0.38 |
| Run1_MDS9 | | -0.1 | 366.92 | 410.66 | 277.14 | 488.79 | 410.18 | 440.85 | 225.16 | 92.6 | -0.2 | -0.24 | -0.23 | -0.17 | 7.01 | 8.52 | 0.87 | 0.64 | 341.92 | 22.2 | -0.33 | -0.27 | 233.93 | 1.06 | 0.44 | -0.25 | -0.38 |
| Run2_MDS12 | Lower Risk | 0.21 | 1813.28 | 617.91 | 1094.3 | 1198.7 | 969.08 | 183.39 | 80.74 | 85.32 | -0.14 | -0.16 | -0.04 | 0.15 | 1.8 | 0.43 | -0.03 | 75.18 | 560.34 | 9.2 | -0.07 | 0.13 | 653.95 | 1.02 | 4.06 | 0.89 | -0.23 |
| Run2_MDS14 | | 0.24 | 36.64 | 10.72 | 120.08 | 65.96 | 390.22 | 335.1 | 68.14 | 96.89 | -0.01 | -0.05 | 0.01 | 0.05 | 1.73 | 0.5 | 0.42 | 147.01 | 353.15 | 4.47 | -0.01 | 0.81 | 311.58 | 0.41 | -0.11 | 0.66 | -0.09 |
| Run2_MDS19 | | 0.02 | 195.21 | 195.37 | 172.02 | 518.43 | 879.93 | 355.86 | 133.67 | 147.24 | -0.02 | -0.07 | -0.05 | -0.04 | 6.23 | 0.8 | -0.17 | 6.63 | 357.73 | 11.49 | -0.2 | 0.44 | 769.79 | 0.48 | 0.08 | 0.38 | -0.28 |
| Run2_MDS20 | | 0.42 | 242.54 | 280.51 | 120.01 | 644.87 | 749.45 | 730.6 | 462.9 | 395.45 | 0.53 | 0.18 | 0.18 | 0.29 | 130.3 | 10.56 | 0.64 | 1.47 | 608.86 | 43.24 | -0.11 | 2.78 | 296.23 | 2.05 | 0.22 | 0.58 | -0.13 |
| Run2_MDS28* | | 0.21 | 276.31 | 231.42 | 272.09 | 356.91 | 1424.88 | 907.2 | 466.69 | 582.28 | 0.09 | -0.08 | -0.04 | -0.01 | 62.33 | 3.32 | 0.37 | 88.5 | 441.74 | 6.35 | -0.06 | 0.17 | 227.27 | 0.93 | 2.96 | 0.78 | -0.24 |
| Run2_MDS5 | | 0.5 | 250.53 | 530.07 | 251.24 | 457 | 529.95 | 145.59 | 61.8 | 107.67 | 0 | -0.02 | 0.2 | 0.33 | 113.16 | 1.37 | 1.01 | 2.27 | 186.32 | 38.54 | 0.31 | 10.07 | 161.71 | 4.61 | 0.83 | 10.63 | -0.11 |
| Run1_ICUS18 | | 0.2 | 139.56 | 133.33 | 133.29 | 199.48 | 470.17 | 440.84 | 225.11 | 160.19 | -0.06 | -0.05 | 0 | 0.11 | 11.6 | 12.72 | 0.61 | 3.61 | 321.28 | 30.08 | -0.21 | -0.17 | 606.15 | 0.47 | -0.03 | 0.42 | -0.27 |
| Run1_ICUS22 | ICUS | 0.26 | 226.79 | 244.37 | 219.13 | 363.82 | 788.44 | 592.45 | 369.14 | 363.13 | -0.06 | -0.12 | -0.1 | -0.02 | 37.17 | 35.7 | 0.64 | 313.3 | 506.86 | 21.21 | -0.06 | -0.03 | 442.86 | 1.73 | 0.52 | 1.9 | -0.2 |
| Run1_ICUS24 | | 0.21 | 149.91 | 101.91 | 165.37 | 444.27 | 523.61 | 123.96 | 52.32 | 86.56 | -0.15 | -0.18 | -0.14 | -0.08 | 7.5 | 5.82 | 0.6 | 362.34 | 611.04 | 4.89 | -0.05 | -0.26 | 571.25 | 0.69 | 0.44 | 0.48 | -0.16 |
| Run1_Nl-1 | | 0.09 | 89.24 | 86.41 | 85.02 | 205.75 | 317.74 | 192.05 | 50.42 | 53.52 | -0.14 | -0.14 | -0.04 | 0.03 | 3.79 | 3.73 | 0.47 | 197.64 | 267.07 | 7.32 | -0.35 | -0.39 | 194.08 | 1.09 | 0.09 | -0.31 | -0.36 |
| Run1_Nl-3 | | 0.34 | 99.23 | 89.09 | 78.19 | 201.63 | 350.51 | 62.9 | 31.57 | 49.78 | -0.05 | -0.04 | 0.16 | 0.25 | 17.42 | 16.34 | 0.64 | 119.84 | 225.87 | 16.69 | -0.24 | -0.27 | 307.67 | 0.91 | -0.01 | -0.06 | -0.34 |
| Run1_Nl-4 | | 0.33 | 244.68 | 328.05 | 160.43 | 386.65 | 440.9 | 82.31 | 37.88 | 80.42 | -0.06 | -0.03 | 0.11 | 0.19 | 31.6 | 19.95 | 0.55 | 170.7 | 437.44 | 28.73 | -0.19 | -0.14 | 503.53 | 1.7 | 0.74 | -0.14 | -0.23 |
| Run1_Nl-6 | | 0.32 | 107.74 | 105.15 | 101.18 | 217.76 | 307.47 | 77.12 | 44.11 | 67.16 | -0.07 | -0.03 | 0.04 | 0.03 | 14.42 | 6.65 | 0.57 | 314.07 | 449.18 | 14.02 | -0.24 | -0.26 | 394.48 | 2.11 | 0.19 | 0 | -0.3 |
| Run2_Nl-3 | Normal | 0.43 | 105.14 | 102.53 | 89.93 | 279.5 | 548.48 | 105.39 | 47.14 | 73.77 | 0.16 | 0.09 | 0.32 | 0.46 | 14.19 | 12.34 | 0.75 | 144.94 | 223.18 | 12.43 | -0.24 | -0.3 | 282.02 | 0.62 | 0.11 | -0.11 | -0.31 |
| Run2_Nl-4 | | 0.49 | 246.02 | 241.3 | 247.12 | 494.48 | 731.66 | 136.07 | 70.47 | 122.92 | 0.27 | 0.14 | 0.36 | 0.44 | 21.16 | 13.32 | 0.71 | 178.27 | 454.08 | 37.54 | -0.18 | -0.18 | 476.36 | 1.35 | 0.64 | -0.15 | -0.2 |
| Run2_Nl-5 | | 0.51 | 103.49 | 99.43 | 75.96 | 241.61 | 372.91 | 81.88 | 43.41 | 55.06 | 0.27 | 0.16 | 0.36 | 0.47 | 5.65 | 2.39 | 0.69 | 172.52 | 153.32 | 6.71 | -0.28 | -0.32 | 110.96 | 1.24 | -0.19 | -0.26 | -0.31 |
| Run2_Nl-6 | | 0.39 | 117.24 | 115.63 | 106.58 | 233.41 | 596.59 | 154.36 | 83.18 | 99.1 | 0.34 | 0.19 | 0.22 | 0.26 | 7.91 | 3.38 | 0.55 | 326.24 | 487.28 | 14.3 | -0.22 | -0.25 | 455.06 | 1.33 | -0.17 | 0.06 | -0.29 |

**Supplemental Table 3**: Median expression of each surface marker on each cell population.

### CD44 Median

| | | Ungated | CD34+CD38low | HSCs | MPP | CMP/GMP | Myelo/Mono-Blasts | ProMonocytes | CD14neg Monocytes | Mature Monocytes | ProMyelocytes | Myelocytes | MetaMyelocytes | MatureGrans | ProErythroblasts | Erythroblasts | LateErythroblasts | PreBcells | Mature Bcells | Plasma Cells | T cells | NK cells | pDCs | Basophils | Platelets | CD8+ T cells | CD8neg T cells |
|---|---|---|---|---|---|---|---|---|---|---|---|---|---|---|---|---|---|---|---|---|---|---|---|---|---|---|---|
| Run1_MDS17 | | 134.56 | 225.04 | 235.05 | 217.9 | 245.11 | 142.17 | 140.02 | 378.72 | 445.03 | 53.84 | 48.68 | 175.8 | 231.8 | 28.04 | 22.22 | 19.88 | 219.99 | 142.29 | 380.55 | 221.73 | 67.79 | 129.31 | 129.24 | 8.48 | 219.85 | 224.42 |
| Run2_MDS15* | AML/RAEB-T | 356.33 | 324.31 | 383.49 | 362.69 | 250.22 | 180.62 | 190.75 | 571.76 | 685.87 | 144.14 | 146.12 | 314.31 | 395.69 | 245.51 | 282.56 | 411.68 | 561.81 | 225.46 | 327.12 | 328.67 | 234.65 | 506.91 | 156.44 | 160.56 | 324.61 | 331.76 |
| Run2_MDS4 | | 208.86 | 77.07 | 75.99 | 50.3 | 8.73 | 110.53 | 441.83 | 668.42 | 698.72 | 31.51 | 42.51 | 184.23 | 310.31 | 13.93 | 15.76 | 8.51 | 68.72 | 36.03 | 129.13 | 360.54 | 477.48 | 86.95 | 74.9 | 12.46 | 401.19 | 324.74 |
| Run1_MDS21 | | 62.48 | 80.67 | 76.75 | 76.93 | 104.15 | 96.48 | 77.43 | 302.55 | 338.56 | 12.32 | 16.31 | 93.45 | 141.43 | 38.95 | 43.25 | 16.83 | 95.72 | 165.85 | 243.03 | 129.46 | 173.93 | 89.97 | 40.69 | 2.21 | 154.21 | 119.13 |
| Run1_MDS3 | | 70.38 | 97.65 | 95.33 | 87 | 129.12 | 117.41 | 97.31 | 319 | 441.59 | 16.61 | 23.78 | 126.57 | 163.39 | 49.13 | 53.18 | 45.88 | 120.17 | 206.29 | 190.64 | 223.89 | 219.75 | 238.69 | 34.31 | 4.73 | 217.17 | 227.07 |
| Run2_MDS13 | | 108.85 | 176.38 | 189.02 | 158.6 | 135.88 | 117.38 | 163.86 | 474.37 | 540.64 | 19.87 | 30.17 | 240.99 | 272.55 | 112.76 | 37.21 | 17.95 | 124.38 | 179.54 | 337.74 | 230.9 | 277.08 | 296.3 | 57.97 | 4.11 | 246.54 | 220.84 |
| Run2_MDS16 | | 145.73 | 145.77 | 151.2 | 138.88 | 197.03 | 408.81 | 590.73 | 596.54 | 750.36 | 5.85 | 8.26 | 80.95 | 157.52 | 101.51 | 17.71 | 73.36 | 261.73 | 127.12 | 394.23 | 285.24 | 92.49 | 308.64 | 163.65 | 4.86 | 272 | 288.4 |
| Run2_MDS1 | | 172.22 | 199.44 | 121.24 | 203.39 | 224.81 | 178.65 | 313.48 | 646.81 | 763.42 | 8.74 | 21.5 | 120.33 | 186.83 | 36.79 | 20.86 | 13.61 | 244.09 | 250.14 | 235.83 | 259.68 | 224.38 | 200.22 | 17.55 | 5.49 | 274.6 | 257.38 |
| Run2_MDS21 | Higher Risk | 75.73 | 88.18 | 92.16 | 84.24 | 117.98 | 97.88 | 79.83 | 406.83 | 423.58 | 15.18 | 18.73 | 192.45 | 247.58 | 42.75 | 26.66 | 13.58 | 107.24 | 179.59 | 310.39 | 174.26 | 204.19 | 148.72 | 45.52 | 2.88 | 214.76 | 160.59 |
| Run2_MDS26* | | 38.36 | 65.76 | 69.33 | 65.31 | 67.25 | 23.76 | 54.33 | 419.19 | 484.12 | 5.6 | 9.18 | 75.02 | 129.37 | 19.87 | 37.56 | 15.4 | 21.58 | 73.46 | 233.97 | 185.93 | 43.72 | 341.31 | 24.44 | 6.32 | 212.54 | 171.1 |
| Run2_MDS27 | | 126.51 | 94.18 | 99.32 | 88.18 | 88.46 | 71.26 | 280.47 | 423.31 | 650.09 | 6.34 | 7.15 | 81.19 | 232.67 | 39.22 | 24.08 | 9.33 | 239.81 | 108.72 | 287.41 | 382.92 | 104.73 | 698.59 | 144.03 | 12.13 | 385.63 | 379.42 |
| Run2_MDS2 | | 237.96 | 552.55 | 558.7 | 547.18 | 435.17 | 382.41 | 590.03 | 616.5 | 805.84 | 35.24 | 30.89 | 193.49 | 207.52 | 354.33 | 211.1 | 95.95 | 309.42 | 563.88 | 456.33 | 281.8 | 72.19 | 430.36 | 109.08 | 1.98 | 288.03 | 281.34 |
| Run2_MDS3 | | 101.28 | 114.15 | 128.31 | 103.58 | 159.69 | 128.37 | 135.24 | 436.22 | 456.36 | 30.21 | 45.03 | 244.12 | 280.93 | 136.26 | 47.75 | 21.12 | 145.5 | 241.25 | 375.61 | 302.41 | 292.57 | 84.16 | 55.65 | 6.06 | 299.89 | 303.38 |
| Run1_MDS23 | | 29.22 | 33.72 | 31.07 | 35.28 | 58.07 | 48.57 | 129.68 | 236.65 | 342.3 | 3.08 | 3.58 | 18.44 | 44.1 | 49.39 | 59.8 | 46.35 | 146.85 | 86.63 | 52.59 | 186.45 | 192.26 | 106.02 | 27.6 | 15.34 | 210.48 | 178.24 |
| Run1_MDS25 | | 93.07 | 59.82 | 28.49 | 97.75 | 71.21 | 257.59 | 306 | 333.62 | 355.18 | 5.87 | 9.86 | 62.8 | 90.23 | 40.52 | 56.82 | 83.85 | 229.03 | 100.48 | 159.25 | 203.5 | 61.94 | 92.67 | 97.61 | 3.18 | 193.97 | 216.1 |
| Run1_MDS5 | | 298.19 | 211.15 | 193.25 | 202.84 | 124.34 | 165.91 | 286.13 | 292.7 | 545 | 10.13 | 111.96 | 245.02 | 357.52 | 177.83 | 36.38 | 71.27 | 307.32 | 116.69 | 271.48 | 212.52 | 302.4 | 165.5 | 98.82 | 16.09 | 253.45 | 192.82 |
| Run1_MDS6 | | 144.69 | 46.48 | 41.61 | 38.3 | 45.02 | 83.68 | 274.06 | 402.34 | 475.44 | 19.65 | 20.43 | 90.48 | 167.73 | 38.69 | 15.17 | 101.48 | 157.68 | 123.91 | 365.26 | 216.06 | 141.08 | 215.05 | 113.27 | 4.2 | 235.37 | 201.07 |
| Run1_MDS8 | | 87.99 | 96.55 | 73.59 | 121.85 | 83.19 | 163.1 | 425.4 | 485.37 | 518.21 | 5.83 | 10.56 | 74.34 | 99.73 | 89.58 | 40.07 | 21.71 | 90.84 | 104.05 | 230.25 | 64.2 | 65.46 | 81.86 | 113.6 | 4.74 | 73.89 | 61.7 |
| Run1_MDS9 | | 53.75 | 77.56 | 82.98 | 63.65 | 81.93 | 94.02 | 362.97 | 405.29 | 386.4 | 3.95 | 6.01 | 32.2 | 58.3 | 33.3 | 29.63 | 31.23 | 138.34 | 74.68 | 246.77 | 133.03 | 73.48 | 322.27 | 142.72 | 6.49 | 176.22 | 111.47 |
| Run2_MDS12 | Lower Risk | 64.85 | 100.79 | 90.57 | 81.48 | 163.23 | 180.99 | 283.31 | 433.92 | 631.11 | 3.14 | 4.89 | 58.59 | 118.27 | 40.87 | 33.32 | 22.58 | 236.02 | 27.61 | 209.34 | 183.78 | 219.45 | 189.48 | 61.91 | 15.71 | 239.52 | 152.53 |
| Run2_MDS14 | | 159.28 | 135.12 | -0.35 | 229.73 | 84.29 | 410.44 | 521.45 | 528.55 | 660.49 | 4.57 | 7.28 | 46.54 | 169.29 | 33.49 | 17.03 | 160.59 | 8 | 124.21 | 496.68 | 324.57 | 91.32 | 170.72 | 110.09 | 8.13 | 297.3 | 342.49 |
| Run2_MDS19 | | 90.57 | 94.64 | 103.8 | 94.11 | 107.42 | 148.56 | 248.66 | 364.12 | 508.16 | 3.94 | 6.61 | 57.34 | 130.92 | 30.32 | 18.39 | 9.22 | 323.32 | 148.47 | 37.23 | 303.49 | 77.55 | 60.6 | 116.67 | 3.54 | 264.68 | 319.43 |
| Run2_MDS20 | | 177.99 | 94.3 | 93.02 | 90.38 | 136.46 | 327.31 | 503.84 | 581.95 | 638.44 | 6.98 | 8.37 | 62.71 | 171.35 | 75.69 | 38.54 | 160.68 | 281.84 | 95.84 | 381.63 | 207.51 | 57.82 | 359.9 | 150.14 | 4.34 | 247.56 | 204.03 |
| Run2_MDS28* | | 94.98 | 75.8 | 88.43 | 57.13 | 95.86 | 72.07 | 149.33 | 361.02 | 581.54 | 3.59 | 4.32 | 42.78 | 127.52 | 45.65 | 25.5 | 70.15 | 322.56 | 101.93 | 76.51 | 250.24 | 197.34 | 418.16 | 142.85 | 5.06 | 252.02 | 248.49 |
| Run2_MDS5 | | 441.85 | 261.38 | 283.46 | 141.82 | 195.97 | 211.68 | 323.75 | 394.87 | 797.72 | 13.13 | 190.4 | 383.65 | 533.12 | 140.77 | 35.1 | 253.63 | 325.13 | 144.92 | 268.07 | 299.52 | 422.76 | 203.23 | 156.76 | 11.22 | 351.64 | 277.82 |
| Run1_ICUS18 | | 109.75 | 82.99 | 76.21 | 69.16 | 76.9 | 319.72 | 427.78 | 482.68 | 513.47 | 6.14 | 6.48 | 43.92 | 105.61 | 57.55 | 43.59 | 134.86 | 214.23 | 143.23 | 56.44 | 214.12 | 77.76 | 226.38 | 120.3 | 2.74 | 223.2 | 212.02 |
| Run1_ICUS22 | ICUS | 65.5 | 65.74 | 80.78 | 60.24 | 74.07 | 108.86 | 309.85 | 330.61 | 388.06 | 5.13 | 5.12 | 23.65 | 66.77 | 33.52 | 39.36 | 71.94 | 28.77 | 55.04 | 111.6 | 215.53 | 21.83 | 347.8 | 76.71 | 2.82 | 217.99 | 214.51 |
| Run1_ICUS24 | | 65.75 | 82.41 | 80.27 | 78.25 | 96.12 | 117.37 | 222.55 | 318.35 | 477.82 | 6.43 | 6.01 | 27.9 | 85.09 | 30.24 | 27.03 | 16.86 | 3.41 | 63.64 | 123.4 | 185.19 | 50.88 | 75.32 | 57.98 | 3.37 | 167.1 | 202.88 |
| Run1_Nl-1 | | 77.28 | 77.77 | 97.44 | 57 | 77.4 | 135.03 | 300.39 | 327.82 | 503.64 | 5.04 | 4.84 | 27.3 | 99.24 | 38.95 | 43.53 | 82.38 | 5.48 | 64.27 | 169.2 | 178.26 | 36.33 | 228.13 | 108.61 | 2.8 | 178.2 | 178.29 |
| Run1_Nl-3 | | 77.7 | 87.31 | 93.56 | 85.09 | 95.19 | 132.23 | 172.22 | 270.84 | 433.98 | 6.4 | 8.66 | 54.7 | 105.56 | 89.8 | 44.57 | 108.41 | 52.32 | 65.55 | 118.05 | 182.93 | 74.17 | 134.91 | 124.31 | 6.61 | 173.48 | 189.84 |
| Run1_Nl-4 | | 87.88 | 84.41 | 89.37 | 77.46 | 96.11 | 130.91 | 196.23 | 317.03 | 467.22 | 7.32 | 9.91 | 64.7 | 122.37 | 104.23 | 45.61 | 119.87 | 63.83 | 118.3 | 347.82 | 198.17 | 67.14 | 298.37 | 113.19 | 3.93 | 194.88 | 200.21 |
| Run1_Nl-6 | | 53.7 | 106.61 | 122.93 | 96.62 | 93.17 | 140.86 | 204.07 | 328.02 | 480.9 | 6.73 | 7.27 | 32.44 | 89.19 | 58.79 | 36.46 | 55.8 | 8.1 | 62.28 | 278.49 | 209.65 | 51.12 | 170.24 | 62.23 | 2.77 | 203.66 | 212.4 |
| Run2_Nl-3 | Normal | 118.88 | 117.96 | 130.42 | 109.55 | 135.71 | 177.96 | 263.27 | 444.22 | 643.26 | 7.38 | 14 | 110.9 | 170.73 | 91.21 | 49.55 | 163.64 | 43.93 | 85.12 | 142.74 | 266.01 | 103.16 | 176.49 | 167.83 | 5.14 | 248.12 | 280.46 |
| Run2_Nl-4 | | 152.56 | 141.68 | 137.36 | 123.07 | 145.75 | 175.76 | 295.09 | 532.67 | 698.28 | 9.01 | 16.9 | 138.54 | 215.6 | 96.44 | 49.64 | 203.66 | 54.7 | 164.47 | 549.08 | 296.88 | 97.02 | 372.66 | 167.12 | 3.9 | 289.76 | 303.9 |
| Run2_Nl-5 | | 122.17 | 89.04 | 112.92 | 69.57 | 111.02 | 193.21 | 297.44 | 554.65 | 714.07 | 5.71 | 9.25 | 88.23 | 184.27 | 77.19 | 32.09 | 160.95 | 4.44 | 93.01 | 247.09 | 234.02 | 72.8 | 278.34 | 171.11 | 4.49 | 224.77 | 244.17 |
| Run2_Nl-6 | | 89.89 | 141.13 | 156.33 | 129.91 | 143.25 | 183.82 | 292.28 | 492.2 | 685.57 | 7.34 | 10.15 | 61.81 | 157.34 | 44.58 | 32.67 | 50.72 | 7.68 | 76.42 | 363.5 | 297.57 | 69.89 | 208.24 | 91.31 | 4.66 | 293.21 | 299.02 |

**Supplemental Table 3**: Median expression of each surface marker on each cell population.

### CD38 Median

| | | Ungated | CD34+CD38low | HSCs | MPP | CMP/GMP | Myelo/Mono-Blasts | ProMonocytes | CD14neg Monocytes | Mature Monocytes | ProMyelocytes | Myelocytes | MetaMyelocytes | MatureGrans | ProErythroblasts | Erythroblasts | LateErythroblasts | PreBcells | Mature Bcells | Plasma Cells | T cells | NK cells | pDCs | Basophils | Platelets | CD8+ T cells | CD8neg T cells |
|---|---|---|---|---|---|---|---|---|---|---|---|---|---|---|---|---|---|---|---|---|---|---|---|---|---|---|---|
| Run1_MDS17 | | 4.03 | 5.78 | 6.07 | 5.33 | 50.56 | 8.2 | 3.09 | 2.41 | 2.43 | 1.47 | 3.3 | 4.08 | 5.49 | 7.04 | 5.7 | 3.19 | 4.38 | 1.67 | 1903.67 | 2.17 | 78.56 | 3.74 | 108.61 | 1.77 | 2.13 | 2.28 |
| Run2_MDS15* | AML/RAEB-T | 10.75 | 1.89 | 4.48 | 4.55 | 87.15 | 57.8 | 29.66 | 24.8 | 27.93 | 27.03 | 12.32 | 5.88 | 5.16 | 28.62 | 15.27 | 13.75 | 16.43 | 0.05 | 1747.52 | 1.45 | 78.84 | 26.78 | 124.17 | 15.81 | 1.94 | 0.98 |
| Run2_MDS4 | | 2.44 | 1.5 | 0.44 | 1.12 | 67.38 | 9.62 | 6.35 | 3.98 | 7.19 | 7.14 | 4.56 | 2.21 | 1.81 | 6.32 | 5.54 | 1.76 | 11.72 | 27.54 | 1836.2 | 0.41 | 9.94 | 10.6 | 52.32 | 4.01 | 0.2 | 0.62 |
| Run1_MDS21 | | 6.1 | 25.32 | 25.17 | 24.9 | 72.15 | 69.92 | 46.25 | 34.3 | 36.36 | 1.36 | 2.95 | 4.46 | 7.33 | 12.86 | 17.26 | 30.87 | 0.05 | 0.65 | 1835.26 | 0.89 | 27.04 | 23.29 | 257.65 | 21.49 | 0.78 | 0.94 |
| Run1_MDS3 | | 3.84 | 26.85 | 25.59 | 26.1 | 76.41 | 73.19 | 39.62 | 13.88 | 31.21 | 1.1 | 1.85 | 3.57 | 6.61 | 12.92 | 12.99 | 8.02 | 0.12 | 0.58 | 1761.88 | 0.98 | 41.18 | 36.74 | 286.35 | 13.14 | 0.81 | 1.08 |
| Run2_MDS13 | | 19.17 | 21.61 | 20.43 | 21.51 | 100.28 | 105.34 | 78.3 | 98.36 | 103.76 | 25.19 | 30.26 | 32.93 | 30.52 | 39.63 | 9.76 | 5.25 | -0.02 | -0.02 | 2005.81 | 0.98 | 21.61 | 158.5 | 228.79 | 9.67 | 1.12 | 0.92 |
| Run2_MDS16 | | 5.94 | 7.24 | 6.53 | 6.05 | 93.8 | 32.44 | 7.74 | 39.1 | 24.22 | 31.92 | 9.47 | 6.41 | 5.38 | 16.69 | 10.21 | 6.29 | 4.45 | 5.02 | 2276.59 | 2.38 | 140.06 | 31.2 | 254.71 | 19.69 | 1.4 | 2.7 |
| Run2_MDS1 | | 5.88 | 25.5 | 17.45 | 27.58 | 94.35 | 57.89 | 21.6 | 8.77 | 8.39 | 4.89 | 2.34 | 3.94 | 3.47 | 38.71 | 6.93 | 2.21 | 12.13 | 2.25 | 1837.55 | 2.95 | 139.86 | 17.57 | 231.16 | 5.44 | 3.7 | 2.84 |
| Run2_MDS21 | Higher Risk | 14.02 | 24.21 | 24.67 | 23.67 | 79.48 | 73.52 | 76.2 | 126.4 | 109.07 | 16.86 | 36.43 | 59.66 | 51.85 | 11.66 | 8.36 | 5.33 | -0.12 | 0.08 | 2086.94 | 0.65 | 2.33 | 10.64 | 286.74 | 17.6 | 0.52 | 0.73 |
| Run2_MDS26* | | 4.75 | 7.72 | 4.77 | 7.09 | 68.39 | 30.26 | 33.95 | 27.79 | 44.68 | 9.28 | 5.73 | 5.29 | 4.91 | 21.88 | 4.63 | 2.03 | 4.47 | 4.89 | 1362.25 | 2.8 | 166.2 | 24.97 | 192.12 | 11.65 | 1.36 | 4.11 |
| Run2_MDS27 | | 6.1 | 5.34 | 4.39 | 5.36 | 56.98 | 47.46 | 26.44 | 45.97 | 28.08 | 47.68 | 17.77 | 9.63 | 6.23 | 23.31 | 8.43 | 1.71 | 1.41 | 1.24 | 2876.1 | 0.19 | 72.71 | -0.04 | 259.29 | 11.19 | 0.05 | 0.37 |
| Run2_MDS2 | | 16.83 | 17.07 | 19.51 | 16.02 | 74.41 | 67.8 | 58.57 | 81.3 | 109.89 | 41.75 | 6.75 | 16.89 | 15.27 | 105.31 | 19.85 | 12.77 | 12.43 | 14.6 | 2010.74 | 2.4 | 237.45 | 114.78 | 125.77 | 15.19 | 1.01 | 2.62 |
| Run2_MDS3 | | 13.14 | 25.85 | 26.69 | 25.23 | 106.26 | 106.41 | 71.74 | 63.84 | 77.6 | 16.16 | 29.07 | 43.09 | 36.23 | 31.46 | 9.64 | 4.73 | -0.09 | 0.24 | 2078.01 | 0.74 | 35.7 | 38.13 | 278.23 | 13.53 | 0.57 | 0.82 |
| Run1_MDS23 | | 5.02 | 15.72 | 13.32 | 15.87 | 112.86 | 30.27 | 30.25 | 28.69 | 25.23 | 5.31 | 3.9 | 3.91 | 4.21 | 22.48 | 21.71 | 9.67 | 21.94 | 1.47 | 1793.97 | 2.55 | 119.03 | 32.43 | 318.62 | 30.24 | 2.39 | 2.63 |
| Run1_MDS25 | | 2.79 | 15.72 | 14.49 | 6.68 | 223.42 | 10.63 | 46.03 | 49.19 | 31.06 | 5.75 | 3.1 | 2.06 | 2.34 | 29.31 | 27.18 | 5.97 | 17.25 | 2.6 | 2636.81 | 4.26 | 249.64 | 58.32 | 376.99 | 4 | 3.51 | 5.17 |
| Run1_MDS5 | | 3.54 | 10.8 | 1.35 | 6.43 | 248.59 | 144.43 | 31.7 | 26.72 | 52.21 | 10.94 | 3.2 | 2.57 | 3.03 | 49.81 | 11.93 | 5.75 | 8.55 | 5.51 | 2047.87 | 2.58 | 46.47 | 174.32 | 305.27 | 4.92 | 39.78 | 1.62 |
| Run1_MDS6 | | 4.15 | 20.25 | 19.13 | 15.96 | 68.02 | 46.46 | 25.45 | 31.15 | 27.65 | 6.67 | 4.76 | 4.09 | 3.31 | 42.49 | 13.53 | 5.9 | 9.69 | 4.34 | 2334.2 | 2.54 | 89.97 | 46.65 | 164.75 | 16.7 | 1.31 | 4.37 |
| Run1_MDS8 | | 3.31 | 21.03 | 23.96 | 21.36 | 164.63 | 61.84 | 12.04 | 35.31 | 32.92 | 9.72 | 5.05 | 2.35 | 2.4 | 38.94 | 20.85 | 5.93 | 10.87 | 5.77 | 1729.74 | 4.28 | 164.76 | 201.63 | 323.2 | 16.36 | 2.42 | 5.01 |
| Run1_MDS9 | | 2.4 | 16.15 | 17.1 | 15.45 | 76.18 | 48.92 | 2.52 | 19.63 | 29.73 | 3.66 | 3.03 | 1.92 | 1.92 | 22.66 | 22.64 | 10.07 | 2.31 | 0.72 | 2208.21 | 0.71 | 72.55 | 1.82 | 246.9 | 8.72 | -0.07 | 1.63 |
| Run2_MDS12 | Lower Risk | 9.82 | 15.13 | 13.89 | 17.82 | 70.97 | 27.86 | 32.05 | 26.62 | 20.18 | 12.85 | 11.91 | 10.11 | 8.43 | 9.72 | 6.5 | 4.64 | 18.42 | -0.09 | 1723.11 | 1.42 | 56.76 | 30.5 | 321.25 | 15.93 | 1.23 | 1.61 |
| Run2_MDS14 | | 4.88 | 5.74 | 0.26 | 3.09 | 295.46 | 20.16 | 5.54 | 37.63 | 31.55 | 20.04 | 7.16 | 5.97 | 3.84 | 20.55 | 12.15 | 4.52 | 393.28 | 19.52 | 1883.21 | 1.8 | 461.74 | 82.49 | 249.47 | 10.88 | 2.94 | 1.43 |
| Run2_MDS19 | | 3.74 | 10.68 | 7.69 | 10.76 | 113.59 | 48.4 | 34.07 | 42.86 | 28.64 | 11.65 | 5.66 | 3.98 | 2.73 | 11 | 5.48 | 1.94 | 6.47 | 0.8 | 2320.43 | 0.47 | 120.84 | 29.94 | 212.17 | 5.71 | 0.27 | 0.55 |
| Run2_MDS20 | | 7.05 | 12.11 | 16.14 | 12.56 | 145.01 | 7.8 | 8.45 | 48.27 | 54.34 | 26.19 | 12.7 | 8.62 | 5.6 | 43.55 | 11.38 | 7.14 | 16.63 | 0.2 | 1903.71 | 7.28 | 63.79 | 1.76 | 232.17 | 9 | 3.73 | 7.57 |
| Run2_MDS28* | | 12.58 | 23.2 | 22.71 | 22.49 | 103.98 | 47.57 | 43.77 | 57.24 | 44.87 | 23.76 | 14.75 | 13.64 | 9.65 | 93.17 | 6.77 | 9.51 | 29.6 | 1.61 | 1213.47 | 0.98 | 127.25 | 57.24 | 141.28 | 25.71 | 0.57 | 1.82 |
| Run2_MDS5 | | 3.21 | 8.25 | 7.8 | 23.71 | 198.37 | 143.82 | 15.89 | 21.7 | 44.11 | 26.33 | 2.73 | 2.39 | 2.9 | 377.8 | 9.29 | 4.4 | 5.05 | 3.85 | 2074.06 | 2.07 | 39.13 | 162.83 | 319.1 | 5.06 | 48.27 | 1.06 |
| Run1_ICUS18 | | 3.82 | 13.36 | 24.12 | 10.25 | 83.25 | 3.85 | 5.73 | 18.26 | 19.99 | 7.27 | 5.74 | 4.6 | 3.25 | 17.37 | 14.68 | 4.16 | 12.3 | 4.19 | 1954.54 | 2.69 | 86.05 | 22.38 | 188.86 | 20.8 | 0.62 | 3.61 |
| Run1_ICUS22 | ICUS | 4.1 | 19.54 | 17.63 | 19.39 | 111.74 | 32.11 | 20.17 | 65.3 | 58.7 | 8.07 | 6.25 | 4.72 | 3.06 | 29.19 | 29.04 | 4.81 | 151.34 | 15.95 | 1694.7 | 1.9 | 436.74 | 3.47 | 164.13 | 8.49 | 1.6 | 2.04 |
| Run1_ICUS24 | | 4.26 | 18.67 | 13.87 | 20.26 | 129.79 | 79.75 | 50.55 | 42.09 | 33.94 | 5.15 | 3.94 | 3.9 | 3.27 | 19.88 | 14.29 | 10.67 | 277.35 | 6 | 1758.26 | 1.46 | 241.37 | 41.86 | 141.46 | 19.41 | 2.02 | 1.07 |
| Run1_Nl-1 | | 3.45 | 18.48 | 14.38 | 19.03 | 189.21 | 83.32 | 32.48 | 39.86 | 29.34 | 5.91 | 3.63 | 3.23 | 2.49 | 20.41 | 18.06 | 5.14 | 372.2 | 17.29 | 1875.65 | 4.8 | 214.48 | 90.87 | 132.64 | 30.1 | 2.46 | 7.35 |
| Run1_Nl-3 | | 7.41 | 20.73 | 17.28 | 18.59 | 114.55 | 65.85 | 48.57 | 47.34 | 30.44 | 15.5 | 9.29 | 7.27 | 5.31 | 35.32 | 14.71 | 8.36 | 58.2 | 7.48 | 2137.68 | 2.29 | 61.25 | 40.65 | 184.47 | 8.86 | 1.26 | 3.86 |
| Run1_Nl-4 | | 6.69 | 19.08 | 19.41 | 18 | 138.33 | 72.98 | 55.64 | 47.41 | 32.67 | 11.96 | 7.77 | 6.26 | 5.15 | 38.05 | 15.77 | 7.31 | 53.76 | 5.23 | 2501.99 | 1.71 | 147.81 | 57.2 | 191.67 | 9.38 | 1.29 | 2.28 |
| Run1_Nl-6 | | 6.87 | 11.7 | 8.35 | 10.75 | 144.68 | 84.87 | 53.38 | 39.61 | 27.07 | 10.78 | 7.52 | 6.58 | 5.19 | 27.17 | 10.66 | 6.88 | 285.35 | 7.29 | 2299.34 | 2.77 | 142.56 | 18.86 | 282.38 | 8.9 | 1.97 | 3.24 |
| Run2_Nl-3 | Normal | 7.26 | 19.27 | 17.54 | 19.67 | 124.23 | 82.58 | 59.72 | 45.04 | 27.31 | 30.12 | 9.46 | 6.53 | 5.26 | 26.18 | 10.76 | 6.11 | 72.03 | 5.74 | 2017.48 | 1.41 | 58.79 | 32.82 | 171.88 | 9.01 | 0.77 | 2.3 |
| Run2_Nl-4 | | 7.97 | 18.83 | 14.45 | 21.35 | 132.45 | 92.79 | 69.01 | 49.14 | 32.47 | 28.99 | 11.15 | 7.35 | 5.63 | 25.99 | 10.79 | 7.32 | 108.12 | 3.61 | 2619.1 | 1.06 | 140.2 | 48.13 | 177.96 | 10.42 | 0.8 | 1.47 |
| Run2_Nl-5 | | 7.81 | 18.05 | 14.12 | 21.46 | 164.01 | 142.23 | 84.02 | 52.17 | 36.62 | 30.1 | 10.32 | 6.63 | 4.91 | 60.22 | 11.3 | 5.86 | 376.94 | 7.1 | 2463.39 | 3.85 | 113.88 | 40.55 | 132.37 | 6.66 | 2.76 | 6.03 |
| Run2_Nl-6 | | 9.47 | 9.28 | 6.37 | 9.02 | 117.53 | 94.76 | 59.32 | 45.42 | 28.21 | 26.04 | 14.14 | 9.83 | 6.89 | 16.05 | 6.97 | 6.09 | 278.77 | 5.38 | 2390.24 | 1.71 | 135.45 | 13.93 | 246.82 | 13.9 | 1.09 | 2.16 |

**Supplemental Table 3**: Median expression of each surface marker on each cell population.

### CD14 Median

| | | Ungated | CD34+CD38low | HSCs | MPP | CMP/GMP | Myelo/Mono-Blasts | ProMonocytes | CD14neg Monocytes | Mature Monocytes | ProMyelocytes | Myelocytes | MetaMyelocytes | MatureGrans | ProErythroblasts | Erythroblasts | LateErythroblasts | PreBcells | Mature Bcells | Plasma Cells | T cells | NK cells | pDCs | Basophils | Platelets | CD8+ T cells | CD8neg T cells |
|---|---|---|---|---|---|---|---|---|---|---|---|---|---|---|---|---|---|---|---|---|---|---|---|---|---|---|---|
| Run1_MDS17 | | 0.1 | -0.02 | 0.04 | -0.06 | 0.54 | 0 | 0.23 | 0.98 | 11.2 | -0.15 | 0.2 | 0.14 | 0.46 | 0.89 | 1.44 | 0.82 | 0.05 | -0.27 | 7.45 | -0.16 | 0.97 | -0.2 | 2.09 | 0.1 | -0.19 | -0.13 |
| Run2_MDS15* | AML/RAEB-T | 1.41 | -0.21 | -0.16 | -0.2 | 0.46 | 0.64 | 1.16 | 2.48 | 35.2 | 0.78 | 0.78 | 1.41 | 1.06 | 0.43 | 3.57 | 3.37 | 7.23 | -0.29 | 4.42 | -0.13 | 0.72 | 0.71 | 1.18 | 0.57 | -0.12 | -0.13 |
| Run2_MDS4 | | 0.06 | -0.22 | -0.25 | -0.25 | -0.53 | -0.12 | 0.35 | 1.01 | 11.42 | 0.05 | -0.02 | -0.07 | 0.06 | 0.58 | 2 | 1.85 | 0.52 | 0.13 | 6.89 | -0.18 | -0.08 | -0.06 | 0.89 | 1.36 | -0.2 | -0.16 |
| Run1_MDS21 | | 0.07 | 0.33 | 0.14 | 0.25 | 0.65 | 0.63 | 0.72 | 2.13 | 11.43 | -0.23 | -0.08 | 0.08 | 0.54 | 0.76 | 1.51 | 1.38 | -0.28 | -0.32 | 9.27 | -0.25 | 0.99 | -0.09 | 2.05 | 0.69 | -0.24 | -0.25 |
| Run1_MDS3 | | 0.02 | 0.47 | 0.41 | 0.37 | 0.71 | 0.75 | 0.68 | 1.28 | 11.26 | -0.22 | -0.1 | 0.08 | 0.61 | 0.83 | 1.22 | 1.04 | -0.24 | -0.3 | 6 | -0.18 | 0.87 | 0.39 | 2.21 | 0.35 | -0.2 | -0.17 |
| Run2_MDS13 | | 0.28 | 0.02 | 0.12 | -0.01 | 0.52 | 0.58 | 0.72 | 1.19 | 9.82 | 0.36 | 0.61 | 0.46 | 0.64 | 1.27 | 1.79 | 1.12 | -0.2 | -0.28 | 4.13 | -0.17 | 0.35 | 1.96 | 1.35 | 0.25 | -0.15 | -0.19 |
| Run2_MDS16 | | 0.3 | -0.03 | -0.05 | -0.12 | 0.4 | 0.26 | 1.25 | 3.47 | 20.01 | 0.54 | 0.2 | 0.19 | 0.35 | 0.42 | 3.02 | 1.45 | 0.03 | -0.21 | 6.45 | -0.16 | 1.37 | 0.41 | 1.87 | 0.71 | -0.2 | -0.14 |
| Run2_MDS1 | | 0.19 | -0.07 | -0.27 | -0.09 | 0.55 | 0.22 | 0.35 | 1.79 | 16.49 | -0.11 | -0.21 | -0.02 | 0.18 | 0.88 | 2.05 | 1.21 | 0.66 | -0.21 | 4.66 | -0.1 | 0.85 | -0.03 | 1.09 | 0.3 | -0.1 | -0.1 |
| Run2_MDS21 | Higher Risk | 0.28 | 0.11 | 0.36 | 0.04 | 0.46 | 0.45 | 0.52 | 1.18 | 10.66 | 0.22 | 0.55 | 0.66 | 0.85 | 0.84 | 1.6 | 1.1 | -0.25 | -0.26 | 5.11 | -0.23 | -0.14 | 0.34 | 2.28 | 0.5 | -0.22 | -0.23 |
| Run2_MDS26* | | 0.18 | 0.13 | 0.21 | 0.01 | 0.77 | 0.44 | 0.8 | 0.96 | 11.88 | 0.11 | 0.06 | -0.02 | 0.11 | 0.86 | 1.89 | 1.2 | 0.26 | -0.2 | 5.48 | -0.18 | 1.41 | 0.4 | 1.6 | -0.22 | -0.22 | -0.15 |
| Run2_MDS27 | | 0.5 | -0.12 | -0.16 | -0.15 | 0.43 | 0.22 | 1.26 | 3.01 | 21.24 | 0.88 | 0.53 | 0.34 | 0.42 | 0.46 | 3.43 | 1.8 | -0.02 | -0.26 | 21.23 | -0.2 | 0.87 | -0.04 | 1.66 | 1.18 | -0.22 | -0.18 |
| Run2_MDS2 | | 0.18 | -0.2 | 0.12 | -0.23 | 0 | 0.01 | 0.88 | 1.78 | 17.06 | 0.46 | 0.12 | 0.21 | 0.48 | 1.44 | 1.93 | 1.39 | 0.48 | -0.19 | 89.89 | -0.12 | 1.17 | 0.31 | 0.76 | 0.32 | -0.12 | -0.12 |
| Run2_MDS3 | | 0.34 | 0.16 | 0.19 | 0.16 | 0.51 | 0.54 | 0.72 | 0.7 | 9.8 | 0.25 | 0.68 | 0.62 | 0.97 | 0.96 | 1.79 | 1.14 | -0.23 | -0.25 | 6.16 | -0.19 | 0.59 | 0.76 | 2.01 | 0.5 | -0.19 | -0.19 |
| Run1_MDS23 | | 0.19 | -0.08 | 0.1 | 0.06 | 0.45 | 0.14 | 1.1 | 2.9 | 21.15 | -0.1 | -0.07 | -0.01 | 0.11 | 0.85 | 1.31 | 1.4 | 1.13 | -0.29 | 5.5 | -0.14 | 0.92 | -0.04 | 2.36 | 0.89 | -0.11 | -0.15 |
| Run1_MDS25 | | 0.35 | 1.03 | 0.56 | 0.81 | 1.27 | 0.02 | 1.4 | 2.81 | 14.35 | 0.21 | 0.14 | 0.07 | 0.26 | 1.06 | 1.94 | 1.42 | 1.24 | -0.3 | 25.18 | -0.06 | 1.94 | 0.21 | 2.44 | 0.11 | -0.1 | 0.01 |
| Run1_MDS5 | | 0.41 | 0.26 | 0.3 | -0.05 | 1.06 | 0.7 | 1.04 | 0.96 | 12.11 | 0.72 | 0.12 | -0.01 | 0.37 | 1.44 | 3.28 | 2.13 | 0.81 | -0.09 | 7.44 | -0.02 | 0.6 | 1.23 | 2.35 | 1.46 | 0.52 | -0.09 |
| Run1_MDS6 | | 0.36 | -0.02 | -0.06 | -0.09 | 0.43 | 0.07 | 0.84 | 2.89 | 18.88 | 0.24 | 0.25 | 0.22 | 0.31 | 0.84 | 2.73 | 0.95 | 0.58 | -0.21 | 11 | -0.11 | 1.01 | 0.6 | 1.59 | 0.76 | -0.13 | -0.09 |
| Run1_MDS8 | | 0.4 | 0.85 | 1.21 | 0.4 | 1.25 | 0.57 | 1.33 | 4.04 | 23.24 | 0.27 | 0.09 | 0.16 | 0.33 | 1.12 | 2.11 | 1.53 | 0.59 | -0.14 | 15.24 | -0.18 | 1.58 | 2.15 | 1.91 | 0.65 | -0.2 | -0.17 |
| Run1_MDS9 | | 0.15 | 0.08 | 0.12 | 0.04 | 0.58 | 0.16 | 0.38 | 2.57 | 13.93 | -0.02 | -0.01 | 0.07 | 0.29 | 0.77 | 1.75 | 1.1 | 0.05 | -0.31 | 6.42 | -0.23 | 1.03 | 0.15 | 1.68 | 0.41 | -0.26 | -0.21 |
| Run2_MDS12 | Lower Risk | 0.3 | -0.21 | 0.17 | -0.12 | 0 | -0.1 | 0.94 | 1.87 | 13.91 | 0.07 | 0.12 | 0.2 | 0.33 | 0.88 | 1.86 | 1.18 | 0.77 | -0.33 | 6.8 | -0.21 | 0.51 | -0.03 | 2.22 | 0.56 | -0.2 | -0.21 |
| Run2_MDS14 | | 0.49 | 0.18 | -0.08 | -0.61 | 1.37 | 0.17 | 2 | 3.65 | 32.73 | 0.57 | 0.22 | 0.29 | 0.33 | 0.99 | 2.19 | 0.83 | 2.87 | -0.02 | 5.42 | -0.1 | 2.34 | 0.22 | 1.59 | 2.49 | -0.09 | -0.1 |
| Run2_MDS19 | | 0.37 | 0.05 | 0.04 | 0.06 | 0.48 | 0.09 | 1.5 | 3 | 20.95 | 0.41 | 0.41 | 0.31 | 0.31 | 1.13 | 2.19 | 0.93 | 0.66 | -0.2 | 7.79 | -0.19 | 1.1 | -0.11 | 1.55 | 0.33 | -0.23 | -0.18 |
| Run2_MDS20 | | 0.38 | -0.11 | 0.05 | -0.25 | 1.14 | -0.05 | 1.08 | 3.59 | 18.14 | 0.61 | 0.38 | 0.32 | 0.4 | 0.71 | 2.93 | 0.9 | 0.66 | -0.27 | 16.38 | -0.06 | 0.65 | -0.09 | 1.29 | 0.37 | -0.1 | -0.06 |
| Run2_MDS28* | | 0.48 | -0.01 | 0.08 | -0.01 | 0.31 | -0.15 | 0.42 | 1.78 | 25.54 | 0.32 | 0.24 | 0.4 | 0.48 | 1.11 | 2.18 | 0.81 | 1.27 | -0.24 | 6.74 | -0.18 | 0.9 | 1.14 | 1.09 | 0.74 | -0.21 | -0.15 |
| Run2_MDS5 | | 0.12 | 0.5 | -0.49 | 0.63 | 0.05 | 0.21 | 0.46 | 0.84 | 10.53 | 0.52 | 0.03 | -0.12 | 0.09 | 1.56 | 2.76 | 1.52 | 0.27 | -0.26 | 6.45 | -0.05 | 0.36 | 0.9 | 1.77 | 1.98 | 0.46 | -0.13 |
| Run1_ICUS18 | | 0.49 | 0.27 | -0.06 | 0.01 | 0.76 | -0.1 | 1.25 | 3.85 | 22.95 | 0.26 | 0.23 | 0.36 | 0.49 | 1.12 | 1.77 | 0.72 | 0.9 | -0.22 | 6.38 | -0.13 | 1.18 | 0.05 | 1.79 | 0.76 | -0.23 | -0.1 |
| Run1_ICUS22 | ICUS | 0.57 | 0.11 | 0.02 | 0.23 | 0.47 | 0.01 | 1.09 | 3.32 | 22.07 | 0.01 | 0.21 | 0.38 | 0.54 | 0.83 | 1.73 | 0.83 | 1.8 | -0.06 | 6.14 | -0.13 | 2.77 | 0.02 | 1.25 | 0.3 | -0.16 | -0.12 |
| Run1_ICUS24 | | 0.33 | 0.1 | 0.2 | 0.02 | 0.77 | 0.5 | 1.18 | 3.47 | 27.08 | -0.03 | 0.09 | 0.2 | 0.32 | 0.85 | 1.55 | 1.07 | 1.66 | -0.21 | 6.01 | -0.19 | 2 | 0.04 | 1.36 | 0.87 | -0.19 | -0.2 |
| Run1_Nl-1 | | 0.37 | 0.32 | -0.17 | 0.33 | 1.19 | 0.52 | 1.46 | 3.17 | 36.79 | -0.03 | -0.01 | 0.2 | 0.38 | 0.81 | 1.62 | 0.74 | 2.22 | 0.2 | 6.5 | -0.09 | 1.9 | 0.81 | 1.47 | 0.87 | -0.17 | -0.05 |
| Run1_Nl-3 | | 0.5 | 0.34 | 0.35 | 0.02 | 0.99 | 0.62 | 1.75 | 3.31 | 24.02 | 0.64 | 0.42 | 0.44 | 0.47 | 1.51 | 2.17 | 0.74 | 1.43 | -0.17 | 7.9 | -0.17 | 0.85 | 0.5 | 1.6 | 0.25 | -0.23 | -0.12 |
| Run1_Nl-4 | | 0.5 | 0.48 | 0.63 | 0.36 | 1.09 | 0.64 | 1.5 | 3.39 | 19.08 | 0.69 | 0.41 | 0.37 | 0.42 | 1.43 | 2.15 | 0.65 | 1.28 | -0.2 | 11.64 | -0.14 | 1.51 | 0.38 | 1.59 | 0.45 | -0.17 | -0.12 |
| Run1_Nl-6 | | 0.51 | 0.1 | -0.03 | 0.08 | 0.98 | 0.69 | 1.58 | 3.23 | 19.97 | 0.45 | 0.31 | 0.35 | 0.44 | 1.35 | 1.88 | 0.77 | 1.78 | -0.14 | 8.59 | -0.13 | 1.45 | 0.11 | 1.95 | 0.23 | -0.19 | -0.11 |
| Run2_Nl-3 | Normal | 0.42 | 0.09 | 0.11 | 0.07 | 0.68 | 0.35 | 1.46 | 3.15 | 18.43 | 0.8 | 0.4 | 0.37 | 0.44 | 1.01 | 1.97 | 0.53 | 1.08 | -0.16 | 7.43 | -0.18 | 0.65 | 0.15 | 1.19 | 0.33 | -0.23 | -0.14 |
| Run2_Nl-4 | | 0.46 | 0.44 | 0.46 | 0.32 | 0.66 | 0.38 | 1.48 | 3.04 | 15.28 | 0.9 | 0.45 | 0.36 | 0.41 | 1.16 | 1.98 | 0.52 | 1.35 | -0.21 | 11.67 | -0.18 | 1.2 | 0.16 | 1.2 | 0.36 | -0.2 | -0.16 |
| Run2_Nl-5 | | 0.57 | 0.27 | -0.2 | 0.47 | 0.91 | 0.62 | 1.43 | 3.14 | 15.03 | 0.92 | 0.5 | 0.42 | 0.41 | 1.23 | 2.39 | 0.67 | 2.11 | 0.02 | 9.94 | -0.1 | 1.09 | 0.32 | 1.13 | 0.17 | -0.15 | -0.06 |
| Run2_Nl-6 | | 0.47 | -0.02 | -0.05 | -0.02 | 0.8 | 0.37 | 1.38 | 2.95 | 17.14 | 0.66 | 0.49 | 0.39 | 0.42 | 1.09 | 1.8 | 0.72 | 1.45 | -0.15 | 10.82 | -0.15 | 1.28 | -0.02 | 1.5 | 0.14 | -0.19 | -0.13 |

**Supplemental Table 3**: Median expression of each surface marker on each cell population.

### Calreticulin Median

| | | Ungated | CD34+CD38low | HSCs | MPP | CMP/GMP | Myelo/Mono-Blasts | ProMonocytes | CD14neg Monocytes | Mature Monocytes | ProMyelocytes | Myelocytes | MetaMyelocytes | MatureGrans | ProErythroblasts | Erythroblasts | LateErythroblasts | PreBcells | Mature Bcells | Plasma Cells | T cells | NK cells | pDCs | Basophils | Platelets | CD8+ T cells | CD8neg T cells |
|---|---|---|---|---|---|---|---|---|---|---|---|---|---|---|---|---|---|---|---|---|---|---|---|---|---|---|---|
| Run1_MDS17 | | -0.1 | -0.21 | -0.22 | -0.24 | -0.17 | -0.21 | 0.36 | -0.01 | 0.14 | 1.31 | 1.74 | 0.41 | 0.64 | -0.35 | -0.38 | -0.2 | -0.18 | -0.47 | -0.43 | -0.4 | -0.32 | 10.19 | 0.21 | 0.19 | -0.4 | -0.4 |
| Run2_MDS15* | AML/RAEB-T | -0.27 | -0.34 | -0.34 | -0.36 | -0.35 | -0.3 | -0.3 | -0.37 | -0.37 | 0.1 | 0.12 | -0.02 | 0 | -0.41 | -0.26 | -0.18 | -0.32 | -0.42 | -0.3 | -0.37 | -0.34 | -0.23 | -0.28 | -0.19 | -0.36 | -0.38 |
| Run2_MDS4 | | -0.12 | -0.23 | -0.19 | -0.21 | -0.6 | -0.32 | -0.34 | -0.36 | -0.28 | 0.58 | 0.76 | 0.72 | 0.87 | -0.38 | -0.36 | -0.07 | -0.36 | -0.28 | -0.27 | -0.37 | -0.33 | -0.28 | -0.15 | 0.04 | -0.37 | -0.38 |
| Run1_MDS21 | | -0.04 | -0.1 | -0.23 | -0.13 | -0.14 | -0.12 | 0.4 | 0.04 | 0.28 | 1.33 | 2.02 | 1.11 | 1.37 | -0.38 | -0.3 | 1.19 | -0.4 | -0.45 | -0.55 | -0.41 | -0.42 | 7.12 | -0.21 | 0.49 | -0.38 | -0.42 |
| Run1_MDS3 | | 0.38 | 0.6 | 0.19 | 0.32 | 0.39 | 0.61 | 1.4 | 0.89 | 0.55 | 1.94 | 2.63 | 1.38 | 2.08 | -0.34 | -0.39 | 0.71 | -0.36 | -0.43 | -0.35 | -0.39 | -0.35 | 4.27 | -0.04 | 0.25 | -0.38 | -0.4 |
| Run2_MDS13 | | -0.12 | -0.23 | -0.26 | -0.23 | -0.25 | -0.24 | -0.26 | -0.3 | -0.23 | 0.95 | 1.35 | 0.57 | 0.66 | -0.36 | -0.37 | -0.17 | -0.35 | -0.36 | -0.27 | -0.38 | -0.4 | 0.07 | -0.21 | 0.14 | -0.37 | -0.38 |
| Run2_MDS16 | | 0.53 | -0.26 | -0.23 | -0.28 | -0.39 | -0.35 | -0.4 | -0.4 | -0.36 | 1.45 | 1.78 | 1.92 | 1.86 | -0.3 | -0.39 | 0.55 | -0.29 | -0.42 | -0.44 | -0.38 | -0.37 | -0.18 | -0.34 | 0.49 | -0.38 | -0.38 |
| Run2_MDS1 | | -0.27 | -0.36 | -0.4 | -0.44 | -0.37 | -0.36 | -0.37 | -0.38 | -0.36 | 0.07 | 0.05 | 0.26 | 0.57 | -0.33 | -0.41 | -0.32 | -0.26 | -0.42 | -0.33 | -0.39 | -0.39 | -0.36 | -0.47 | 0.23 | -0.39 | -0.39 |
| Run2_MDS21 | Higher Risk | -0.19 | -0.34 | -0.3 | -0.34 | -0.31 | -0.32 | -0.33 | -0.34 | -0.28 | 1.19 | 1.46 | 0.7 | 0.93 | -0.39 | -0.41 | -0.25 | -0.36 | -0.41 | -0.25 | -0.4 | -0.37 | 3.73 | -0.42 | 0.21 | -0.38 | -0.4 |
| Run2_MDS26* | | -0.05 | -0.31 | -0.22 | -0.35 | -0.31 | -0.31 | -0.24 | -0.38 | -0.34 | 0.12 | 0.22 | 0.53 | 0.61 | -0.32 | -0.4 | -0.37 | -0.3 | -0.38 | -0.54 | -0.4 | -0.34 | -0.12 | -0.38 | -0.05 | -0.39 | -0.4 |
| Run2_MDS27 | | -0.15 | -0.31 | -0.29 | -0.33 | -0.3 | -0.33 | -0.42 | -0.37 | -0.35 | 1.06 | 1.64 | 1.79 | 1.82 | -0.35 | -0.4 | -0.3 | -0.35 | -0.42 | -0.42 | -0.37 | -0.35 | -0.46 | -0.38 | 1.04 | -0.36 | -0.38 |
| Run2_MDS2 | | -0.23 | -0.37 | -0.26 | -0.4 | -0.35 | -0.35 | -0.43 | -0.39 | -0.43 | -0.03 | 0.12 | 0.03 | 0.46 | -0.26 | -0.15 | 0.12 | -0.34 | -0.28 | -0.37 | -0.39 | -0.37 | -0.14 | -0.32 | -0.1 | -0.35 | -0.4 |
| Run2_MDS3 | | -0.08 | -0.25 | -0.28 | -0.26 | -0.3 | -0.29 | -0.2 | -0.27 | -0.05 | 1.55 | 2.1 | 1.04 | 1.79 | -0.35 | -0.38 | -0.31 | -0.38 | -0.42 | -0.47 | -0.39 | -0.35 | -0.05 | -0.2 | 0.22 | -0.37 | -0.39 |
| Run1_MDS23 | | 0.59 | -0.32 | -0.24 | -0.33 | -0.4 | -0.27 | -0.18 | -0.31 | -0.33 | 1.73 | 1.76 | 1.24 | 0.99 | -0.4 | -0.35 | 0.06 | -0.25 | -0.44 | -0.47 | -0.42 | -0.37 | -0.41 | -0.45 | 0.36 | -0.43 | -0.42 |
| Run1_MDS25 | | 1.22 | 1.15 | -0.62 | 1.18 | -0.39 | -0.46 | -0.31 | -0.28 | -0.18 | 1.62 | 1.83 | 1.77 | 1.67 | -0.44 | -0.22 | 2.12 | -0.39 | -0.48 | -0.43 | -0.43 | -0.35 | -0.41 | -0.36 | -0.04 | -0.42 | -0.43 |
| Run1_MDS5 | | 0.1 | 0.19 | -0.52 | 0.35 | -0.16 | -0.31 | -0.39 | -0.06 | -0.07 | 1.75 | 0.63 | 0.24 | 0.36 | -0.08 | -0.19 | 0.63 | -0.24 | -0.37 | -0.45 | -0.37 | -0.34 | -0.2 | -0.26 | 0.4 | -0.4 | -0.36 |
| Run1_MDS6 | | 1.06 | -0.26 | -0.25 | -0.3 | -0.3 | -0.28 | -0.28 | -0.28 | -0.29 | 2.59 | 2.89 | 2.16 | 1.51 | -0.3 | -0.41 | 0.95 | -0.35 | -0.42 | -0.52 | -0.39 | -0.33 | -0.33 | -0.21 | 0.63 | -0.39 | -0.39 |
| Run1_MDS8 | | 1.33 | 0.09 | -0.35 | -0.19 | -0.36 | -0.34 | -0.25 | -0.36 | -0.26 | 2.19 | 2.67 | 2.14 | 2.16 | -0.29 | -0.29 | 0.81 | -0.35 | -0.39 | -0.41 | -0.38 | -0.33 | -0.04 | -0.12 | 0.9 | -0.37 | -0.38 |
| Run1_MDS9 | | 0.82 | -0.27 | -0.24 | -0.28 | -0.31 | -0.33 | -0.42 | -0.39 | -0.24 | 1.94 | 2.3 | 2.2 | 2.17 | -0.38 | -0.39 | 1.35 | -0.35 | -0.44 | -0.51 | -0.42 | -0.33 | -0.3 | -0.4 | 0.27 | -0.42 | -0.41 |
| Run2_MDS12 | Lower Risk | 0.3 | -0.41 | -0.38 | -0.41 | -0.38 | -0.41 | -0.37 | -0.36 | -0.35 | 0.53 | 0.6 | 0.78 | 0.84 | -0.37 | -0.4 | -0.22 | -0.36 | -0.39 | -0.38 | -0.38 | -0.34 | -0.38 | -0.33 | 0.14 | -0.37 | -0.38 |
| Run2_MDS14 | | 0.87 | -0.11 | -0.08 | -0.31 | -0.35 | -0.44 | -0.36 | -0.41 | -0.37 | 2.96 | 2.33 | 2.11 | 1.34 | -0.32 | -0.37 | 1.14 | -0.38 | -0.4 | -0.47 | -0.38 | -0.35 | -0.39 | -0.27 | 0.25 | -0.39 | -0.37 |
| Run2_MDS19 | | 1.6 | -0.34 | -0.36 | -0.32 | -0.3 | -0.4 | -0.38 | -0.37 | -0.35 | 1.76 | 2.48 | 2.63 | 2.38 | -0.35 | -0.38 | -0.19 | -0.39 | -0.41 | -0.52 | -0.39 | -0.36 | -0.37 | -0.34 | 0.52 | -0.38 | -0.39 |
| Run2_MDS20 | | 0.22 | -0.41 | -0.38 | -0.61 | -0.45 | -0.41 | -0.42 | -0.41 | -0.38 | 1.32 | 1.49 | 1.61 | 1.39 | -0.34 | -0.34 | 0.81 | -0.38 | -0.38 | -0.4 | -0.38 | -0.39 | -0.31 | -0.36 | 0.24 | -0.39 | -0.38 |
| Run2_MDS28* | | 1.06 | -0.25 | -0.19 | -0.29 | -0.37 | -0.38 | -0.38 | -0.38 | -0.38 | 1.42 | 1.78 | 1.93 | 1.6 | -0.26 | -0.32 | 0.92 | -0.38 | -0.41 | -0.4 | -0.4 | -0.36 | -0.37 | -0.31 | 0.74 | -0.4 | -0.4 |
| Run2_MDS5 | | -0.1 | -0.08 | -0.67 | -0.46 | -0.37 | -0.45 | -0.38 | -0.27 | -0.31 | 0.22 | 0.04 | -0.05 | 0.08 | -0.33 | -0.3 | 0.3 | -0.23 | -0.32 | -0.37 | -0.37 | -0.37 | -0.29 | -0.38 | 0.47 | -0.39 | -0.36 |
| Run1_ICUS18 | | 1.32 | -0.2 | -0.59 | -0.29 | -0.46 | -0.43 | -0.34 | -0.38 | -0.31 | 2.31 | 2.76 | 2.58 | 2.11 | -0.36 | -0.32 | 1.88 | -0.3 | -0.43 | -0.38 | -0.42 | -0.34 | -0.28 | -0.32 | 0.33 | -0.41 | -0.42 |
| Run1_ICUS22 | ICUS | 1.41 | -0.38 | -0.34 | -0.4 | -0.33 | -0.43 | -0.41 | -0.42 | -0.41 | 2.31 | 2.92 | 2.78 | 2.11 | -0.39 | -0.34 | 1.78 | -0.42 | -0.47 | -0.55 | -0.44 | -0.41 | -0.42 | -0.39 | 0.33 | -0.42 | -0.44 |
| Run1_ICUS24 | | 0.7 | -0.25 | -0.18 | -0.35 | -0.32 | -0.38 | -0.32 | -0.35 | -0.4 | 2.41 | 2.54 | 1.76 | 1.22 | -0.41 | -0.44 | 0.71 | -0.46 | -0.46 | -0.53 | -0.43 | -0.41 | -0.41 | -0.39 | 0.6 | -0.44 | -0.43 |
| Run1_Nl-1 | | 1.04 | -0.45 | -0.63 | -0.45 | -0.39 | -0.33 | -0.29 | -0.28 | -0.36 | 1.89 | 2.15 | 2.13 | 1.52 | -0.41 | -0.4 | 1.29 | -0.44 | -0.44 | -0.33 | -0.42 | -0.38 | -0.37 | -0.38 | 0.71 | -0.42 | -0.43 |
| Run1_Nl-3 | | 1.46 | -0.15 | -0.13 | -0.22 | -0.34 | -0.23 | -0.1 | -0.23 | -0.29 | 1.68 | 2.26 | 2.39 | 2.16 | -0.26 | -0.4 | 2.14 | -0.38 | -0.43 | -0.5 | -0.41 | -0.32 | -0.3 | -0.28 | 0.18 | -0.41 | -0.42 |
| Run1_Nl-4 | | 1.35 | -0.1 | -0.21 | -0.2 | -0.29 | -0.23 | -0.09 | -0.18 | -0.27 | 1.76 | 2.1 | 2.15 | 1.76 | -0.24 | -0.39 | 1.91 | -0.43 | -0.44 | -0.55 | -0.4 | -0.34 | -0.38 | -0.3 | -0.02 | -0.41 | -0.4 |
| Run1_Nl-6 | | 1.13 | -0.28 | -0.14 | -0.35 | -0.33 | -0.25 | -0.14 | -0.3 | -0.31 | 1.7 | 2.28 | 2.2 | 1.57 | -0.38 | -0.44 | 1.06 | -0.43 | -0.45 | -0.46 | -0.42 | -0.32 | -0.38 | -0.37 | -0.07 | -0.42 | -0.42 |
| Run2_Nl-3 | Normal | 1.32 | -0.19 | -0.09 | -0.23 | -0.32 | -0.36 | -0.3 | -0.33 | -0.34 | 1.1 | 2 | 2.21 | 2.29 | -0.32 | -0.36 | 1.73 | -0.37 | -0.39 | -0.3 | -0.38 | -0.36 | -0.34 | -0.29 | 0.24 | -0.39 | -0.38 |
| Run2_Nl-4 | | 1.18 | -0.22 | -0.24 | -0.28 | -0.33 | -0.36 | -0.28 | -0.33 | -0.32 | 1.12 | 1.64 | 1.93 | 1.84 | -0.32 | -0.36 | 1.65 | -0.36 | -0.39 | -0.47 | -0.38 | -0.34 | -0.39 | -0.29 | 0.34 | -0.38 | -0.39 |
| Run2_Nl-5 | | 0.86 | -0.18 | -0.18 | -0.24 | -0.31 | -0.35 | -0.32 | -0.33 | -0.32 | 1.1 | 2.1 | 2.37 | 2.02 | -0.3 | -0.4 | 1.24 | -0.4 | -0.39 | -0.4 | -0.38 | -0.32 | -0.33 | -0.29 | 0.22 | -0.38 | -0.38 |
| Run2_Nl-6 | | 1.07 | -0.3 | -0.3 | -0.28 | -0.33 | -0.37 | -0.32 | -0.38 | -0.35 | 0.87 | 1.52 | 1.88 | 1.73 | -0.38 | -0.39 | 0.36 | -0.4 | -0.41 | -0.4 | -0.4 | -0.35 | -0.37 | -0.34 | -0.09 | -0.39 | -0.4 |

**Supplemental Table 3**: Median expression of each surface marker on each cell population.

### CD321 Median

| | | Ungated | CD34+CD38low | HSCs | MPP | CMP/GMP | Myelo/Mono-Blasts | ProMonocytes | CD14neg Monocytes | Mature Monocytes | ProMyelocytes | Myelocytes | MetaMyelocytes | MatureGrans | ProErythroblasts | Erythroblasts | LateErythroblasts | PreBcells | Mature Bcells | Plasma Cells | T cells | NK cells | pDCs | Basophils | Platelets | CD8+ T cells | CD8neg T cells |
|---|---|---|---|---|---|---|---|---|---|---|---|---|---|---|---|---|---|---|---|---|---|---|---|---|---|---|---|
| Run1_MDS17 | | 14.31 | 78.2 | 80.63 | 75.27 | 144.25 | 57.47 | 32.46 | 28.55 | 31.71 | 35.39 | 16.36 | 4.07 | 4.5 | 14.9 | 13.97 | 10.3 | 16.39 | 2.08 | 1.17 | 6.66 | 9.12 | 23.3 | 43.69 | 500.51 | 6.3 | 7.28 |
| Run2_MDS15* | AML/RAEB-T | 28.17 | 52.28 | 67.44 | 71.18 | 80.76 | 38.43 | 30.95 | 19.67 | 32.98 | 41.82 | 28.88 | 17.49 | 13.25 | 12.5 | 25.93 | 38.77 | 28.19 | 2.76 | 0.53 | 7.49 | 9.34 | 71.01 | 66.76 | 342.98 | 6.31 | 8.86 |
| Run2_MDS4 | | 8.36 | 33.16 | 38.99 | 25.16 | 13.5 | 9.64 | 10.98 | 14.72 | 24.38 | 36.43 | 18.21 | 5.58 | 6.28 | 4.59 | 9.28 | 7.58 | 6.4 | 5.44 | 0.7 | 5.98 | 4.75 | 7.01 | 25.26 | 304.1 | 5.9 | 6.05 |
| Run1_MDS21 | | 15.19 | 78.81 | 77.95 | 78.71 | 112.32 | 97.99 | 43.66 | 31.55 | 35.88 | 21.2 | 15.26 | 3.26 | 3.71 | 15.14 | 17.68 | 1106.2 | 9.77 | 3.04 | 1.98 | 9.42 | 11.73 | 35.9 | 33.44 | 1402.16 | 10.52 | 8.92 |
| Run1_MDS3 | | 16.22 | 71.08 | 76.96 | 66.13 | 108.42 | 95.52 | 48.77 | 41.88 | 49.49 | 23.63 | 14.39 | 4.37 | 6.52 | 16.12 | 18.28 | 37.24 | 11.11 | 3.15 | 1.62 | 11.18 | 13.19 | 20.48 | 49.89 | 928.19 | 11.99 | 10.88 |
| Run2_MDS13 | | 16.06 | 59.31 | 61.78 | 56.64 | 88.14 | 75.31 | 30.47 | 38.47 | 46.44 | 34.74 | 22.67 | 5.78 | 5.35 | 7.18 | 10.18 | 6.35 | 9.36 | 1.87 | 0.22 | 8.17 | 8.54 | 7.44 | 43.76 | 519.95 | 8.67 | 7.92 |
| Run2_MDS16 | | 6.04 | 54.42 | 53.04 | 50 | 54.64 | 6.6 | 6.58 | 13.04 | 13.71 | 34.58 | 12.03 | 4.15 | 4.01 | 29.44 | 6.89 | 6.87 | 9.83 | 0.91 | 1.14 | 7.89 | 9.73 | 8.52 | 13.98 | 1310.27 | 6.63 | 8.23 |
| Run2_MDS1 | | 7.55 | 38.58 | 46.65 | 33.24 | 64.23 | 29.61 | 14.8 | 21.33 | 24.07 | 14.02 | 4.55 | 5.71 | 5.41 | 5.7 | 6.69 | 5.5 | 13.59 | 4.03 | 0.43 | 6.52 | 5.32 | 11.05 | 39.06 | 1056.48 | 6.76 | 6.48 |
| Run2_MDS21 | Higher Risk | 11.58 | 73.58 | 77.36 | 73.18 | 109.85 | 83.47 | 17.48 | 25.65 | 28.53 | 30.74 | 18.1 | 3.27 | 3.01 | 8.92 | 8.35 | 6.33 | 9.31 | 3.43 | 1.67 | 8.41 | 10.31 | 32.26 | 24.75 | 1163.03 | 9.45 | 7.98 |
| Run2_MDS26* | | 4.94 | 35.32 | 39.94 | 33.91 | 30.33 | 10.01 | 11.53 | 22 | 12.88 | 13.08 | 10.38 | 4.22 | 4.2 | 9.02 | 8.68 | 4.14 | 4.98 | 1.34 | 3.64 | 6.52 | 6.87 | 10.87 | 16.82 | 48.36 | 6.29 | 6.69 |
| Run2_MDS27 | | 9.02 | 39.46 | 43.41 | 35.21 | 34.81 | 17.38 | 7.03 | 11.6 | 11.91 | 35.13 | 14.61 | 3.64 | 3.42 | 21.42 | 7.64 | 3.3 | 13.52 | 4.41 | 0.53 | 12.22 | 11.7 | 3.94 | 17.15 | 855.87 | 11.25 | 13.68 |
| Run2_MDS2 | | 5.05 | 31.84 | 40.51 | 30.04 | 40.5 | 30.75 | 10.54 | 10.25 | 24.72 | 37.78 | 6.1 | 1.92 | 2.9 | 13.71 | 13.2 | 23.09 | 12.03 | 17.57 | 3.01 | 6.72 | 10.16 | 22.3 | 76.55 | 844.7 | 5.8 | 6.8 |
| Run2_MDS3 | | 16.93 | 63.43 | 67.65 | 59.95 | 103.66 | 82.32 | 29.97 | 41.09 | 49.17 | 42.65 | 23.82 | 5.75 | 5.74 | 13.93 | 11.48 | 5.65 | 11.61 | 3.66 | 0.76 | 10.44 | 12.28 | 27.5 | 49.95 | 816.79 | 11.58 | 9.92 |
| Run1_MDS23 | | 7.73 | 20.82 | 28.06 | 18.51 | 21.75 | 4.69 | 7.2 | 10.31 | 13.6 | 30.58 | 17.32 | 6.17 | 5.43 | 10.98 | 14.97 | 13.49 | 10 | 3.77 | 0.57 | 7.85 | 8.68 | 5.82 | 26.42 | 414.43 | 7.58 | 7.96 |
| Run1_MDS25 | | 4.31 | 21.37 | 43.99 | 9.43 | 11.01 | 4.62 | 7.85 | 15.08 | 17.43 | 29.82 | 14.27 | 3.83 | 3.55 | 12.64 | 18.56 | 11.2 | 9.2 | 1.23 | 0.97 | 5.45 | 12.24 | 13 | 17.5 | 785.4 | 4.9 | 6.15 |
| Run1_MDS5 | | 9.33 | 47.2 | 49.09 | 44.1 | 22.39 | 5.95 | 7.08 | 17.92 | 15.53 | 55.13 | 10.43 | 8.47 | 7.88 | 6.07 | 15.56 | 186.94 | 9.33 | 4.07 | -0.08 | 6.88 | 12.44 | 16.26 | 22.99 | 398.66 | 4.12 | 8.19 |
| Run1_MDS6 | | 9.25 | 27.32 | 25.82 | 21.52 | 22.88 | 19.63 | 23.44 | 37.64 | 45.11 | 39.07 | 19.18 | 7.56 | 7.53 | 6.8 | 15.04 | 13.66 | 7.01 | 2.17 | 0.78 | 8.25 | 11.08 | 21.31 | 28 | 1330.74 | 6.76 | 9.72 |
| Run1_MDS8 | | 11.11 | 34.22 | 29.44 | 31.31 | 17.03 | 12.32 | 16.48 | 23.74 | 29.85 | 27.58 | 17.19 | 9.01 | 10.6 | 13.21 | 13.76 | 13.95 | 8.9 | 1.82 | 0.82 | 7.67 | 10.01 | 17.9 | 29.53 | 1405.68 | 8 | 7.58 |
| Run1_MDS9 | | 8.57 | 28.09 | 30.01 | 24.68 | 28.7 | 20.48 | 22.04 | 37.82 | 40.62 | 26.43 | 15.62 | 6.12 | 6.17 | 12.15 | 14.32 | 76.03 | 10.03 | 2.56 | 0.87 | 7.51 | 10.78 | 14.85 | 18.8 | 604.34 | 7.34 | 7.61 |
| Run2_MDS12 | Lower Risk | 6.15 | 7.71 | 37.24 | 18.34 | 42.05 | 3.32 | 7.05 | 12.12 | 16.33 | 22.16 | 21.64 | 6.24 | 4.07 | 3.45 | 4.11 | 3.24 | 8.22 | 2.4 | 1.43 | 5.18 | 6.73 | 4.91 | 21.27 | 700.45 | 5.21 | 5.16 |
| Run2_MDS14 | | 4.96 | 30.52 | 41.64 | 21.28 | 8.41 | 4.26 | 6.09 | 10.55 | 13.24 | 14.12 | 8.99 | 4.42 | 4.34 | 3.47 | 5.21 | 5.66 | 3.71 | 1.71 | 0.07 | 9.31 | 4.79 | 19.24 | 19.57 | 1288.45 | 7.14 | 10.61 |
| Run2_MDS19 | | 5.67 | 47.35 | 55.26 | 47.32 | 23.76 | 9.65 | 20.91 | 33.87 | 33.82 | 20.36 | 10.71 | 3.72 | 4.25 | 3.86 | 5.32 | 2.69 | 17.24 | 3.23 | 1.52 | 13.42 | 9.92 | 9.91 | 20.71 | 1353.16 | 9.38 | 15.22 |
| Run2_MDS20 | | 6.27 | 28.05 | 28.86 | 35.02 | 20.04 | 4.22 | 3.84 | 8.03 | 12.29 | 31 | 19.71 | 5.37 | 4.73 | 3.04 | 8.28 | 6.62 | 9.89 | 5.8 | 0.59 | 7.79 | 3.83 | 10.04 | 13.98 | 1064.43 | 7.84 | 7.79 |
| Run2_MDS28* | | 7.65 | 26.12 | 42.27 | 25.84 | 33.17 | 4.73 | 7.31 | 9.81 | 8.95 | 36.23 | 19.54 | 6.37 | 6.13 | 12.33 | 10.07 | 5.94 | 11.44 | 3.01 | 1.75 | 8.05 | 7.26 | 11.09 | 34.37 | 1404.74 | 7.27 | 8.77 |
| Run2_MDS5 | | 6.27 | 37.84 | 94.19 | 38.83 | 37.65 | 3.86 | 4.44 | 15.24 | 12.81 | 44.66 | 5.26 | 5.29 | 5.79 | 2.09 | 11.29 | 30.74 | 8.09 | 3.03 | 0.33 | 5.97 | 10.64 | 16.94 | 23.93 | 564.48 | 3.61 | 6.94 |
| Run1_ICUS18 | | 5.07 | 42.12 | 37.87 | 41.11 | 35.87 | 8.94 | 8.92 | 14.07 | 19.64 | 29.78 | 17.74 | 4.37 | 3.83 | 12.31 | 14.61 | 4.33 | 11 | 2.95 | 2.14 | 9.17 | 10.31 | 13.29 | 16.75 | 1054.15 | 8.14 | 9.39 |
| Run1_ICUS22 | ICUS | 4.22 | 39.41 | 42.26 | 36.84 | 29.48 | 3.69 | 5.17 | 7.57 | 9.57 | 26.03 | 16.19 | 3.4 | 3.11 | 12.66 | 17.78 | 4.74 | 4.97 | 1.03 | 0.22 | 7.94 | 8.4 | 5.25 | 13.29 | 806.37 | 7.31 | 8.21 |
| Run1_ICUS24 | | 5.14 | 31.07 | 32.76 | 30 | 13.68 | 6.71 | 8.93 | 11.81 | 12.2 | 23.41 | 14.98 | 4.74 | 3.79 | 8.55 | 10.86 | 38.64 | 3.17 | 0.71 | 0.43 | 3.3 | 3.87 | 5.97 | 16.52 | 1522.99 | 2.79 | 3.74 |
| Run1_Nl-1 | | 5.72 | 30.45 | 48.48 | 26.57 | 12.11 | 6.66 | 9.7 | 15.01 | 15.84 | 29.77 | 16.08 | 5.35 | 4.2 | 11.28 | 15.77 | 8.09 | 2.41 | 1.35 | 0.78 | 6.81 | 8.33 | 12.98 | 25.03 | 1793.27 | 5.93 | 7.37 |
| Run1_Nl-3 | | 8.94 | 46.29 | 43.36 | 48.24 | 26.63 | 10.34 | 13.61 | 19.56 | 20.14 | 29.17 | 13.99 | 6.61 | 7.61 | 14.77 | 11.66 | 8.67 | 5.65 | 1.78 | 0.41 | 8.15 | 9.16 | 9.05 | 15.95 | 599.99 | 7.05 | 9 |
| Run1_Nl-4 | | 9.77 | 32.54 | 41.2 | 27.96 | 14.95 | 7.88 | 13.17 | 19.89 | 20.37 | 34.99 | 20.03 | 7.62 | 8.55 | 15.41 | 14.25 | 9.26 | 3.9 | 1.03 | 0.43 | 5.72 | 5.48 | 5.79 | 17.17 | 510.7 | 4.91 | 6.51 |
| Run1_Nl-6 | | 8.71 | 40.66 | 43.99 | 35.49 | 20.92 | 11.84 | 14.67 | 17.83 | 17.56 | 32.48 | 16.46 | 6.7 | 6.98 | 11.69 | 12.22 | 7.46 | 3.34 | 1.09 | 0.6 | 8.49 | 8.94 | 9.8 | 18.02 | 443.56 | 7.32 | 8.98 |
| Run2_Nl-3 | Normal | 7.17 | 42.82 | 47.5 | 38.9 | 20.38 | 7.45 | 12.52 | 18.09 | 18.07 | 26.03 | 10.37 | 5.04 | 6.28 | 9.97 | 7.92 | 5.5 | 4.48 | 1.3 | 1.03 | 6.91 | 7.92 | 7.63 | 15.17 | 838.68 | 6.04 | 7.6 |
| Run2_Nl-4 | | 8.1 | 29.27 | 36.57 | 27.27 | 13.3 | 5.66 | 10.92 | 18.78 | 18.82 | 34.21 | 15.59 | 6.18 | 7.65 | 9.41 | 9.44 | 6.34 | 3.69 | 0.76 | 0.44 | 4.76 | 4.82 | 5.12 | 16.79 | 746.82 | 4.05 | 5.41 |
| Run2_Nl-5 | | 8.25 | 31.61 | 35.91 | 25.44 | 14.61 | 7.01 | 11.11 | 18.14 | 20.1 | 27.5 | 12.15 | 5.09 | 7.72 | 10.61 | 5.71 | 5.8 | 2.63 | 1.62 | 0.81 | 9.09 | 11.28 | 8.28 | 20 | 842.28 | 8.34 | 9.97 |
| Run2_Nl-6 | | 6.68 | 38.27 | 39.07 | 35.69 | 23.56 | 8.4 | 10.95 | 15.21 | 14.99 | 28.33 | 14.84 | 5.33 | 5.62 | 5.34 | 7.05 | 4.4 | 2.6 | 0.85 | 0.49 | 7.41 | 8.36 | 8.83 | 17.1 | 802.84 | 6.33 | 7.83 |

**Supplemental Table 3**: Median expression of each surface marker on each cell population.

### CD99 Median

| Sample | Group | Ungated | CD34+CD38low | HSCs | MPP | CMP/GMP | Myelo/Mono-Blasts | ProMonocytes | CD14neg Monocytes | Mature Monocytes | ProMyelocytes | Myelocytes | MetaMyelocytes | MatureGrans | ProErythroblasts | Erythroblasts | LateErythroblasts | PreBcells | Mature Bcells | Plasma Cells | T cells | NK cells | pDCs | Basophils | Platelets | CD8+ T cells | CD8neg T cells |
|---|---|---|---|---|---|---|---|---|---|---|---|---|---|---|---|---|---|---|---|---|---|---|---|---|---|---|---|
| Run1_MDS17 | | 8.2 | 135.34 | 147.58 | 129.14 | 237.58 | 67.44 | 28.09 | 25.11 | 37.29 | 3.79 | 3.52 | 2.23 | 2.61 | 10.47 | 5.92 | 5.12 | 51.24 | 22.17 | 7.91 | 126.84 | 113.41 | 23.23 | 37.03 | 57.14 | 130.83 | 120.19 |
| Run2_MDS15* | AML/RAEB-T | 13.05 | 156.02 | 228.89 | 220.04 | 190.6 | 31.49 | 13.56 | 22.52 | 71.06 | 8.67 | 5.64 | 3.55 | 2.46 | 71 | 13.35 | 16.79 | 21.91 | 30.93 | 76.67 | 182.69 | 147.09 | 103.57 | 37.16 | 229.61 | 178.73 | 185.37 |
| Run2_MDS4 | | 5.36 | 312.15 | 257.3 | 356.98 | 86.89 | 36.8 | 59.96 | 32.33 | 34.72 | 3.18 | 2.2 | 0.48 | 0.47 | 2.59 | 1.39 | 2.63 | 45.93 | 75.88 | 7.4 | 179.08 | 165.56 | 130.48 | 26.97 | 84.77 | 218.74 | 138.05 |
| Run1_MDS21 | | 12.44 | 115.12 | 133.08 | 109.18 | 163.17 | 131.68 | 43.08 | 30.02 | 31.44 | 1.73 | 1.64 | 0.66 | 0.92 | 7.95 | 9.24 | 21.61 | 19.41 | 14.49 | 9.02 | 45.19 | 45.7 | 47.62 | 26.63 | 20.44 | 61.38 | 39.18 |
| Run1_MDS3 | | 9.22 | 156.72 | 166.3 | 147.35 | 227.34 | 174.64 | 52.29 | 30.07 | 44.34 | 1.79 | 1.39 | 0.79 | 1.44 | 11.44 | 11.25 | 11.88 | 25.75 | 21.43 | 5.61 | 81.32 | 56.17 | 67.22 | 15.7 | 25.45 | 95.88 | 74.51 |
| Run2_MDS13 | | 11.55 | 213.96 | 243.33 | 192.49 | 232.95 | 168.55 | 24.06 | 36.23 | 50.08 | 4.14 | 3.64 | 2.06 | 2.25 | 24.18 | 5.6 | 4.15 | 33.22 | 21.14 | 23.24 | 81.08 | 67.03 | 150.11 | 13.2 | 21.08 | 102.17 | 71.12 |
| Run2_MDS16 | | 2.15 | 195.77 | 181.43 | 172.41 | 284.64 | 51.38 | 21.82 | 47.57 | 59.76 | 1.5 | 1 | 0.24 | 0.33 | 101.62 | 30.69 | 6.82 | 90.8 | 20.76 | 18.46 | 87.95 | 178.15 | 174.56 | 12.63 | 239.68 | 89.36 | 87.63 |
| Run2_MDS1 | | 15.96 | 59.98 | 51.63 | 55.75 | 77.54 | 49.62 | 56.06 | 82.27 | 69.44 | 1.14 | 0.39 | 0.23 | 0.28 | 12.33 | 3.75 | 2.8 | 55.2 | 28.22 | 3.19 | 58.64 | 93.99 | 56.18 | 9.35 | 282.65 | 94.83 | 51.66 |
| Run2_MDS21 | Higher Risk | 8.3 | 120.87 | 135.68 | 117.86 | 168.77 | 113.51 | 7.71 | 12.17 | 16.66 | 1.65 | 1.51 | 0.52 | 0.66 | 10.79 | 4.52 | 4.68 | 21.03 | 15.27 | 3.54 | 45.49 | 45.01 | 71.46 | 18.95 | 25.12 | 63.82 | 38.95 |
| Run2_MDS26* | | 0.45 | 48.59 | 54.4 | 45.85 | 67.79 | 11.88 | 6.68 | 5.78 | 7.84 | 0.72 | 0.69 | 0 | -0.03 | 11.49 | 2.56 | 1.31 | 3.41 | 5 | 2.48 | 24.34 | 35.35 | 15.06 | 5.88 | 2.09 | 39.71 | 17.63 |
| Run2_MDS27 | | 33.67 | 98.94 | 94.53 | 93.45 | 96.24 | 59.65 | 40.05 | 15.78 | 21.8 | 1.75 | 1.31 | 0.21 | 0.15 | 33.47 | 33.41 | 10.92 | 144.56 | 37.95 | 10.54 | 214.05 | 204.11 | 6.03 | 19.85 | 321.73 | 232.22 | 190.22 |
| Run2_MDS2 | | 5.24 | 457.97 | 526.81 | 471.86 | 437.85 | 224.49 | 80.18 | 11.33 | 24 | 12.1 | 0.74 | 0.46 | 0.7 | 44.39 | 11.64 | 52.09 | 127.15 | 75.35 | 3.59 | 109.91 | 153.66 | 42.32 | 60.84 | 126.84 | 162.62 | 106.5 |
| Run2_MDS3 | | 7.48 | 173.37 | 209.44 | 153.96 | 237.67 | 141.17 | 13.52 | 13.05 | 30.51 | 2.53 | 2.01 | 1.05 | 1.46 | 37.7 | 7.56 | 3.29 | 31.98 | 21.02 | 8.28 | 87.54 | 73.54 | 53.98 | 13.39 | 28.99 | 109.76 | 78.52 |
| Run1_MDS23 | | 0.67 | 5.3 | 7.15 | 4.95 | 6.18 | 1.61 | 2.23 | 1.44 | 2.05 | 1.3 | 0.87 | 0.1 | -0.05 | 2.43 | 2.92 | 2.34 | 3.7 | 5.27 | 1.31 | 22.18 | 6.31 | 6.55 | 1.85 | 22.23 | 29.38 | 19.17 |
| Run1_MDS25 | | 0.55 | 27.29 | 82.76 | 14.89 | 61.44 | 12.65 | 40.79 | 28.64 | 15.25 | 1.71 | 1.07 | 0.08 | 0.09 | 6.83 | 10.19 | 2.89 | 38.22 | 9.42 | 3.77 | 99.52 | 58.9 | 79.1 | 7.31 | 44.66 | 106.39 | 92.27 |
| Run1_MDS5 | | 2.4 | 140.81 | 196.6 | 148.48 | 127.67 | 58.12 | 24.81 | 12 | 26.28 | 2.66 | 1.75 | 1.11 | 1.36 | 5.58 | 15.37 | 32.3 | 11.6 | 21.99 | 6.54 | 123.5 | 240.12 | 23.02 | 19.99 | 50.77 | 178.09 | 97.62 |
| Run1_MDS6 | | 1.49 | 103.99 | 82.8 | 55.53 | 94.81 | 35.14 | 25.58 | 27.17 | 24.84 | 2.34 | 1.75 | 0.7 | 0.62 | 24.5 | 18.99 | 4.75 | 24.37 | 17.96 | 5.35 | 84.8 | 108.58 | 67.98 | 11.74 | 180.97 | 100.53 | 73.54 |
| Run1_MDS8 | | 1.66 | 83.3 | 98.54 | 69.5 | 136.09 | 32.67 | 14.38 | 51.24 | 49.59 | 2.18 | 1.71 | 0.64 | 0.65 | 43.1 | 36.36 | 18.57 | 35.8 | 22.71 | 4.04 | 41.77 | 107.22 | 43.5 | 16.47 | 167.16 | 51.57 | 39.69 |
| Run1_MDS9 | | 1.88 | 115.13 | 133.13 | 84.43 | 134.85 | 48.82 | 7.57 | 19.4 | 24.64 | 1.52 | 1.21 | 0.23 | 0.16 | 9.41 | 8.2 | 25.33 | 52.91 | 13.58 | 7.7 | 53 | 80.54 | 9.82 | 10.43 | 49.95 | 82.75 | 39.26 |
| Run2_MDS12 | Lower Risk | 0.84 | 15.76 | 58.17 | 22.26 | 56.3 | 15.36 | 28.65 | 28.93 | 40.16 | 0.98 | 1.37 | 0.33 | 0.1 | 9.49 | 9.76 | 3.66 | 22.94 | 14.21 | 5.57 | 37.31 | 60.57 | 43.97 | 10.39 | 65.02 | 58.63 | 27.91 |
| Run2_MDS14 | | 0.54 | 30.89 | 22.8 | 53.25 | 56.16 | 5.88 | 6.46 | 9.74 | 12.87 | 0.52 | 0.49 | 0.11 | 0.08 | 4.21 | 3.58 | 0.86 | 5.23 | 11.15 | 1.64 | 75.21 | 30.33 | 119.43 | 5.79 | 73.19 | 70.46 | 77.56 |
| Run2_MDS19 | | 0.57 | 62.84 | 64.46 | 60.61 | 69.38 | 12.99 | 17.77 | 13.35 | 17.51 | 0.98 | 0.71 | 0 | -0.01 | 8.66 | 9.26 | 3.17 | 44.11 | 16.96 | 5.34 | 80.88 | 62.17 | 115.64 | 9.49 | 83.94 | 96.83 | 74.89 |
| Run2_MDS20 | | 1.96 | 63.88 | 70.14 | 55.43 | 102.35 | 6.18 | 7.59 | 12.99 | 19.78 | 1.51 | 1.17 | 0.13 | 0.05 | 18.87 | 8.53 | 1.04 | 59 | 32.03 | 12.73 | 72.98 | 41.24 | 7.68 | 6.63 | 109.27 | 138.95 | 67.92 |
| Run2_MDS28* | | 0.42 | 23.2 | 31.19 | 18.02 | 38.55 | 1.79 | 3.07 | 2.06 | 4.81 | 1.08 | 0.82 | -0.01 | -0.07 | 24.35 | 7.9 | 0.81 | 10.59 | 7.48 | 5.13 | 38.87 | 24.18 | 7.14 | 5.96 | 90.77 | 48.37 | 31.81 |
| Run2_MDS5 | | 1.81 | 124.29 | 284.12 | 161.92 | 147.83 | 58.81 | 19.51 | 11.88 | 26.62 | 2.05 | 1.28 | 0.87 | 1.18 | 11.57 | 14.09 | 9.02 | 12.63 | 23.15 | 8.71 | 144.14 | 300.41 | 52.38 | 22.27 | 77.18 | 202.42 | 115.17 |
| Run1_ICUS18 | | 0.58 | 55.6 | 44.61 | 46.06 | 72.86 | 7.66 | 10.99 | 23.16 | 18.25 | 1.47 | 1.14 | 0.04 | -0.07 | 19.09 | 18.1 | 0.13 | 49.14 | 15.4 | 2.78 | 58.77 | 73.97 | 110.68 | 6.31 | 54.83 | 88.46 | 54 |
| Run1_ICUS22 | ICUS | 0.27 | 13.8 | 14.09 | 12.23 | 17.37 | 2.06 | 2.77 | 2.93 | 3.84 | 0.98 | 0.84 | -0.06 | -0.19 | 3.09 | 3.99 | 0.44 | 4.16 | 3.13 | 1.5 | 22.21 | 12.6 | 2.5 | 2.76 | 25.34 | 25.5 | 20.92 |
| Run1_ICUS24 | | 0.85 | 22.59 | 18.67 | 22.44 | 27.05 | 6.38 | 7.67 | 4.83 | 5.47 | 1.05 | 0.9 | 0.16 | -0.05 | 4.16 | 3.88 | 9.38 | 2.66 | 3.84 | 1.04 | 27.65 | 14.58 | 22.96 | 4.83 | 34.7 | 26.25 | 29.06 |
| Run1_Nl-1 | | 0.69 | 37.03 | 36.1 | 32.3 | 63.29 | 8.86 | 10.91 | 12.68 | 16.66 | 1.45 | 1.03 | 0.2 | -0.05 | 7.59 | 8.82 | 1.15 | 3.71 | 5.12 | 3.13 | 22.98 | 34.74 | 62.13 | 4.88 | 75.5 | 22.38 | 23.31 |
| Run1_Nl-3 | | 1.76 | 105.16 | 92.6 | 89.13 | 116.12 | 30.65 | 34.11 | 31.18 | 30.59 | 2.42 | 1.79 | 0.74 | 0.72 | 33.62 | 9.82 | 1.14 | 14.35 | 11.58 | 4.84 | 56.11 | 93.3 | 86.88 | 7.71 | 46.07 | 58.85 | 54.53 |
| Run1_Nl-4 | | 2.11 | 90.51 | 148.13 | 81.81 | 134.54 | 34.86 | 41.54 | 35.74 | 38.38 | 3.55 | 2.66 | 1.02 | 1.03 | 42.84 | 10.48 | 1.37 | 14.55 | 18.9 | 6.44 | 75.06 | 78.1 | 94.94 | 16.74 | 67.49 | 63.81 | 84.07 |
| Run1_Nl-6 | | 1.11 | 39.04 | 39.5 | 34.5 | 64.42 | 21.23 | 26.71 | 21.58 | 26.97 | 2.18 | 1.44 | 0.41 | 0.35 | 10.26 | 6.26 | 1.1 | 6.01 | 8.04 | 1.86 | 42.44 | 57.78 | 40.95 | 5.71 | 32.81 | 53.55 | 38.41 |
| Run2_Nl-3 | Normal | 1.31 | 99.34 | 110.88 | 91.62 | 104.28 | 25.93 | 31.14 | 24.25 | 31.46 | 1.97 | 1.18 | 0.46 | 0.54 | 27.38 | 7.94 | 0.69 | 12.62 | 12.05 | 6.02 | 61.3 | 107.4 | 105.7 | 8.08 | 65.02 | 64.49 | 59.6 |
| Run2_Nl-4 | | 1.75 | 115.66 | 153.56 | 92.23 | 132.79 | 25.76 | 36.3 | 28.62 | 40.21 | 3.05 | 1.99 | 0.77 | 0.93 | 33.8 | 7.72 | 0.94 | 13.5 | 19.42 | 6.32 | 83.72 | 88.69 | 113.58 | 18.13 | 91.83 | 72.78 | 91.98 |
| Run2_Nl-5 | | 1.75 | 138.47 | 160.69 | 113.96 | 184.9 | 37.09 | 55.61 | 49.36 | 51.38 | 2.08 | 1.15 | 0.4 | 0.63 | 18.31 | 5.96 | 0.85 | 9.51 | 21.21 | 7.48 | 71.82 | 169.34 | 140.03 | 10.78 | 56.81 | 63.67 | 78.41 |
| Run2_Nl-6 | | 0.69 | 31.86 | 34.3 | 29.5 | 55.19 | 10.8 | 16.9 | 13.23 | 20.35 | 1.65 | 1.1 | 0.21 | 0.18 | 5.56 | 3.54 | 0.77 | 4.79 | 7.2 | 1.58 | 38.02 | 58.01 | 39.82 | 4.43 | 50.77 | 54.05 | 33.61 |

**Supplemental Table 3**: Median expression of each surface marker on each cell population.

### CD13 Median

| Sample | Group | Ungated | CD34+CD38low | HSCs | MPP | CMP/GMP | Myelo/Mono-Blasts | ProMonocytes | CD14neg Monocytes | Mature Monocytes | ProMyelocytes | Myelocytes | MetaMyelocytes | MatureGrans | ProErythroblasts | Erythroblasts | LateErythroblasts | PreBcells | Mature Bcells | Plasma Cells | T cells | NK cells | pDCs | Basophils | Platelets | CD8+ T cells | CD8neg T cells |
|---|---|---|---|---|---|---|---|---|---|---|---|---|---|---|---|---|---|---|---|---|---|---|---|---|---|---|---|
| Run1_MDS17 | | 1.53 | 3.82 | 4.57 | 3.6 | 4.77 | 2.72 | 3.22 | 2.8 | 7.17 | 0.55 | 0.43 | 1.16 | 2.39 | 0.26 | -0.1 | 0.01 | 2.97 | 0.83 | -0.19 | 3.02 | 2.54 | 1.87 | 9.53 | 1.75 | 2.93 | 3.14 |
| Run2_MDS15* | AML/RAEB-T | 3.03 | 4.99 | 5.73 | 4.44 | 3.59 | 3.51 | 2.43 | 4.13 | 7.04 | 3.48 | 1.32 | 1.56 | 2.57 | 0.88 | 1.65 | 3.86 | 6.77 | 0.62 | -0.12 | 2.6 | 2.1 | 4.58 | 5.6 | 3.88 | 2.49 | 2.74 |
| Run2_MDS4 | | 0.87 | 4.3 | 3.99 | 3.26 | 1.25 | 1.19 | 1.34 | 1.22 | 2.4 | 0.22 | -0.06 | 0.14 | 0.57 | -0.22 | -0.24 | -0.08 | 1.37 | 1.55 | -0.21 | 2.68 | 1.78 | 2.13 | 10.12 | 1.76 | 2.96 | 2.36 |
| Run1_MDS21 | | 1.4 | 7.82 | 8.79 | 7.43 | 7.71 | 6.77 | 5.57 | 4.4 | 6.65 | -0.09 | 0.09 | 0.93 | 1.79 | 0.04 | 0.33 | 1.16 | 1.77 | 1.01 | -0.07 | 2.66 | 2.1 | 3.24 | 10.86 | 0.99 | 2.8 | 2.61 |
| Run1_MDS3 | | 1.07 | 7.11 | 7.84 | 6.55 | 6.23 | 5.73 | 4.86 | 4.47 | 9.39 | -0.02 | -0.02 | 0.75 | 1.73 | 0.22 | 0.4 | 0.76 | 1.84 | 1 | -0.02 | 2.8 | 1.81 | 2.51 | 14.81 | 1.01 | 2.93 | 2.73 |
| Run2_MDS13 | | 1.2 | 5.41 | 6.56 | 5.02 | 5.26 | 4.15 | 2.05 | 2.49 | 4.55 | 0.35 | 0.31 | 0.84 | 1.65 | 0.35 | -0.11 | -0.04 | 1.69 | 0.86 | -0.01 | 2.15 | 1.48 | 3.32 | 10.12 | 0.81 | 2.3 | 2.09 |
| Run2_MDS16 | | 0.64 | 5.14 | 5.51 | 4.14 | 5.12 | 3.99 | 6.07 | 7.54 | 13.34 | -0.1 | -0.17 | -0.14 | 0.31 | 1.79 | 0.47 | 0.56 | 3.48 | 0.33 | -0.17 | 1.99 | 2.08 | 2.87 | 6.13 | 1.76 | 1.94 | 2 |
| Run2_MDS1 | | 1.78 | 8 | 6.49 | 7.83 | 9.35 | 4.41 | 4.96 | 13.5 | 23.97 | -0.03 | 0.27 | 1.86 | 2.62 | 0.29 | -0.09 | -0.04 | 3.45 | 1 | -0.25 | 1.56 | 1.39 | 1.38 | 8.64 | 1.85 | 1.87 | 1.49 |
| Run2_MDS21 | Higher Risk | 0.81 | 6.74 | 8.17 | 6.34 | 6.77 | 5.07 | 1.44 | 1.77 | 3.95 | -0.09 | -0.08 | 0.32 | 0.92 | 0 | -0.19 | -0.14 | 1.29 | 0.62 | -0.23 | 1.93 | 1.25 | -0.08 | 10.24 | 0.72 | 2.07 | 1.89 |
| Run2_MDS26* | | 0.1 | 4.75 | 5.1 | 4.41 | 5.18 | 3.12 | 2.19 | 5.73 | 6.08 | -0.09 | -0.05 | 0.29 | 0.73 | 1.01 | -0.22 | -0.27 | 1.3 | 0.07 | 0.27 | 1.39 | 0.95 | 2.32 | 2.78 | 0.24 | 1.66 | 1.22 |
| Run2_MDS27 | | 0.9 | 3 | 3.75 | 2.27 | 2.36 | 1.54 | 2 | 2.7 | 6.11 | 0.06 | -0.17 | -0.22 | 0.08 | 0.7 | 0.4 | 0.15 | 3.58 | 0.85 | -0.27 | 3.04 | 2.58 | 2.41 | 4.48 | 1.25 | 3.17 | 2.89 |
| Run2_MDS2 | | 1.62 | 3.61 | 5.46 | 2.85 | 3.95 | 2.28 | 1.18 | 1.41 | 3.98 | 2.04 | 0.47 | 1.13 | 1.61 | 2.01 | 1.6 | 1.61 | 2.69 | 1.26 | -0.63 | 2.2 | 1.82 | 1.57 | 3.97 | 1.77 | 2.51 | 2.18 |
| Run2_MDS3 | | 0.73 | 5.22 | 5.62 | 4.94 | 5.21 | 3.79 | 1.47 | 1.53 | 5.71 | 0.13 | 0.06 | 0.25 | 0.97 | 0.62 | -0.06 | -0.19 | 1.33 | 0.82 | -0.3 | 2.1 | 1.15 | 4.47 | 12.03 | 0.8 | 2.28 | 2.03 |
| Run1_MDS23 | | 0.37 | 2.09 | 3.45 | 1.83 | 1.82 | 1.91 | 5.73 | 5.58 | 9.14 | -0.14 | -0.23 | -0.06 | 0.61 | -0.21 | -0.13 | 0.88 | 2.51 | 1.12 | 0.34 | 1.72 | 0.5 | 2.31 | 10.71 | 0.98 | 1.93 | 1.67 |
| Run1_MDS25 | | 2.06 | 4.36 | 3.38 | 2.97 | 2.14 | 4.53 | 6.98 | 7.08 | 12.63 | 0.21 | 0.3 | 1.46 | 1.92 | 0.04 | 0.51 | 2.03 | 5.78 | -0.03 | -0.29 | 2.57 | 1.56 | 2.66 | 4.04 | 1.9 | 2.63 | 2.51 |
| Run1_MDS5 | | 1.94 | 4.88 | 3.97 | 6.59 | 2.7 | 0.71 | 1.91 | 4.44 | 6.06 | 0.47 | 1.14 | 1.13 | 2.18 | 0.26 | 0.73 | 1.93 | 3.14 | 0.82 | -0.17 | 2.16 | 3.54 | 3.45 | 3.26 | 1.71 | 2.24 | 2.15 |
| Run1_MDS6 | | 0.96 | 3.41 | 3.47 | 2.75 | 2.69 | 1.96 | 2.42 | 2.5 | 5.05 | 0.06 | -0.13 | -0.07 | 1.1 | 0.45 | 0.37 | 0.99 | 1.52 | 0.38 | -0.33 | 2.15 | 2.2 | 3.51 | 7.75 | 1.94 | 2.28 | 2.07 |
| Run1_MDS8 | | 0.93 | 4.23 | 5.2 | 2.66 | 2.78 | 2.48 | 3.14 | 3.99 | 6.98 | -0.06 | -0.12 | 0.4 | 0.77 | 0.64 | 0.93 | 1.06 | 2.35 | 0.36 | -0.39 | 1.54 | 2.01 | 1.74 | 3.78 | 1.9 | 1.82 | 1.41 |
| Run1_MDS9 | | 0.91 | 4.37 | 5.69 | 3.71 | 4.43 | 4.16 | 9.15 | 10.67 | 16.3 | -0.09 | -0.17 | 0.23 | 0.87 | 0.26 | 0.09 | 1.14 | 2.98 | 0.61 | -0.38 | 2.11 | 1.9 | 5.51 | 4.7 | 1.45 | 2.43 | 1.93 |
| Run2_MDS12 | Lower Risk | 0.42 | 0.98 | 3.11 | 1.25 | 2.56 | 1.54 | 3.37 | 4.26 | 9.26 | -0.11 | -0.08 | 0.19 | 0.88 | -0.04 | -0.05 | 0.13 | 2.77 | 0.79 | 0.38 | 1.57 | 1.21 | 1.14 | 3.28 | 1.13 | 1.8 | 1.4 |
| Run2_MDS14 | | 1.12 | 6.32 | 4.72 | 5.7 | 1.32 | 4.72 | 14.08 | 7.65 | 17.23 | 0.59 | -0.09 | -0.09 | 1.31 | -0.08 | -0.2 | 1 | 0.59 | 0.1 | -0.4 | 1.79 | 0.55 | 1.43 | 4.56 | 0.71 | 1.74 | 1.81 |
| Run2_MDS19 | | 0.18 | 3.68 | 4.65 | 3.38 | 4.06 | 1.88 | 2.61 | 2.96 | 5.36 | -0.17 | -0.22 | -0.19 | 0.37 | 0.02 | -0.03 | -0.14 | 3.31 | 0.74 | -0.01 | 1.97 | 1.32 | 2.12 | 3.99 | 1.26 | 2.21 | 1.9 |
| Run2_MDS20 | | 0.8 | 2.16 | 1.02 | 2.81 | 3.35 | 0.48 | 0.85 | 1.41 | 3.34 | -0.08 | -0.18 | -0.16 | 0.4 | -0.06 | 0.08 | 0.46 | 2.39 | 0.95 | -0.51 | 1.84 | 0.83 | 1.57 | 3.85 | 1.48 | 2.41 | 1.79 |
| Run2_MDS28* | | 0.29 | 3.82 | 5.43 | 3.26 | 4.23 | 0.65 | 1.46 | 4.37 | 9.69 | -0.1 | -0.18 | -0.13 | 0.41 | 0.6 | -0.04 | -0.02 | 3.43 | -0.04 | -0.38 | 1.05 | 0.4 | 3.37 | 6.82 | 0.71 | 1.21 | 0.93 |
| Run2_MDS5 | | 1.41 | 7.01 | 2.26 | 6.28 | 2.38 | 0.89 | 1.12 | 3.61 | 4.75 | 0.32 | 0.6 | 0.67 | 1.65 | -0.11 | 0.62 | 1.56 | 2.39 | 0.23 | -0.34 | 2.04 | 3.19 | 2.79 | 3.07 | 1.36 | 2.2 | 1.97 |
| Run1_ICUS18 | | 0.87 | 4.86 | 5.83 | 3.69 | 2.61 | 3.26 | 4.45 | 3.9 | 7.23 | -0.12 | -0.16 | -0.13 | 0.67 | 0.56 | 0.62 | 0.51 | 3.4 | 0.57 | -0.02 | 2.02 | 1.72 | 3.37 | 4.94 | 1.77 | 2.41 | 1.94 |
| Run1_ICUS22 | ICUS | 0.48 | 3.55 | 3.84 | 2.92 | 3.37 | 3.55 | 10.5 | 7.06 | 13.86 | -0.16 | -0.24 | -0.27 | 0.43 | -0.12 | 0.01 | 0.31 | 1.4 | -0.14 | -0.17 | 1.18 | 0.29 | 19.74 | 7.34 | 1.52 | 1.23 | 1.17 |
| Run1_ICUS24 | | 0.47 | 5.76 | 7.01 | 5.03 | 4.24 | 3.59 | 5.8 | 7.97 | 16.97 | -0.1 | -0.19 | -0.14 | 0.63 | -0.1 | -0.17 | 0.67 | 0.37 | -0.12 | -0.13 | 1.22 | 0.19 | 2.02 | 5.83 | 1.14 | 1.18 | 1.24 |
| Run1_Nl-1 | | 0.53 | 4.19 | 4.86 | 3.37 | 2.65 | 2.72 | 5.07 | 4.16 | 8.69 | -0.01 | -0.18 | -0.19 | 0.78 | 0.05 | 0.2 | 0.56 | 0.25 | -0.08 | 0.01 | 1.29 | 0.85 | 3.01 | 6.53 | 1.48 | 1.38 | 1.25 |
| Run1_Nl-3 | | 1.15 | 6.1 | 6.38 | 5.46 | 4.72 | 4.61 | 5.01 | 6.23 | 13.46 | 0.3 | 0.1 | 0.46 | 2.14 | 1.06 | 0.3 | 1.09 | 1.21 | 0.08 | -0.23 | 1.79 | 2.19 | 2.61 | 7.56 | 1.69 | 1.74 | 1.83 |
| Run1_Nl-4 | | 1.14 | 4.64 | 5.84 | 3.18 | 3.5 | 2.94 | 3.49 | 4.18 | 7.37 | 0.45 | 0.31 | 0.61 | 1.96 | 1.08 | 0.35 | 0.97 | 0.91 | 0.37 | -0.17 | 2.18 | 1.64 | 1.91 | 4.57 | 1.97 | 1.97 | 2.36 |
| Run1_Nl-6 | | 1.15 | 3.63 | 4.49 | 2.72 | 3.37 | 4.06 | 5.69 | 7.41 | 17.09 | 0.42 | 0.43 | 0.61 | 2.28 | 0.13 | -0.06 | 0.79 | 0.51 | -0.02 | -0.34 | 1.96 | 1.37 | 3.3 | 5.48 | 1.17 | 2.15 | 1.88 |
| Run2_Nl-3 | Normal | 0.69 | 4.95 | 6.5 | 4.12 | 3.65 | 2.43 | 3.52 | 6.27 | 13.83 | 0.07 | -0.01 | 0.3 | 1.48 | 0.6 | 0.11 | 0.6 | 0.99 | 0.08 | -0.24 | 1.51 | 1.84 | 2.26 | 5.19 | 1.3 | 1.52 | 1.51 |
| Run2_Nl-4 | | 0.58 | 4.29 | 5.38 | 3.57 | 3.06 | 1.31 | 1.91 | 3.19 | 6.57 | 0.21 | 0.02 | 0.17 | 1.17 | 0.65 | 0.09 | 0.38 | 0.72 | 0.26 | -0.29 | 1.81 | 1.47 | 1.83 | 3.07 | 1.94 | 1.72 | 1.89 |
| Run2_Nl-5 | | 0.61 | 3.97 | 4.29 | 3.29 | 3.02 | 1.43 | 2.22 | 3.51 | 7.36 | -0.07 | -0.11 | -0.05 | 1.1 | 0.54 | -0.1 | 0.37 | 0.35 | 0.5 | -0.14 | 1.76 | 2.48 | 2.16 | 3.42 | 1.33 | 1.7 | 1.83 |
| Run2_Nl-6 | | 0.59 | 2.89 | 3.56 | 2.46 | 3.46 | 2.28 | 3.85 | 8.11 | 18.65 | 0.08 | 0 | 0.2 | 1.3 | -0.05 | -0.18 | 0 | 0.65 | 0.03 | -0.21 | 1.85 | 1.2 | 2.92 | 4.25 | 1.09 | 2.11 | 1.76 |

**Supplemental Table 3**: Median expression of each surface marker on each cell population.

### CXCR4 Median

| | | Ungated | CD34+CD38low | HSCs | MPP | CMP/GMP | Myelo/Mono-Blasts | ProMonocytes | CD14neg Monocytes | Mature Monocytes | ProMyelocytes | Myelocytes | MetaMyelocytes | MatureGrans | ProErythroblasts | Erythroblasts | LateErythroblasts | PreBcells | Mature Bcells | Plasma Cells | T cells | NK cells | pDCs | Basophils | Platelets | CD8+ T cells | CD8neg T cells |
|---|---|---|---|---|---|---|---|---|---|---|---|---|---|---|---|---|---|---|---|---|---|---|---|---|---|---|---|
| Run1_MDS17 | | 0.67 | 0.22 | 0.34 | 0.14 | 0.41 | 0.14 | 3.02 | 1.44 | 0.48 | 0.69 | 2.17 | 1.99 | 2.22 | -0.31 | -0.42 | -0.19 | 0.9 | 3.99 | 1.37 | 1.01 | 0.04 | 9.46 | 7.65 | 16.62 | 0.94 | 1.11 |
| Run2_MDS15* | AML/RAEB-T | 1.21 | -0.11 | -0.22 | -0.21 | 0.02 | 1.57 | 4.79 | 0.85 | 0.31 | 4 | 5.12 | 3.66 | 0.79 | -0.02 | 0.56 | 1.31 | 1.06 | 2.92 | 0.48 | 0.02 | -0.04 | 0.8 | 7.23 | 3.03 | -0.1 | 0.33 |
| Run2_MDS4 | | 0.35 | -0.2 | -0.22 | -0.28 | -0.23 | 0.6 | 0.08 | -0.07 | 0.11 | 1.83 | 1.95 | 0.91 | 0.54 | -0.29 | -0.35 | 0.18 | 0.98 | 2.15 | 0.91 | 0 | -0.13 | 0.68 | 8.42 | 19.46 | -0.04 | 0.1 |
| Run1_MDS21 | | -0.08 | -0.08 | -0.1 | -0.09 | 0.06 | 0.14 | 1.06 | 0.64 | 0.25 | -0.37 | -0.18 | -0.21 | -0.16 | -0.39 | -0.28 | 49.01 | 2.31 | 1.85 | 1.24 | 0.53 | -0.03 | 13.25 | 7.03 | 66.87 | 0.04 | 0.75 |
| Run1_MDS3 | | -0.16 | -0.13 | -0.13 | -0.17 | -0.03 | 0.03 | 0.23 | -0.14 | 0.55 | -0.37 | -0.32 | -0.3 | -0.12 | -0.35 | -0.29 | 1.41 | 2.22 | 2.54 | 1.55 | 0.91 | 0.11 | 6.06 | 10.58 | 48.51 | 0.55 | 1.06 |
| Run2_MDS13 | | 0.2 | -0.14 | -0.13 | -0.15 | 0 | 0.13 | 1.03 | 0.5 | 1.31 | 0.1 | 0.74 | 0.25 | 0.26 | -0.05 | -0.34 | -0.17 | 4.12 | 2.68 | 1.09 | 0.88 | -0.05 | 1.13 | 3.07 | 32.84 | 0.42 | 1.14 |
| Run2_MDS16 | | 0.06 | 0.35 | 0.51 | 0.22 | 0.25 | -0.03 | -0.12 | 0.06 | 0.05 | 2.3 | 0.99 | 0.1 | -0.19 | 0.42 | -0.36 | -0.01 | 0.73 | 2.67 | 0.86 | 0.96 | -0.08 | 0.89 | 9.64 | 5.79 | 0.86 | 0.98 |
| Run2_MDS1 | | 0.47 | 0.78 | 1.01 | 0.63 | 0.87 | 0.99 | 0.74 | -0.1 | -0.07 | 0.44 | 0.28 | 0.63 | 0.5 | -0.08 | -0.31 | -0.24 | 1.15 | 2.06 | 1.37 | 1.4 | 0.06 | 1.37 | 11.97 | 9.51 | 0.37 | 1.69 |
| Run2_MDS21 | Higher Risk | 0.21 | -0.12 | -0.1 | -0.14 | 0.09 | 0.12 | 1.97 | 0.75 | 0.96 | 0.21 | 0.96 | 0.61 | 0.37 | -0.23 | -0.36 | -0.19 | 2.26 | 2.93 | 1.73 | 0.54 | -0.14 | 14.4 | 6.83 | 78.82 | 0.13 | 0.73 |
| Run2_MDS26* | | 0.98 | 0.97 | 0.87 | 0.85 | 1.7 | 0.83 | 1.64 | 1.15 | 0.99 | 2 | 1.83 | 1.79 | 1.84 | -0.08 | -0.37 | -0.36 | 1.02 | 6.46 | 0.68 | 1.48 | -0.05 | 1.17 | 28.34 | 3.99 | 0.28 | 3.17 |
| Run2_MDS27 | | -0.13 | -0.12 | -0.11 | -0.16 | 0.18 | 0.13 | 1.23 | 0.35 | 0 | 2.63 | 2.1 | 0.44 | -0.19 | 0.12 | -0.39 | -0.36 | 0.05 | 2.21 | 0.83 | -0.06 | -0.12 | -0.23 | 10.63 | 3.82 | -0.14 | 0.06 |
| Run2_MDS2 | | -0.18 | -0.33 | -0.29 | -0.35 | -0.28 | -0.26 | -0.18 | -0.2 | 0 | 0.39 | -0.16 | -0.25 | -0.2 | 0.02 | -0.05 | 0.79 | -0.05 | 4.62 | -0.1 | 0.33 | -0.02 | 1.2 | 0.52 | 5.32 | -0.01 | 0.36 |
| Run2_MDS3 | | 0.22 | -0.05 | -0.06 | -0.06 | 0.06 | 0.22 | 1.07 | 0.82 | 1.06 | -0.08 | 0.5 | 0.43 | 0.53 | -0.09 | -0.32 | -0.3 | 3.66 | 3.87 | 1.17 | 1.32 | 0.28 | 4.16 | 10.93 | 64.64 | 0.95 | 1.5 |
| Run1_MDS23 | | 0.55 | 0.68 | 0.72 | 0.63 | 0.9 | 1.07 | 1.92 | 0.6 | 0.04 | 0.79 | 1.06 | 0.75 | 0.48 | -0.34 | -0.34 | 0.76 | 0.73 | 1.72 | 1.95 | 0.22 | -0.05 | 0.3 | 1.67 | 17.63 | -0.14 | 0.46 |
| Run1_MDS25 | | -0.24 | 1.09 | 0.66 | 0.21 | 0.73 | -0.1 | 0.66 | 0.34 | 0.1 | 0.52 | 0.66 | -0.24 | -0.31 | -0.29 | -0.29 | 0.8 | 0.63 | 2.19 | 0.99 | -0.02 | 0.2 | 0.67 | 16.31 | 8.16 | -0.13 | 0.21 |
| Run1_MDS5 | | 0.07 | 0.59 | -0.54 | 0.08 | -0.05 | 0.09 | 0.07 | 0.19 | 0.69 | 2.62 | 0.52 | -0.03 | -0.08 | 0.39 | -0.24 | 3.06 | 0.49 | 2.79 | 1.09 | 0.6 | -0.07 | 4.29 | 8.47 | 4.88 | 0.24 | 0.72 |
| Run1_MDS6 | | -0.03 | -0.14 | -0.26 | -0.23 | 0.24 | 0.1 | 0.52 | 0.34 | 0.1 | 0.41 | 1.12 | 0.74 | -0.17 | -0.12 | -0.42 | 0.46 | 0.4 | 1.27 | 1.03 | -0.03 | -0.07 | 1.52 | 3.54 | 5.59 | -0.09 | 0.04 |
| Run1_MDS8 | | -0.21 | 0.49 | -0.42 | 0.6 | 0.61 | 0.11 | -0.18 | -0.1 | 0.01 | 0.1 | 0.59 | -0.21 | -0.33 | -0.26 | -0.32 | 0.27 | 0.47 | 1.66 | 0.43 | 0.08 | -0.06 | 7.53 | 5.78 | 10.82 | 0.1 | 0.07 |
| Run1_MDS9 | | -0.02 | -0.11 | 0.03 | -0.21 | -0.05 | -0.12 | 0.26 | 0.54 | 0.98 | 0.53 | 1.08 | 0 | -0.24 | -0.3 | -0.34 | 5.95 | 0.49 | 2.12 | 1.18 | 0.26 | -0.02 | 0.45 | 36.3 | 16.17 | -0.09 | 0.54 |
| Run2_MDS12 | Lower Risk | 1.45 | 0.87 | 1.38 | 0.64 | 0.77 | 3.26 | 6.77 | 3.54 | 2.39 | 1.7 | 2.23 | 1.77 | 1.11 | -0.28 | -0.39 | -0.08 | 2.72 | 2.06 | 1.32 | 0.33 | -0.01 | 1.5 | 21.61 | 47.36 | 0 | 0.54 |
| Run2_MDS14 | | -0.03 | 0.09 | -0.2 | 1.08 | 0.92 | -0.06 | -0.14 | 0.09 | -0.08 | 1.2 | 1.19 | 0.74 | -0.1 | -0.19 | -0.39 | -0.03 | 2.06 | 3.54 | 0.65 | 0.25 | 0.35 | 0.68 | 11.85 | 17.8 | 0.14 | 0.29 |
| Run2_MDS19 | | 0.38 | 0.7 | 1.5 | 0.55 | 0.79 | 1.15 | 1.34 | 0.83 | 0.34 | 1.89 | 1.85 | 0.76 | -0.04 | -0.21 | -0.37 | -0.31 | 0.51 | 2.39 | 0.83 | 0.26 | -0.08 | 0.54 | 8.44 | 22.19 | -0.1 | 0.41 |
| Run2_MDS20 | | -0.03 | 0.58 | 0 | 0.26 | 0.8 | -0.16 | -0.18 | -0.13 | 0.11 | 2.23 | 1.99 | 0.64 | -0.17 | 0.07 | -0.28 | 0.01 | 0.58 | 2.06 | 0.99 | 0.22 | -0.18 | -0.14 | 18.51 | 6.95 | -0.04 | 0.25 |
| Run2_MDS28* | | 1.3 | 0.77 | 1.7 | 0.65 | 0.88 | 3.52 | 5.38 | 1.21 | 0.34 | 1.83 | 2.31 | 1.86 | 1.04 | 0.08 | -0.34 | 0.49 | 1.27 | 2.63 | 1.36 | 0.41 | 0.25 | 2.75 | 23.99 | 24.11 | 0.01 | 0.83 |
| Run2_MDS5 | | -0.02 | 0.1 | -0.66 | -0.26 | 0.28 | 0.16 | -0.1 | 0.13 | 0.26 | 3.41 | 0.07 | -0.04 | -0.06 | 0.78 | -0.34 | 1.49 | 0.33 | 2.89 | 0.56 | 0.41 | -0.12 | 3.08 | 8.6 | 6.01 | 0.1 | 0.62 |
| Run1_ICUS18 | | -0.19 | 0.42 | 0.56 | 0.01 | 0.88 | -0.19 | -0.1 | -0.04 | -0.06 | 1.34 | 1.75 | 0.35 | -0.32 | -0.33 | -0.39 | -0.23 | 0.53 | 2.42 | 2.19 | 0.84 | 0 | 0.23 | 4.26 | 21.97 | 0.17 | 0.98 |
| Run1_ICUS22 | ICUS | -0.19 | 0.47 | 1.15 | 0.34 | 0.68 | 0.05 | 0.3 | 0.45 | 0.24 | 0.89 | 1.26 | 0.19 | -0.36 | -0.3 | -0.32 | -0.1 | 1.42 | 2.46 | 1.57 | 0.4 | 0.56 | -0.13 | 29.86 | 12.56 | -0.07 | 0.59 |
| Run1_ICUS24 | | 0.36 | 0.52 | 0.33 | 0.57 | 1.04 | 1.44 | 2.96 | 0.97 | -0.01 | 0.98 | 1.61 | 0.86 | -0.08 | -0.33 | -0.43 | 3.82 | 2.26 | 2.51 | 1.39 | 0.18 | 0.22 | 1.35 | 29.64 | 14.05 | 0.12 | 0.22 |
| Run1_Nl-1 | | 0.32 | 0.58 | -0.43 | -0.07 | 1.43 | 1.73 | 1.56 | 1.19 | 0.16 | 1.2 | 1.81 | 0.72 | -0.18 | -0.32 | -0.38 | 0.57 | 3.98 | 4.02 | 1.8 | 1.65 | 0.33 | 1.41 | 169.21 | 12.59 | 2.06 | 1.44 |
| Run1_Nl-3 | | 0.47 | 0.7 | 0.65 | 0.64 | 0.92 | 2.15 | 3.19 | 1.72 | 0.47 | 1.24 | 1.33 | 0.61 | -0.07 | 0.08 | -0.33 | 0.38 | 1.43 | 2.17 | 1.11 | 0.23 | -0.06 | 0.79 | 13.42 | 8.63 | 0.13 | 0.28 |
| Run1_Nl-4 | | 0.33 | 0.7 | 0.41 | 0.78 | 0.93 | 1.92 | 3.94 | 1.41 | 0.52 | 1.19 | 1.2 | 0.39 | -0.13 | 0.32 | -0.36 | 0.22 | 1.59 | 1.75 | 1.41 | 0.35 | 0.44 | 1.04 | 20.29 | 13.32 | 0.44 | 0.29 |
| Run1_Nl-6 | | 0.33 | 0.75 | 0.93 | 0.65 | 0.87 | 2.11 | 3.34 | 0.89 | 0.1 | 1.18 | 1.15 | 0.56 | -0.05 | -0.19 | -0.41 | 0.08 | 2.08 | 2.3 | 1.25 | 0.5 | 0.1 | 0.61 | 14.62 | 8.59 | 0.13 | 0.6 |
| Run2_Nl-3 | Normal | 0.5 | 0.6 | 0.65 | 0.47 | 0.75 | 1.96 | 3.04 | 1.12 | 0.43 | 1.63 | 1.16 | 0.42 | -0.03 | -0.02 | -0.32 | 0.07 | 1.46 | 2.33 | 0.91 | 0.26 | -0.1 | 0.52 | 13.44 | 11.46 | 0.2 | 0.3 |
| Run2_Nl-4 | | 0.43 | 0.81 | 1.56 | 0.56 | 0.75 | 1.87 | 3.25 | 0.93 | 0.61 | 1.77 | 1.17 | 0.35 | -0.06 | 0.32 | -0.28 | 0.05 | 1.64 | 2.01 | 0.93 | 0.36 | 0.33 | 0.89 | 23.5 | 19.59 | 0.54 | 0.23 |
| Run2_Nl-5 | | 0.38 | 0.47 | 0.74 | 0.46 | 1.23 | 2.47 | 3.11 | 0.77 | 0.53 | 1.94 | 1.28 | 0.37 | -0.06 | 0.33 | -0.33 | -0.06 | 2.05 | 2.21 | 1.06 | 0.36 | -0.03 | 0.88 | 10.62 | 14.85 | 0.36 | 0.35 |
| Run2_Nl-6 | | 0.4 | 0.54 | 0.91 | 0.47 | 0.82 | 1.82 | 2.57 | 0.55 | 0.23 | 1.71 | 1.21 | 0.51 | -0.04 | -0.12 | -0.36 | -0.15 | 1.95 | 2.52 | 1.11 | 0.47 | 0 | 0.45 | 10.95 | 14.72 | 0.1 | 0.61 |

**Supplemental Table 3**: Median expression of each surface marker on each cell population.

### CD10 Median

| | | Ungated | CD34+CD38low | HSCs | MPP | CMP/GMP | Myelo/Mono-Blasts | ProMonocytes | CD14neg Monocytes | Mature Monocytes | ProMyelocytes | Myelocytes | MetaMyelocytes |
|---|---|---|---|---|---|---|---|---|---|---|---|---|---|
| Run1_MDS17 | | -0.07 | -0.06 | 0 | -0.09 | -0.15 | -0.16 | -0.03 | 0.34 | 1.03 | -0.21 | -0.15 | 0.47 |
| Run2_MDS15* | AML/RAEB-T | 0.39 | -0.09 | -0.13 | -0.16 | -0.2 | -0.18 | -0.14 | 1.59 | 1.08 | -0.16 | -0.09 | 0.59 |
| Run2_MDS4 | | 0.06 | -0.29 | -0.33 | -0.32 | 0.77 | -0.24 | 0.03 | 0.37 | 0.8 | -0.33 | -0.28 | 0.36 |
| Run1_MDS21 | | -0.24 | -0.22 | -0.27 | -0.25 | -0.29 | -0.28 | 0.03 | 0.57 | 0.59 | -0.37 | -0.29 | -0.17 |
| Run1_MDS3 | | -0.23 | -0.23 | -0.27 | -0.25 | -0.26 | -0.27 | -0.05 | 0.28 | 1.11 | -0.4 | -0.34 | -0.14 |
| Run2_MDS13 | | -0.23 | -0.22 | -0.23 | -0.24 | -0.26 | -0.27 | 0.04 | 0.02 | 0.71 | -0.35 | -0.29 | -0.15 |
| Run2_MDS16 | | 0.01 | -0.2 | -0.17 | -0.26 | -0.12 | 0.01 | 0.33 | 0.04 | 0.41 | -0.39 | -0.31 | -0.18 |
| Run2_MDS1 | | -0.07 | -0.07 | -0.04 | -0.12 | -0.13 | -0.21 | -0.12 | -0.08 | 0.3 | -0.35 | -0.28 | 0.49 |
| Run2_MDS21 | Higher Risk | -0.25 | -0.25 | -0.19 | -0.26 | -0.27 | -0.28 | -0.2 | -0.03 | 0.35 | -0.35 | -0.32 | -0.2 |
| Run2_MDS26* | | -0.25 | -0.2 | -0.15 | -0.21 | -0.35 | -0.35 | -0.23 | 0.08 | -0.13 | -0.39 | -0.37 | -0.16 |
| Run2_MDS27 | | -0.2 | -0.23 | -0.19 | -0.26 | -0.2 | -0.21 | -0.05 | -0.09 | 0.23 | -0.38 | -0.35 | -0.3 |
| Run2_MDS2 | | -0.13 | -0.19 | -0.02 | -0.23 | -0.19 | -0.22 | -0.09 | -0.09 | 0.35 | -0.37 | -0.25 | -0.14 |
| Run2_MDS3 | | -0.22 | -0.23 | -0.26 | -0.22 | -0.27 | -0.27 | -0.26 | 0.1 | 0.48 | -0.33 | -0.27 | -0.15 |
| Run1_MDS23 | | -0.05 | -0.32 | -0.3 | -0.29 | -0.27 | -0.28 | -0.04 | 0.01 | 0.38 | -0.38 | -0.37 | -0.14 |
| Run1_MDS25 | | 3.57 | 1.18 | 0.95 | 5.06 | -0.59 | -0.15 | 0.18 | 0.22 | 0.53 | -0.34 | -0.2 | 2.7 |
| Run1_MDS5 | | 0.18 | 0.38 | 0.51 | 0.19 | -0.47 | -0.28 | -0.03 | -0.06 | 0.97 | -0.17 | 0.04 | 0.02 |
| Run1_MDS6 | | 1.15 | -0.27 | -0.27 | -0.37 | -0.35 | -0.22 | 0.08 | 0.18 | 0.54 | -0.33 | -0.31 | -0.11 |
| Run1_MDS8 | | 1.43 | 0.04 | -0.58 | 0.05 | -0.18 | -0.09 | 0.29 | -0.04 | 0.32 | -0.36 | -0.26 | 1.37 |
| Run1_MDS9 | | 0.17 | -0.23 | -0.26 | -0.26 | -0.2 | -0.15 | 0.39 | 0.21 | 0.49 | -0.35 | -0.33 | 0.1 |
| Run2_MDS12 | Lower Risk | -0.12 | -0.28 | -0.04 | -0.32 | -0.22 | -0.17 | 0.01 | 0.08 | 0.42 | -0.38 | -0.38 | -0.17 |
| Run2_MDS14 | | 0.68 | 0.17 | -0.07 | 2.09 | 0.14 | 0.17 | 0.29 | 0 | 0.24 | -0.4 | -0.34 | -0.27 |
| Run2_MDS19 | | -0.12 | -0.22 | -0.11 | -0.22 | -0.19 | -0.21 | -0.06 | -0.06 | 0.31 | -0.39 | -0.36 | -0.28 |
| Run2_MDS20 | | 0.37 | -0.28 | -0.14 | 0.87 | -0.16 | -0.13 | 0.08 | -0.04 | 0.33 | -0.37 | -0.32 | -0.22 |
| Run2_MDS28* | | -0.16 | -0.28 | -0.18 | -0.26 | -0.21 | -0.3 | -0.14 | -0.1 | 0.15 | -0.38 | -0.38 | -0.33 |
| Run2_MDS5 | | 0.08 | 0.13 | -0.37 | -0.28 | -0.37 | -0.33 | -0.14 | -0.1 | 0.37 | -0.34 | -0.11 | -0.06 |
| Run1_ICUS18 | | 1.35 | -0.03 | -0.24 | -0.06 | -0.26 | 0.11 | 0.32 | 0.01 | 0.46 | -0.34 | -0.3 | -0.05 |
| Run1_ICUS22 | ICUS | 0.4 | -0.31 | -0.4 | -0.31 | -0.14 | -0.13 | 0.23 | -0.05 | 0.25 | -0.41 | -0.39 | -0.35 |
| Run1_ICUS24 | | 0.15 | -0.19 | -0.08 | -0.28 | 0.24 | -0.02 | 0.12 | 0.07 | 0.44 | -0.38 | -0.35 | -0.24 |
| Run1_Nl-1 | | 0.37 | -0.05 | -0.26 | -0.05 | 0.65 | 0.2 | 0.34 | 0.03 | 0.43 | -0.37 | -0.38 | -0.33 |
| Run1_Nl-3 | | 0.57 | -0.17 | -0.04 | -0.16 | -0.16 | -0.11 | -0.03 | -0.02 | 0.35 | -0.24 | -0.24 | 0.03 |
| Run1_Nl-4 | | 0.72 | -0.09 | -0.19 | -0.39 | -0.24 | -0.15 | 0.11 | 0.04 | 0.51 | -0.19 | -0.19 | 0.31 |
| Run1_Nl-6 | | 0.44 | -0.14 | -0.02 | -0.22 | -0.03 | -0.09 | 0.15 | 0.06 | 0.51 | -0.18 | -0.13 | -0.07 |
| Run2_Nl-3 | Normal | 0.25 | -0.27 | -0.12 | -0.3 | -0.09 | -0.13 | -0.01 | 0.1 | 0.44 | -0.32 | -0.21 | 0.12 |
| Run2_Nl-4 | | 0.31 | -0.29 | -0.36 | -0.28 | -0.19 | -0.16 | 0.02 | 0.18 | 0.6 | -0.33 | -0.22 | 0.14 |
| Run2_Nl-5 | | 0 | -0.18 | -0.33 | -0.41 | 0.21 | -0.09 | 0.06 | 0.1 | 0.52 | -0.35 | -0.28 | -0.15 |
| Run2_Nl-6 | | -0.08 | -0.09 | 0.02 | -0.13 | -0.03 | -0.14 | 0.02 | 0.09 | 0.46 | -0.34 | -0.32 | -0.24 |

| | MatureGrans | ProErythroblasts | Erythroblasts | LateErythroblasts | PreBcells | Mature Bcells | Plasma Cells | T cells | NK cells | pDCs | Basophils | Platelets | CD8+ T cells | CD8neg T cells |
|---|---|---|---|---|---|---|---|---|---|---|---|---|---|---|
| Run1_MDS17 | 1.18 | -0.37 | -0.38 | -0.32 | 0.33 | 0.42 | -0.4 | -0.07 | -0.3 | -0.32 | 0.72 | -0.41 | -0.07 | -0.06 |
| Run2_MDS15* | 3.34 | -0.27 | 0.04 | 0.6 | 3.57 | 0.62 | -0.1 | -0.05 | -0.09 | 0.2 | -0.11 | -0.24 | -0.06 | -0.04 |
| Run2_MDS4 | 2.22 | -0.41 | -0.35 | -0.3 | 0.71 | 0.21 | -0.43 | 0 | 0.09 | 0.3 | -0.26 | -0.15 | 0 | 0 |
| Run1_MDS21 | 0.22 | -0.33 | -0.27 | -0.2 | 0.28 | 0.58 | -0.28 | -0.13 | -0.04 | -0.42 | -0.29 | -0.42 | -0.09 | -0.15 |
| Run1_MDS3 | 0.4 | -0.31 | -0.29 | -0.15 | 0.27 | 0.69 | -0.51 | 0.02 | 0.15 | 0.18 | -0.28 | -0.41 | -0.01 | 0.03 |
| Run2_MDS13 | -0.02 | -0.3 | -0.33 | -0.32 | 0.11 | 0.3 | -0.28 | -0.07 | -0.03 | -0.16 | -0.33 | -0.44 | -0.07 | -0.07 |
| Run2_MDS16 | 0.68 | -0.28 | -0.35 | -0.16 | 0.14 | 0.37 | -0.26 | -0.01 | -0.3 | 0.09 | -0.19 | -0.42 | -0.04 | 0 |
| Run2_MDS1 | 1.26 | -0.33 | -0.27 | -0.27 | 0.21 | 0.3 | -0.34 | -0.01 | -0.08 | -0.02 | -0.37 | -0.29 | 0 | -0.01 |
| Run2_MDS21 | -0.06 | -0.31 | -0.33 | -0.35 | 0.11 | 0.6 | -0.27 | -0.1 | -0.16 | -0.05 | -0.18 | -0.39 | -0.07 | -0.12 |
| Run2_MDS26* | 0.48 | -0.35 | -0.32 | -0.35 | -0.02 | 0.22 | -0.3 | -0.09 | -0.33 | -0.09 | -0.35 | -0.37 | -0.07 | -0.11 |
| Run2_MDS27 | 0.02 | -0.34 | -0.38 | -0.34 | 0.23 | 0.38 | -0.4 | 0.03 | -0.27 | 0.4 | -0.18 | 0.53 | -0.01 | 0.07 |
| Run2_MDS2 | -0.03 | -0.15 | -0.13 | -0.14 | 0 | 0.96 | -0.03 | -0.03 | -0.29 | -0.28 | -0.33 | -0.41 | -0.02 | -0.03 |
| Run2_MDS3 | 0.17 | -0.24 | -0.31 | -0.36 | 0.21 | 0.56 | -0.34 | 0.07 | -0.03 | -0.37 | -0.34 | -0.37 | 0.07 | 0.07 |
| Run1_MDS23 | 1.79 | -0.32 | -0.24 | -0.08 | 0.31 | 0.41 | -0.49 | -0.01 | -0.06 | -0.15 | -0.32 | -0.34 | 0.01 | -0.02 |
| Run1_MDS25 | 4.95 | -0.25 | -0.09 | 2.02 | 0.15 | 0.42 | -0.57 | -0.1 | -0.3 | -0.05 | -0.2 | -0.5 | -0.13 | -0.05 |
| Run1_MDS5 | 0.54 | -0.2 | -0.25 | -0.14 | 0.27 | 0.32 | -0.52 | -0.1 | 0.03 | 0.04 | -0.35 | -0.36 | -0.14 | -0.08 |
| Run1_MDS6 | 3 | -0.33 | -0.39 | 0.17 | 0.5 | 0.43 | -0.56 | -0.05 | -0.19 | -0.04 | -0.22 | -0.45 | -0.06 | -0.04 |
| Run1_MDS8 | 2.98 | -0.29 | -0.31 | -0.15 | 0.08 | 0.42 | -0.47 | -0.26 | -0.29 | 0.08 | -0.28 | -0.47 | -0.22 | -0.27 |
| Run1_MDS9 | 1.25 | -0.33 | -0.34 | -0.14 | 0.28 | 0.23 | -0.41 | -0.12 | -0.27 | 0.18 | -0.2 | -0.43 | -0.08 | -0.15 |
| Run2_MDS12 | 1.07 | -0.35 | -0.34 | -0.28 | 0.09 | 0.05 | -0.39 | -0.06 | -0.12 | -0.17 | -0.3 | -0.37 | -0.02 | -0.09 |
| Run2_MDS14 | 1.31 | -0.31 | -0.35 | 0.69 | 1.39 | 0.55 | -0.33 | 0.1 | -0.24 | 0.05 | -0.24 | -0.37 | -0.01 | 0.17 |
| Run2_MDS19 | 0.47 | -0.36 | -0.37 | -0.34 | 0.36 | 0.58 | -0.54 | 0.04 | -0.25 | -0.28 | -0.3 | -0.39 | 0 | 0.05 |
| Run2_MDS20 | 1.12 | -0.28 | -0.26 | 0.31 | 0.28 | 0.44 | -0.39 | -0.08 | -0.26 | 0.18 | -0.17 | -0.41 | -0.11 | -0.08 |
| Run2_MDS28* | 0.17 | -0.27 | -0.34 | -0.21 | 0.22 | 0.26 | -0.43 | -0.01 | -0.11 | -0.06 | -0.1 | -0.28 | -0.02 | 0.02 |
| Run2_MDS5 | 0.37 | -0.36 | -0.27 | -0.11 | 0.28 | 0.37 | -0.46 | -0.07 | -0.05 | -0.25 | -0.19 | -0.36 | -0.07 | -0.07 |
| Run1_ICUS18 | 2.32 | -0.28 | -0.25 | 1.7 | 0.38 | 0.56 | -0.32 | -0.02 | -0.27 | 0.08 | -0.18 | -0.46 | -0.02 | -0.02 |
| Run1_ICUS22 | 1.03 | -0.36 | -0.29 | 0.31 | 1.34 | 0.45 | -0.52 | 0.06 | -0.38 | 0.32 | -0.26 | -0.45 | 0.04 | 0.06 |
| Run1_ICUS24 | 1.05 | -0.35 | -0.37 | -0.25 | 2.04 | 0.43 | -0.34 | -0.01 | -0.3 | -0.23 | -0.32 | -0.47 | -0.06 | 0.06 |
| Run1_Nl-1 | 1.64 | -0.35 | -0.29 | 0.18 | 1.8 | 0.73 | -0.48 | -0.02 | -0.32 | 0.33 | -0.23 | -0.42 | -0.02 | -0.02 |
| Run1_Nl-3 | 3.63 | -0.23 | -0.27 | 1.17 | 0.89 | 0.39 | -0.45 | -0.06 | -0.25 | -0.05 | -0.2 | -0.48 | -0.09 | -0.05 |
| Run1_Nl-4 | 3.88 | -0.21 | -0.26 | 1.15 | 0.84 | 0.43 | -0.45 | -0.04 | -0.27 | 0.24 | -0.19 | -0.46 | -0.03 | -0.04 |
| Run1_Nl-6 | 2.35 | -0.3 | -0.33 | 0.33 | 2.22 | 0.4 | -0.41 | 0.01 | -0.28 | 0.11 | -0.33 | -0.43 | -0.01 | 0.01 |
| Run2_Nl-3 | 2.53 | -0.29 | -0.27 | 0.81 | 0.84 | 0.37 | -0.42 | -0.01 | -0.24 | -0.1 | -0.14 | -0.42 | -0.02 | -0.01 |
| Run2_Nl-4 | 2.56 | -0.28 | -0.26 | 0.6 | 1.05 | 0.45 | -0.35 | 0.02 | -0.25 | 0.3 | -0.19 | -0.39 | -0.01 | 0.06 |
| Run2_Nl-5 | 1.42 | -0.27 | -0.33 | 0.16 | 2.15 | 0.63 | -0.35 | -0.07 | -0.3 | 0.06 | -0.1 | -0.41 | -0.08 | -0.06 |
| Run2_Nl-6 | 0.82 | -0.34 | -0.36 | -0.19 | 2.12 | 0.37 | -0.35 | 0.1 | -0.25 | -0.01 | -0.29 | -0.44 | 0.06 | 0.12 |

**Supplemental Table 3**: Median expression of each surface marker on each cell population.

### CD19 Median

| | | Ungated | CD34+CD38low | HSCs | MPP | CMP/GMP | Myelo/Mono-Blasts | ProMonocytes | CD14neg Monocytes | Mature Monocytes | ProMyelocytes | Myelocytes | MetaMyelocytes | MatureGrans | ProErythroblasts | Erythroblasts | LateErythroblasts | PreBcells | Mature Bcells | Plasma Cells | T cells | NK cells | pDCs | Basophils | Platelets | CD8+ T cells | CD8neg T cells |
|---|---|---|---|---|---|---|---|---|---|---|---|---|---|---|---|---|---|---|---|---|---|---|---|---|---|---|---|
| Run1_MDS17 | | -0.46 | -0.46 | -0.46 | -0.47 | -0.47 | -0.48 | -0.53 | -0.46 | -0.46 | -0.49 | -0.48 | -0.46 | -0.42 | -0.5 | -0.5 | -0.48 | 2.7 | 13.83 | -0.33 | -0.46 | -0.48 | -0.17 | -0.56 | -0.44 | -0.45 | -0.47 |
| Run2_MDS15* | AML/RAEB-T | -0.38 | -0.41 | -0.41 | -0.42 | -0.43 | -0.42 | -0.42 | -0.37 | -0.36 | -0.43 | -0.43 | -0.38 | -0.3 | -0.38 | -0.33 | -0.32 | 2.79 | 24.69 | -0.12 | -0.42 | -0.39 | -0.41 | -0.41 | -0.49 | -0.42 | -0.42 |
| Run2_MDS4 | | -0.39 | -0.43 | -0.45 | -0.44 | -0.07 | -0.42 | -0.4 | -0.4 | -0.36 | -0.43 | -0.41 | -0.42 | -0.35 | -0.45 | -0.43 | -0.43 | 3.55 | 5.41 | -0.28 | -0.41 | -0.4 | -0.32 | -0.45 | -0.22 | -0.41 | -0.42 |
| Run1_MDS21 | | -0.46 | -0.47 | -0.42 | -0.49 | -0.47 | -0.47 | -0.45 | -0.44 | -0.45 | -0.49 | -0.51 | -0.49 | -0.47 | -0.51 | -0.49 | -0.45 | 10.65 | 16.46 | -0.66 | -0.47 | -0.44 | -0.54 | -0.51 | -0.48 | -0.47 | -0.47 |
| Run1_MDS3 | | -0.46 | -0.46 | -0.48 | -0.5 | -0.49 | -0.48 | -0.46 | -0.53 | -0.34 | -0.5 | -0.48 | -0.5 | -0.46 | -0.5 | -0.47 | -0.51 | 9.42 | 16.11 | -0.1 | -0.47 | -0.46 | -0.35 | -0.37 | -0.47 | -0.45 | -0.48 |
| Run2_MDS13 | | -0.41 | -0.42 | -0.43 | -0.42 | -0.43 | -0.43 | -0.36 | -0.4 | -0.33 | -0.44 | -0.43 | -0.42 | -0.41 | -0.36 | -0.44 | -0.37 | 8.09 | 12.74 | -0.09 | -0.42 | -0.41 | -0.06 | -0.4 | -0.41 | -0.41 | -0.43 |
| Run2_MDS16 | | -0.4 | -0.41 | -0.43 | -0.42 | -0.4 | -0.47 | -0.46 | -0.44 | -0.41 | -0.44 | -0.44 | -0.44 | -0.4 | -0.43 | -0.44 | -0.42 | 2.85 | 17.3 | 0.01 | -0.41 | -0.43 | -0.41 | -0.43 | -0.43 | -0.42 | -0.41 |
| Run2_MDS1 | | -0.41 | -0.42 | -0.35 | -0.47 | -0.4 | -0.42 | -0.45 | -0.43 | -0.43 | -0.43 | -0.47 | -0.41 | -0.36 | -0.45 | -0.42 | -0.41 | 2.84 | 8.94 | -0.3 | -0.41 | -0.43 | -0.43 | -0.36 | -0.51 | -0.4 | -0.41 |
| Run2_MDS21 | Higher Risk | -0.41 | -0.41 | -0.44 | -0.41 | -0.42 | -0.43 | -0.41 | -0.42 | -0.45 | -0.43 | -0.42 | -0.44 | -0.43 | -0.47 | -0.45 | -0.46 | 10.77 | 17.11 | -0.15 | -0.43 | -0.39 | -0.47 | -0.53 | -0.41 | -0.43 | -0.43 |
| Run2_MDS26* | | -0.43 | -0.44 | -0.42 | -0.46 | -0.4 | -0.41 | -0.34 | -0.41 | -0.27 | -0.45 | -0.45 | -0.44 | -0.41 | -0.43 | -0.44 | -0.42 | 2.78 | 10.52 | -0.31 | -0.42 | -0.42 | -0.51 | -0.43 | -0.47 | -0.4 | -0.44 |
| Run2_MDS27 | | -0.42 | -0.42 | -0.42 | -0.42 | -0.45 | -0.4 | -0.42 | -0.38 | -0.4 | -0.45 | -0.44 | -0.43 | -0.43 | -0.44 | -0.43 | -0.46 | 2.97 | 17.63 | -0.22 | -0.41 | -0.43 | -0.39 | -0.47 | -0.43 | -0.41 | -0.41 |
| Run2_MDS2 | | -0.42 | -0.4 | -0.36 | -0.41 | -0.45 | -0.44 | -0.46 | -0.42 | -0.39 | -0.42 | -0.5 | -0.43 | -0.43 | -0.4 | -0.34 | -0.41 | 2.82 | 6.32 | -0.38 | -0.4 | -0.42 | -0.59 | -0.48 | -0.43 | -0.38 | -0.4 |
| Run2_MDS3 | | -0.41 | -0.44 | -0.36 | -0.49 | -0.41 | -0.42 | -0.42 | -0.44 | -0.37 | -0.44 | -0.42 | -0.42 | -0.42 | -0.43 | -0.45 | -0.42 | 11.02 | 17.18 | -0.14 | -0.41 | -0.5 | -0.51 | -0.48 | -0.4 | -0.42 | -0.41 |
| Run1_MDS23 | | -0.46 | -0.55 | -0.72 | -0.47 | -0.48 | -0.48 | -0.48 | -0.48 | -0.46 | -0.49 | -0.5 | -0.48 | -0.42 | -0.5 | -0.49 | -0.46 | 2.78 | 14.03 | -0.32 | -0.48 | -0.48 | -0.52 | -0.44 | -0.45 | -0.45 | -0.48 |
| Run1_MDS25 | | -0.32 | -0.42 | -0.77 | -0.71 | -0.33 | -0.45 | -0.51 | -0.47 | -0.47 | -0.51 | -0.52 | -0.39 | -0.32 | -0.45 | -0.46 | -0.22 | 2.94 | 18.1 | -0.25 | -0.47 | -0.5 | -0.48 | -0.48 | -0.55 | -0.47 | -0.47 |
| Run1_MDS5 | | -0.45 | -0.32 | -0.12 | -0.48 | -0.52 | -0.53 | -0.43 | -0.46 | -0.44 | -0.51 | -0.51 | -0.47 | -0.46 | -0.37 | -0.5 | -0.4 | 2.74 | 13.39 | -0.12 | -0.49 | -0.45 | -0.42 | -0.45 | -0.44 | -0.5 | -0.48 |
| Run1_MDS6 | | -0.39 | -0.5 | -0.54 | -0.47 | -0.46 | -0.46 | -0.46 | -0.49 | -0.45 | -0.49 | -0.5 | -0.48 | -0.37 | -0.46 | -0.5 | -0.42 | 3.63 | 12.21 | -0.01 | -0.47 | -0.48 | -0.43 | -0.44 | -0.48 | -0.46 | -0.48 |
| Run1_MDS8 | | -0.4 | -0.5 | -0.61 | -0.69 | -0.4 | -0.44 | -0.53 | -0.49 | -0.45 | -0.49 | -0.51 | -0.44 | -0.38 | -0.48 | -0.47 | -0.43 | 2.74 | 15.88 | -0.49 | -0.49 | -0.5 | -0.28 | -0.51 | -0.48 | -0.49 | -0.49 |
| Run1_MDS9 | | -0.45 | -0.49 | -0.49 | -0.49 | -0.46 | -0.47 | -0.48 | -0.46 | -0.48 | -0.5 | -0.5 | -0.49 | -0.44 | -0.5 | -0.49 | -0.44 | 2.65 | 10.44 | -0.57 | -0.48 | -0.46 | -0.55 | -0.47 | -0.47 | -0.48 | -0.49 |
| Run2_MDS12 | Lower Risk | -0.41 | -0.45 | -0.37 | -0.43 | -0.39 | -0.43 | -0.42 | -0.41 | -0.4 | -0.44 | -0.44 | -0.43 | -0.38 | -0.44 | -0.45 | -0.42 | 2.96 | 11.53 | -0.18 | -0.4 | -0.43 | -0.42 | -0.38 | -0.4 | -0.41 | -0.39 |
| Run2_MDS14 | | -0.37 | -0.15 | -0.36 | -0.15 | -0.18 | -0.37 | -0.42 | -0.42 | -0.42 | -0.45 | -0.45 | -0.45 | -0.37 | -0.38 | -0.46 | -0.34 | 4.68 | 14.9 | 0.08 | -0.42 | -0.46 | -0.41 | -0.42 | -0.41 | -0.42 | -0.41 |
| Run2_MDS19 | | -0.42 | -0.39 | -0.37 | -0.39 | -0.45 | -0.44 | -0.42 | -0.41 | -0.39 | -0.44 | -0.45 | -0.44 | -0.41 | -0.38 | -0.46 | -0.43 | 2.73 | 24.34 | -0.31 | -0.41 | -0.43 | -0.46 | -0.45 | -0.43 | -0.42 | -0.4 |
| Run2_MDS20 | | -0.39 | -0.49 | -0.04 | -0.77 | -0.36 | -0.41 | -0.44 | -0.43 | -0.4 | -0.43 | -0.45 | -0.45 | -0.38 | -0.4 | -0.43 | -0.36 | 2.76 | 26.27 | -0.04 | -0.42 | -0.44 | -0.39 | -0.4 | -0.45 | -0.4 | -0.42 |
| Run2_MDS28* | | -0.42 | -0.36 | -0.31 | -0.46 | -0.43 | -0.43 | -0.42 | -0.43 | -0.4 | -0.43 | -0.45 | -0.44 | -0.42 | -0.41 | -0.44 | -0.41 | 2.82 | 13.8 | -0.31 | -0.42 | -0.43 | -0.44 | -0.39 | -0.47 | -0.42 | -0.42 |
| Run2_MDS5 | | -0.4 | -0.33 | 0.45 | -0.64 | -0.23 | -0.41 | -0.47 | -0.41 | -0.4 | -0.44 | -0.44 | -0.43 | -0.4 | -0.36 | -0.42 | -0.37 | 2.77 | 14.41 | -0.13 | -0.4 | -0.4 | -0.52 | -0.35 | -0.4 | -0.4 | -0.4 |
| Run1_ICUS18 | | -0.41 | -0.43 | -0.35 | -0.31 | -0.62 | -0.45 | -0.48 | -0.49 | -0.47 | -0.48 | -0.48 | -0.48 | -0.41 | -0.47 | -0.48 | -0.39 | 2.72 | 12.53 | -0.17 | -0.47 | -0.48 | -0.42 | -0.51 | -0.45 | -0.46 | -0.47 |
| Run1_ICUS22 | ICUS | -0.45 | -0.5 | -0.54 | -0.54 | -0.45 | -0.48 | -0.46 | -0.48 | -0.47 | -0.5 | -0.5 | -0.51 | -0.46 | -0.51 | -0.46 | -0.44 | 3.72 | 11.8 | -0.31 | -0.48 | -0.51 | -0.47 | -0.5 | -0.44 | -0.48 | -0.48 |
| Run1_ICUS24 | | -0.43 | -0.42 | -0.36 | -0.48 | -0.41 | -0.43 | -0.46 | -0.47 | -0.46 | -0.48 | -0.5 | -0.49 | -0.45 | -0.5 | -0.5 | -0.45 | 6.01 | 13.67 | -0.26 | -0.47 | -0.5 | -0.47 | -0.46 | -0.49 | -0.46 | -0.48 |
| Run1_Nl-1 | | -0.42 | -0.54 | -0.77 | -0.56 | -0.28 | -0.38 | -0.46 | -0.5 | -0.45 | -0.49 | -0.5 | -0.5 | -0.42 | -0.5 | -0.48 | -0.42 | 5.5 | 12.2 | -0.08 | -0.48 | -0.5 | -0.46 | -0.47 | -0.45 | -0.48 | -0.48 |
| Run1_Nl-3 | | -0.39 | -0.48 | -0.58 | -0.49 | -0.39 | -0.45 | -0.47 | -0.47 | -0.46 | -0.48 | -0.47 | -0.45 | -0.35 | -0.45 | -0.48 | -0.36 | 3.76 | 14.21 | -0.11 | -0.47 | -0.48 | -0.46 | -0.5 | -0.48 | -0.48 | -0.46 |
| Run1_Nl-4 | | -0.39 | -0.47 | -0.74 | -0.47 | -0.48 | -0.44 | -0.45 | -0.46 | -0.44 | -0.47 | -0.47 | -0.45 | -0.33 | -0.41 | -0.49 | -0.38 | 4.5 | 12.55 | -0.35 | -0.47 | -0.49 | -0.46 | -0.47 | -0.45 | -0.48 | -0.47 |
| Run1_Nl-6 | | -0.42 | -0.44 | -0.41 | -0.44 | -0.41 | -0.44 | -0.46 | -0.46 | -0.44 | -0.47 | -0.47 | -0.48 | -0.39 | -0.51 | -0.49 | -0.43 | 6.15 | 10.49 | -0.07 | -0.47 | -0.48 | -0.41 | -0.46 | -0.48 | -0.45 | -0.48 |
| Run2_Nl-3 | Normal | -0.36 | -0.44 | -0.5 | -0.39 | -0.35 | -0.41 | -0.41 | -0.4 | -0.41 | -0.42 | -0.42 | -0.39 | -0.32 | -0.42 | -0.44 | -0.36 | 4.22 | 14.78 | -0.18 | -0.42 | -0.43 | -0.38 | -0.43 | -0.44 | -0.42 | -0.42 |
| Run2_Nl-4 | | -0.36 | -0.36 | -0.26 | -0.48 | -0.4 | -0.39 | -0.41 | -0.42 | -0.38 | -0.42 | -0.41 | -0.39 | -0.31 | -0.42 | -0.44 | -0.37 | 4.57 | 13.53 | -0.26 | -0.41 | -0.43 | -0.43 | -0.42 | -0.41 | -0.41 | -0.4 |
| Run2_Nl-5 | | -0.38 | -0.42 | -0.47 | -0.37 | -0.33 | -0.38 | -0.39 | -0.41 | -0.38 | -0.43 | -0.42 | -0.42 | -0.35 | -0.41 | -0.43 | -0.4 | 5.53 | 12.67 | -0.19 | -0.42 | -0.42 | -0.39 | -0.42 | -0.43 | -0.42 | -0.42 |
| Run2_Nl-6 | | -0.39 | -0.4 | -0.41 | -0.38 | -0.4 | -0.41 | -0.41 | -0.41 | -0.37 | -0.43 | -0.44 | -0.43 | -0.38 | -0.43 | -0.44 | -0.42 | 6.27 | 10.97 | -0.06 | -0.41 | -0.44 | -0.4 | -0.4 | -0.44 | -0.4 | -0.41 |

**Supplemental Table 3**: Median expression of each surface marker on each cell population.

### CD20 Median

| | | Ungated | CD34+CD38low | HSCs | MPP | CMP/GMP | Myelo/Mono-Blasts | ProMonocytes | CD14neg Monocytes | Mature Monocytes | ProMyelocytes | Myelocytes | MetaMyelocytes | MatureGrans | ProErythroblasts | Erythroblasts | LateErythroblasts | PreBcells | Mature Bcells | Plasma Cells | T cells | NK cells | pDCs | Basophils | Platelets | CD8+ T cells | CD8neg T cells |
|---|---|---|---|---|---|---|---|---|---|---|---|---|---|---|---|---|---|---|---|---|---|---|---|---|---|---|---|
| Run1_MDS17 | | -0.41 | -0.45 | -0.44 | -0.45 | -0.43 | -0.43 | -0.33 | -0.38 | -0.55 | -0.42 | -0.34 | -0.41 | -0.39 | -0.29 | -0.34 | -0.39 | -0.39 | 50.07 | -0.43 | -0.4 | -0.46 | -0.38 | -0.25 | -0.28 | -0.37 | -0.45 |
| Run2_MDS15* | AML/RAEB-T | -0.33 | -0.37 | -0.39 | -0.42 | -0.4 | -0.28 | -0.22 | -0.36 | -0.38 | -0.21 | -0.25 | -0.34 | -0.33 | -0.3 | -0.14 | -0.25 | -0.29 | 137.6 | -0.23 | -0.34 | -0.36 | -0.39 | -0.36 | -0.35 | -0.28 | -0.39 |
| Run2_MDS4 | | -0.34 | -0.35 | -0.34 | -0.34 | -0.52 | -0.33 | -0.39 | -0.41 | -0.4 | -0.34 | -0.36 | -0.4 | -0.38 | -0.18 | -0.15 | -0.16 | 0.42 | 38.63 | -0.43 | -0.3 | -0.31 | 0.08 | -0.42 | -0.15 | -0.26 | -0.34 |
| Run1_MDS21 | | -0.41 | -0.45 | -0.39 | -0.46 | -0.46 | -0.44 | -0.3 | -0.45 | -0.41 | -0.45 | -0.42 | -0.45 | -0.43 | -0.35 | -0.33 | -0.18 | 1.78 | 32.69 | -0.62 | -0.47 | -0.45 | -0.56 | -0.57 | -0.24 | -0.45 | -0.48 |
| Run1_MDS3 | | -0.4 | -0.45 | -0.46 | -0.44 | -0.45 | -0.44 | -0.31 | -0.43 | -0.34 | -0.46 | -0.41 | -0.45 | -0.38 | -0.28 | -0.27 | -0.33 | 1.27 | 35.94 | -0.51 | -0.46 | -0.39 | -0.47 | -0.58 | -0.2 | -0.42 | -0.48 |
| Run2_MDS13 | | -0.36 | -0.4 | -0.42 | -0.4 | -0.41 | -0.39 | -0.36 | -0.38 | -0.33 | -0.39 | -0.37 | -0.39 | -0.38 | -0.17 | -0.21 | -0.24 | 1.92 | 63 | -0.3 | -0.38 | -0.3 | -0.17 | -0.42 | -0.24 | -0.36 | -0.4 |
| Run2_MDS16 | | -0.36 | -0.36 | -0.39 | -0.37 | -0.47 | -0.4 | -0.41 | -0.42 | -0.41 | -0.35 | -0.37 | -0.38 | -0.37 | -0.25 | -0.22 | -0.31 | -0.31 | 69.92 | -0.26 | -0.4 | -0.43 | -0.36 | -0.42 | -0.26 | -0.37 | -0.41 |
| Run2_MDS1 | | -0.38 | -0.41 | -0.49 | -0.47 | -0.4 | -0.38 | -0.33 | -0.41 | -0.4 | -0.32 | -0.39 | -0.41 | -0.4 | -0.24 | -0.24 | -0.33 | -0.28 | 84.39 | -0.17 | -0.38 | -0.25 | -0.43 | -0.46 | -0.28 | -0.28 | -0.4 |
| Run2_MDS21 | Higher Risk | -0.35 | -0.41 | -0.36 | -0.43 | -0.41 | -0.39 | -0.28 | -0.42 | -0.29 | -0.4 | -0.37 | -0.41 | -0.39 | -0.16 | -0.24 | -0.29 | 4.32 | 74.97 | -0.48 | -0.37 | -0.26 | -0.18 | -0.41 | -0.19 | -0.3 | -0.4 |
| Run2_MDS26* | | -0.38 | -0.33 | -0.29 | -0.34 | -0.32 | -0.34 | -0.37 | -0.37 | -0.36 | -0.37 | -0.35 | -0.42 | -0.41 | -0.25 | -0.29 | -0.31 | -0.29 | 53.1 | -0.36 | -0.35 | -0.43 | -0.41 | -0.41 | -0.29 | -0.23 | -0.41 |
| Run2_MDS27 | | -0.31 | -0.31 | -0.32 | -0.31 | -0.27 | -0.28 | -0.27 | -0.41 | -0.38 | -0.29 | -0.28 | -0.36 | -0.36 | -0.12 | -0.2 | -0.37 | -0.2 | 88.44 | -0.43 | -0.33 | -0.4 | -0.44 | -0.38 | -0.18 | -0.28 | -0.38 |
| Run2_MDS2 | | -0.41 | -0.4 | -0.51 | -0.39 | -0.45 | -0.44 | -0.41 | -0.4 | -0.39 | -0.2 | -0.33 | -0.41 | -0.4 | -0.42 | -0.26 | -0.26 | -0.39 | 34.67 | -0.4 | -0.41 | -0.43 | -0.41 | -0.43 | -0.22 | -0.43 | -0.41 |
| Run2_MDS3 | | -0.34 | -0.4 | -0.43 | -0.38 | -0.41 | -0.39 | -0.34 | -0.34 | -0.28 | -0.38 | -0.35 | -0.37 | -0.36 | -0.11 | -0.18 | -0.28 | 2.75 | 82.2 | -0.35 | -0.37 | -0.22 | -0.4 | -0.43 | -0.19 | -0.3 | -0.39 |
| Run1_MDS23 | | -0.43 | -0.44 | -0.44 | -0.47 | -0.46 | -0.43 | -0.4 | -0.46 | -0.48 | -0.43 | -0.4 | -0.44 | -0.44 | -0.37 | -0.36 | -0.37 | -0.29 | 73.76 | -0.44 | -0.41 | -0.42 | -0.47 | -0.53 | -0.19 | -0.27 | -0.45 |
| Run1_MDS25 | | -0.39 | -0.21 | -0.38 | -0.42 | -0.32 | -0.41 | -0.42 | -0.43 | -0.45 | -0.35 | -0.38 | -0.44 | -0.39 | -0.34 | -0.22 | -0.2 | -0.36 | 50.96 | -0.59 | -0.45 | -0.38 | -0.41 | -0.48 | -0.36 | -0.45 | -0.46 |
| Run1_MDS5 | | -0.38 | -0.15 | -0.6 | -0.13 | -0.17 | -0.28 | -0.36 | -0.38 | -0.35 | -0.21 | -0.34 | -0.4 | -0.39 | -0.35 | -0.22 | -0.22 | -0.33 | 76.15 | -0.43 | -0.38 | -0.39 | -0.35 | -0.45 | -0.21 | -0.29 | -0.4 |
| Run1_MDS6 | | -0.36 | -0.4 | -0.44 | -0.42 | -0.45 | -0.4 | -0.4 | -0.46 | -0.46 | -0.36 | -0.36 | -0.41 | -0.36 | -0.2 | -0.33 | -0.34 | 0.32 | 58.68 | -0.49 | -0.41 | -0.46 | -0.42 | -0.47 | -0.32 | -0.36 | -0.45 |
| Run1_MDS8 | | -0.4 | -0.41 | -0.5 | -0.42 | -0.4 | -0.4 | -0.39 | -0.46 | -0.44 | -0.37 | -0.41 | -0.43 | -0.41 | -0.28 | -0.22 | -0.32 | -0.31 | 60.93 | -0.39 | -0.46 | -0.45 | -0.27 | -0.48 | -0.32 | -0.41 | -0.48 |
| Run1_MDS9 | | -0.41 | -0.39 | -0.36 | -0.42 | -0.38 | -0.39 | -0.44 | -0.42 | -0.37 | -0.36 | -0.38 | -0.45 | -0.43 | -0.36 | -0.31 | -0.33 | -0.3 | 86.86 | -0.43 | -0.42 | -0.45 | -0.6 | -0.45 | -0.3 | -0.33 | -0.47 |
| Run2_MDS12 | Lower Risk | -0.37 | -0.4 | -0.2 | -0.4 | -0.35 | -0.35 | -0.34 | -0.37 | -0.38 | -0.39 | -0.38 | -0.4 | -0.39 | -0.1 | -0.26 | -0.32 | -0.26 | 152.79 | -0.43 | -0.22 | -0.28 | -0.41 | -0.41 | -0.32 | -0.06 | -0.3 |
| Run2_MDS14 | | -0.33 | -0.18 | -0.49 | 0.86 | -0.05 | -0.37 | -0.46 | -0.4 | -0.4 | -0.34 | -0.35 | -0.36 | -0.34 | -0.2 | -0.29 | -0.25 | 0.83 | 109.08 | -0.35 | -0.35 | -0.4 | -0.29 | -0.4 | -0.34 | -0.3 | -0.36 |
| Run2_MDS19 | | -0.36 | -0.32 | -0.25 | -0.32 | -0.33 | -0.33 | -0.33 | -0.36 | -0.41 | -0.31 | -0.31 | -0.38 | -0.38 | 0 | -0.23 | -0.32 | -0.26 | 126.91 | -0.49 | -0.36 | -0.4 | -0.44 | -0.46 | -0.31 | -0.25 | -0.38 |
| Run2_MDS20 | | -0.37 | -0.54 | -0.61 | -0.41 | -0.4 | -0.4 | -0.38 | -0.44 | -0.42 | -0.27 | -0.29 | -0.37 | -0.36 | 0.04 | -0.02 | -0.31 | -0.31 | 188.42 | -0.39 | -0.39 | -0.38 | -0.37 | -0.41 | -0.27 | -0.35 | -0.4 |
| Run2_MDS28* | | -0.4 | -0.42 | -0.32 | -0.44 | -0.42 | -0.41 | -0.41 | -0.42 | -0.41 | -0.39 | -0.39 | -0.41 | -0.41 | -0.3 | -0.29 | -0.4 | -0.32 | 72.19 | -0.25 | -0.32 | -0.43 | -0.41 | -0.4 | -0.24 | -0.23 | -0.37 |
| Run2_MDS5 | | -0.34 | -0.15 | -0.23 | -0.28 | -0.31 | -0.4 | -0.5 | -0.38 | -0.11 | -0.15 | -0.33 | -0.38 | -0.34 | -0.06 | -0.13 | -0.24 | -0.29 | 115.83 | -0.29 | -0.35 | -0.34 | -0.35 | -0.4 | -0.18 | -0.26 | -0.39 |
| Run1_ICUS18 | | -0.4 | -0.32 | -0.36 | -0.32 | -0.38 | -0.43 | -0.44 | -0.48 | -0.46 | -0.31 | -0.31 | -0.41 | -0.4 | -0.19 | -0.23 | -0.36 | -0.31 | 78.81 | -0.56 | -0.45 | -0.47 | -0.49 | -0.48 | -0.3 | -0.38 | -0.46 |
| Run1_ICUS22 | ICUS | -0.44 | -0.32 | -0.3 | -0.32 | -0.3 | -0.43 | -0.46 | -0.46 | -0.48 | -0.41 | -0.4 | -0.46 | -0.46 | -0.36 | -0.35 | -0.41 | 0.06 | 87.69 | -0.36 | -0.41 | -0.43 | -0.45 | -0.5 | -0.33 | -0.28 | -0.45 |
| Run1_ICUS24 | | -0.38 | -0.38 | -0.37 | -0.45 | -0.29 | -0.31 | -0.39 | -0.44 | -0.48 | -0.4 | -0.36 | -0.42 | -0.42 | -0.38 | -0.39 | -0.35 | 1.3 | 66.04 | -0.39 | -0.44 | -0.47 | -0.49 | -0.45 | -0.33 | -0.41 | -0.45 |
| Run1_Nl-1 | | -0.39 | -0.39 | -0.31 | -0.53 | -0.28 | -0.34 | -0.39 | -0.47 | -0.46 | -0.36 | -0.35 | -0.42 | -0.42 | -0.32 | -0.35 | -0.35 | 1.63 | 74.29 | -0.47 | -0.46 | -0.47 | -0.39 | -0.33 | -0.3 | -0.43 | -0.47 |
| Run1_Nl-3 | | -0.27 | -0.38 | -0.37 | -0.44 | -0.28 | -0.33 | -0.34 | -0.39 | -0.45 | -0.19 | -0.22 | -0.3 | -0.25 | -0.1 | -0.21 | -0.24 | 0.41 | 48.86 | -0.5 | -0.46 | -0.47 | -0.43 | -0.46 | -0.44 | -0.44 | -0.47 |
| Run1_Nl-4 | | -0.24 | -0.42 | -0.49 | -0.39 | -0.38 | -0.31 | -0.27 | -0.37 | -0.42 | -0.13 | -0.16 | -0.26 | -0.21 | -0.11 | -0.18 | -0.18 | 1.07 | 37.98 | -0.46 | -0.44 | -0.45 | -0.34 | -0.45 | -0.45 | -0.43 | -0.45 |
| Run1_Nl-6 | | -0.31 | -0.45 | -0.49 | -0.46 | -0.3 | -0.28 | -0.35 | -0.36 | -0.43 | -0.24 | -0.26 | -0.36 | -0.34 | -0.15 | -0.28 | -0.32 | 1.75 | 39.6 | -0.42 | -0.44 | -0.44 | -0.39 | -0.44 | -0.45 | -0.38 | -0.46 |
| Run2_Nl-3 | Normal | -0.25 | -0.36 | -0.37 | -0.38 | -0.32 | -0.32 | -0.27 | -0.35 | -0.38 | -0.18 | -0.22 | -0.27 | -0.22 | -0.09 | -0.09 | -0.23 | 0.47 | 58.55 | -0.41 | -0.4 | -0.41 | -0.37 | -0.39 | -0.38 | -0.38 | -0.41 |
| Run2_Nl-4 | | -0.21 | -0.34 | -0.36 | -0.32 | -0.23 | -0.29 | -0.27 | -0.35 | -0.37 | -0.13 | -0.16 | -0.23 | -0.18 | -0.01 | -0.05 | -0.19 | 0.7 | 46.52 | -0.35 | -0.39 | -0.43 | -0.31 | -0.42 | -0.38 | -0.39 | -0.4 |
| Run2_Nl-5 | | -0.25 | -0.27 | -0.32 | -0.22 | -0.3 | -0.25 | -0.29 | -0.35 | -0.36 | -0.13 | -0.2 | -0.29 | -0.23 | -0.08 | -0.14 | -0.23 | 1.8 | 65.94 | -0.36 | -0.41 | -0.42 | -0.3 | -0.4 | -0.36 | -0.4 | -0.41 |
| Run2_Nl-6 | | -0.29 | -0.38 | -0.42 | -0.37 | -0.31 | -0.3 | -0.28 | -0.38 | -0.37 | -0.21 | -0.23 | -0.33 | -0.31 | 0.04 | -0.17 | -0.26 | 1.76 | 48.04 | -0.39 | -0.37 | -0.4 | -0.36 | -0.43 | -0.29 | -0.29 | -0.4 |

**Supplemental Table 3**: Median expression of each surface marker on each cell population.

### CD69 Median

| | | Ungated | CD34+CD38low | HSCs | MPP | CMP/GMP | Myelo/Mono-Blasts | ProMonocytes | CD14neg Monocytes | Mature Monocytes | ProMyelocytes | Myelocytes | MetaMyelocytes | MatureGrans | ProErythroblasts | Erythroblasts | LateErythroblasts | PreBcells | Mature Bcells | Plasma Cells | T cells | NK cells | pDCs | Basophils | Platelets | CD8+ T cells | CD8neg T cells |
|---|---|---|---|---|---|---|---|---|---|---|---|---|---|---|---|---|---|---|---|---|---|---|---|---|---|---|---|
| Run1_MDS17 | | -0.27 | -0.13 | -0.05 | -0.15 | 0.06 | -0.25 | -0.24 | -0.25 | -0.28 | -0.4 | -0.34 | -0.31 | -0.25 | -0.37 | -0.44 | -0.31 | -0.18 | -0.02 | -0.53 | -0.15 | -0.06 | -0.3 | -0.21 | 13.4 | -0.06 | -0.24 |
| Run2_MDS15* | AML/RAEB-T | -0.25 | -0.02 | -0.15 | -0.21 | -0.14 | -0.16 | -0.24 | -0.35 | -0.28 | -0.25 | -0.26 | -0.29 | -0.36 | -0.26 | -0.27 | -0.13 | -0.23 | -0.04 | -0.52 | -0.11 | 0.34 | -0.08 | -0.08 | 1.41 | 0.21 | -0.23 |
| Run2_MDS4 | | -0.23 | 1.44 | 1.03 | 1.53 | 4.27 | -0.06 | -0.22 | -0.31 | -0.17 | -0.35 | -0.36 | -0.38 | -0.39 | -0.36 | -0.34 | 0.05 | 0.83 | 1.38 | -0.39 | 0.14 | 0.68 | 2.64 | 0.17 | 22.34 | 0.32 | 0.01 |
| Run1_MDS21 | | -0.3 | -0.14 | -0.06 | -0.17 | -0.13 | -0.16 | -0.22 | -0.26 | -0.33 | -0.43 | -0.4 | -0.44 | -0.41 | -0.43 | -0.36 | 25.33 | 0.38 | -0.11 | -0.5 | -0.26 | 1.42 | -0.32 | 0.22 | 37.56 | -0.15 | -0.3 |
| Run1_MDS3 | | -0.32 | 0.04 | -0.01 | 0.02 | 0.12 | 0.03 | -0.13 | -0.26 | -0.1 | -0.44 | -0.42 | -0.45 | -0.34 | -0.43 | -0.37 | 0.79 | 0.45 | 0.07 | -0.42 | -0.21 | 0.97 | -0.34 | 0.56 | 31.93 | -0.11 | -0.25 |
| Run2_MDS13 | | -0.31 | -0.1 | -0.04 | -0.12 | -0.1 | -0.15 | -0.3 | -0.33 | -0.02 | -0.4 | -0.37 | -0.39 | -0.36 | -0.34 | -0.4 | -0.3 | 0.34 | -0.03 | -0.36 | -0.2 | 0.94 | -0.39 | -0.15 | 28.9 | -0.15 | -0.22 |
| Run2_MDS16 | | -0.33 | 1 | 1.19 | 0.85 | 0.62 | -0.21 | -0.36 | -0.35 | -0.29 | -0.33 | -0.39 | -0.41 | -0.41 | 0.36 | -0.41 | -0.22 | -0.15 | -0.08 | -0.43 | -0.22 | -0.11 | -0.23 | -0.13 | 28.5 | -0.12 | -0.24 |
| Run2_MDS1 | | -0.23 | 0.58 | 1.64 | 0.48 | 0.38 | -0.03 | 0.11 | -0.25 | -0.23 | -0.38 | -0.4 | -0.39 | -0.38 | -0.31 | -0.37 | -0.24 | 0.31 | 0.49 | -0.36 | 0.01 | 0.51 | 0.02 | 0.17 | 19.83 | 1.89 | -0.08 |
| Run2_MDS21 | Higher Risk | -0.27 | -0.14 | -0.17 | -0.14 | -0.09 | -0.16 | -0.43 | -0.26 | -0.21 | -0.38 | -0.36 | -0.39 | -0.39 | -0.38 | -0.41 | -0.29 | 0.46 | 0.19 | -0.27 | -0.21 | 0.93 | 1.14 | -0.15 | 43.37 | -0.1 | -0.25 |
| Run2_MDS26* | | -0.37 | 1.64 | 2.6 | 1.58 | 0.89 | -0.09 | -0.32 | -0.34 | -0.35 | -0.36 | -0.37 | -0.38 | -0.36 | -0.26 | -0.42 | -0.39 | -0.1 | 0.03 | -0.41 | 0 | 0.43 | -0.32 | 0.26 | -0.23 | 0.87 | -0.15 |
| Run2_MDS27 | | -0.23 | 1.72 | 2.01 | 1.39 | 1.25 | 0.55 | -0.16 | -0.43 | -0.35 | -0.32 | -0.34 | -0.41 | -0.42 | 1.04 | -0.41 | -0.39 | 0.43 | 0.01 | -0.4 | 0.01 | -0.02 | -0.46 | 0.35 | 13.43 | 0.23 | -0.07 |
| Run2_MDS2 | | -0.33 | -0.3 | -0.12 | -0.32 | -0.32 | -0.33 | -0.35 | -0.38 | -0.22 | -0.3 | -0.36 | -0.39 | -0.39 | -0.22 | -0.24 | 0.44 | -0.24 | 0.24 | -0.29 | -0.23 | -0.14 | -0.37 | -0.44 | 41.68 | -0.12 | -0.23 |
| Run2_MDS3 | | -0.29 | 0.31 | 0.45 | 0.24 | 0.36 | 0.01 | -0.22 | -0.29 | -0.1 | -0.4 | -0.39 | -0.38 | -0.39 | -0.31 | -0.41 | -0.4 | 0.63 | 0.38 | -0.32 | -0.19 | 0.66 | 0.44 | 0.81 | 34.67 | -0.11 | -0.22 |
| Run1_MDS23 | | -0.38 | 0.71 | 1.4 | 0.73 | 0.36 | -0.33 | -0.32 | -0.37 | -0.3 | -0.38 | -0.41 | -0.43 | -0.44 | -0.42 | -0.38 | -0.01 | -0.14 | 0.14 | -0.36 | 0.02 | 1.52 | -0.34 | -0.19 | 17.95 | 1.02 | -0.1 |
| Run1_MDS25 | | -0.41 | 0.59 | 1.18 | -0.27 | 1.01 | -0.43 | -0.38 | -0.39 | -0.34 | -0.42 | -0.4 | -0.46 | -0.45 | -0.43 | -0.39 | -0.04 | -0.18 | -0.13 | -0.35 | -0.07 | 2.91 | -0.32 | -0.08 | 16.07 | -0.04 | -0.1 |
| Run1_MDS5 | | -0.18 | 1.14 | -0.19 | 1.05 | 0.75 | 0.12 | 0.04 | -0.09 | 0.28 | -0.25 | -0.31 | -0.34 | -0.31 | -0.35 | -0.28 | 7.34 | 0.14 | 0.16 | -0.45 | 0.16 | 3.72 | 1.78 | 1.13 | 13.77 | 0.53 | 0.04 |
| Run1_MDS6 | | -0.39 | 0.51 | 0.65 | -0.02 | 0.43 | -0.2 | -0.26 | -0.31 | -0.14 | -0.42 | -0.43 | -0.43 | -0.44 | -0.35 | -0.41 | -0.09 | -0.16 | -0.06 | -0.42 | -0.2 | -0.08 | -0.13 | -0.23 | 19.92 | -0.05 | -0.29 |
| Run1_MDS8 | | -0.38 | 1.5 | 1.69 | 0.7 | 0.52 | -0.16 | -0.14 | -0.11 | 0.13 | -0.43 | -0.41 | -0.46 | -0.44 | -0.41 | -0.4 | 0.01 | -0.18 | -0.01 | -0.32 | -0.27 | -0.1 | 0.3 | 0.51 | 25.13 | -0.08 | -0.32 |
| Run1_MDS9 | | -0.34 | 1.11 | 1.91 | 0.72 | 1.01 | 0.15 | 0.1 | 0.37 | 0.64 | -0.4 | -0.41 | -0.45 | -0.46 | -0.34 | -0.37 | 4.59 | -0.05 | 0.11 | -0.3 | -0.22 | 1.01 | -0.12 | 1.02 | 19.1 | 0.03 | -0.31 |
| Run2_MDS12 | Lower Risk | -0.35 | 0.07 | 1.73 | 0.36 | 0.87 | -0.26 | -0.18 | -0.22 | -0.12 | -0.35 | -0.36 | -0.37 | -0.38 | -0.37 | -0.41 | -0.35 | -0.07 | 0.01 | -0.25 | 2.23 | 9.14 | -0.26 | 0.39 | 10.66 | 12.56 | 0.79 |
| Run2_MDS14 | | -0.38 | 0.02 | -0.64 | 0.43 | 0.51 | -0.33 | -0.39 | -0.4 | -0.3 | -0.36 | -0.39 | -0.4 | -0.41 | -0.34 | -0.41 | -0.32 | -0.1 | 0 | -0.61 | -0.11 | -0.05 | -0.43 | -0.04 | 9.69 | 0.19 | -0.18 |
| Run2_MDS19 | | -0.38 | 1.04 | 1.76 | 0.98 | 0.41 | -0.29 | -0.34 | -0.35 | -0.34 | -0.36 | -0.37 | -0.4 | -0.41 | -0.35 | -0.39 | -0.4 | -0.15 | 0 | -0.48 | -0.09 | 0.22 | -0.31 | -0.17 | 26.22 | 0.72 | -0.17 |
| Run2_MDS20 | | -0.29 | 1.07 | 2.02 | 0.73 | 0.43 | -0.35 | -0.38 | -0.36 | -0.31 | -0.34 | -0.34 | -0.41 | -0.41 | -0.35 | -0.37 | -0.23 | 0.08 | 0.28 | -0.3 | 0.14 | 0.27 | -0.37 | 0.15 | 25.1 | -0.01 | 0.16 |
| Run2_MDS28* | | -0.35 | 0.48 | 1.37 | 0.44 | 0.54 | -0.28 | -0.31 | -0.36 | -0.28 | -0.37 | -0.35 | -0.38 | -0.39 | -0.34 | -0.39 | -0.38 | -0.12 | -0.01 | -0.28 | -0.02 | 0.04 | -0.06 | 0.29 | 55.48 | 0.33 | -0.13 |
| Run2_MDS5 | | -0.23 | 0.48 | 0.02 | 0.66 | 0.45 | -0.02 | -0.25 | -0.12 | -0.07 | -0.26 | -0.35 | -0.32 | -0.29 | -0.43 | -0.36 | 1.08 | -0.07 | -0.03 | -0.4 | 0 | 3.38 | 0.37 | 0.86 | 15.12 | 0.16 | -0.04 |
| Run1_ICUS18 | | -0.41 | 0.12 | 0.32 | -0.13 | 0.56 | -0.42 | -0.38 | -0.39 | -0.35 | -0.38 | -0.41 | -0.44 | -0.47 | -0.41 | -0.4 | -0.39 | -0.12 | 0.03 | -0.32 | -0.22 | 0.18 | -0.45 | -0.29 | 14.8 | -0.04 | -0.25 |
| Run1_ICUS22 | ICUS | -0.42 | 1.24 | 2.15 | 0.94 | 0.75 | -0.37 | -0.41 | -0.41 | -0.33 | -0.4 | -0.41 | -0.46 | -0.48 | -0.44 | -0.42 | -0.34 | 0.04 | 0.08 | -0.38 | 0.14 | 2.04 | -0.44 | 0.41 | 14.67 | 0.6 | -0.02 |
| Run1_ICUS24 | | -0.37 | 1.4 | 1.87 | 1.27 | 0.5 | -0.22 | -0.3 | -0.41 | -0.33 | -0.4 | -0.4 | -0.43 | -0.45 | -0.39 | -0.43 | 2.35 | -0.01 | -0.02 | -0.45 | -0.14 | 0.19 | -0.29 | 0.41 | 20.49 | -0.02 | -0.22 |
| Run1_Nl-1 | | -0.38 | 1.44 | 0.75 | 1.33 | 1 | -0.19 | -0.26 | -0.3 | -0.26 | -0.38 | -0.39 | -0.43 | -0.45 | -0.38 | -0.43 | -0.15 | 0.4 | 0.18 | -0.44 | -0.16 | 0.35 | -0.31 | 2.76 | 29.87 | -0.04 | -0.21 |
| Run1_Nl-3 | | -0.37 | 0.72 | 0.86 | 0.68 | 0.23 | -0.28 | -0.32 | -0.35 | -0.31 | -0.38 | -0.39 | -0.41 | -0.42 | -0.35 | -0.42 | -0.33 | -0.16 | -0.06 | -0.49 | -0.25 | -0.04 | -0.39 | 0.14 | 4.89 | -0.1 | -0.33 |
| Run1_Nl-4 | | -0.38 | 0.4 | 1.4 | 0.8 | 0.17 | -0.26 | -0.33 | -0.38 | -0.33 | -0.37 | -0.38 | -0.41 | -0.42 | -0.34 | -0.42 | -0.36 | -0.13 | -0.12 | -0.43 | -0.22 | -0.12 | -0.37 | 0.36 | 5.29 | -0.18 | -0.25 |
| Run1_Nl-6 | | -0.39 | 1 | 1.38 | 0.88 | 0.28 | -0.25 | -0.3 | -0.37 | -0.3 | -0.37 | -0.39 | -0.43 | -0.43 | -0.35 | -0.44 | -0.36 | -0.06 | -0.06 | -0.46 | -0.15 | -0.05 | -0.35 | -0.07 | 6.45 | 0.5 | -0.23 |
| Run2_Nl-3 | Normal | -0.36 | 0.33 | 0.83 | 0.08 | -0.01 | -0.29 | -0.29 | -0.33 | -0.3 | -0.35 | -0.37 | -0.39 | -0.38 | -0.36 | -0.41 | -0.36 | -0.19 | -0.13 | -0.4 | -0.26 | -0.08 | -0.39 | -0.01 | 5.66 | -0.16 | -0.32 |
| Run2_Nl-4 | | -0.36 | 0.41 | 0.91 | 0.15 | -0.05 | -0.28 | -0.28 | -0.35 | -0.33 | -0.33 | -0.36 | -0.39 | -0.38 | -0.32 | -0.39 | -0.38 | -0.08 | -0.15 | -0.42 | -0.22 | -0.15 | -0.3 | 0.29 | 6.07 | -0.21 | -0.23 |
| Run2_Nl-5 | | -0.34 | 0.37 | 0.31 | 0.29 | 0.26 | -0.26 | -0.27 | -0.35 | -0.32 | -0.33 | -0.37 | -0.4 | -0.39 | -0.3 | -0.4 | -0.38 | -0.02 | -0.07 | -0.42 | -0.24 | -0.09 | -0.35 | -0.08 | 8.48 | -0.21 | -0.27 |
| Run2_Nl-6 | | -0.38 | 0.68 | 1 | 0.64 | 0.26 | -0.24 | -0.31 | -0.37 | -0.31 | -0.35 | -0.38 | -0.4 | -0.4 | -0.33 | -0.4 | -0.39 | -0.09 | -0.1 | -0.43 | -0.16 | -0.04 | -0.33 | -0.07 | 8.06 | 0.51 | -0.24 |

**Supplemental Table 4**: Most common surface marker aberrancies by cell type.
Number indicates the number of samples (of 23 in total; from MDS and sAML) which demonstrated aberrant expression within the indicated group of cell populations (columns) for each marker (rows).
HSPC = HSC, MPP and CD33+MPP (all CD34+CD38low).
Boxes are colored from green (no aberrancy) to red (maximum for each cell type).
The final column on the right is a sum of the first four groups with a possible score of up to 92 (23x4).
Note that the analysis includes samples MDS3, MDS13, and MDS21 which came from serial biopsies of the same patient (each several months apart) and demonstrate consistent properties.

| | HSPC | Blasts | Monocyte Lineage | Granulocyte Lineage | Other | Total: Progenitor, Monocyte, and Granulocyte |
|---|---|---|---|---|---|---|
| **CD45RA** | 2 | 0 | 0 | 0 | 1 | 2 |
| **CD7** | 3 | 0 | 0 | 0 | 14 | 3 |
| **CD71** | 1 | 2 | 0 | 1 | 19 | 4 |
| **CD235** | 4 | 0 | 0 | 0 | 16 | 4 |
| **CD47** | 8 | 1 | 2 | 6 | 10 | 17 |
| **CD8** | 4 | 0 | 0 | 1 | 9 | 5 |
| **CD34** | 16 | 8 | 1 | 0 | 6 | 25 |
| **CD117** | 11 | 5 | 0 | 0 | 3 | 16 |
| **CD56** | 0 | 0 | 2 | 2 | 8 | 4 |
| **CD90** | 3 | 0 | 0 | 0 | 0 | 3 |
| **CD33** | 3 | 13 | 1 | 2 | 7 | 19 |
| **CD64** | 5 | 2 | 2 | 5 | 5 | 14 |
| **CD16** | 5 | 0 | 3 | 3 | 11 | 11 |
| **CD11b** | 6 | 0 | 8 | 8 | 19 | 22 |
| **CD15** | 5 | 0 | 0 | 18 | 18 | 23 |
| **CD123** | 2 | 0 | 0 | 1 | 10 | 3 |
| **CD3** | 0 | 0 | 0 | 0 | 6 | 0 |
| **CD45** | 11 | 4 | 3 | 2 | 19 | 20 |
| **CD133** | 1 | 0 | 0 | 0 | 0 | 1 |
| **HLADR** | 13 | 1 | 12 | 2 | 14 | 28 |
| **CD44** | 11 | 10 | 6 | 9 | 20 | 36 |
| **CD38** | 3 | 0 | 15 | 3 | 13 | 21 |
| **CD14** | 1 | 0 | 0 | 0 | 4 | 1 |
| **Calreticulin** | 1 | 0 | 0 | 0 | 3 | 1 |
| **CD321** | 7 | 10 | 11 | 5 | 20 | 33 |
| **CD99** | 4 | 5 | 1 | 2 | 18 | 12 |
| **CD13** | 0 | 0 | 2 | 1 | 4 | 3 |
| **CXCR4** | 1 | 0 | 0 | 2 | 8 | 3 |
| **CD10** | 3 | 0 | 0 | 1 | 0 | 4 |
| **CD19** | 0 | 0 | 0 | 0 | 4 | 0 |
| **CD20** | 0 | 0 | 0 | 0 | 5 | 0 |
| **CD69** | 0 | 0 | 0 | 0 | 7 | 0 |

**Supplemental Table 5**: Fequency of surface marker aberrancies by sample. Number of marker abberrancies is shown for each marker and population (HSPC, Blasts, Monocytes, Granulocytes, and Other), in each sample.
Samples MDS3, 13 & 21 are serial samples from the same patient each several months apart.

| | | AML/RAEB-T | | | Higher Risk MDS | | | | | | | |
|---|---|---|---|---|---|---|---|---|---|---|---|---|
| | | MDS17 | MDS15 | MDS4 | MDS13 | MDS16 | MDS1 | MDS21 | MDS26 | MDS27 | MDS2 | MDS3 |
| **CD45RA Median** | **HSPC** | 0 | 0 | 0 | 0 | 0 | 0 | 0 | 0 | 0 | 0 | 0 |
| | **Blasts** | 0 | 0 | 0 | 0 | 0 | 0 | 0 | 0 | 0 | 0 | 0 |
| | **Mono** | 0 | 0 | 0 | 0 | 0 | 0 | 0 | 0 | 0 | 0 | 0 |
| | **Gran** | 0 | 0 | 0 | 0 | 0 | 0 | 0 | 0 | 0 | 0 | 0 |
| | **Other** | 0 | 0 | 0 | 0 | 0 | 0 | 0 | 0 | 0 | 0 | 0 |
| **CD7 Median** | **HSPC** | 0 | 0 | 3 | 0 | 0 | 0 | 0 | 0 | 0 | 0 | 0 |
| | **Blasts** | 0 | 0 | 0 | 0 | 0 | 0 | 0 | 0 | 0 | 0 | 0 |
| | **Mono** | 0 | 0 | 0 | 0 | 0 | 0 | 0 | 0 | 0 | 0 | 0 |
| | **Gran** | 0 | 0 | 0 | 0 | 0 | 0 | 0 | 0 | 0 | 0 | 0 |
| | **Other** | 2 | 0 | 1 | 0 | 4 | 1 | 2 | 0 | 1 | 2 | 1 |
| **CD71 Median** | **HSPC** | 0 | 0 | 0 | 0 | 0 | 0 | 0 | 0 | 0 | 0 | 0 |
| | **Blasts** | 0 | 0 | 0 | 0 | 0 | 0 | 0 | 0 | 0 | 1 | 0 |
| | **Mono** | 0 | 0 | 0 | 0 | 0 | 0 | 0 | 0 | 0 | 0 | 0 |
| | **Gran** | 0 | 0 | 0 | 0 | 0 | 0 | 0 | 0 | 0 | 0 | 0 |
| | **Other** | 2 | 1 | 0 | 3 | 2 | 1 | 1 | 1 | 2 | 1 | 1 |
| **CD235 Median** | **HSPC** | 0 | 0 | 0 | 0 | 0 | 0 | 0 | 0 | 0 | 0 | 0 |
| | **Blasts** | 0 | 0 | 0 | 0 | 0 | 0 | 0 | 0 | 0 | 0 | 0 |
| | **Mono** | 0 | 0 | 0 | 0 | 0 | 0 | 0 | 0 | 0 | 0 | 0 |
| | **Gran** | 0 | 0 | 0 | 0 | 0 | 0 | 0 | 0 | 0 | 0 | 0 |
| | **Other** | 1 | 1 | 2 | 1 | 2 | 1 | 1 | 1 | 2 | 1 | 1 |
| **CD47 Median** | **HSPC** | 0 | 0 | 0 | 2 | 0 | 1 | 2 | 0 | 0 | 2 | 1 |
| | **Blasts** | 0 | 0 | 0 | 0 | 0 | 0 | 0 | 0 | 0 | 0 | 0 |
| | **Mono** | 0 | 0 | 0 | 0 | 0 | 0 | 0 | 0 | 0 | 0 | 0 |
| | **Gran** | 1 | 0 | 0 | 0 | 0 | 1 | 0 | 0 | 0 | 0 | 0 |
| | **Other** | 0 | 2 | 1 | 1 | 0 | 0 | 1 | 1 | 0 | 1 | 1 |
| **CD8 Median** | **HSPC** | 0 | 0 | 0 | 0 | 0 | 0 | 0 | 0 | 0 | 0 | 0 |
| | **Blasts** | 0 | 0 | 0 | 0 | 0 | 0 | 0 | 0 | 0 | 0 | 0 |
| | **Mono** | 0 | 0 | 0 | 0 | 0 | 0 | 0 | 0 | 0 | 0 | 0 |
| | **Gran** | 3 | 0 | 0 | 0 | 0 | 0 | 0 | 0 | 0 | 0 | 0 |
| | **Other** | 1 | 2 | 1 | 0 | 0 | 0 | 1 | 0 | 1 | 0 | 0 |
| **CD34 Median** | **HSPC** | 1 | 3 | 0 | 0 | 0 | 2 | 1 | 2 | 0 | 0 | 0 |
| | **Blasts** | 1 | 1 | 0 | 1 | 0 | 0 | 1 | 0 | 1 | 1 | 1 |
| | **Mono** | 0 | 0 | 0 | 0 | 0 | 0 | 0 | 0 | 0 | 0 | 0 |
| | **Gran** | 0 | 0 | 0 | 0 | 0 | 0 | 0 | 0 | 0 | 0 | 0 |
| | **Other** | 1 | 4 | 0 | 0 | 1 | 0 | 1 | 0 | 1 | 0 | 0 |

**Supplemental Table 5**: Fequency of surface marker aberrancies by sample. Number of marker abberrancies is shown for each marker and population (HSPC, Blasts, Monocytes, Granulocytes, and Other), in each sample. Samples MDS3, 13 & 21 are serial samples from the same patient each several months apart.

| | | AML/RAEB-T | | | Higher Risk MDS | | | | | | | |
|---|---|---|---|---|---|---|---|---|---|---|---|---|
| | | MDS17 | MDS15 | MDS4 | MDS13 | MDS16 | MDS1 | MDS21 | MDS26 | MDS27 | MDS2 | MDS3 |
| CD117 Median | HSPC | 4 | 4 | 3 | 0 | 0 | 3 | 3 | 4 | 0 | 0 | 0 |
| | Blasts | 1 | 0 | 0 | 1 | 0 | 0 | 1 | 1 | 0 | 0 | 1 |
| | Mono | 0 | 0 | 0 | 0 | 0 | 0 | 0 | 0 | 0 | 0 | 0 |
| | Gran | 0 | 0 | 0 | 0 | 0 | 0 | 0 | 0 | 0 | 0 | 0 |
| | Other | 0 | 3 | 0 | 0 | 0 | 0 | 1 | 0 | 0 | 0 | 0 |
| CD56 Median | HSPC | 0 | 0 | 0 | 0 | 0 | 0 | 0 | 0 | 0 | 0 | 0 |
| | Blasts | 0 | 0 | 0 | 0 | 0 | 0 | 0 | 0 | 0 | 0 | 0 |
| | Mono | 0 | 0 | 0 | 0 | 0 | 0 | 0 | 0 | 0 | 0 | 0 |
| | Gran | 0 | 1 | 0 | 0 | 0 | 0 | 0 | 0 | 0 | 0 | 0 |
| | Other | 0 | 1 | 0 | 0 | 0 | 1 | 0 | 0 | 0 | 0 | 0 |
| CD90 Median | HSPC | 0 | 0 | 0 | 0 | 0 | 0 | 0 | 0 | 0 | 0 | 0 |
| | Blasts | 0 | 0 | 0 | 0 | 0 | 0 | 0 | 0 | 0 | 0 | 0 |
| | Mono | 0 | 0 | 0 | 0 | 0 | 0 | 0 | 0 | 0 | 0 | 0 |
| | Gran | 0 | 0 | 0 | 0 | 0 | 0 | 0 | 0 | 0 | 0 | 0 |
| | Other | 0 | 0 | 0 | 0 | 0 | 0 | 0 | 0 | 0 | 0 | 0 |
| CD33 Median | HSPC | 0 | 2 | 2 | 0 | 0 | 0 | 0 | 0 | 0 | 0 | 0 |
| | Blasts | 1 | 0 | 1 | 1 | 0 | 1 | 1 | 1 | 1 | 1 | 1 |
| | Mono | 0 | 0 | 1 | 0 | 0 | 0 | 0 | 0 | 0 | 0 | 0 |
| | Gran | 0 | 3 | 1 | 0 | 0 | 0 | 0 | 0 | 0 | 0 | 0 |
| | Other | 0 | 6 | 0 | 1 | 0 | 0 | 0 | 0 | 0 | 0 | 0 |
| CD64 Medians | HSPC | 0 | 0 | 0 | 0 | 0 | 1 | 0 | 0 | 0 | 0 | 0 |
| | Blasts | 0 | 0 | 0 | 0 | 0 | 0 | 0 | 0 | 0 | 0 | 0 |
| | Mono | 0 | 0 | 0 | 0 | 0 | 0 | 0 | 0 | 0 | 3 | 0 |
| | Gran | 1 | 0 | 1 | 0 | 0 | 0 | 0 | 0 | 0 | 1 | 0 |
| | Other | 0 | 1 | 0 | 0 | 0 | 0 | 0 | 0 | 0 | 2 | 1 |
| CD16 Median | HSPC | 0 | 0 | 0 | 0 | 0 | 0 | 0 | 0 | 0 | 0 | 0 |
| | Blasts | 0 | 0 | 0 | 0 | 0 | 0 | 0 | 0 | 0 | 0 | 0 |
| | Mono | 0 | 1 | 0 | 0 | 0 | 0 | 0 | 0 | 0 | 0 | 0 |
| | Gran | 0 | 0 | 0 | 0 | 0 | 0 | 0 | 0 | 0 | 0 | 0 |
| | Other | 0 | 1 | 0 | 0 | 1 | 0 | 0 | 2 | 2 | 1 | 0 |
| CD11b Median | HSPC | 0 | 0 | 0 | 0 | 0 | 0 | 0 | 0 | 0 | 0 | 0 |
| | Blasts | 0 | 0 | 0 | 0 | 0 | 0 | 0 | 0 | 0 | 0 | 0 |
| | Mono | 0 | 1 | 0 | 0 | 0 | 2 | 1 | 2 | 0 | 1 | 0 |
| | Gran | 1 | 0 | 0 | 2 | 0 | 0 | 3 | 1 | 0 | 1 | 1 |
| | Other | 1 | 3 | 1 | 1 | 0 | 1 | 1 | 1 | 0 | 2 | 1 |

**Supplemental Table 5**: Fequency of surface marker aberrancies by sample. Number of marker abberrancies is shown for each marker and population (HSPC, Blasts, Monocytes, Granulocytes, and Other), in each sample.
Samples MDS3, 13 & 21 are serial samples from the same patient each several months apart.

| | | AML/RAEB-T | | | Higher Risk MDS | | | | | | | |
|---|---|---|---|---|---|---|---|---|---|---|---|---|
| | | **MDS17** | **MDS15** | **MDS4** | **MDS13** | **MDS16** | **MDS1** | **MDS21** | **MDS26** | **MDS27** | **MDS2** | **MDS3** |
| **CD15** | **HSPC** | 0 | 0 | 0 | 0 | 0 | 0 | 0 | 0 | 0 | 0 | 0 |
| **Median** | **Blasts** | 0 | 0 | 0 | 0 | 0 | 0 | 0 | 0 | 0 | 0 | 0 |
| | **Mono** | 0 | 0 | 0 | 0 | 0 | 0 | 0 | 0 | 0 | 0 | 0 |
| | **Gran** | 1 | 2 | 0 | 1 | 1 | 2 | 1 | 1 | 1 | 1 | 1 |
| | **Other** | 1 | 1 | 1 | 1 | 1 | 1 | 1 | 2 | 1 | 1 | 1 |
| **CD123** | **HSPC** | 0 | 0 | 0 | 0 | 0 | 0 | 0 | 0 | 0 | 0 | 0 |
| **Median** | **Blasts** | 0 | 0 | 0 | 0 | 0 | 0 | 0 | 0 | 0 | 0 | 0 |
| | **Mono** | 0 | 0 | 0 | 0 | 0 | 0 | 0 | 0 | 0 | 0 | 0 |
| | **Gran** | 0 | 1 | 0 | 0 | 0 | 0 | 0 | 0 | 0 | 0 | 0 |
| | **Other** | 1 | 1 | 0 | 1 | 0 | 0 | 0 | 1 | 1 | 1 | 0 |
| **CD3** | **HSPC** | 0 | 0 | 0 | 0 | 0 | 0 | 0 | 0 | 0 | 0 | 0 |
| **Median** | **Blasts** | 0 | 0 | 0 | 0 | 0 | 0 | 0 | 0 | 0 | 0 | 0 |
| | **Mono** | 0 | 0 | 0 | 0 | 0 | 0 | 0 | 0 | 0 | 0 | 0 |
| | **Gran** | 0 | 0 | 0 | 0 | 0 | 0 | 0 | 0 | 0 | 0 | 0 |
| | **Other** | 0 | 0 | 0 | 0 | 1 | 0 | 0 | 0 | 1 | 0 | 0 |
| **CD45** | **HSPC** | 1 | 1 | 0 | 2 | 0 | 0 | 3 | 0 | 0 | 2 | 2 |
| **Median** | **Blasts** | 0 | 0 | 0 | 0 | 0 | 0 | 0 | 0 | 0 | 0 | 0 |
| | **Mono** | 0 | 0 | 0 | 0 | 0 | 0 | 0 | 0 | 0 | 0 | 0 |
| | **Gran** | 0 | 0 | 0 | 0 | 0 | 0 | 0 | 0 | 0 | 0 | 0 |
| | **Other** | 1 | 5 | 2 | 2 | 0 | 2 | 4 | 0 | 2 | 7 | 0 |
| **CD133** | **HSPC** | 0 | 0 | 0 | 0 | 0 | 0 | 0 | 0 | 0 | 0 | 0 |
| **Median** | **Blasts** | 0 | 0 | 0 | 0 | 0 | 0 | 0 | 0 | 0 | 0 | 0 |
| | **Mono** | 0 | 0 | 0 | 0 | 0 | 0 | 0 | 0 | 0 | 0 | 0 |
| | **Gran** | 0 | 0 | 0 | 0 | 0 | 0 | 0 | 0 | 0 | 0 | 0 |
| | **Other** | 0 | 0 | 0 | 0 | 0 | 0 | 0 | 0 | 0 | 0 | 0 |
| **HLADR** | **HSPC** | 0 | 2 | 4 | 1 | 0 | 3 | 0 | 0 | 4 | 4 | 0 |
| **Median** | **Blasts** | 0 | 0 | 0 | 0 | 0 | 0 | 0 | 0 | 0 | 0 | 0 |
| | **Mono** | 0 | 0 | 3 | 0 | 0 | 2 | 0 | 0 | 2 | 3 | 0 |
| | **Gran** | 0 | 1 | 0 | 0 | 0 | 0 | 0 | 0 | 0 | 1 | 0 |
| | **Other** | 0 | 5 | 1 | 1 | 1 | 0 | 0 | 0 | 2 | 2 | 0 |
| **CD44** | **HSPC** | 4 | 4 | 0 | 1 | 0 | 3 | 0 | 0 | 0 | 4 | 0 |
| **Median** | **Blasts** | 0 | 0 | 0 | 0 | 1 | 0 | 0 | 1 | 1 | 1 | 0 |
| | **Mono** | 0 | 0 | 1 | 0 | 1 | 0 | 0 | 1 | 0 | 1 | 0 |
| | **Gran** | 2 | 4 | 3 | 3 | 0 | 0 | 1 | 0 | 0 | 2 | 3 |
| | **Other** | 1 | 7 | 4 | 2 | 1 | 3 | 2 | 0 | 3 | 4 | 3 |

**Supplemental Table 5**: Fequency of surface marker aberrancies by sample. Number of marker abberrancies is shown for each marker and population (HSPC, Blasts, Monocytes, Granulocytes, and Other), in each sample. Samples MDS3, 13 & 21 are serial samples from the same patient each several months apart.

| | | AML/RAEB-T | | | Higher Risk MDS | | | | | | | |
|---|---|---|---|---|---|---|---|---|---|---|---|---|
| | | MDS17 | MDS15 | MDS4 | MDS13 | MDS16 | MDS1 | MDS21 | MDS26 | MDS27 | MDS2 | MDS3 |
| **CD38 Median** | **HSPC** | 0 | 1 | 1 | 0 | 0 | 0 | 0 | 0 | 0 | 0 | 0 |
| | **Blasts** | 0 | 0 | 0 | 0 | 0 | 0 | 0 | 0 | 0 | 0 | 0 |
| | **Mono** | 3 | 1 | 2 | 2 | 0 | 2 | 2 | 2 | 0 | 2 | 2 |
| | **Gran** | 0 | 0 | 0 | 3 | 0 | 0 | 3 | 0 | 0 | 2 | 3 |
| | **Other** | 0 | 2 | 2 | 2 | 0 | 0 | 1 | 1 | 0 | 3 | 1 |
| **CD14 Median** | **HSPC** | 0 | 0 | 0 | 0 | 0 | 0 | 0 | 0 | 0 | 0 | 0 |
| | **Blasts** | 0 | 0 | 0 | 0 | 0 | 0 | 0 | 0 | 0 | 0 | 0 |
| | **Mono** | 0 | 0 | 0 | 0 | 0 | 0 | 0 | 0 | 0 | 0 | 0 |
| | **Gran** | 0 | 0 | 0 | 0 | 0 | 0 | 0 | 0 | 0 | 0 | 0 |
| | **Other** | 0 | 2 | 0 | 0 | 0 | 0 | 0 | 0 | 1 | 1 | 0 |
| **CD321 Median** | **HSPC** | 4 | 2 | 0 | 2 | 0 | 0 | 4 | 0 | 0 | 0 | 3 |
| | **Blasts** | 1 | 1 | 0 | 1 | 0 | 1 | 1 | 0 | 1 | 1 | 1 |
| | **Mono** | 3 | 2 | 0 | 3 | 0 | 0 | 1 | 0 | 0 | 0 | 3 |
| | **Gran** | 0 | 2 | 0 | 0 | 0 | 1 | 0 | 1 | 0 | 0 | 0 |
| | **Other** | 3 | 6 | 1 | 1 | 2 | 2 | 1 | 1 | 2 | 6 | 3 |
| **CD99 Median** | **HSPC** | 0 | 1 | 3 | 3 | 0 | 0 | 0 | 0 | 0 | 4 | 0 |
| | **Blasts** | 1 | 0 | 0 | 1 | 0 | 0 | 1 | 0 | 0 | 1 | 1 |
| | **Mono** | 0 | 0 | 0 | 0 | 0 | 1 | 0 | 0 | 0 | 0 | 0 |
| | **Gran** | 0 | 3 | 0 | 0 | 0 | 0 | 0 | 0 | 0 | 1 | 0 |
| | **Other** | 4 | 9 | 4 | 3 | 6 | 2 | 1 | 0 | 8 | 7 | 1 |
| **CD13 Median** | **HSPC** | 0 | 0 | 0 | 0 | 0 | 0 | 0 | 0 | 0 | 0 | 0 |
| | **Blasts** | 0 | 0 | 0 | 0 | 0 | 0 | 0 | 0 | 0 | 0 | 0 |
| | **Mono** | 0 | 0 | 0 | 0 | 0 | 1 | 0 | 0 | 0 | 0 | 0 |
| | **Gran** | 0 | 1 | 0 | 0 | 0 | 0 | 0 | 0 | 0 | 0 | 0 |
| | **Other** | 0 | 2 | 0 | 0 | 0 | 0 | 0 | 0 | 0 | 0 | 0 |
| **CXCR4 Median** | **HSPC** | 0 | 0 | 0 | 0 | 0 | 0 | 0 | 0 | 0 | 0 | 0 |
| | **Blasts** | 0 | 0 | 0 | 0 | 0 | 0 | 0 | 0 | 0 | 0 | 0 |
| | **Mono** | 0 | 0 | 0 | 0 | 0 | 0 | 0 | 0 | 0 | 0 | 0 |
| | **Gran** | 1 | 2 | 0 | 0 | 0 | 0 | 0 | 0 | 0 | 0 | 0 |
| | **Other** | 1 | 0 | 0 | 1 | 0 | 0 | 2 | 0 | 0 | 0 | 2 |
| **CD10 Median** | **HSPC** | 0 | 0 | 0 | 0 | 0 | 0 | 0 | 0 | 0 | 0 | 0 |
| | **Blasts** | 0 | 0 | 0 | 0 | 0 | 0 | 0 | 0 | 0 | 0 | 0 |
| | **Mono** | 0 | 0 | 0 | 0 | 0 | 0 | 0 | 0 | 0 | 0 | 0 |
| | **Gran** | 0 | 0 | 0 | 0 | 0 | 0 | 0 | 0 | 0 | 0 | 0 |
| | **Other** | 0 | 0 | 0 | 0 | 0 | 0 | 0 | 0 | 0 | 0 | 0 |

**Supplemental Table 5**: Fequency of surface marker aberrancies by sample. Number of marker abberrancies is shown for each marker and population (HSPC, Blasts, Monocytes, Granulocytes, and Other), in each sample.
Samples MDS3, 13 & 21 are serial samples from the same patient each several months apart.

| | | AML/RAEB-T | | | | Higher Risk MDS | | | | | | | |
|---|---|---|---|---|---|---|---|---|---|---|---|---|---|
| | | MDS17 | MDS15 | MDS4 | | MDS13 | MDS16 | MDS1 | MDS21 | MDS26 | MDS27 | MDS2 | MDS3 |
| **CD19** | **HSPC** | 0 | 0 | 0 | | 0 | 0 | 0 | 0 | 0 | 0 | 0 | 0 |
| **Median** | **Blasts** | 0 | 0 | 0 | | 0 | 0 | 0 | 0 | 0 | 0 | 0 | 0 |
| | **Mono** | 0 | 0 | 0 | | 0 | 0 | 0 | 0 | 0 | 0 | 0 | 0 |
| | **Gran** | 0 | 0 | 0 | | 0 | 0 | 0 | 0 | 0 | 0 | 0 | 0 |
| | **Other** | 0 | 1 | 0 | | 0 | 0 | 0 | 1 | 0 | 0 | 0 | 1 |
| **CD20** | **HSPC** | 0 | 0 | 0 | | 0 | 0 | 0 | 0 | 0 | 0 | 0 | 0 |
| **Median** | **Blasts** | 0 | 0 | 0 | | 0 | 0 | 0 | 0 | 0 | 0 | 0 | 0 |
| | **Mono** | 0 | 0 | 0 | | 0 | 0 | 0 | 0 | 0 | 0 | 0 | 0 |
| | **Gran** | 0 | 0 | 0 | | 0 | 0 | 0 | 0 | 0 | 0 | 0 | 0 |
| | **Other** | 0 | 1 | 0 | | 0 | 0 | 0 | 0 | 0 | 0 | 0 | 0 |
| **CD69** | **HSPC** | 0 | 0 | 0 | | 0 | 0 | 0 | 0 | 0 | 0 | 0 | 0 |
| **Median** | **Blasts** | 0 | 0 | 0 | | 0 | 0 | 0 | 0 | 0 | 0 | 0 | 0 |
| | **Mono** | 0 | 0 | 0 | | 0 | 0 | 0 | 0 | 0 | 0 | 0 | 0 |
| | **Gran** | 0 | 0 | 0 | | 0 | 0 | 0 | 0 | 0 | 0 | 0 | 0 |
| | **Other** | 0 | 0 | 1 | | 0 | 0 | 0 | 1 | 0 | 0 | 0 | 0 |

**Supplemental Table 5**: Fequency of surface marker aberrancies by sample. Number of marker abberrancies is shown for each marker and population (HSPC, Blasts, Monocytes, Granulocytes, and Other), in each sample.
Samples MDS3, 13 & 21 are serial samples from the same patient each several months apart.

| | | Lower Risk MDS | | | | | | | | | | | | ICUS | | |
|---|---|---|---|---|---|---|---|---|---|---|---|---|---|---|---|---|
| | | MDS23 | MDS25 | MDS5 | MDS6 | MDS8 | MDS9 | MDS12 | MDS14 | MDS19 | MDS20 | MDS28 | MDS11 | ICUS18 | ICUS22 | ICUS24 |
| **CD45RA Median** | **HSPC** | 0 | 1 | 0 | 0 | 1 | 0 | 0 | 0 | 0 | 0 | 0 | 0 | 0 | 0 | 0 |
| | **Blasts** | 0 | 0 | 0 | 0 | 0 | 0 | 0 | 0 | 0 | 0 | 0 | 0 | 0 | 0 | 0 |
| | **Mono** | 0 | 0 | 0 | 0 | 0 | 0 | 0 | 0 | 0 | 0 | 0 | 0 | 0 | 0 | 0 |
| | **Gran** | 0 | 0 | 0 | 0 | 0 | 0 | 0 | 0 | 0 | 0 | 0 | 0 | 0 | 0 | 0 |
| | **Other** | 0 | 0 | 0 | 0 | 1 | 0 | 0 | 0 | 0 | 0 | 0 | 0 | 0 | 0 | 0 |
| **CD7 Median** | **HSPC** | 0 | 1 | 0 | 0 | 0 | 0 | 0 | 1 | 0 | 0 | 0 | 0 | 0 | 0 | 0 |
| | **Blasts** | 0 | 0 | 0 | 0 | 0 | 0 | 0 | 0 | 0 | 0 | 0 | 0 | 0 | 0 | 0 |
| | **Mono** | 0 | 0 | 0 | 0 | 0 | 0 | 0 | 0 | 0 | 0 | 0 | 0 | 0 | 0 | 0 |
| | **Gran** | 0 | 0 | 0 | 0 | 0 | 0 | 0 | 0 | 0 | 0 | 0 | 0 | 0 | 0 | 0 |
| | **Other** | 0 | 0 | 3 | 1 | 2 | 2 | 0 | 0 | 0 | 2 | 0 | 3 | 1 | 0 | 1 |
| **CD71 Median** | **HSPC** | 0 | 0 | 0 | 0 | 0 | 1 | 0 | 0 | 0 | 0 | 0 | 0 | 0 | 0 | 0 |
| | **Blasts** | 0 | 0 | 0 | 0 | 0 | 1 | 0 | 0 | 0 | 0 | 0 | 0 | 0 | 0 | 0 |
| | **Mono** | 0 | 0 | 0 | 0 | 0 | 0 | 0 | 0 | 0 | 0 | 0 | 0 | 0 | 0 | 0 |
| | **Gran** | 0 | 0 | 1 | 0 | 0 | 0 | 0 | 0 | 0 | 0 | 0 | 0 | 0 | 0 | 0 |
| | **Other** | 0 | 2 | 3 | 1 | 3 | 3 | 1 | 1 | 1 | 0 | 0 | 2 | 0 | 0 | 1 |
| **CD235 Median** | **HSPC** | 0 | 1 | 0 | 0 | 1 | 0 | 0 | 2 | 0 | 0 | 0 | 2 | 0 | 0 | 0 |
| | **Blasts** | 0 | 0 | 0 | 0 | 0 | 0 | 0 | 0 | 0 | 0 | 0 | 0 | 0 | 0 | 0 |
| | **Mono** | 0 | 0 | 0 | 0 | 0 | 0 | 0 | 0 | 0 | 0 | 0 | 0 | 0 | 0 | 0 |
| | **Gran** | 0 | 0 | 0 | 0 | 0 | 0 | 0 | 0 | 0 | 0 | 0 | 0 | 0 | 0 | 0 |
| | **Other** | 1 | 0 | 3 | 0 | 1 | 0 | 1 | 0 | 1 | 0 | 0 | 2 | 0 | 0 | 1 |
| **CD47 Median** | **HSPC** | 0 | 2 | 4 | 0 | 0 | 0 | 0 | 1 | 0 | 0 | 0 | 2 | 0 | 0 | 0 |
| | **Blasts** | 0 | 0 | 1 | 0 | 0 | 0 | 0 | 0 | 0 | 0 | 0 | 0 | 0 | 1 | 0 |
| | **Mono** | 0 | 0 | 1 | 0 | 0 | 3 | 0 | 0 | 0 | 0 | 0 | 0 | 0 | 1 | 0 |
| | **Gran** | 1 | 0 | 1 | 0 | 0 | 0 | 0 | 0 | 0 | 0 | 2 | 0 | 0 | 0 | 0 |
| | **Other** | 0 | 0 | 4 | 0 | 1 | 4 | 0 | 0 | 1 | 0 | 0 | 0 | 2 | 1 | 1 |
| **CD8 Median** | **HSPC** | 0 | 1 | 1 | 0 | 1 | 0 | 0 | 0 | 0 | 0 | 0 | 1 | 0 | 0 | 0 |
| | **Blasts** | 0 | 0 | 0 | 0 | 0 | 0 | 0 | 0 | 0 | 0 | 0 | 0 | 0 | 0 | 0 |
| | **Mono** | 0 | 0 | 0 | 0 | 0 | 0 | 0 | 0 | 0 | 0 | 0 | 0 | 0 | 0 | 0 |
| | **Gran** | 0 | 0 | 0 | 0 | 0 | 0 | 0 | 0 | 0 | 0 | 0 | 0 | 0 | 0 | 0 |
| | **Other** | 0 | 2 | 0 | 1 | 1 | 2 | 0 | 0 | 0 | 0 | 0 | 0 | 0 | 0 | 0 |
| **CD34 Median** | **HSPC** | 1 | 3 | 2 | 1 | 0 | 1 | 1 | 1 | 0 | 2 | 1 | 3 | 0 | 0 | 0 |
| | **Blasts** | 0 | 0 | 0 | 0 | 0 | 1 | 0 | 0 | 0 | 0 | 0 | 0 | 0 | 0 | 0 |
| | **Mono** | 0 | 0 | 0 | 0 | 0 | 0 | 0 | 0 | 0 | 0 | 0 | 0 | 0 | 0 | 0 |
| | **Gran** | 0 | 0 | 0 | 0 | 0 | 0 | 0 | 0 | 0 | 0 | 0 | 0 | 0 | 0 | 0 |
| | **Other** | 0 | 0 | 1 | 0 | 0 | 0 | 0 | 0 | 0 | 0 | 0 | 0 | 0 | 0 | 0 |

**Supplemental Table 5**: Fequency of surface marker aberrancies by sample. Number of marker abberrancies is shown for each marker and population (HSPC, Blasts, Monocytes, Granulocytes, and Other), in each sample.
Samples MDS3, 13 & 21 are serial samples from the same patient each several months apart.

| | | Lower Risk MDS | | | | | | | | | | | | ICUS | | |
|---|---|---|---|---|---|---|---|---|---|---|---|---|---|---|---|---|
| | | MDS23 | MDS25 | MDS5 | MDS6 | MDS8 | MDS9 | MDS12 | MDS14 | MDS19 | MDS20 | MDS28 | MDS11 | ICUS18 | ICUS22 | ICUS24 |
| **CD117** | **HSPC** | 1 | 0 | 0 | 0 | 2 | 0 | 4 | 0 | 0 | 0 | 3 | 1 | 0 | 0 | 0 |
| **Median** | **Blasts** | 0 | 0 | 0 | 0 | 0 | 0 | 0 | 0 | 0 | 0 | 0 | 0 | 0 | 0 | 0 |
| | **Mono** | 0 | 0 | 0 | 0 | 0 | 0 | 0 | 0 | 0 | 0 | 0 | 0 | 0 | 0 | 0 |
| | **Gran** | 0 | 0 | 0 | 0 | 0 | 0 | 0 | 0 | 0 | 0 | 0 | 0 | 0 | 0 | 0 |
| | **Other** | 0 | 0 | 0 | 0 | 0 | 0 | 0 | 0 | 0 | 0 | 1 | 0 | 0 | 0 | 0 |
| **CD56** | **HSPC** | 0 | 0 | 0 | 0 | 0 | 0 | 0 | 0 | 0 | 0 | 0 | 0 | 0 | 0 | 0 |
| **Median** | **Blasts** | 0 | 0 | 0 | 0 | 0 | 0 | 0 | 0 | 0 | 0 | 0 | 0 | 0 | 0 | 0 |
| | **Mono** | 0 | 0 | 0 | 1 | 0 | 0 | 3 | 0 | 0 | 0 | 0 | 0 | 0 | 0 | 0 |
| | **Gran** | 0 | 0 | 0 | 2 | 0 | 0 | 0 | 0 | 0 | 0 | 0 | 0 | 0 | 0 | 0 |
| | **Other** | 0 | 0 | 1 | 0 | 0 | 0 | 1 | 1 | 1 | 1 | 0 | 0 | 0 | 0 | 0 |
| **CD90** | **HSPC** | 0 | 1 | 0 | 2 | 0 | 0 | 0 | 1 | 0 | 0 | 0 | 0 | 0 | 0 | 0 |
| **Median** | **Blasts** | 0 | 0 | 0 | 0 | 0 | 0 | 0 | 0 | 0 | 0 | 0 | 0 | 0 | 0 | 0 |
| | **Mono** | 0 | 0 | 0 | 0 | 0 | 0 | 0 | 0 | 0 | 0 | 0 | 0 | 0 | 0 | 0 |
| | **Gran** | 0 | 0 | 0 | 0 | 0 | 0 | 0 | 0 | 0 | 0 | 0 | 0 | 0 | 0 | 0 |
| | **Other** | 0 | 0 | 0 | 0 | 0 | 0 | 0 | 0 | 0 | 0 | 0 | 0 | 0 | 0 | 0 |
| **CD33** | **HSPC** | 0 | 0 | 0 | 2 | 0 | 0 | 0 | 0 | 0 | 0 | 0 | 0 | 0 | 0 | 0 |
| **Median** | **Blasts** | 1 | 0 | 0 | 0 | 1 | 1 | 0 | 0 | 0 | 0 | 0 | 1 | 0 | 0 | 0 |
| | **Mono** | 0 | 0 | 0 | 0 | 0 | 0 | 0 | 0 | 0 | 0 | 0 | 0 | 0 | 0 | 0 |
| | **Gran** | 0 | 0 | 0 | 0 | 0 | 0 | 0 | 0 | 0 | 0 | 0 | 0 | 0 | 0 | 0 |
| | **Other** | 0 | 0 | 1 | 2 | 0 | 0 | 1 | 0 | 0 | 0 | 3 | 0 | 0 | 0 | 0 |
| **CD64** | **HSPC** | 0 | 1 | 2 | 0 | 1 | 0 | 0 | 0 | 0 | 0 | 0 | 1 | 0 | 0 | 0 |
| **Medians** | **Blasts** | 0 | 0 | 0 | 0 | 0 | 0 | 1 | 0 | 0 | 0 | 1 | 0 | 0 | 0 | 0 |
| | **Mono** | 0 | 0 | 3 | 0 | 0 | 0 | 0 | 0 | 0 | 0 | 0 | 0 | 0 | 0 | 0 |
| | **Gran** | 1 | 0 | 4 | 0 | 0 | 0 | 0 | 0 | 0 | 0 | 0 | 0 | 0 | 0 | 0 |
| | **Other** | 0 | 0 | 2 | 0 | 0 | 0 | 2 | 0 | 0 | 0 | 3 | 0 | 0 | 0 | 0 |
| **CD16** | **HSPC** | 0 | 3 | 2 | 0 | 1 | 0 | 0 | 2 | 0 | 1 | 0 | 0 | 0 | 0 | 0 |
| **Median** | **Blasts** | 0 | 0 | 0 | 0 | 0 | 0 | 0 | 0 | 0 | 0 | 0 | 0 | 0 | 0 | 0 |
| | **Mono** | 0 | 0 | 0 | 0 | 0 | 1 | 0 | 0 | 0 | 1 | 0 | 0 | 0 | 0 | 0 |
| | **Gran** | 0 | 0 | 1 | 0 | 0 | 0 | 0 | 1 | 0 | 1 | 0 | 0 | 0 | 0 | 0 |
| | **Other** | 0 | 1 | 1 | 0 | 1 | 2 | 0 | 1 | 0 | 1 | 0 | 0 | 2 | 0 | 0 |
| **CD11b** | **HSPC** | 0 | 3 | 2 | 0 | 1 | 0 | 0 | 2 | 0 | 1 | 0 | 2 | 0 | 0 | 0 |
| **Median** | **Blasts** | 0 | 0 | 0 | 0 | 0 | 0 | 0 | 0 | 0 | 0 | 0 | 0 | 0 | 0 | 0 |
| | **Mono** | 0 | 0 | 1 | 0 | 0 | 0 | 0 | 0 | 0 | 0 | 2 | 0 | 0 | 1 | 0 |
| | **Gran** | 0 | 0 | 0 | 0 | 0 | 0 | 2 | 0 | 0 | 0 | 2 | 0 | 0 | 1 | 0 |
| | **Other** | 2 | 1 | 3 | 1 | 1 | 1 | 1 | 0 | 2 | 0 | 3 | 1 | 1 | 1 | 0 |

**Supplemental Table 5**: Fequency of surface marker aberrancies by sample. Number of marker abberrancies is shown for each marker and population (HSPC, Blasts, Monocytes, Granulocytes, and Other), in each sample.
Samples MDS3, 13 & 21 are serial samples from the same patient each several months apart.

| | | Lower Risk MDS | | | | | | | | | | | | ICUS | | |
|---|---|---|---|---|---|---|---|---|---|---|---|---|---|---|---|---|
| | | MDS23 | MDS25 | MDS5 | MDS6 | MDS8 | MDS9 | MDS12 | MDS14 | MDS19 | MDS20 | MDS28 | MDS11 | ICUS18 | ICUS22 | ICUS24 |
| **CD15 Median** | **HSPC** | 0 | 3 | 1 | 0 | 1 | 0 | 0 | 1 | 0 | 1 | 0 | 0 | 0 | 0 | 0 |
| | **Blasts** | 0 | 0 | 0 | 0 | 0 | 0 | 0 | 0 | 0 | 0 | 0 | 0 | 0 | 0 | 0 |
| | **Mono** | 0 | 0 | 0 | 0 | 0 | 0 | 0 | 0 | 0 | 0 | 0 | 0 | 0 | 0 | 0 |
| | **Gran** | 0 | 0 | 1 | 1 | 1 | 1 | 0 | 2 | 4 | 1 | 1 | 1 | 1 | 4 | 1 |
| | **Other** | 1 | 0 | 2 | 0 | 1 | 1 | 1 | 0 | 1 | 0 | 0 | 2 | 1 | 1 | 1 |
| **CD123 Median** | **HSPC** | 1 | 0 | 0 | 0 | 0 | 0 | 3 | 0 | 0 | 0 | 0 | 0 | 0 | 0 | 0 |
| | **Blasts** | 0 | 0 | 0 | 0 | 0 | 0 | 0 | 0 | 0 | 0 | 0 | 0 | 0 | 0 | 0 |
| | **Mono** | 0 | 0 | 0 | 0 | 0 | 0 | 0 | 0 | 0 | 0 | 0 | 0 | 0 | 0 | 0 |
| | **Gran** | 0 | 0 | 0 | 0 | 0 | 0 | 0 | 0 | 0 | 0 | 0 | 0 | 0 | 0 | 0 |
| | **Other** | 0 | 1 | 0 | 0 | 0 | 1 | 0 | 0 | 0 | 0 | 1 | 0 | 1 | 1 | 0 |
| **CD3 Median** | **HSPC** | 0 | 0 | 0 | 0 | 0 | 0 | 0 | 0 | 0 | 0 | 0 | 0 | 0 | 0 | 0 |
| | **Blasts** | 0 | 0 | 0 | 0 | 0 | 0 | 0 | 0 | 0 | 0 | 0 | 0 | 0 | 0 | 0 |
| | **Mono** | 0 | 0 | 0 | 0 | 0 | 0 | 0 | 0 | 0 | 0 | 0 | 0 | 0 | 0 | 0 |
| | **Gran** | 0 | 0 | 0 | 0 | 0 | 0 | 0 | 0 | 0 | 0 | 0 | 0 | 0 | 0 | 0 |
| | **Other** | 0 | 0 | 0 | 0 | 1 | 1 | 0 | 0 | 0 | 1 | 0 | 1 | 0 | 0 | 0 |
| **CD45 Median** | **HSPC** | 0 | 4 | 3 | 0 | 1 | 0 | 0 | 3 | 0 | 0 | 0 | 2 | 0 | 0 | 0 |
| | **Blasts** | 0 | 1 | 0 | 0 | 0 | 0 | 0 | 1 | 0 | 1 | 0 | 1 | 1 | 1 | 0 |
| | **Mono** | 0 | 0 | 0 | 0 | 2 | 2 | 0 | 1 | 0 | 0 | 0 | 0 | 0 | 2 | 0 |
| | **Gran** | 0 | 1 | 4 | 0 | 0 | 0 | 0 | 0 | 0 | 0 | 0 | 0 | 0 | 0 | 0 |
| | **Other** | 2 | 1 | 7 | 3 | 1 | 2 | 0 | 0 | 1 | 3 | 3 | 1 | 1 | 1 | 0 |
| **CD133 Median** | **HSPC** | 0 | 0 | 0 | 0 | 1 | 0 | 0 | 0 | 0 | 0 | 0 | 0 | 0 | 0 | 0 |
| | **Blasts** | 0 | 0 | 0 | 0 | 0 | 0 | 0 | 0 | 0 | 0 | 0 | 0 | 0 | 0 | 0 |
| | **Mono** | 0 | 0 | 0 | 0 | 0 | 0 | 0 | 0 | 0 | 0 | 0 | 0 | 0 | 0 | 0 |
| | **Gran** | 0 | 0 | 0 | 0 | 0 | 0 | 0 | 0 | 0 | 0 | 0 | 0 | 0 | 0 | 0 |
| | **Other** | 0 | 0 | 0 | 0 | 0 | 0 | 0 | 0 | 0 | 0 | 0 | 0 | 0 | 0 | 0 |
| **HLADR Median** | **HSPC** | 2 | 1 | 1 | 3 | 0 | 1 | 4 | 0 | 0 | 0 | 1 | 0 | 0 | 0 | 0 |
| | **Blasts** | 0 | 0 | 0 | 0 | 0 | 0 | 0 | 0 | 0 | 0 | 1 | 0 | 0 | 0 | 0 |
| | **Mono** | 3 | 0 | 0 | 2 | 2 | 2 | 0 | 1 | 2 | 3 | 3 | 0 | 2 | 3 | 0 |
| | **Gran** | 0 | 0 | 0 | 0 | 0 | 0 | 0 | 0 | 0 | 0 | 0 | 0 | 0 | 0 | 0 |
| | **Other** | 4 | 3 | 2 | 2 | 0 | 0 | 1 | 0 | 0 | 2 | 2 | 1 | 0 | 1 | 0 |
| **CD44 Median** | **HSPC** | 1 | 1 | 3 | 0 | 0 | 0 | 0 | 3 | 0 | 1 | 0 | 1 | 0 | 0 | 0 |
| | **Blasts** | 1 | 1 | 0 | 1 | 0 | 0 | 0 | 1 | 0 | 1 | 1 | 0 | 1 | 0 | 0 |
| | **Mono** | 0 | 0 | 0 | 0 | 0 | 0 | 0 | 1 | 0 | 1 | 0 | 0 | 0 | 0 | 0 |
| | **Gran** | 0 | 0 | 3 | 1 | 0 | 0 | 0 | 0 | 0 | 0 | 0 | 0 | 0 | 0 | 0 |
| | **Other** | 3 | 1 | 4 | 2 | 0 | 1 | 3 | 0 | 1 | 1 | 2 | 1 | 1 | 0 | 0 |

**Supplemental Table 5**: Fequency of surface marker aberrancies by sample. Number of marker abberrancies is shown for each marker and population (HSPC, Blasts, Monocytes, Granulocytes, and Other), in each sample.
Samples MDS3, 13 & 21 are serial samples from the same patient each several months apart.

| | | Lower Risk MDS | | | | | | | | | | | | ICUS | | |
|---|---|---|---|---|---|---|---|---|---|---|---|---|---|---|---|---|
| | | **MDS23** | **MDS25** | **MDS5** | **MDS6** | **MDS8** | **MDS9** | **MDS12** | **MDS14** | **MDS19** | **MDS20** | **MDS28** | **MDS11** | **ICUS18** | **ICUS22** | **ICUS24** |
| **CD38 Median** | **HSPC** | 0 | 0 | 0 | 0 | 0 | 0 | 0 | 0 | 0 | 0 | 0 | 1 | 0 | 0 | 0 |
| | **Blasts** | 0 | 0 | 0 | 0 | 0 | 0 | 0 | 0 | 0 | 0 | 0 | 0 | 1 | 0 | 0 |
| | **Mono** | 0 | 0 | 2 | 0 | 0 | 2 | 1 | 0 | 0 | 1 | 1 | 1 | 1 | 2 | 0 |
| | **Gran** | 0 | 0 | 0 | 0 | 0 | 0 | 0 | 0 | 0 | 0 | 0 | 0 | 0 | 0 | 0 |
| | **Other** | 0 | 1 | 2 | 0 | 1 | 0 | 0 | 2 | 1 | 0 | 3 | 1 | 0 | 3 | 0 |
| **CD14 Median** | **HSPC** | 0 | 0 | 0 | 0 | 0 | 0 | 0 | 0 | 0 | 0 | 0 | 1 | 0 | 0 | 0 |
| | **Blasts** | 0 | 0 | 0 | 0 | 0 | 0 | 0 | 0 | 0 | 0 | 0 | 0 | 0 | 0 | 0 |
| | **Mono** | 0 | 0 | 0 | 0 | 0 | 0 | 0 | 0 | 0 | 0 | 0 | 0 | 0 | 0 | 0 |
| | **Gran** | 0 | 0 | 0 | 0 | 0 | 0 | 0 | 0 | 0 | 0 | 0 | 0 | 0 | 0 | 0 |
| | **Other** | 0 | 1 | 0 | 0 | 0 | 0 | 0 | 0 | 0 | 0 | 0 | 0 | 0 | 0 | 0 |
| **CD321 Median** | **HSPC** | 1 | 0 | 0 | 0 | 0 | 0 | 2 | 0 | 0 | 0 | 0 | 0 | 0 | 0 | 0 |
| | **Blasts** | 0 | 0 | 0 | 1 | 0 | 1 | 0 | 0 | 0 | 0 | 0 | 0 | 0 | 0 | 0 |
| | **Mono** | 0 | 0 | 0 | 3 | 1 | 3 | 0 | 0 | 3 | 2 | 1 | 0 | 0 | 1 | 0 |
| | **Gran** | 0 | 0 | 1 | 0 | 0 | 0 | 0 | 1 | 0 | 0 | 0 | 0 | 0 | 0 | 0 |
| | **Other** | 1 | 0 | 1 | 1 | 0 | 2 | 0 | 1 | 1 | 2 | 2 | 1 | 1 | 0 | 1 |
| **CD99 Median** | **HSPC** | 0 | 0 | 0 | 0 | 0 | 0 | 0 | 0 | 0 | 0 | 0 | 0 | 0 | 0 | 0 |
| | **Blasts** | 0 | 0 | 0 | 0 | 0 | 0 | 0 | 0 | 0 | 0 | 0 | 0 | 0 | 0 | 0 |
| | **Mono** | 0 | 0 | 0 | 0 | 0 | 0 | 0 | 0 | 0 | 0 | 0 | 0 | 0 | 0 | 0 |
| | **Gran** | 0 | 0 | 0 | 0 | 0 | 0 | 0 | 0 | 0 | 0 | 0 | 0 | 0 | 0 | 0 |
| | **Other** | 0 | 1 | 3 | 3 | 4 | 2 | 0 | 0 | 1 | 3 | 0 | 3 | 2 | 0 | 1 |
| **CD13 Median** | **HSPC** | 0 | 0 | 0 | 0 | 0 | 0 | 0 | 0 | 0 | 0 | 0 | 0 | 0 | 0 | 0 |
| | **Blasts** | 0 | 0 | 0 | 0 | 0 | 0 | 0 | 0 | 0 | 0 | 0 | 0 | 0 | 0 | 0 |
| | **Mono** | 0 | 0 | 0 | 0 | 0 | 0 | 0 | 1 | 0 | 0 | 0 | 0 | 0 | 0 | 0 |
| | **Gran** | 0 | 0 | 0 | 0 | 0 | 0 | 0 | 0 | 0 | 0 | 0 | 0 | 0 | 0 | 0 |
| | **Other** | 0 | 1 | 0 | 0 | 0 | 0 | 0 | 0 | 0 | 0 | 1 | 0 | 0 | 1 | 0 |
| **CXCR4 Median** | **HSPC** | 0 | 1 | 0 | 0 | 0 | 0 | 0 | 0 | 0 | 0 | 0 | 0 | 0 | 0 | 0 |
| | **Blasts** | 0 | 0 | 0 | 0 | 0 | 0 | 0 | 0 | 0 | 0 | 0 | 0 | 0 | 0 | 0 |
| | **Mono** | 0 | 0 | 0 | 0 | 0 | 0 | 0 | 0 | 0 | 0 | 0 | 0 | 0 | 0 | 0 |
| | **Gran** | 0 | 0 | 0 | 0 | 0 | 0 | 0 | 0 | 0 | 0 | 0 | 0 | 0 | 0 | 0 |
| | **Other** | 0 | 0 | 2 | 0 | 1 | 1 | 1 | 0 | 0 | 0 | 0 | 0 | 0 | 0 | 1 |
| **CD10 Median** | **HSPC** | 0 | 1 | 0 | 0 | 1 | 0 | 0 | 1 | 0 | 0 | 0 | 0 | 0 | 0 | 0 |
| | **Blasts** | 0 | 0 | 0 | 0 | 0 | 0 | 0 | 0 | 0 | 0 | 0 | 0 | 0 | 0 | 0 |
| | **Mono** | 0 | 0 | 0 | 0 | 0 | 0 | 0 | 0 | 0 | 0 | 0 | 0 | 0 | 0 | 0 |
| | **Gran** | 0 | 1 | 0 | 0 | 0 | 0 | 0 | 0 | 0 | 0 | 0 | 0 | 0 | 0 | 0 |
| | **Other** | 0 | 0 | 0 | 0 | 0 | 0 | 0 | 0 | 0 | 0 | 0 | 0 | 0 | 0 | 0 |

**Supplemental Table 5**: Fequency of surface marker aberrancies by sample. Number of marker abberrancies is shown for each marker and population (HSPC, Blasts, Monocytes, Granulocytes, and Other), in each sample.
Samples MDS3, 13 & 21 are serial samples from the same patient each several months apart.

| | | Lower Risk MDS | | | | | | | | | | | | ICUS | | |
|---|---|---|---|---|---|---|---|---|---|---|---|---|---|---|---|---|
| | | MDS23 | MDS25 | MDS5 | MDS6 | MDS8 | MDS9 | MDS12 | MDS14 | MDS19 | MDS20 | MDS28 | MDS11 | ICUS18 | ICUS22 | ICUS24 |
| CD19 Median | HSPC | 0 | 0 | 0 | 0 | 0 | 0 | 0 | 0 | 0 | 0 | 0 | 0 | 0 | 0 | 0 |
| | Blasts | 0 | 0 | 0 | 0 | 0 | 0 | 0 | 0 | 0 | 0 | 0 | 0 | 0 | 0 | 0 |
| | Mono | 0 | 0 | 0 | 0 | 0 | 0 | 0 | 0 | 0 | 0 | 0 | 0 | 0 | 0 | 0 |
| | Gran | 0 | 0 | 0 | 0 | 0 | 0 | 0 | 0 | 0 | 0 | 0 | 0 | 0 | 0 | 0 |
| | Other | 0 | 0 | 0 | 0 | 0 | 0 | 0 | 0 | 1 | 1 | 0 | 0 | 0 | 0 | 0 |
| CD20 Median | HSPC | 0 | 0 | 0 | 0 | 0 | 0 | 0 | 0 | 0 | 0 | 0 | 0 | 0 | 0 | 0 |
| | Blasts | 0 | 0 | 0 | 0 | 0 | 0 | 0 | 0 | 0 | 0 | 0 | 0 | 0 | 0 | 0 |
| | Mono | 0 | 0 | 0 | 0 | 0 | 0 | 0 | 0 | 0 | 0 | 0 | 0 | 0 | 0 | 0 |
| | Gran | 0 | 0 | 0 | 0 | 0 | 0 | 0 | 0 | 0 | 0 | 0 | 0 | 0 | 0 | 0 |
| | Other | 0 | 0 | 0 | 0 | 0 | 0 | 1 | 1 | 1 | 1 | 0 | 0 | 0 | 0 | 0 |
| CD69 Median | HSPC | 0 | 0 | 0 | 0 | 0 | 0 | 0 | 0 | 0 | 0 | 0 | 0 | 0 | 0 | 0 |
| | Blasts | 0 | 0 | 0 | 0 | 0 | 0 | 0 | 0 | 0 | 0 | 0 | 0 | 0 | 0 | 0 |
| | Mono | 0 | 0 | 0 | 0 | 0 | 0 | 0 | 0 | 0 | 0 | 0 | 0 | 0 | 0 | 0 |
| | Gran | 0 | 0 | 0 | 0 | 0 | 0 | 0 | 0 | 0 | 0 | 0 | 0 | 0 | 0 | 0 |
| | Other | 0 | 1 | 3 | 0 | 0 | 1 | 3 | 0 | 0 | 0 | 1 | 0 | 0 | 0 | 1 |

**Supplemental Table 6:** Frequency of MDS sample surface marker aberrances in gated cell populations and by using an appoximation of the LeukemiaNet criteria.
"Yes" indicates the presence of observed aberrancy or aberrancies in the indicated population

| | | Gated aberrancy analysis | | Modified European LeukemiaNet MDS criteria | | | | |
|---|---|---|---|---|---|---|---|---|
| | | At least 1 HSPC abberancy | At least 2 HSPC abberancies | Immature B cell Frequency <normal | Abnormal Lymphocyte : Myeloblast CD45 ratio | Abnormal Myeloblast Frequency | Immuno-phenotypically abnormal Granulocytes | 2 or more abnormalities |
| MDS17 | AML / RAEB-T | Yes | Yes | Yes | Yes | Yes | Yes | Yes |
| MDS15 | | Yes | Yes | Yes | Yes | Yes | Yes | Yes |
| MDS4 | | Yes | Yes | No | Yes | Yes | Yes | Yes |
| MDS13 | Higher Risk MDS | Yes | Yes | No | Yes | Yes | Yes | Yes |
| MDS16 | | Yes | Yes | Yes | No | Yes | No | Yes |
| MDS1 | | Yes | Yes | Yes | Yes | Yes | No | Yes |
| MDS21 | | Yes | Yes | No | Yes | Yes | Yes | Yes |
| MDS26 | | Yes | Yes | Yes | Yes | No | Yes | Yes |
| MDS27 | | Yes | Yes | Yes | Yes | Yes | No | Yes |
| MDS2 | | Yes | Yes | Yes | Yes | Yes | Yes | Yes |
| MDS3 | | Yes | Yes | No | Yes | Yes | Yes | Yes |
| MDS23 | Lower Risk MDS | Yes | Yes | Yes | Yes | Yes | Yes | Yes |
| MDS25 | | Yes | Yes | Yes | No | No | Yes | Yes |
| MDS5 | | Yes | Yes | Yes | No | No | Yes | Yes |
| MDS6 | | Yes | Yes | Yes | Yes | No | No | Yes |
| MDS8 | | Yes | Yes | Yes | No | No | No | No |
| MDS9 | | Yes | Yes | Yes | Yes | Yes | No | Yes |
| MDS12 | | Yes | Yes | Yes | No | Yes | Yes | Yes |
| MDS14 | | Yes | Yes | No | No | No | No | No |
| MDS19 | | Yes | No | Yes | No | No | Yes | Yes |
| MDS20 | | Yes | Yes | Yes | No | No | No | No |
| MDS28 | | Yes | Yes | Yes | Yes | No | Yes | Yes |
| MDS11 | | Yes | Yes | Yes | No | No | No | No |
| ICUS18 | ICUS | Yes | No | Yes | No | No | No | No |
| ICUS22 | | No | No | No | No | No | Yes | No |
| ICUS24 | | No | No | No | No | No | No | No |
| Nl-1 | Healthy Controls | No | No | No | No | No | No | No |
| Nl-3 | | No | No | No | No | No | No | No |
| Nl-4 | | No | No | No | No | No | No | No |
| Nl-6 | | No | No | No | No | No | No | No |
| Nl-3 | | No | No | No | No | No | No | No |
| Nl-4 | | No | No | No | No | No | No | No |
| Nl-5 | | No | No | No | No | No | No | No |
| Nl-6 | | No | No | No | No | No | No | No |

**Supplemental Table 7**: Immunophenotype of Aberrant Myeloid Cells.
The normalized expression level of each marker on the gated IAMC population (See Supplemental Figure 1 for gating) is shown.
The expression level of each marker is expressed as a percentage of the average median expression level of that marker in the normal cell population with the highest expression level of the marker.
Percent values of <0 were set to 0 to facilitate heat plot of data.
Note that samples MDS3, MDS13, and MDS21 come from serial biopsies of the same patient (each several months apart) and demonstrate consistent properties.

| Sample | Group | CD19 | CD3 | CD69 | CD20 | CD133 | CD117 | CD7 | CD123 | CD10 | CD90 | CD34 | CD16 | CXCR4 | CD14 | CD15 | CD71 | CD56 | CD13 | CD45RA | CD235 | HLA-DR | CD8 | CD99 | CD33 | CD321 | CD38 | CD47 | CD45 | CD64 | CD11b | CD44 |
|---|---|---|---|---|---|---|---|---|---|---|---|---|---|---|---|---|---|---|---|---|---|---|---|---|---|---|---|---|---|---|---|---|
| MDS17 | AML/RAE B-T | 0.0 | 0.0 | 0.0 | 0.0 | 0.0 | 0.0 | 0.0 | 0.0 | 20.0 | 0.0 | 0.2 | 12.6 | 4.2 | 0.0 | 1.3 | 0.1 | 8.8 | 10.1 | 41.2 | 3.8 | 0.0 | 3.5 | 1.8 | 13.6 | 0.4 | 0.2 | 31.7 | 31.4 | 12.4 | 126.8 | 37.6 |
| MDS15* | | 0.0 | 0.0 | 0.0 | 0.0 | 0.0 | 0.0 | 0.0 | 1.4 | 14.9 | 0.0 | 0.3 | 5.8 | 9.2 | 1.0 | 1.0 | 0.1 | 12.0 | 13.3 | 31.7 | 2.2 | 0.1 | 1.0 | 3.8 | 23.2 | 2.7 | 0.3 | 13.7 | 14.4 | 2.7 | 41.8 | 48.8 |
| MDS4 | | 0.0 | 0.0 | 0.0 | 0.0 | 0.0 | 0.0 | 0.0 | 0.0 | 17.0 | 0.0 | 0.0 | 13.9 | 3.0 | 0.1 | 0.8 | 0.0 | 5.4 | 8.3 | 36.0 | 1.8 | 0.1 | 1.1 | 1.5 | 14.2 | 1.2 | 0.1 | 13.8 | 19.6 | 43.7 | 64.1 | 29.2 |
| MDS13 | Higher Risk MDS | 0.0 | 0.0 | 0.0 | 0.0 | 0.0 | 0.0 | 0.0 | 0.0 | 0.0 | 0.0 | 0.0 | 8.3 | 0.2 | 0.4 | 1.4 | 0.0 | 9.8 | 10.3 | 27.6 | 2.7 | 0.0 | 1.5 | 2.7 | 15.7 | 0.8 | 1.6 | 20.2 | 24.4 | 4.3 | 88.1 | 42.6 |
| MDS16 | | 0.0 | 0.0 | 0.0 | 0.0 | 0.0 | 0.0 | 0.0 | 0.0 | 0.0 | 0.0 | 0.2 | 3.4 | 1.0 | 2.3 | 0.1 | 0.3 | 13.6 | 37.1 | 26.0 | 1.6 | 1.1 | 1.2 | 19.7 | 57.4 | 1.3 | 1.5 | 11.9 | 34.4 | 69.5 | 28.7 | 54.3 |
| MDS1 | | 0.0 | 0.0 | 0.0 | 0.0 | 0.0 | 0.0 | 0.0 | 0.0 | 0.0 | 0.0 | 0.1 | 9.3 | 0.7 | 0.1 | 0.7 | 0.0 | 12.0 | 8.7 | 20.4 | 0.9 | 0.0 | 1.2 | 1.3 | 0.0 | 0.5 | 0.3 | 18.8 | 19.8 | 0.9 | 36.4 | 19.5 |
| MDS21 | | 0.0 | 0.0 | 0.0 | 0.0 | 0.0 | 0.0 | 0.0 | 0.0 | 0.0 | 0.0 | 0.0 | 7.6 | 0.5 | 0.1 | 1.3 | 0.0 | 5.8 | 5.3 | 16.1 | 1.5 | 0.1 | 0.9 | 0.9 | 11.9 | 0.5 | 1.8 | 18.4 | 17.5 | 4.8 | 86.2 | 23.0 |
| MDS26* | | 0.0 | 0.0 | 0.0 | 0.0 | 0.0 | 0.0 | 0.0 | 0.0 | 0.0 | 0.0 | 0.0 | 6.1 | 3.5 | 0.0 | 1.8 | 0.0 | 4.8 | 2.9 | 9.3 | 0.6 | 0.0 | 0.7 | 0.4 | 1.2 | 0.8 | 0.2 | 11.4 | 6.3 | 1.2 | 25.6 | 2.9 |
| MDS27 | | 0.0 | 0.0 | 0.0 | 0.0 | 0.0 | 0.0 | 0.5 | 0.0 | 1.7 | 0.0 | 0.2 | 14.9 | 0.0 | 1.0 | 0.2 | 0.1 | 8.3 | 19.1 | 28.4 | 1.4 | 0.1 | 1.2 | 7.4 | 0.0 | 0.8 | 1.0 | 8.5 | 21.5 | 1.7 | 16.6 | 18.3 |
| MDS2 | | 0.0 | 0.0 | 0.0 | 0.0 | 0.0 | 0.0 | 0.0 | 0.0 | 0.0 | 0.0 | 0.0 | 17.9 | 0.0 | 0.0 | 1.5 | 0.0 | 3.7 | 7.7 | 31.0 | 0.9 | 0.0 | 1.1 | 0.3 | 5.1 | 0.1 | 0.8 | 25.3 | 17.3 | 28.3 | 147.9 | 35.0 |
| MDS3 | | 0.0 | 0.0 | 0.0 | 0.0 | 0.0 | 0.0 | 0.0 | 0.0 | 0.0 | 0.0 | 0.0 | 5.9 | 1.3 | 0.2 | 1.2 | 0.1 | 7.6 | 7.8 | 39.1 | 3.0 | 0.1 | 1.9 | 2.9 | 17.2 | 2.1 | 1.5 | 17.7 | 20.8 | 6.8 | 47.1 | 30.6 |
| MDS23 | Lower Risk MDS | 0.0 | 0.0 | 0.0 | 0.0 | 0.0 | 0.0 | 0.0 | 0.0 | 0.0 | 0.0 | 0.0 | 11.1 | 2.3 | 0.0 | 2.0 | 0.1 | 3.1 | 0.0 | 21.3 | 2.2 | 0.0 | 1.9 | 0.6 | 0.1 | 1.5 | 0.1 | 24.9 | 8.2 | 15.7 | 17.2 | 1.2 |
| MDS25 | | 0.0 | 0.0 | 0.0 | 0.0 | 0.0 | 0.0 | 0.0 | 0.0 | 7.2 | 0.0 | 0.3 | 4.4 | 0.8 | 2.3 | 0.3 | 1.1 | 13.2 | 49.4 | 16.9 | 1.6 | 1.7 | 1.0 | 4.5 | 33.5 | 2.3 | 0.9 | 12.8 | 57.7 | 59.1 | 56.5 | 37.2 |
| MDS5 | | 0.0 | 0.0 | 0.0 | 0.0 | 0.0 | 0.0 | 0.0 | 0.0 | 1.3 | 0.0 | 0.2 | 125.2 | 0.0 | 0.0 | 1.7 | 0.3 | 11.5 | 11.9 | 39.1 | 4.5 | 0.0 | 1.9 | 0.9 | 5.9 | 0.8 | 0.1 | 31.8 | 78.3 | 354.8 | 54.9 | 58.0 |
| MDS6 | | 0.0 | 0.0 | 0.0 | 0.0 | 0.0 | 0.0 | 0.0 | 0.0 | 41.6 | 0.0 | 0.4 | 8.4 | 1.9 | 1.0 | 0.4 | 0.2 | 32.1 | 22.6 | 45.6 | 2.5 | 0.8 | 2.1 | 3.0 | 38.6 | 2.2 | 0.4 | 16.1 | 47.9 | 58.4 | 89.1 | 36.0 |
| MDS8 | | 0.0 | 0.0 | 0.0 | 0.0 | 0.0 | 0.0 | 0.0 | 0.0 | 0.0 | 0.0 | 0.3 | 9.3 | 1.9 | 1.7 | 0.2 | 0.5 | 10.0 | 26.5 | 42.3 | 2.3 | 0.4 | 1.8 | 10.0 | 3.0 | 2.5 | 1.3 | 15.1 | 33.8 | 1.3 | 20.6 | 16.5 |
| MDS9 | | 0.0 | 0.0 | 0.0 | 0.0 | 0.0 | 0.0 | 0.0 | 0.0 | 0.0 | 0.0 | 0.2 | 11.5 | 1.7 | 0.5 | 1.2 | 0.3 | 9.1 | 21.5 | 29.5 | 2.3 | 0.2 | 1.6 | 2.4 | 0.1 | 1.7 | 0.2 | 18.9 | 23.0 | 5.5 | 19.7 | 4.7 |
| MDS12 | | 0.0 | 0.0 | 0.0 | 0.0 | 0.0 | 0.0 | 0.0 | 0.0 | 0.0 | 0.0 | 0.0 | 7.0 | 5.4 | 0.1 | 1.9 | 0.1 | 4.4 | 1.1 | 16.9 | 1.8 | 0.0 | 1.2 | 0.9 | 4.1 | 1.6 | 0.4 | 20.5 | 8.2 | 34.1 | 24.1 | 1.5 |
| MDS14 | | 0.0 | 0.0 | 0.0 | 0.0 | 0.0 | 0.0 | 0.0 | 0.0 | 0.8 | 0.0 | 0.1 | 5.0 | 0.7 | 2.1 | 0.2 | 0.4 | 11.8 | 22.6 | 36.0 | 2.2 | 0.8 | 1.9 | 3.1 | 14.8 | 1.1 | 1.3 | 11.4 | 34.2 | 57.8 | 39.2 | 50.8 |
| MDS19 | | 0.0 | 0.0 | 0.0 | 0.0 | 0.0 | 0.0 | 0.0 | 0.0 | 0.0 | 0.0 | 0.0 | 5.5 | 2.3 | 1.7 | 0.2 | 0.1 | 10.7 | 18.8 | 30.4 | 1.0 | 0.7 | 1.3 | 3.8 | 13.3 | 1.9 | 1.4 | 11.2 | 23.9 | 8.2 | 34.6 | 28.4 |
| MDS20 | | 0.0 | 0.0 | 0.0 | 0.0 | 0.0 | 0.0 | 0.3 | 0.0 | 18.3 | 0.0 | 0.1 | 7.4 | 0.0 | 0.4 | 0.4 | 0.0 | 13.4 | 7.7 | 37.8 | 1.8 | 0.2 | 1.1 | 3.6 | 0.3 | 0.6 | 0.5 | 12.2 | 20.3 | 0.7 | 10.6 | 24.0 |
| MDS28* | | 0.0 | 0.0 | 0.0 | 0.0 | 0.0 | 0.0 | 0.0 | 0.0 | 0.0 | 0.0 | 0.1 | 7.9 | 3.0 | 0.5 | 0.4 | 0.0 | 13.4 | 5.7 | 48.8 | 1.4 | 0.1 | 1.8 | 2.9 | 2.2 | 1.3 | 1.3 | 19.3 | 26.9 | 3.1 | 25.4 | 19.3 |
| MDS11 | | 0.0 | 0.0 | 0.0 | 0.0 | 0.0 | 0.0 | 0.0 | 0.0 | 5.5 | 0.0 | 0.3 | 5.8 | 1.5 | 1.6 | 0.5 | 0.4 | 10.3 | 59.9 | 48.0 | 3.4 | 1.0 | 2.6 | 8.3 | 6.4 | 3.0 | 1.6 | 16.6 | 48.7 | 65.0 | 56.7 | 37.8 |
| ICUS18 | ICUS | 0.0 | 0.0 | 0.0 | 0.0 | 0.0 | 0.0 | 0.0 | 0.0 | 0.0 | 0.0 | 0.2 | 8.8 | 1.6 | 1.4 | 0.2 | 0.4 | 14.6 | 21.2 | 30.8 | 1.6 | 0.3 | 1.9 | 7.4 | 8.3 | 1.5 | 0.9 | 12.9 | 28.8 | 4.0 | 16.9 | 13.8 |
| ICUS22 | | 0.0 | 0.0 | 0.0 | 0.0 | 0.0 | 0.0 | 0.0 | 0.0 | 0.0 | 0.0 | 0.0 | 11.0 | 0.9 | 1.3 | 0.7 | 0.1 | 4.9 | 18.1 | 14.1 | 0.8 | 0.9 | 1.2 | 0.8 | 9.7 | 1.0 | 0.6 | 14.4 | 39.1 | 26.3 | 80.7 | 10.9 |
| ICUS24 | | 0.0 | 0.0 | 0.0 | 0.0 | 0.0 | 0.0 | 0.0 | 0.0 | 0.0 | 0.0 | 0.0 | 5.9 | 2.9 | 1.7 | 0.3 | 0.4 | 6.9 | 34.3 | 18.9 | 1.6 | 1.1 | 1.3 | 1.9 | 14.4 | 1.6 | 0.7 | 13.9 | 38.4 | 76.5 | 40.9 | 17.6 |
| Nl-1 | Normal | 0.0 | 0.0 | 0.0 | 0.0 | 0.0 | 0.0 | 0.0 | 0.3 | 0.0 | 0.0 | 0.1 | 4.3 | 3.9 | 1.4 | 0.2 | 1.3 | 6.8 | 19.4 | 21.3 | 2.1 | 1.1 | 1.3 | 3.3 | 58.1 | 1.9 | 0.8 | 13.3 | 36.2 | 71.5 | 30.4 | 18.3 |
| Nl-3 | | 0.0 | 0.0 | 0.0 | 0.0 | 0.0 | 5.4 | 0.1 | 0.2 | 0.0 | 11.6 | 0.3 | 4.8 | 4.9 | 2.0 | 0.5 | 0.6 | 19.0 | 40.9 | 108.9 | 6.6 | 1.3 | 4.4 | 6.7 | 27.5 | 2.4 | 1.2 | 13.0 | 37.3 | 45.0 | 28.2 | 22.1 |
| Nl-4 | | 0.0 | 0.0 | 0.0 | 0.0 | 0.0 | 1.0 | 0.0 | 0.1 | 0.0 | 11.4 | 0.2 | 4.9 | 4.2 | 1.4 | 0.8 | 0.4 | 17.6 | 23.7 | 92.0 | 6.7 | 0.6 | 4.0 | 6.9 | 21.3 | 2.8 | 0.9 | 13.1 | 31.5 | 46.4 | 29.4 | 20.4 |
| Nl-6 | | 0.0 | 0.0 | 0.0 | 0.0 | 0.0 | 0.0 | 0.0 | 0.0 | 5.5 | 3.9 | 0.2 | 5.4 | 3.6 | 1.1 | 0.8 | 0.5 | 15.8 | 27.8 | 74.6 | 5.7 | 0.6 | 3.2 | 3.3 | 24.3 | 2.4 | 0.7 | 13.9 | 29.1 | 86.9 | 21.2 | 16.1 |
| Nl-3 | | 0.0 | 0.0 | 0.0 | 0.0 | 0.0 | 0.0 | 0.0 | 0.0 | 2.5 | 7.1 | 0.1 | 2.8 | 3.7 | 1.8 | 0.3 | 0.2 | 16.3 | 32.6 | 64.0 | 5.6 | 1.5 | 3.0 | 4.0 | 21.4 | 2.1 | 1.4 | 9.7 | 28.9 | 36.0 | 33.5 | 42.7 |
| Nl-4 | | 0.0 | 0.0 | 0.0 | 0.0 | 0.0 | 0.0 | 0.0 | 0.0 | 3.4 | 4.8 | 0.1 | 3.1 | 3.6 | 1.3 | 0.7 | 0.1 | 16.7 | 12.8 | 53.4 | 6.2 | 0.6 | 2.6 | 4.4 | 16.5 | 2.5 | 1.1 | 9.0 | 19.7 | 33.6 | 25.2 | 36.7 |
| Nl-5 | | 0.0 | 0.0 | 0.0 | 0.0 | 0.0 | 3.8 | 0.1 | 0.0 | 8.9 | 16.4 | 0.2 | 2.1 | 3.5 | 2.2 | 0.2 | 0.3 | 18.9 | 23.9 | 73.8 | 6.9 | 1.5 | 4.0 | 7.9 | 34.3 | 2.3 | 2.0 | 7.2 | 34.5 | 52.0 | 50.0 | 60.6 |
| Nl-6 | | 0.0 | 0.0 | 0.0 | 0.0 | 0.0 | 0.0 | 0.0 | 0.0 | 0.0 | 0.0 | 0.1 | 3.8 | 2.4 | 1.1 | 0.7 | 0.2 | 13.3 | 23.4 | 43.4 | 4.7 | 0.5 | 1.8 | 1.9 | 17.8 | 2.2 | 1.0 | 11.2 | 18.8 | 78.8 | 22.8 | 23.5 |

***CMML**
