## Supplemental Figures 2-13 for "Profiling Myelodysplastic Syndromes by Mass Cytometry Demonstrates Abnormal Progenitor Cell Phenotype and Differentiation"

### Supplemental Figure 2

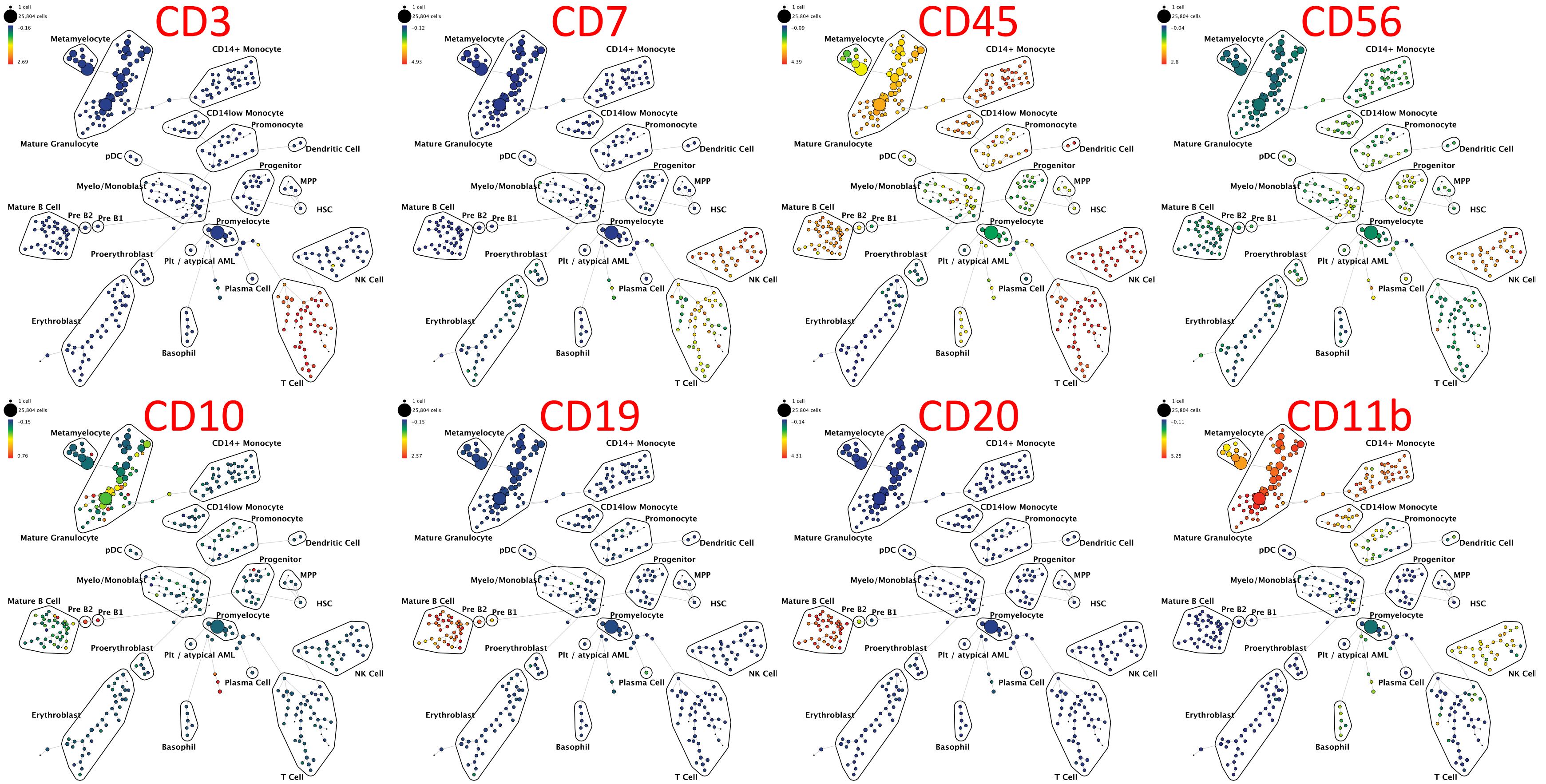

**Supplemental Figure 2:** SPADE plots of normal bone marrow sample #6 (additional markers from analysis shown in Figure 1A). SPADE clustering was performed on all samples (normal and MDS) simultaneously to generate a single tree structure for all samples. All of the cell events from each sample were then mapped to the common tree structure. Each cell cluster (node) of the SPADE tree is colored for the median expression of the indicated markers from low (blue) to high (red). The size of each cell node is correlated to the fraction of cells mapping to the node; however, a minimum size was enforced for most nodes to allow visualization of node color. Immunophenotypic grouping of nodes was performed manually on the basis of the median marker expression level of each node, and based on analysis of the relevant biaxial plots (e.g., CD38 vs. CD34).

### Supplemental Figure 3A

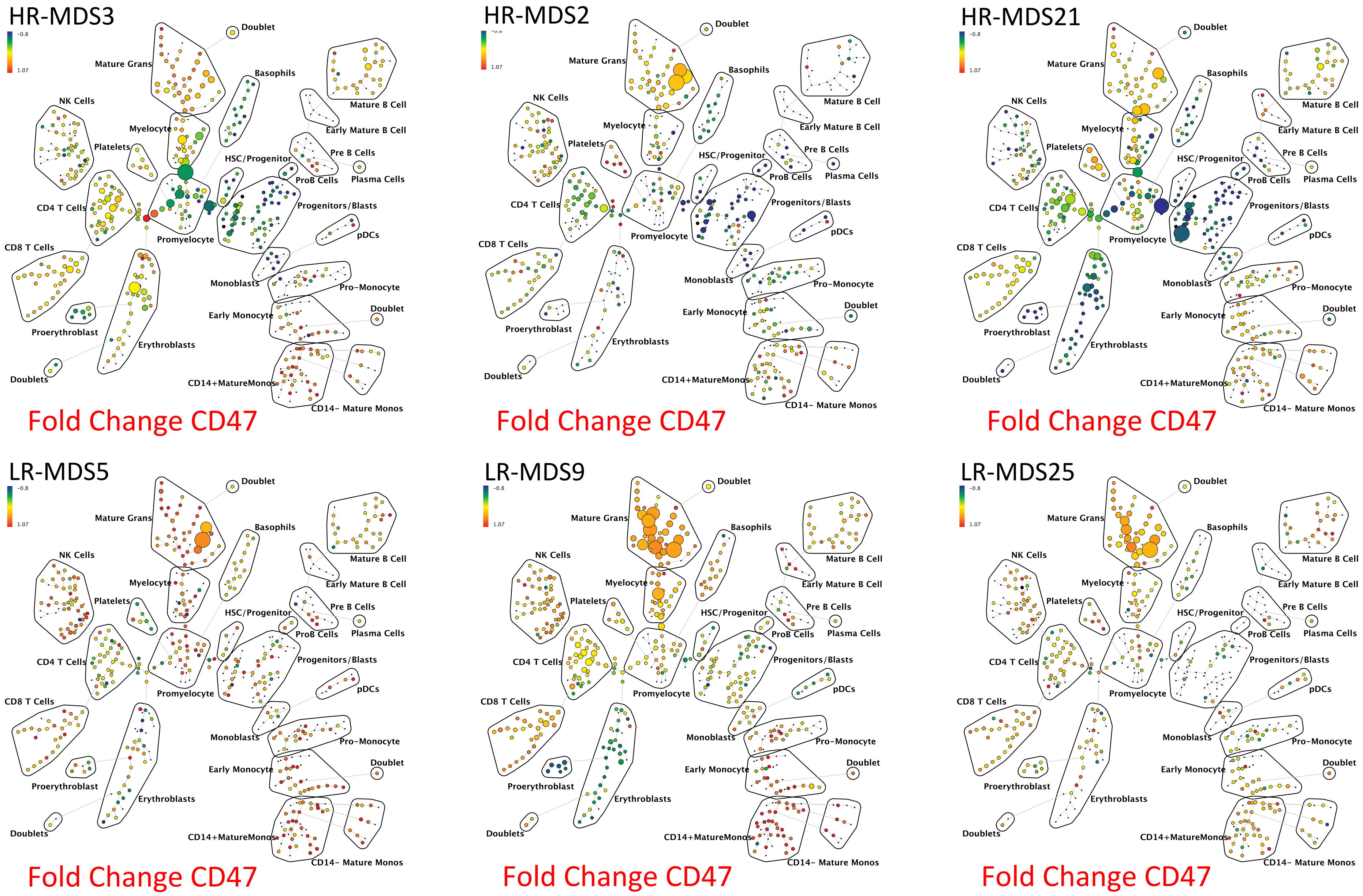

### Supplemental Figure 3B

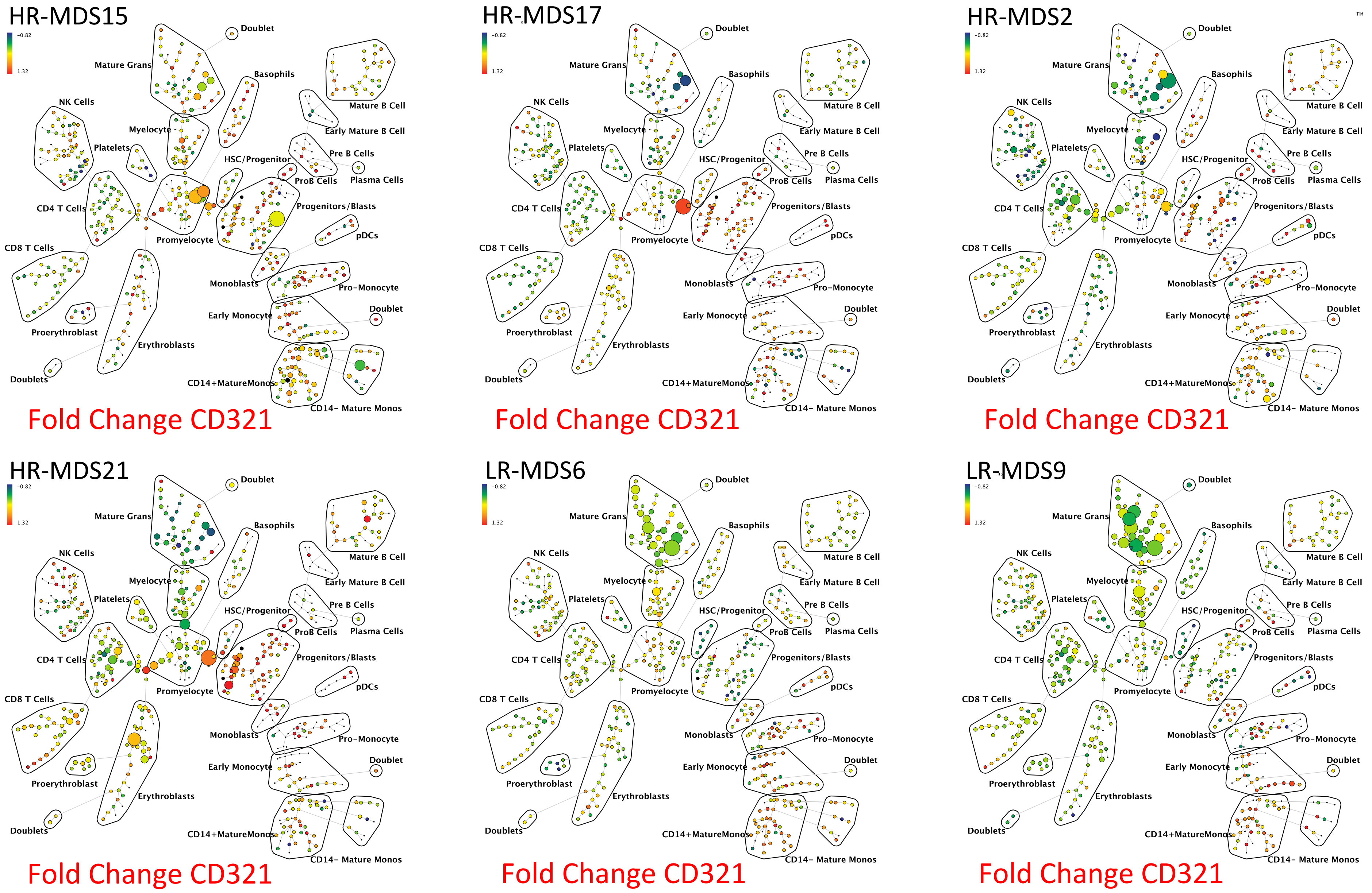

### Supplemental Figure 3C

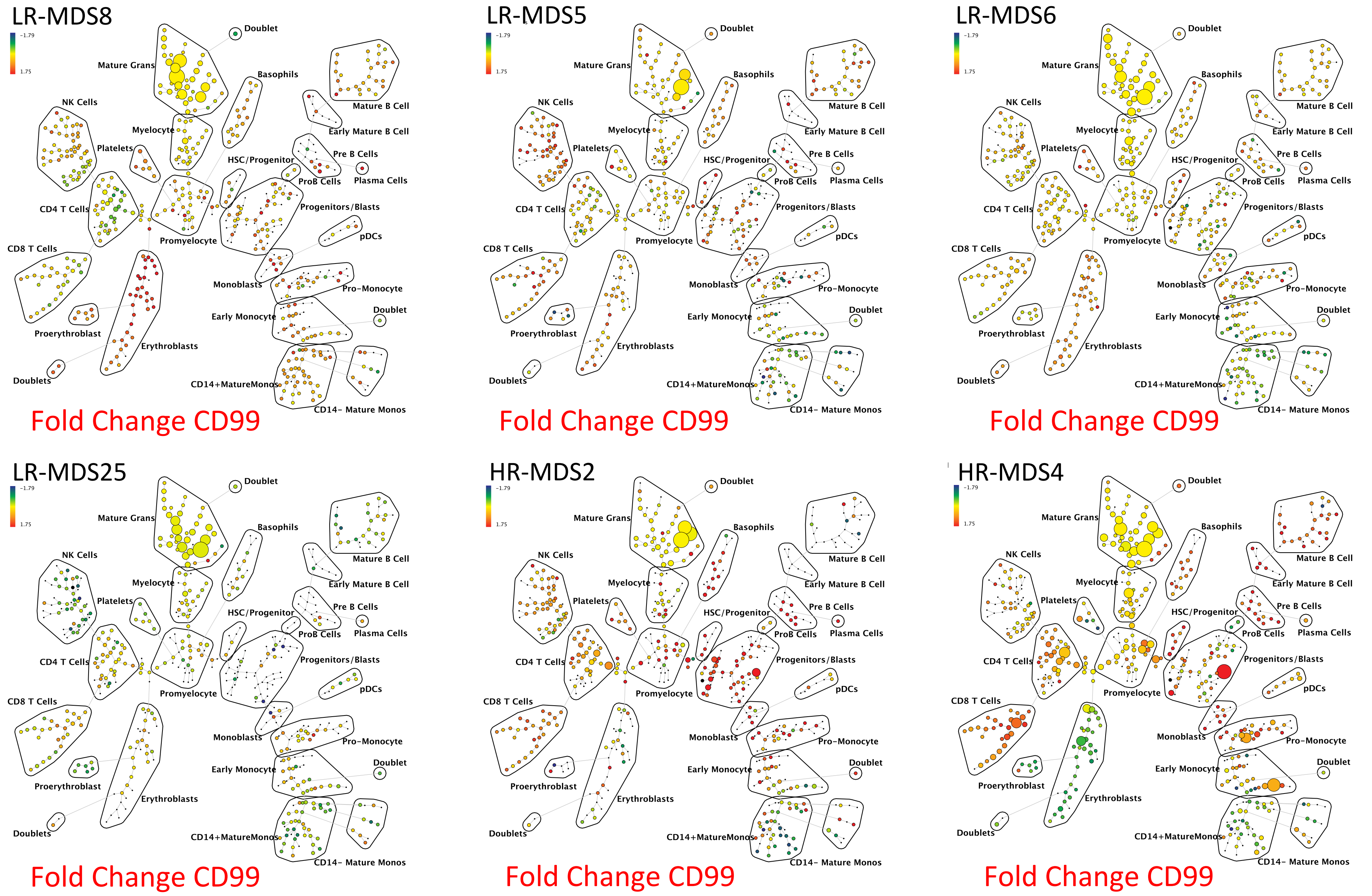

**Supplemental Figure 3**: SPADE analysis enables detection of novel aberrant surface marker expression patterns. SPADE trees colored for the fold change in each cell node for each of the indicated markers CD47 **(A)**, CD321 **(B)**, and CD99 **(C)** relative to the average of the 8 healthy donor samples for the same node. Cell nodes are colored from lowest expression relative to normal (blue) to highest expression relative to normal (red); yellow and light green colors indicate no change in expression relative the average of the control samples. These can be compared to the normal samples shown in Supplemental Figure 4. HR indicates higher-risk MDS (IPSS of Int-2 or sAML/High) LR indicates lower-risk MDS (IPSS of low or Int-1). The 8 replicate control samples came from 5 healthy donors. The size of each node is correlated to the fraction of cells mapping to the node; however, a minimum size was enforced for most nodes to allow visualization of node color. Immunophenotypic grouping of nodes was performed manually on the basis of the median marker expression level of each node, and based on analysis of the relevant biaxial plots (e.g., CD38 vs. CD34).

### Supplemental Figure 4

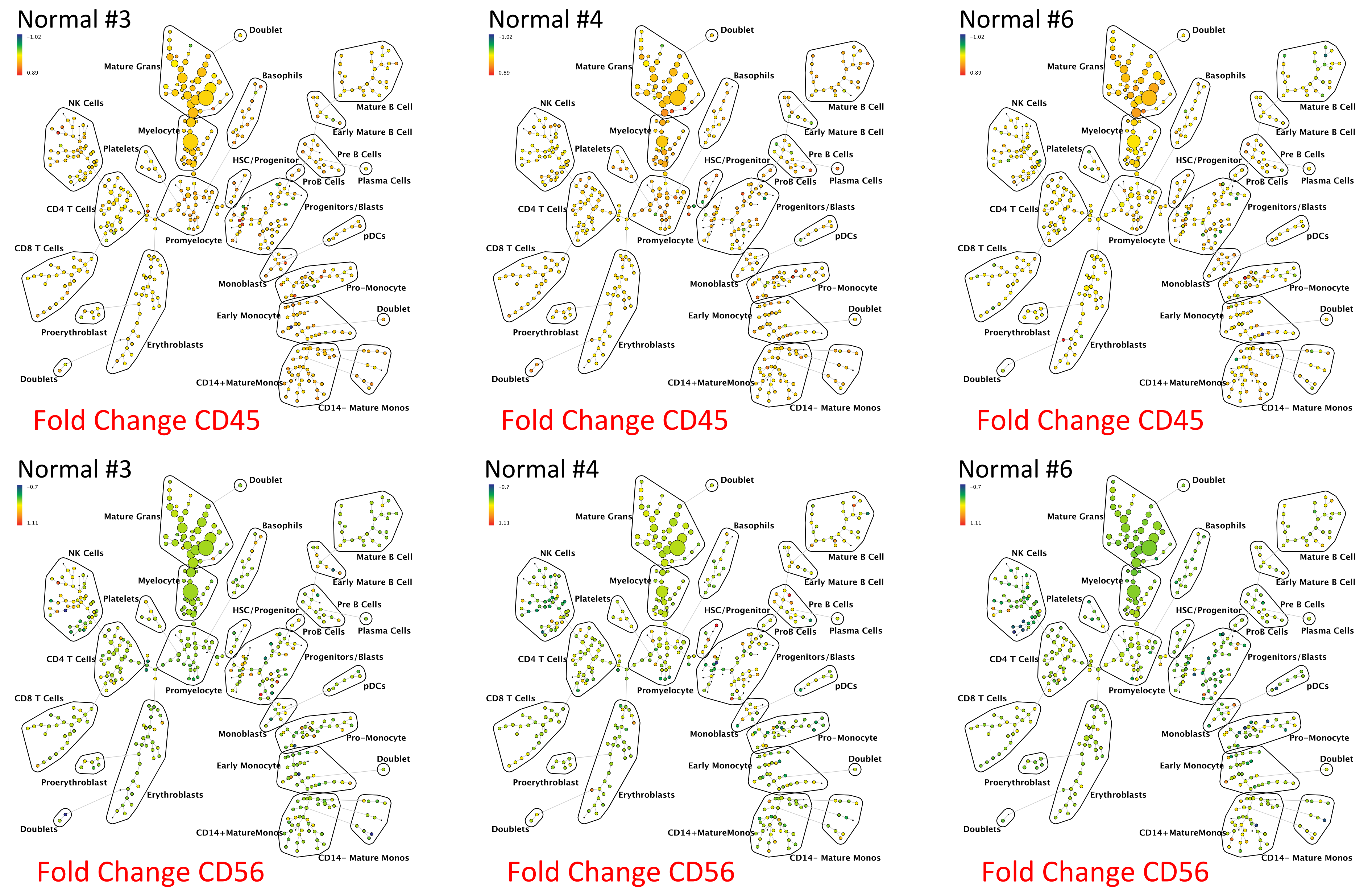

### Supplemental Figure 4

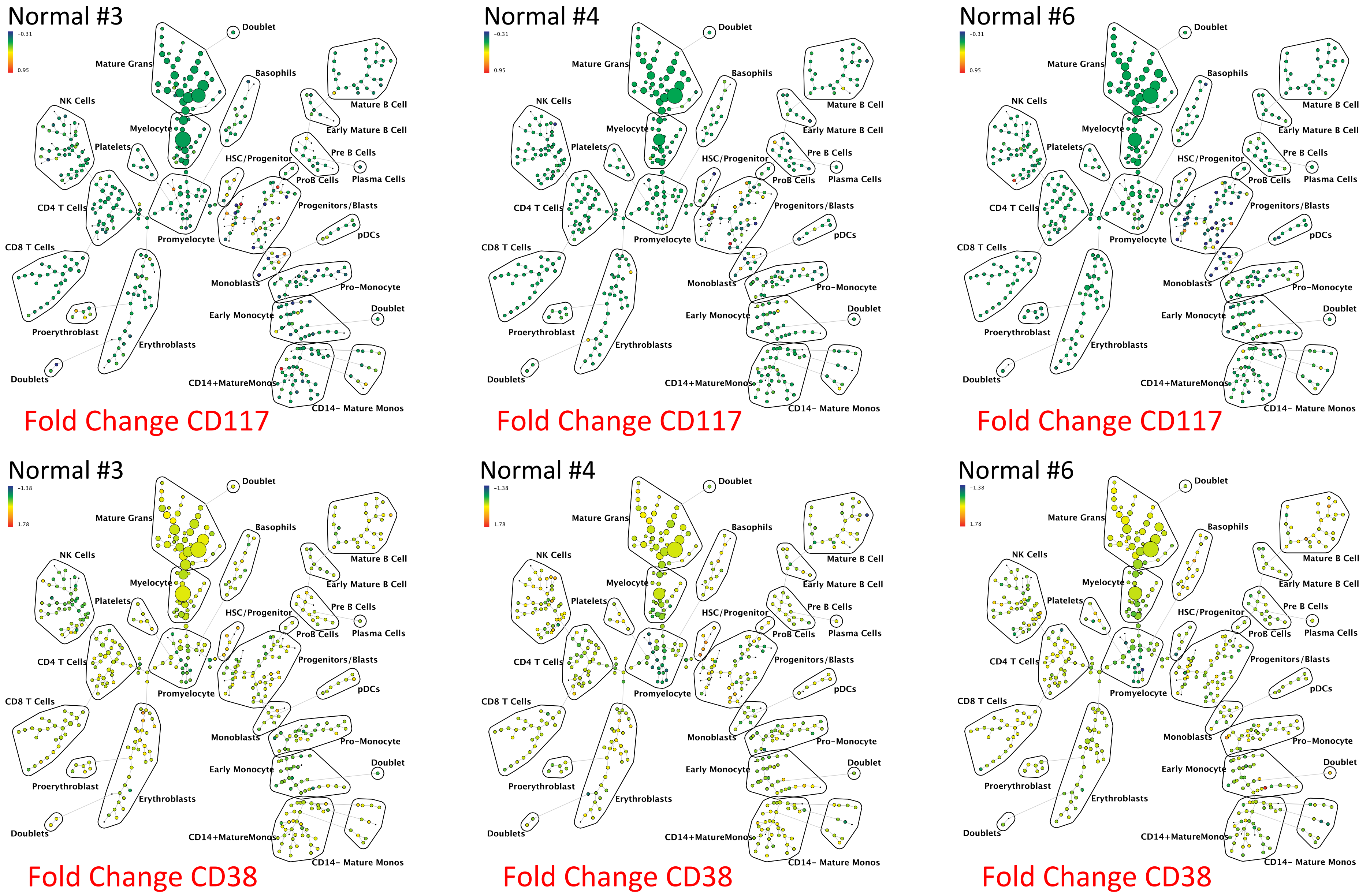

### Supplemental Figure 4

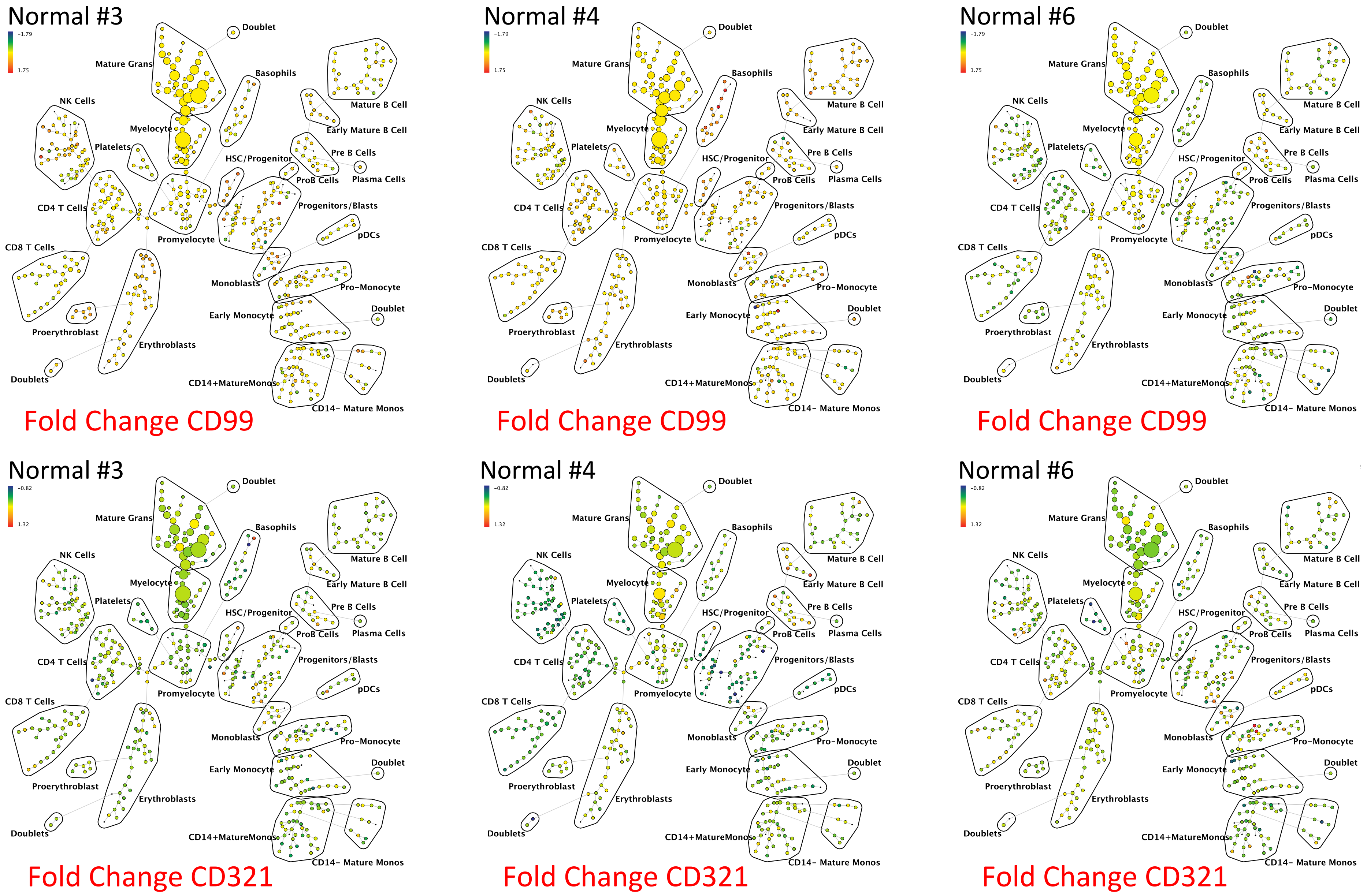

**Supplemental Figure 4**: Normal surface marker expression variance in 3 of the 5 healthy control samples. SPADE trees are colored for the fold change in each node for each of the indicated markers from lowest (blue) to highest (red); each healthy donor control sample node is compared to the average of all 8 control sample aliquots for the same node; yellow represents no change. The healthy control samples demonstrated relatively low levels inter-sample variance (as compared to aberrant expression patterns shown in Figures 1B and Supplemental Figure 3). The size of each node is correlated to the fraction of cells mapping to the node; however, a minimum size was enforced for most nodes to allow visualization of node color.

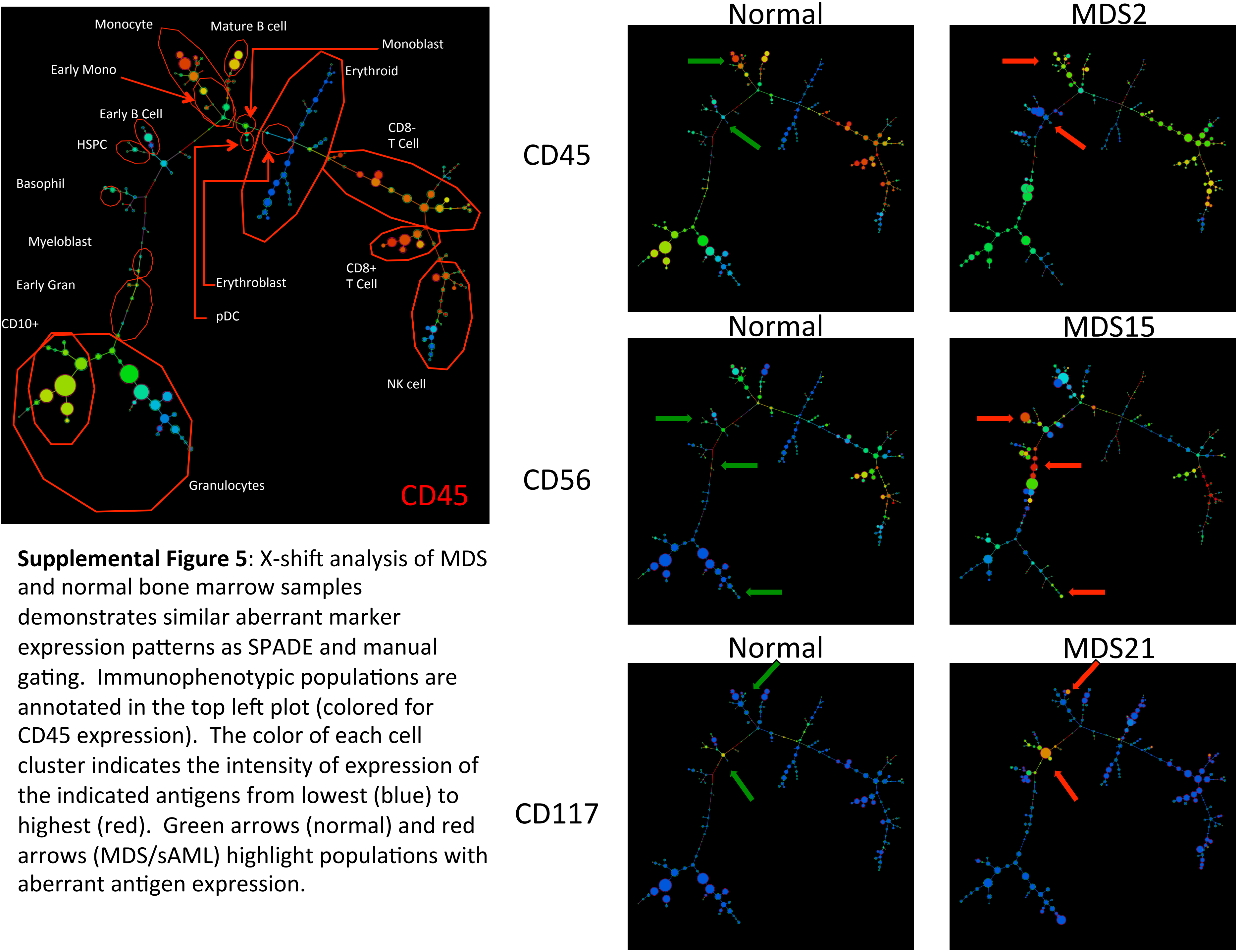


**Supplemental Figure 5**: X-shift analysis of MDS and normal bone marrow samples demonstrates similar aberrant marker expression patterns as SPADE and manual gating. Immunophenotypic populations are annotated in the top left plot (colored for CD45 expression). The color of each cell cluster indicates the intensity of expression of the indicated antigens from lowest (blue) to highest (red). Green arrows (normal) and red arrows (MDS/sAML) highlight populations with aberrant antigen expression.

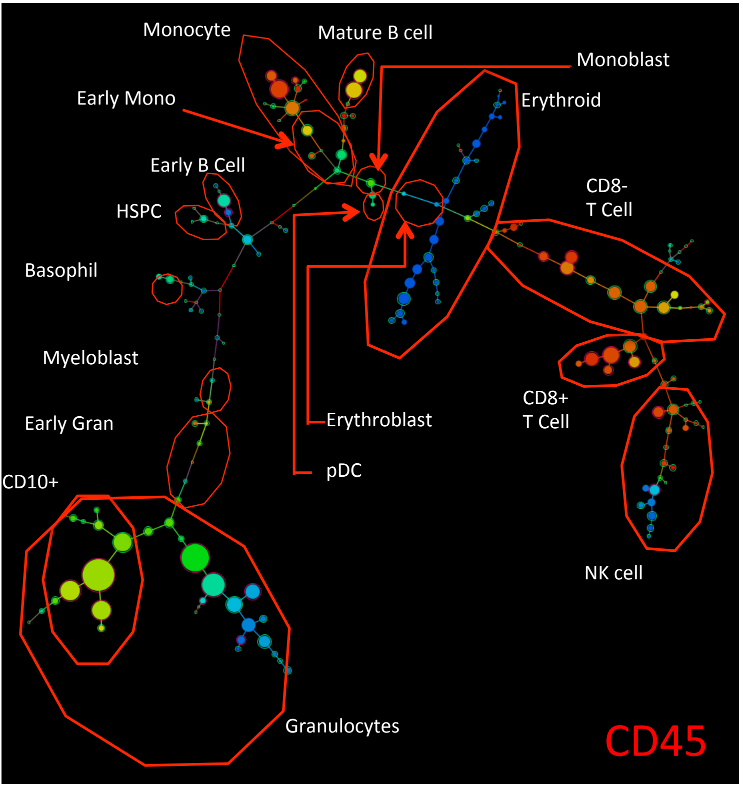


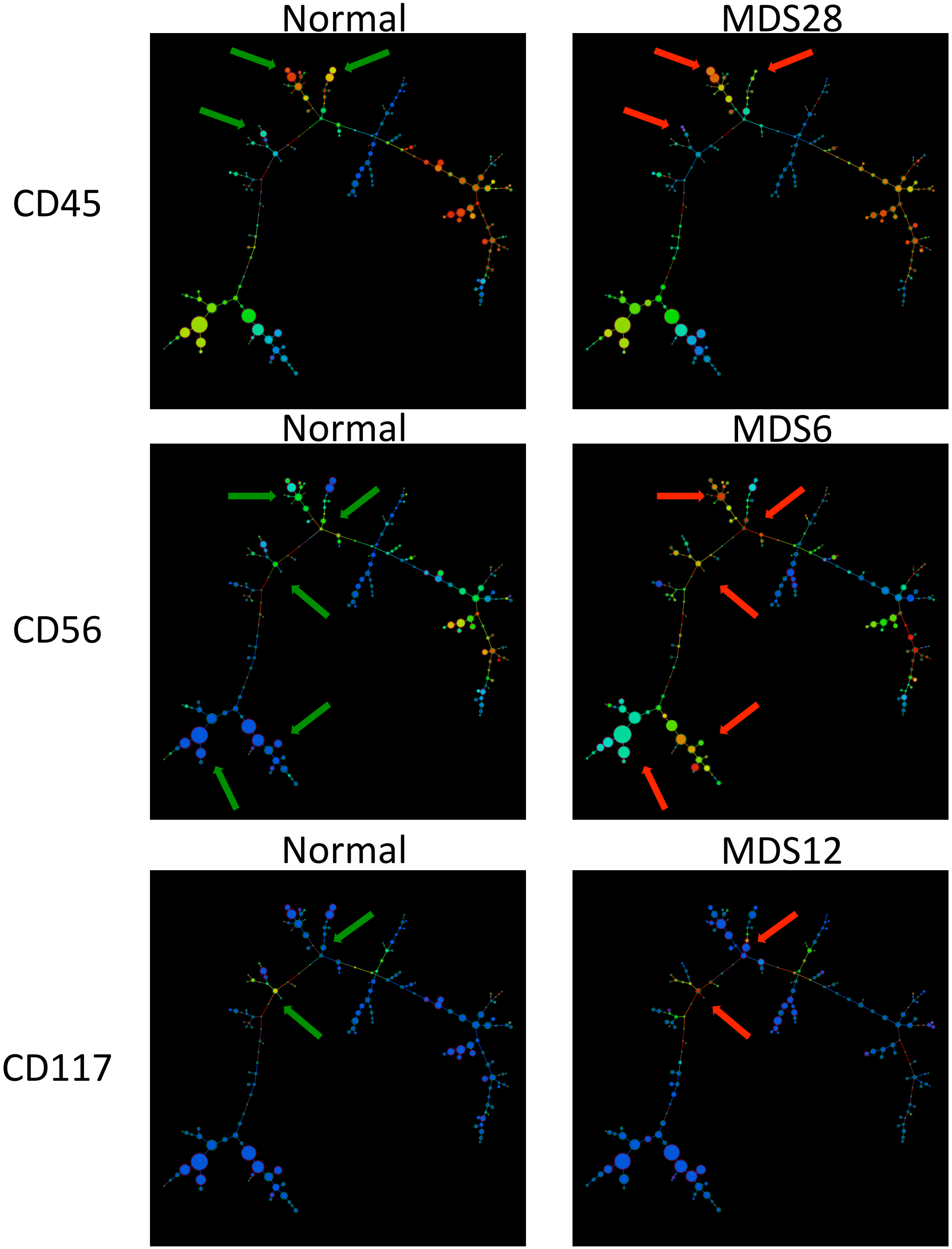


**Supplemental Figure 5**: X-shift analysis of MDS and normal bone marrow samples demonstrates similar aberrant marker expression patterns as SPADE and manual gating. Immunophenotypic populations are annotated in the top left plot (colored for CD45 expression). The color of each cell cluster indicates the intensity of expression of the indicated antigens from lowest (blue) to highest (red). Green arrows (normal) and red arrows (MDS/sAML) highlight populations with aberrant antigen expression.

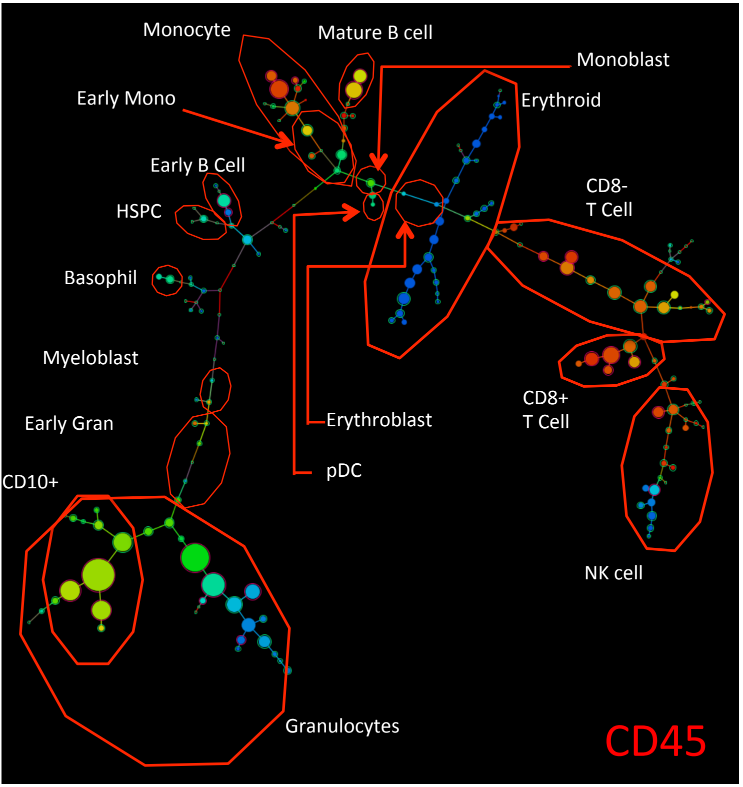


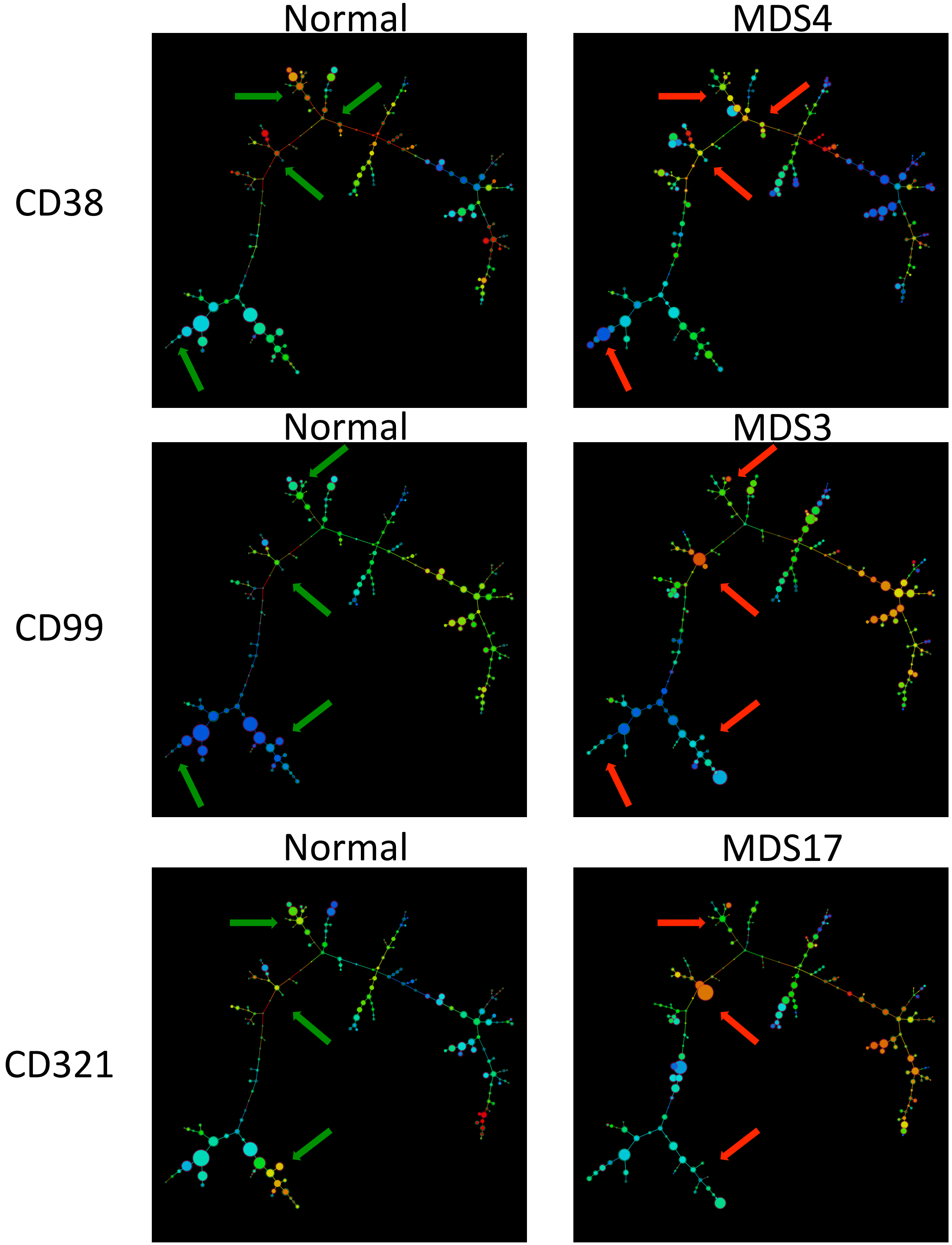


**Supplemental Figure 5**: X-shift analysis of MDS and normal bone marrow samples demonstrates similar aberrant marker expression patterns as SPADE and manual gating. Immunophenotypic populations are annotated in the top left plot (colored for CD45 expression). The color of each cell cluster indicates the intensity of expression of the indicated antigens from lowest (blue) to highest (red). Green arrows (normal) and red arrows (MDS/sAML) highlight populations with aberrant antigen expression.

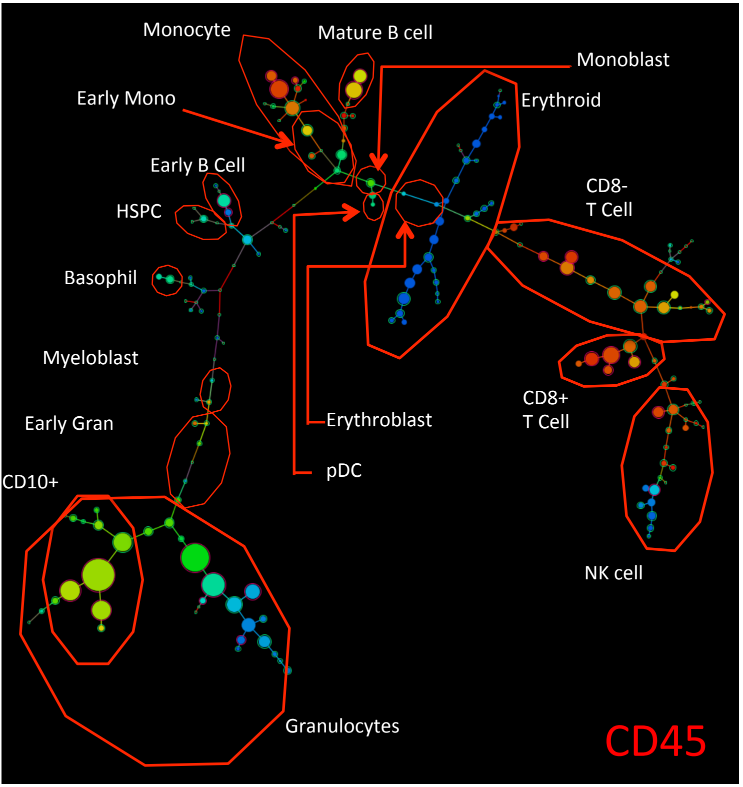


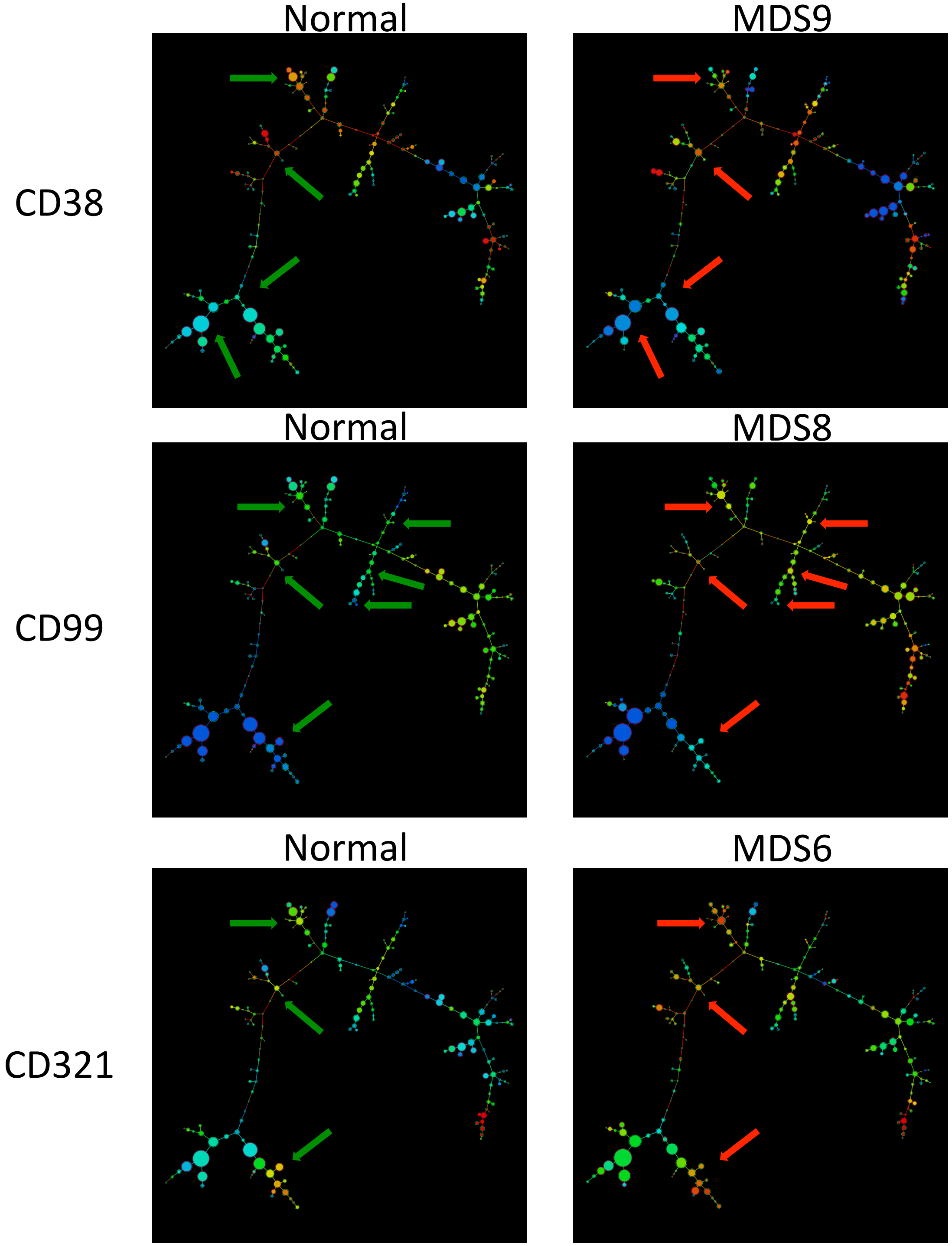


**Supplemental Figure 5**: X-shift analysis of MDS and normal bone marrow samples demonstrates similar aberrant marker expression patterns as SPADE and manual gating. Immunophenotypic populations are annotated in the top left plot (colored for CD45 expression). The color of each cell cluster indicates the intensity of expression of the indicated antigens from lowest (blue) to highest (red). Green arrows (normal) and red arrows (MDS/sAML) highlight populations with aberrant antigen expression.

### Supplemental Figure 6

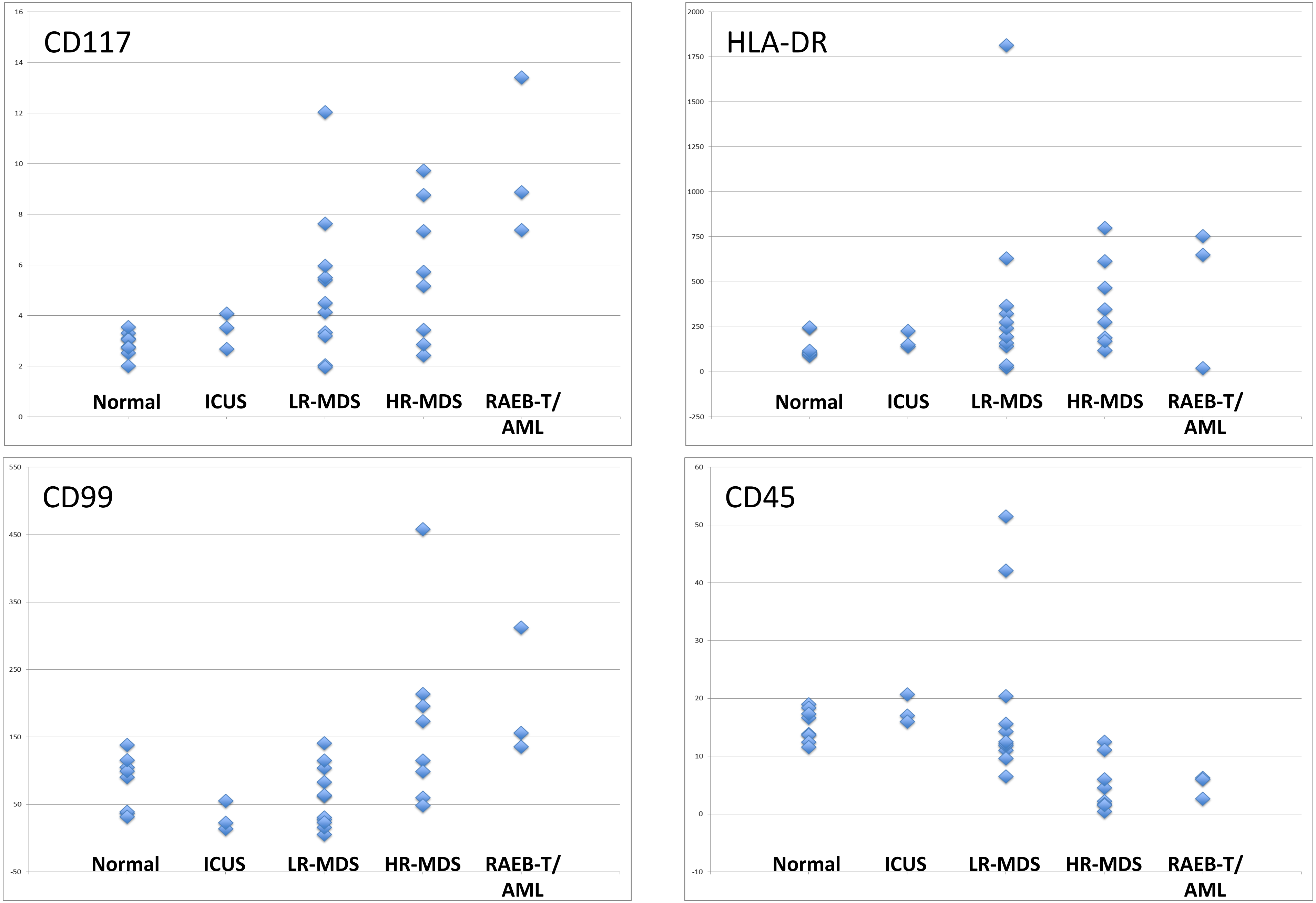

**Supplemental Figure 6:** Specific marker aberrances in $CD34^+CD38^{low}$ cells from bone marrow are correlated with clinical risk in MDS. Median expression level of the indicated markers in the total $CD34^+CD38^{\mathbf{low}}$ population, each data point represents the median expression level of one patient sample or one of the 8 sample aliquots from the five healthy donors. Normal = healthy donor; LR-MDS = lower-risk MDS (IPSS low or Int-1); HR-MDS = higher-risk MDS (IPSS Int-2 or high). Note that the analysis includes samples MDS3, MDS13, and MDS21 (all higher-risk MDS) that come from serial biopsies of the same patient (each several months apart) and demonstrate consistent properties.

Supplemental Figure 7

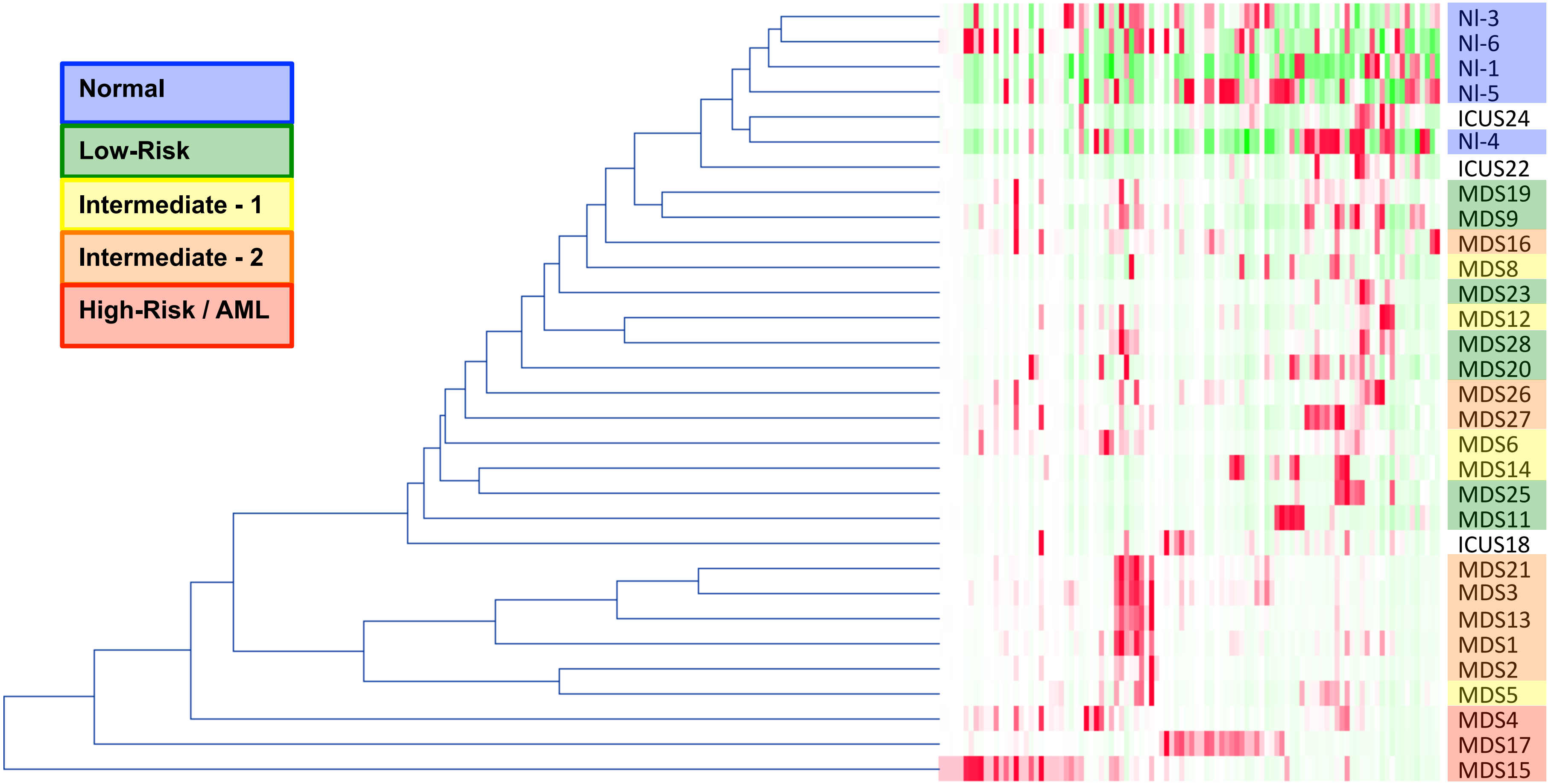

**Supplemental Figure 7**: Multi-dimensional binning analysis of the $CD34^{+}CD38^{low}$ subset confirmed that MDS immunophenotypic patterns are observed in high-dimensional space. Hierarchical grouping of normal and MDS samples based on the pairwise correlation of the distribution of cells across the 100 multi-dimensional bins are shown. Samples from patients with higher IPSS risk clustered further from the normal samples than those from patients with lower IPSS risk. Note that samples MDS3, MDS13, and MDS21 come from serial biopsies of the same patient (each several months apart) and demonstrate consistent properties.

**Supplemental Figure 8**: Summed surface marker aberrancy in the $CD34^{+}CD38^{low}$ cell subset is inversely correlated with survival. Summed fold change in marker expression across the 31 measured surface marker is shown versus patient survival in months (from the time that the analyzed sample was collected). Survival outcomes were reached for 11 patients (13 total samples). Pearson's R= -0.56, $p < 0.05$. Note that samples MDS3, MDS13, and MDS21 come from serial biopsies of the same patient (each several months apart) and demonstrate consistent properties.

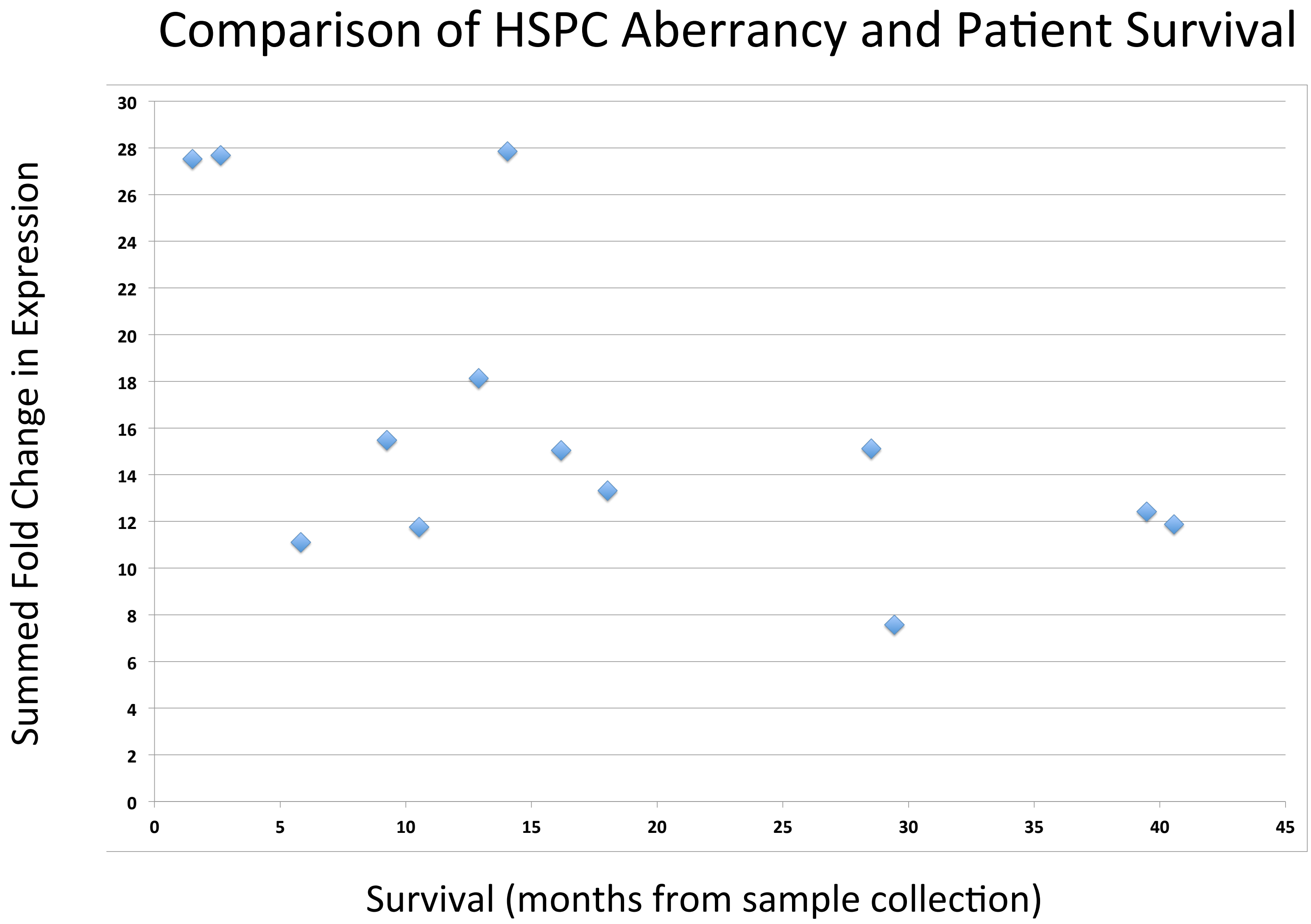

### Supplemental Figure 9

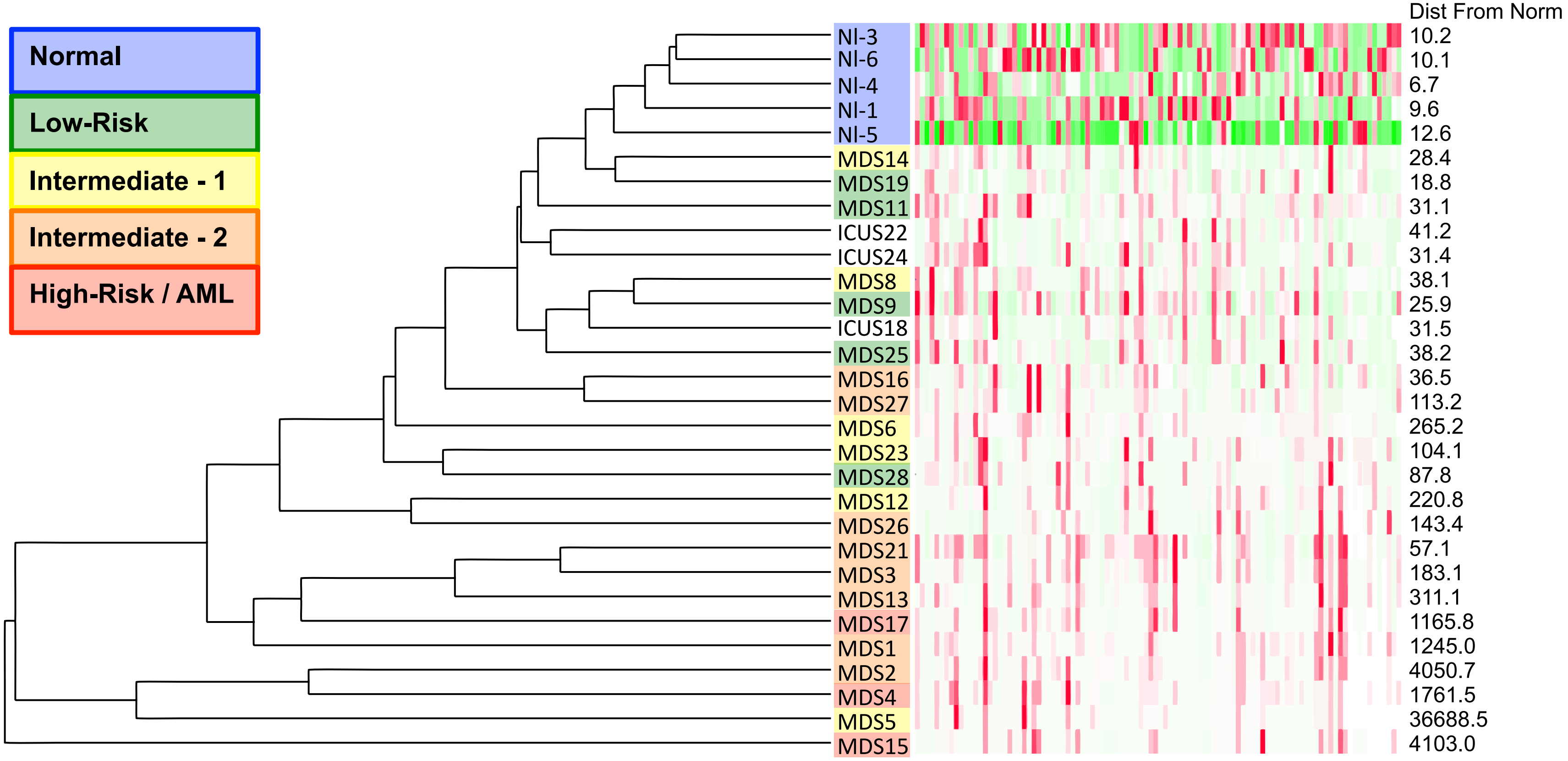

**Supplemental Figure 9**: Multi-dimensional binning analysis of total cell events subset confirmed that distinct MDS immunophenotypic patterns are observed in high-dimensional space. Hierarchical grouping of normal and MDS samples based on the pairwise correlation of the distribution of cells across the 100 multi-dimensional bins are shown. Samples from patients with higher IPSS risk clustered further from the normal samples than those from patients with lower IPSS risk. Note that samples MDS3, MDS13, and MDS21 come from serial biopsies of the same patient (each several months apart) and demonstrate consistent properties.

### Supplemental Figure 10A

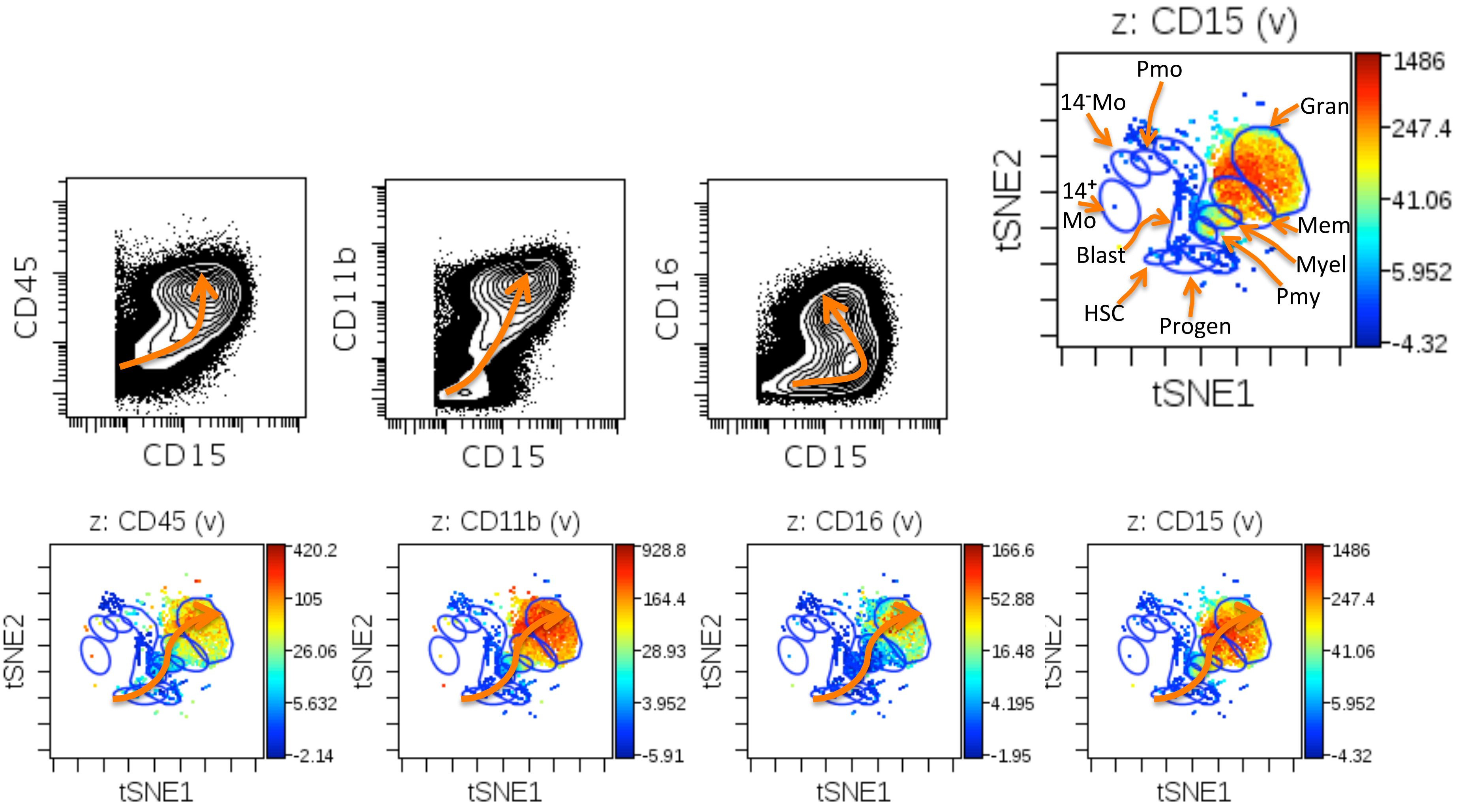

### Supplemental Figure 10B

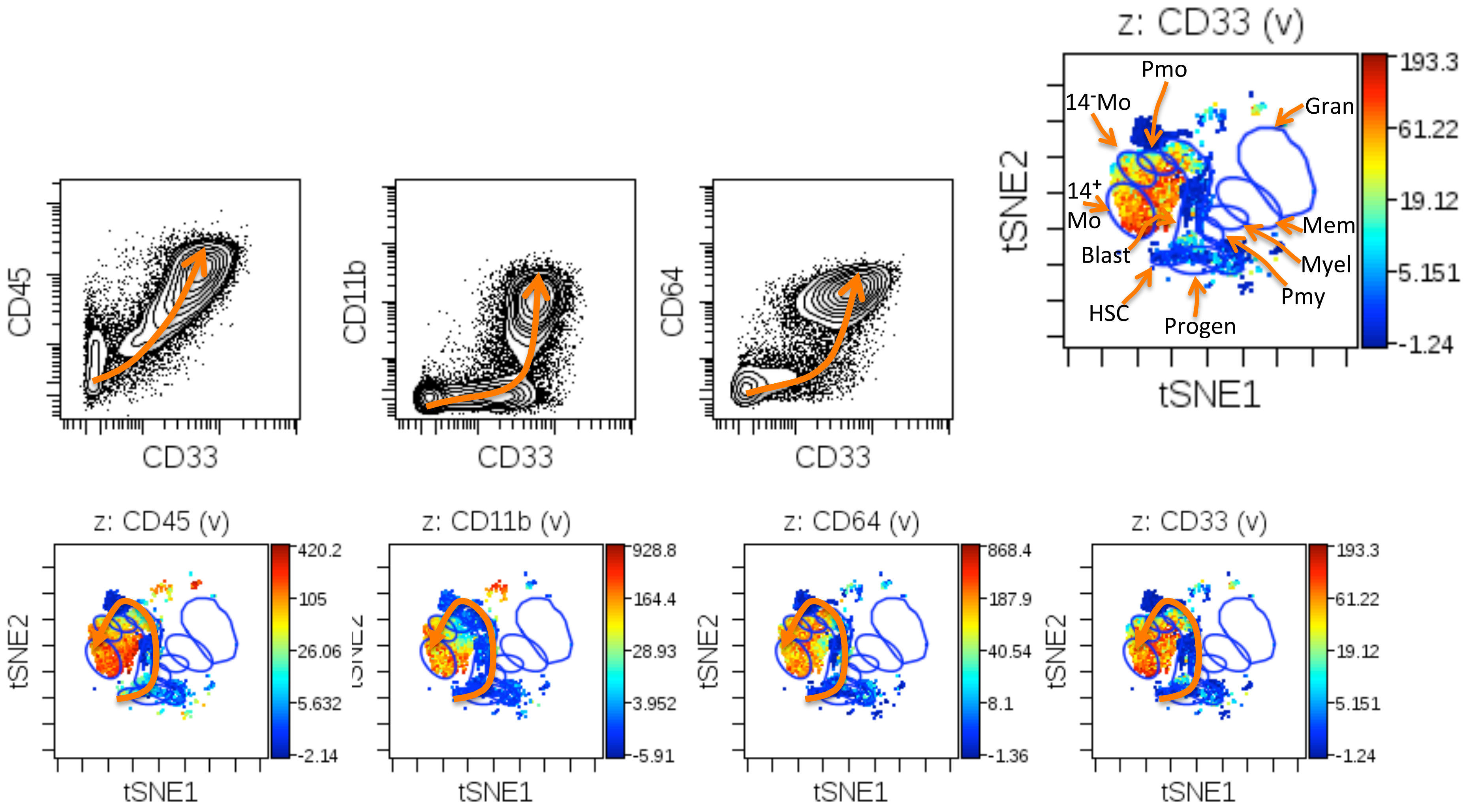

**Supplemental Figure 10**: Normal developmental progression in during granulocyte (A) and monocyte development (B). The normal immunophenotypic progression is shown by the arrow in both biaxial plots and viSNE projections colored for the indicated surface markers. Blue gates on the viSNE projection show the position of the following cell populations: HSC=immunphenotypic hematopoietic stem cells ($CD34^{+}CD38^{lo}lin^{-}$), Progen=early progenitor cells, Blast=Myelo/monoblasts, Pmy=promyelocytes, Myel=myelocytes, Mem=metamyelocytes, Gran=mature granulocytes, Pmo=promonocytes, $14^{-}$Mo=CD14 negative monocytes, $14^{+}$Mo=CD14+ monocytes.

Supplemental Figure 11A

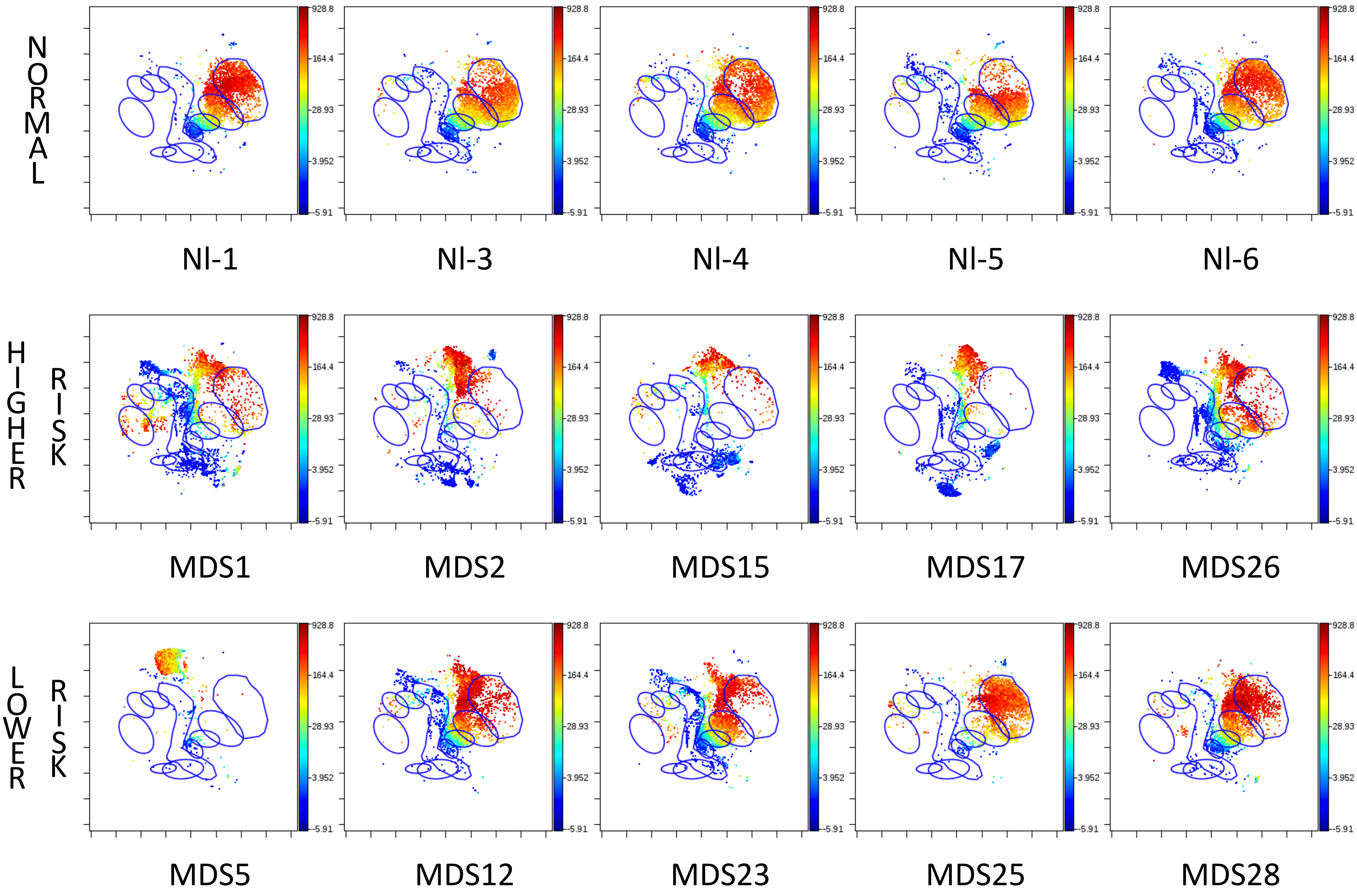

Supplemental Figure 11B

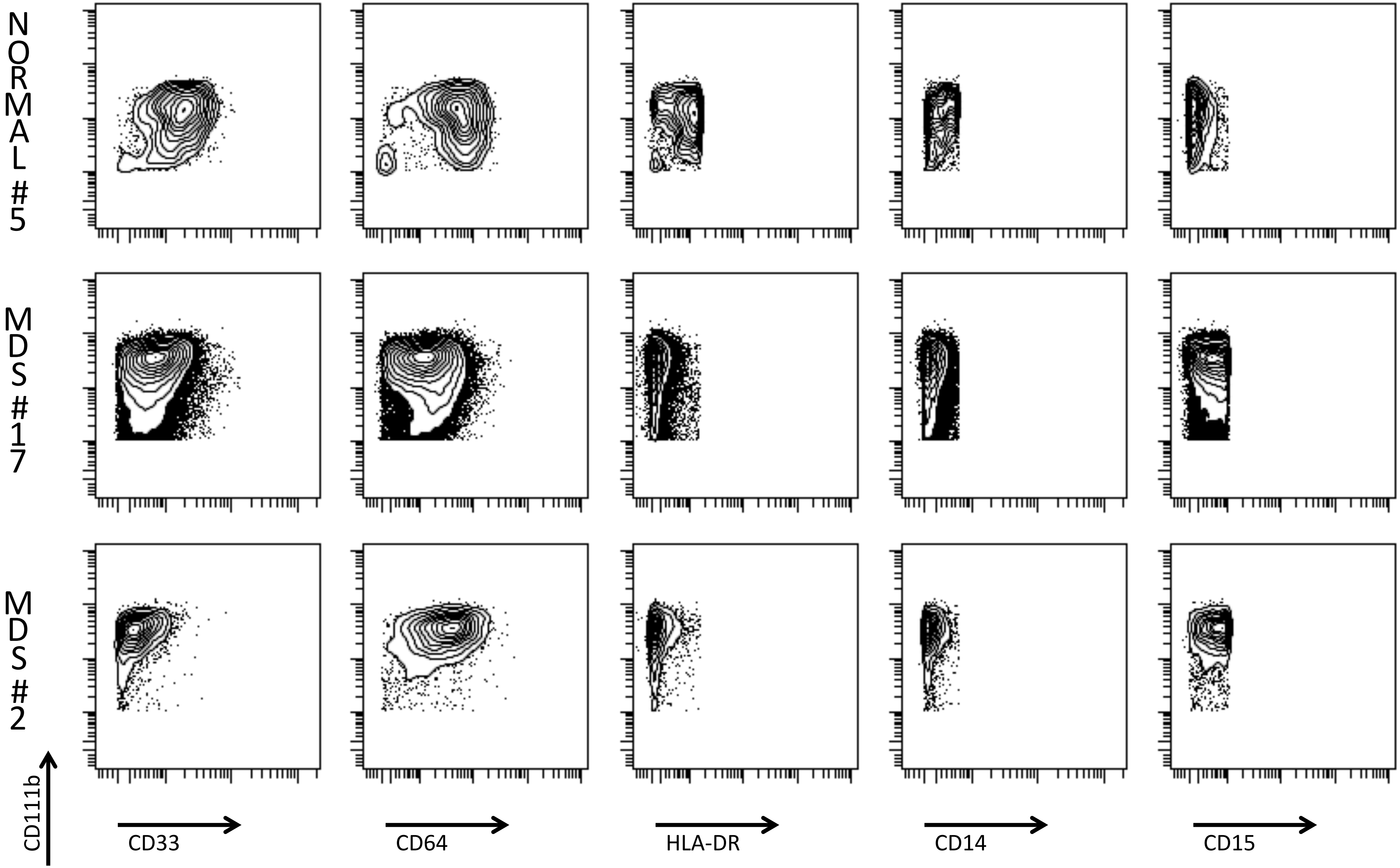

**Supplemental Figure 11**: Immunophenotypically aberrant myeloid cells (IAMCs) are more frequent in patients with higher-risk MDS and sAML. (A) viSNE analysis of granulocyte development (differentiated cells of other lineages have been removed). Blue circles show positions of normal developmental stages (as shown in Supplemental Figure 10). Plots are colored for CD11b expression from low (blue) to high (red). (B) Biaxial plots of the manually gated IAMC cells (see Supplemental Figure 1 for gating). IAMCs from healthy donor sample #5 and MDS samples 2 and 17 are shown.

Supplemental Figure 12

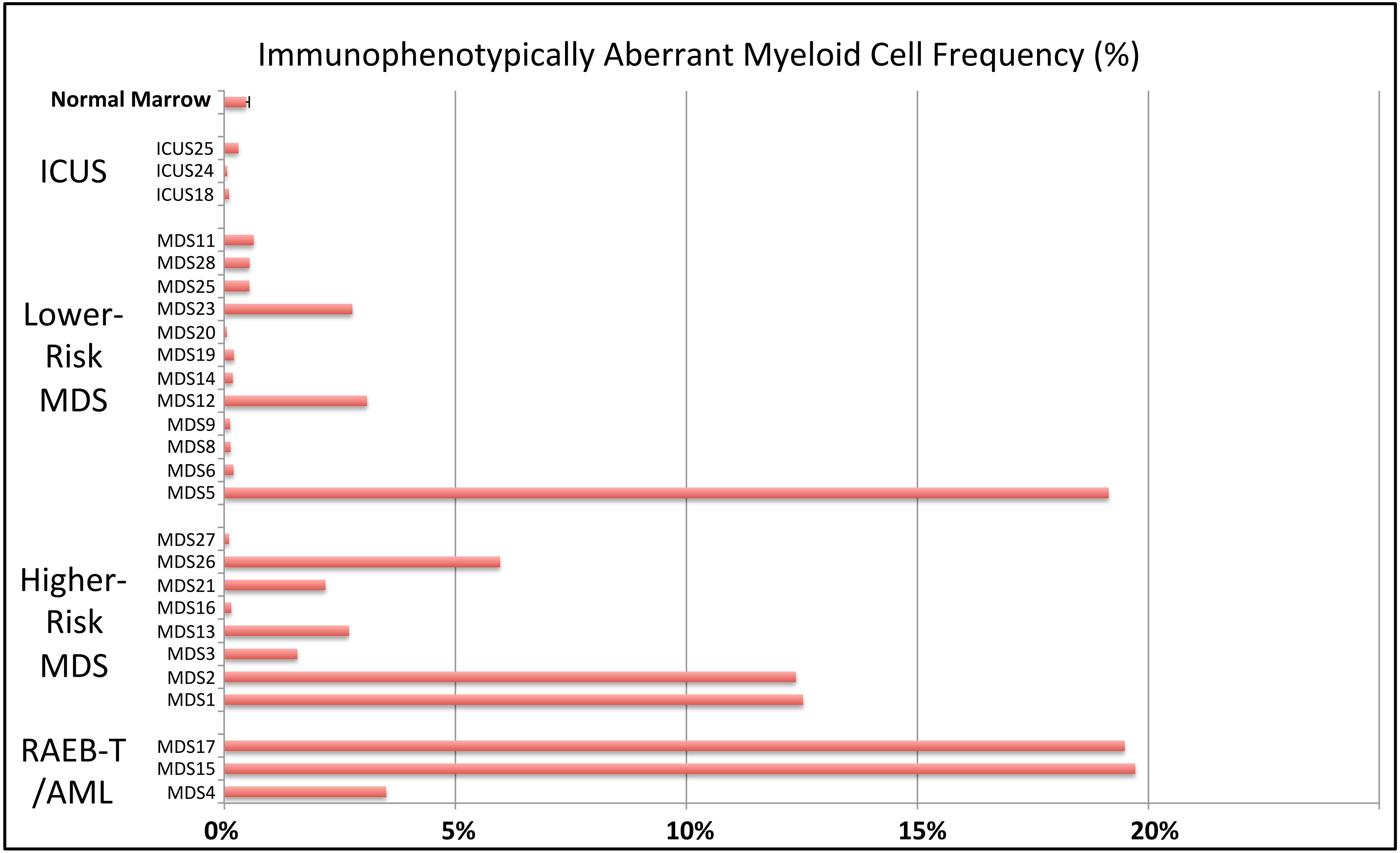


**Supplemental Figure 12**: MDS bone marrow samples demonstrate an increase in immunophenotypically aberrant myeloid cells. The frequency immunophenotypically aberrant myeloid cells (% of total cell events) is shown for the average of all healthy donor samples and for each individual patient sample. Error bar (for normal samples only) indicates the standard error of the mean for the 8 healthy donor samples (derived from 5 healthy donors). Note that samples MDS3, MDS13, and MDS21 come from serial biopsies of the same patient (each several months apart) and demonstrate consistent properties.

**Supplemental Figure 13**: Use of markers for both lineage gating and aberrant marker detection. CD123 was used to both exclude pDCs (green arrows) and basophils (blue arrows) while still allowing for detection of aberrant increases in CD123 among immature progenitor cells (such as $CD34^{+}CD38^{low}$ cells shown on the right).

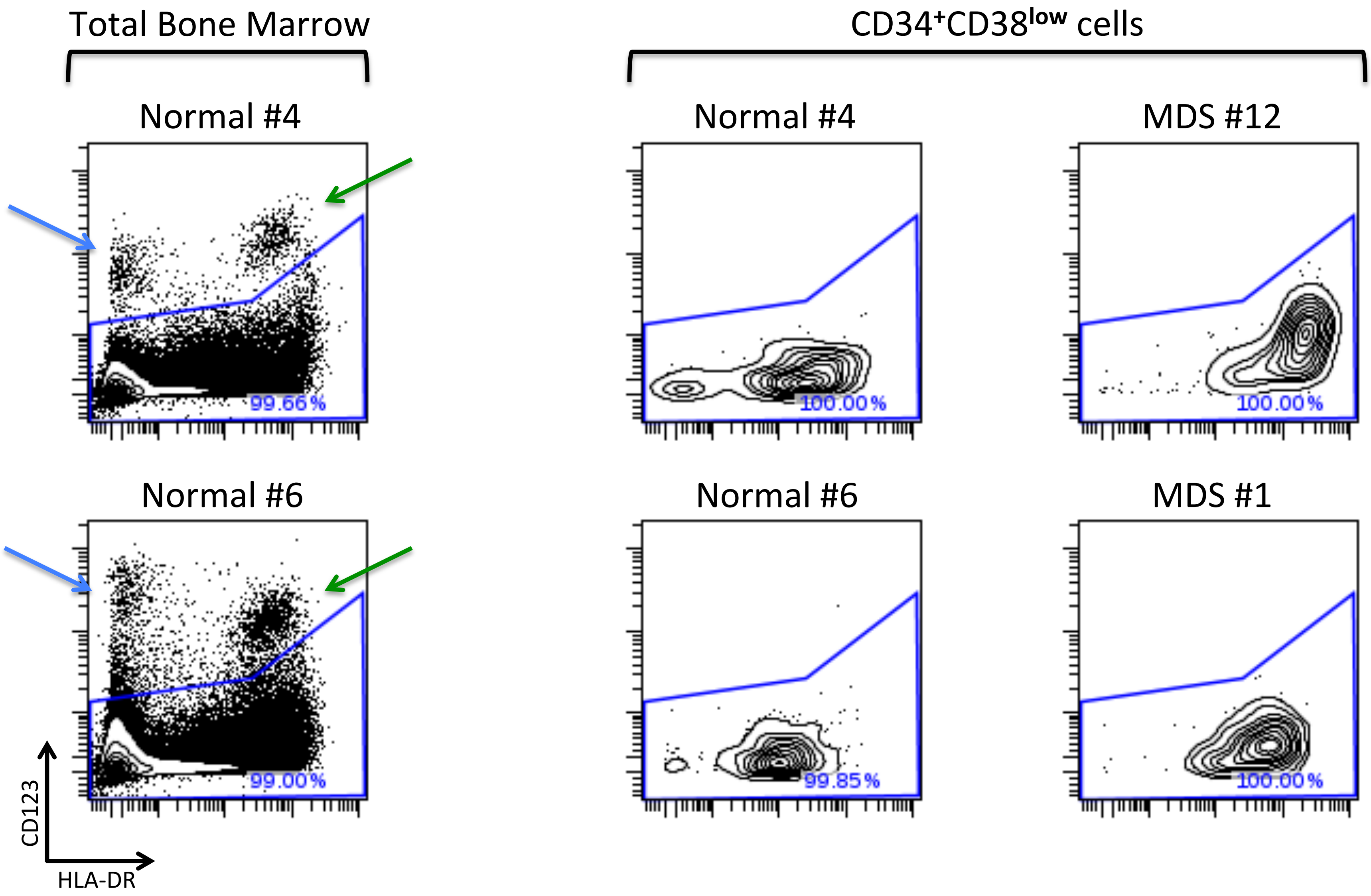
